# Macroevolutionary dynamics of beetles reveal long-term coupling with vascular plant diversification

**DOI:** 10.64898/2026.05.18.725901

**Authors:** Jules Ferreira, David Peris, Corentin Jouault, Fabien L. Condamine

**Affiliations:** Institut Botànic de Barcelona (CSIC-CMCNB), 08038 Barcelona, Spain; Facultat de Ciències de la Terra, Universitat de Barcelona, 08028 Barcelona, Spain; Oxford University Museum of Natural History, University of Oxford, Parks Road, Oxford OX1 3PW, United-Kingdom; CNRS, Institut des Sciences de l’Évolution de Montpellier, Université de Montpellier, 475 rue du Truel, 34090 Montpellier, France

## Abstract

Beetles (Coleoptera) represent the most species-rich group of organisms, yet the macroevolutionary processes underlying their exceptional diversification remain unresolved. Here, we estimated their origination and extinction dynamics as well as the potential drivers shaping these patterns, using Bayesian birth-death models applied to a comprehensive fossil occurrence dataset. We find that beetles have experienced low extinction rates and exhibited high resilience through major extinction events. Vascular plant diversities emerge as a key driver of beetle diversification, with origination rates positively correlated with angiosperms, and extinction rates negatively correlated, especially for Polyphaga, the most diverse beetle clade. Together, our results provide quantitative evidence that the angiosperm radiation not only promoted beetle origination, but also buffered them against extinction, illustrating how ecological interactions can shape macroevolution.

---

The oft-cited remark that “the Creator has an inordinate fondness for beetles” attributed to J.B.S. Haldane, reflects the extraordinary dominance of beetles in global biodiversity. Indeed, with more than 440,000 described species, beetles (Coleoptera) account for about a quarter of all known organisms (Mora et al., 2011; Stork, 2018; Barclay and Bouchard, 2023). They occupy virtually all terrestrial and freshwater ecosystems and exhibit a wide array of ecologies, including herbivory, predation, saprophagy, coprophagy, and fungivory (Beutel et al., 2024). Beetles are also an ancient lineage, with a fossil record extending back to the Permian, 298.9–293.52 million years ago (Ma) (Beutel et al., 2024; Kirejtshuk et al., 2014), and is structured into four extant suborders: Archostemata, Polyphaga, Adephaga, and Myxophaga (Bouchard et al., 2017; Cai et al., 2022). Despite this ecological breadth and deep evolutionary history, the processes underlying their exceptional diversification remain a central unresolved question in macroevolutionary biology (Farrell, 1998; Smith and Marcot, 2015).

Several hypotheses have been proposed to explain the extraordinary diversification of beetles. Intrinsic innovations, such as the evolution of elytra and holometabolous development, may have enabled ecological expansion and niche partitioning (Zhang et al., 2018; Goczał and Beutel, 2023; Goczał et al., 2024). Genomic changes facilitating shifts to new food resources have also been proposed (McKenna et al., 2019; Seppey et al., 2019), alongside high lineage persistence and relatively low extinction rates, which may have promoted the long-term accumulation of diversity (Smith and Marcot, 2015; Hunt et al., 2007) (Fig. 1A, D, E). Another, long-standing alternative/complementary hypothesis links beetle diversification to the rise and ecological dominance of angiosperms during the Cretaceous and Cenozoic (McKenna et al., 2015; Benton et al., 2022; Peris and Condamine, 2024) (Fig. 1A–C). This idea is notably supported by the prevalence of phytophagous lineages, particularly within Polyphaga, which comprises more than 90% of extant beetle species and dominates post-Cretaceous diversity (Beutel et al., 2024; McKenna et al., 2019; Seppey et al., 2019) (Fig. 2B, D). However, whether the expansion of flowering plants directly influenced beetle origination, extinction, and lineage turnover over deep time remains unclear. Disentangling these effects is challenging because of the inherent structure of the fossil record and potential sampling biases, particularly those associated with amber preservation (Smith and Marcot, 2015; Clapham et al., 2016; Schachat et al., 2019).

**Figure 1.**
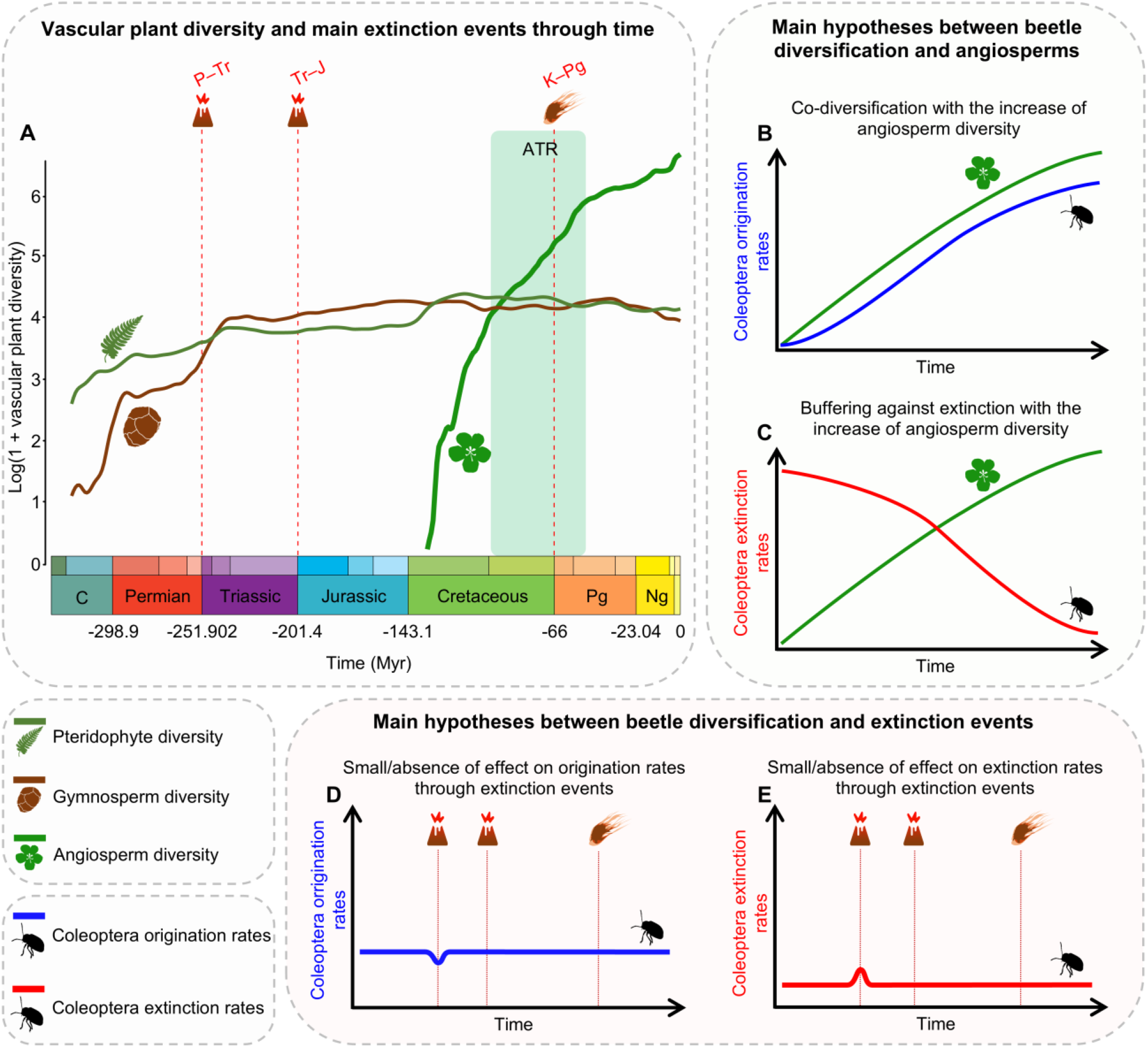
Main hypotheses linking the diversification of Coleoptera with the angiosperm radiation and some major extinction events. (A) Diversity of angiosperms, pteridophytes, and gymnosperms through time, as well as three of the major extinction crises: P–Tr, Permian–Triassic; Tr–J, Triassic–Jurassic; K–Pg, Cretaceous–Paleogene. The main hypotheses include (B) the co-diversification with angiosperms, where beetle origination increases with the radiation of angiosperms, (C) the buffering against extinction, where the radiation of angiosperms reduces beetle extinction by providing more stable and diverse ecological resources, and (D, E) a small, or absence of effect of extinction events on (D) origination rates (i.e., no significant decrease) and (E) extinction rates (i.e., no significant increase). Dark-green area in panel (A) represents the Angiosperm Terrestrial Revolution (ATR). Time is in millions of years (Myr). Abbreviations: C, Carboniferous; Ng, Neogene; Pg, Paleogene. Insect and plant silhouettes are from http://phylopic.org/.

**Figure 2.**
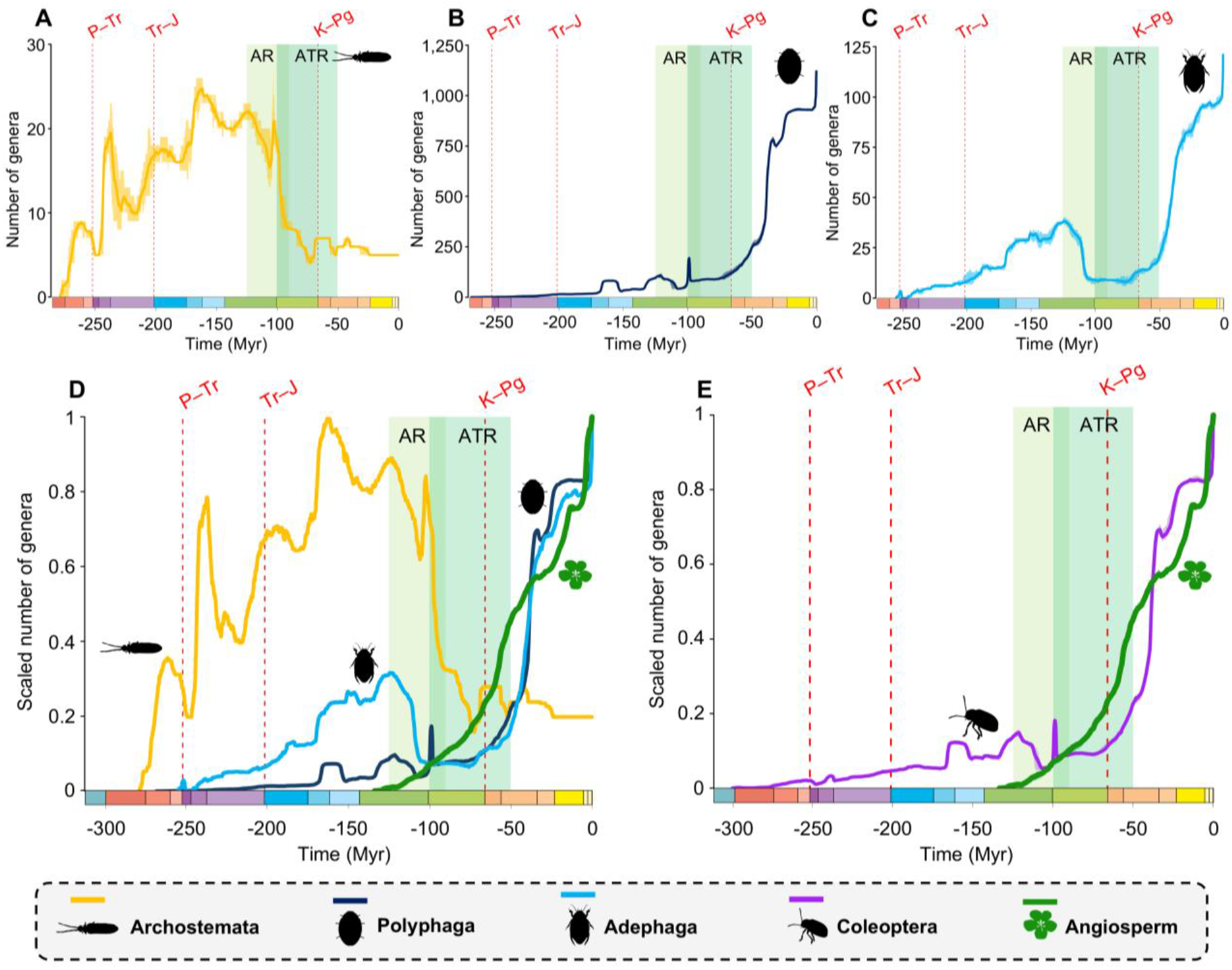
Lineage-through-time plots of Coleoptera, Archostemata, Polyphaga, and Adephaga genera. Number of genera through time computed by summing up the lifespans of all genera for, (A) Archostemata, (B) Polyphaga, and (C) Adephaga. (D) Rescaled number of genera through time for all Coleoptera suborders, overlaid with angiosperm diversity. (E) Rescaled number of genera through time for all Coleoptera genera, overlaid with angiosperm diversity. Solid lines indicate mean diversity at each point in time and shaded areas show estimations of different replications that incorporate age uncertainties of fossil occurrences. Age uncertainties are not shown in panel D as they are shown in panels A–C. Light-green area represents the AR, angiosperm radiation, and dark-green area represents the ATR, angiosperm terrestrial revolution. Red-dashed vertical lines indicate major crises: P–Tr, Permian–Triassic; Tr–J, Triassic–Jurassic; K–Pg, Cretaceous–Paleogene. Time is in millions of years (Myr). Silhouettes are from http://phylopic.org/.

Here, we assembled the most comprehensive genus-level fossil occurrence dataset to infer the macroevolutionary dynamics of Coleoptera across their evolutionary history. This dataset encompasses 32,199 occurrences spanning 300 million years and includes 187 families (out of 240 extant and extinct known families) and 3,398 genera (Data S1–S25). Because beetles originated before the establishment of modern terrestrial ecosystems (*i.e*., the first fossils are found before the expansion of flowering plants), their evolutionary history spans major phases and changes in plant lineages dominance, with the gymnosperm heyday (Triassic–Jurassic), the rise of angiosperms (Cretaceous), the decline of gymnosperms (Late Cretaceous–Cenozoic), and the ecological dominance of angiosperms (Cenozoic).

We therefore tested whether changes in vascular plant diversities influenced beetle diversification dynamics, with the expectation that the radiation of angiosperms may have promoted beetle diversification (McKenna et al., 2015; Benton et al., 2022) by increasing origination rates and/or buffering against extinction (Fig. 1B, C). To do so, we evaluated the effects of plant diversity (pteridophytes, gymnosperms, and angiosperms), alongside environmental variables such as temperature, sea level, atmospheric composition, and continental fragmentation, as well as diversity dependence. We first inferred origination and extinction dynamics over the past 300 million years, capturing temporal variation in diversification rates at both genus and family levels (Silvestro et al., 2014a, 2014b, 2019) (Fig. 2, 3 and figs. S1–S72). We then assessed whether these dynamics could be explained by major biotic and/or abiotic drivers (figs. S73–S76, tables S1–S16, Data S26). Finally, to account for potential preservation biases, we tested the robustness of our results by repeating all analyses after excluding occurrences from amber deposits, which are known to disproportionately preserve certain taxa (Smith and Marcot, 2015; Clapham et al., 2016; Schachat et al., 2019).

**Figure 3.**
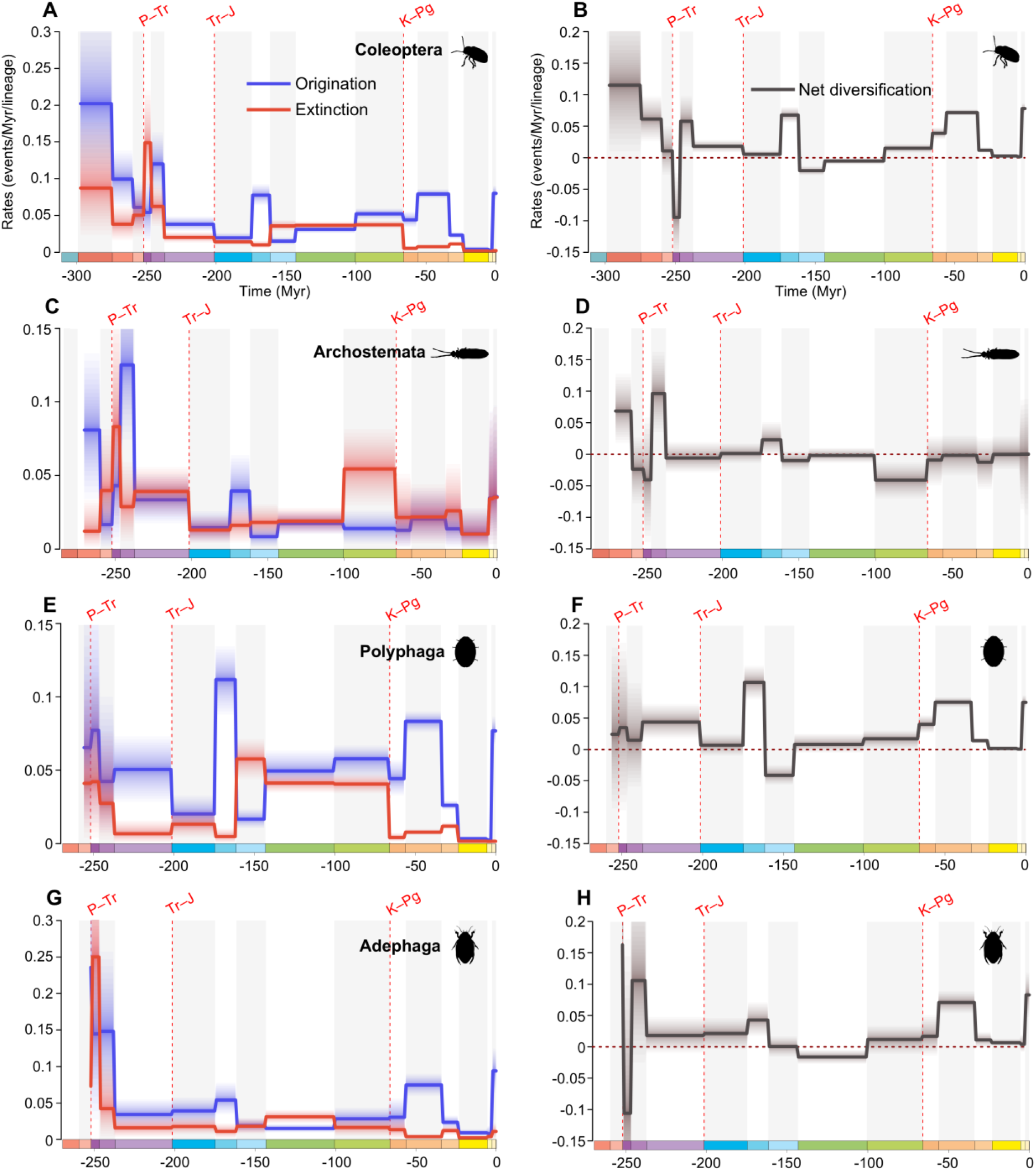
Diversification and diversity dynamics of Coleoptera, Archostemata, Polyphaga, and Adephaga genera. Bayesian fossil-based inferences of Coleoptera (A), Archostemata (D), Polyphaga (G), and Adephaga (J) origination and extinction rates at the genus level under the birth-death model with epochs as constrained shifts (BDCS), without singletons and considering amber occurrences. Net diversification rates for Coleoptera (B), Archostemata (E), Polyphaga (H), and Adephaga (K) obtained from the difference between origination and extinction rates (rates above 0 indicate increasing diversity, and rates below 0 indicate declining diversity). Solid lines indicate mean posterior rates, and the shaded areas show 95% HPD. Red-dashed vertical lines indicate major crises: P–Tr, Permian–Triassic; Tr–J, Triassic–Jurassic; K–Pg, Cretaceous–Paleogene. Time is in millions of years (Myr). The color of each geological period in the chronostratigraphic scale follows that of the International Chronostratigraphic Chart (v2024/12). Silhouettes are from http://phylopic.org/.

## An ancient clade with high resilience through time

Coleoptera may have originated around the uppermost Carboniferous–lowermost Permian, approximately 300 Ma (Fig. 3A, B). Despite this old age, it seems that beetle diversification does not conform to a simple model of continuous expansion or to a single major radiation event (Figs. 2, 3 and figs. S1–S72). Instead, genus-level analyses reveal a long-term trajectory characterized by early establishment, episodic turnover, and a late ecological restructuring that ultimately shaped modern diversity. The earliest positive diversification signal, detected in the Cisuralian (Permian), likely reflects the emergence of stem-lineages (Beutel et al., 2024; Smith and Marcot, 2015). During this interval, net diversification is positive (Fig. 3B), indicating that origination exceeds extinction at this early stage of beetle evolutionary history. However, this early phase does not translate into sustained expansion. Diversification rates decline toward the Guadalupian and Lopingian (Permian, Fig. 3B), despite the origination of Archostemata during the Guadalupian. Early beetles were thus established but not yet flourishing, likely restricted to gymnosperm-associated niches (Ponomarenko, 2003; Feng et al., 2017). This decoupling between early origin and later success is a central feature of beetle macroevolution. It suggests that the extraordinary modern diversity of Coleoptera may not be explained solely by intrinsic innovations such as elytra or holometabolous development, which were already established early (Zhang et al., 2018; Goczał and Beutel, 2023; Goczał et al., 2024). Instead, these traits may have provided latent potential evolutionary potential. In this context, early beetle evolutionary history represents a phase of lineage establishment, whereas ecological dominance emerged later through interactions with changing environments and biotic parameters.

Across their evolutionary history, beetles show an overall low extinction relative to origination. This is especially true during three of the major extinction crises such as the Permian– Triassic (∼252 Ma), the Triassic–Jurassic (∼201 Ma), and the Cretaceous–Paleogene (∼66 Ma) (Figs. 2, 3). Extinction signals remain low or poorly resolved (Fig. 3A, B and fig. S19). During the P–Tr, extinction rates do not exceed background extinction (fig. S17). However, this likely reflects limitations of the fossil record, as fossil occurrences are sparse during this period. That said, we observe the same trend during the Tr–J and the K–Pg, suggesting that mass extinction events weakly affected beetles (fig. S17 and table S17). Nevertheless, significant episodic peaks of extinction are still observed throughout their history. The Ladinian–Carnian transition (Triassic, ∼238–236 Ma) was the most severe, with ∼51.3% of standing genus diversity lost, much later followed by a Late Jurassic (∼154.4–151.6 Ma) turnover that eliminated ∼43.9% of the standing diversity. Nevertheless, the Ladinian–Carnian transition appeared to have had a more limited impact on Coleoptera than on insects as a whole, which experienced a loss of ∼74.8% genera (Jouault et al., 2022). In comparison, Cretaceous extinctions were moderate, with ∼13.6% of lineages lost during the Barremian–Aptian (∼122.4–120.1 Ma) transition and only ∼33.1% in the lowermost-Albian (∼113.6–110.8 Ma). Finally, during the Eocene–Oligocene transition (∼35.5–32.5 Ma), losses were minimal (∼3.4%), highlighting the high resilience of beetle genera during this period. Despite these episodic peaks of extinction, Coleoptera have never experienced a significant collapse or more than a 51% loss of genera, which only occurred early in their history. Importantly, these broad-scale patterns remain stable with and without singleton taxa and amber-preserved occurrences, indicating that the inferred macroevolutionary dynamics are robust to potential sampling and taphonomic biases.

Major mass extinction events appear to have had only a limited impact on beetles, suggesting that other factors, most likely long-term environmental changes, played a more important role in shaping their evolutionary history. In our analyses, extinction rates are consistently associated with a limited set of drivers (gymnosperm and pteridophyte diversity, atmospheric O_2_; Fig. 4A, B and tables S1–S2), whereas origination rates are influenced by a broader range of drivers (angiosperm, gymnosperm, and pteridophyte diversity, atmospheric O_2_ and CO_2_). This asymmetry suggests that beetle diversification has been driven more strongly by processes promoting origination than by those increasing extinction, further highlighting the resilience of the group in the face of repeated environmental changes.

**Figure 4.**
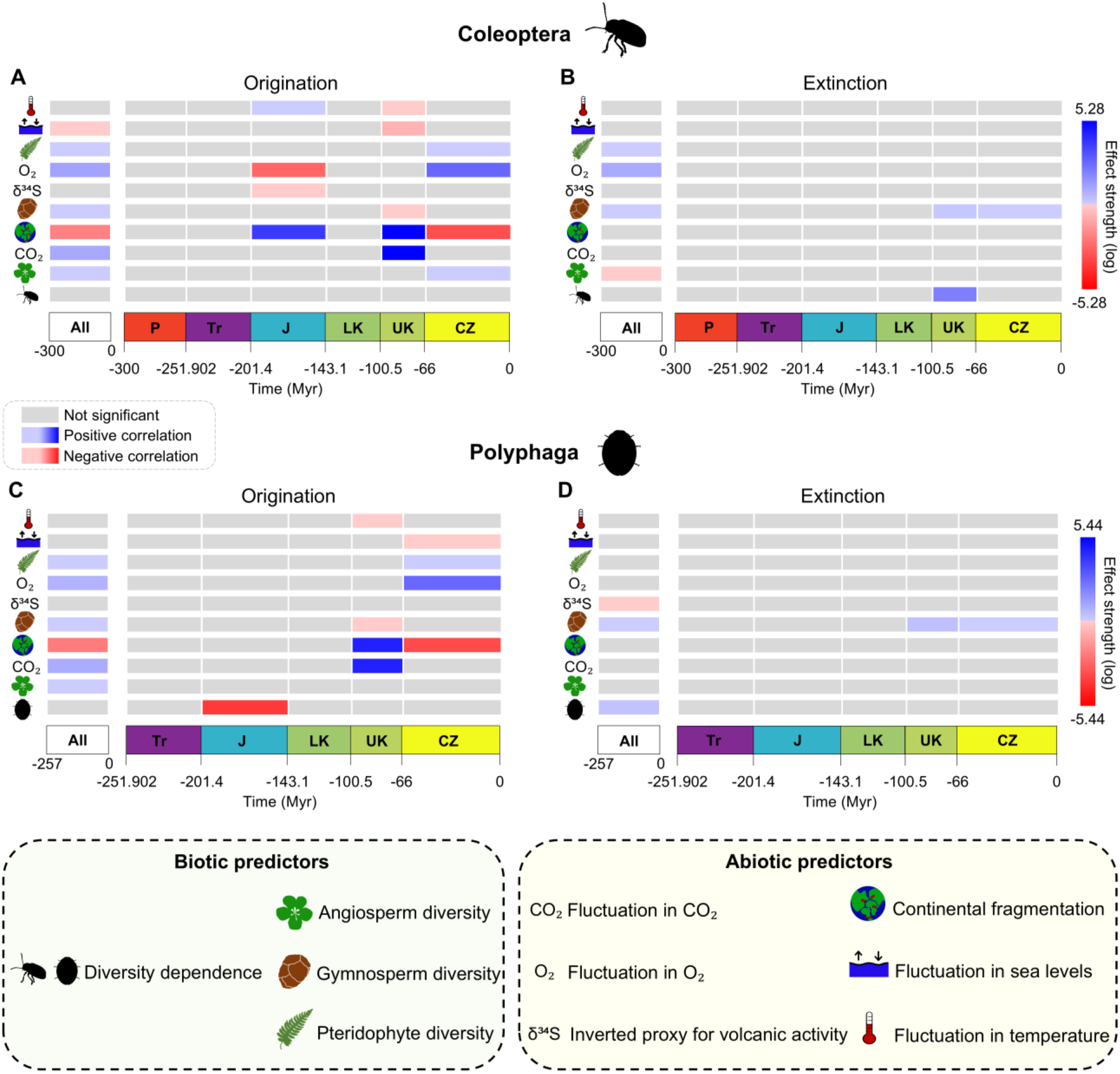
Bayesian estimates of correlations between origination and extinction rates and biotic and abiotic predictors inferred under the Multivariate Birth-Death (MBD) model for Coleoptera and Polyphaga. MBD results for Coleoptera (A, B), and Polyphaga (C, D), for origination (A, C) and extinction (B, D). Results presented are those significant in frameworks with and without singletons. Data includes amber occurrences. Analyses were performed across multiple temporal windows. The strength effect corresponds to the median value of the correlation parameter of the origination/extinction rate. A variable was considered to have a significant effect when its shrinkage weight exceeded 0.5 and when the 95% HPD interval of the corresponding correlation parameter did not overlap with zero. When a significant correlation is positive, it is represented by a filled blue square; when a significant correlation is negative, it is represented by a filled red square; according to the sign of the median value of the correlation parameter. When a correlation is considered insignificant, it is represented by a light-grey filled square. “All” corresponds to the time window encompassing the entire evolutionary history of the clade. The other time intervals are defined as follows: P, Permian (298.9–251.902 Ma); Tr, Triassic (251.902–201.4 Ma); J, Jurassic (201.4–143.1 Ma); LK, Lower Cretaceous (143.1–100.5 Ma); UK, Upper Cretaceous (100.5–66 Ma); CZ, Cenozoic (66 Ma to the present). Time is in millions of years (Myr). Silhouettes are from http://phylopic.org/.

## Multifactorial drivers shaped early beetle diversification

Rather than reflecting a steady accumulation of diversity, genus-level diversification analyses reveal multiple turnover phases in beetle evolutionary history (Fig. 3A, B). The first example is the drop in diversification across the P–Tr boundary, where net diversification becomes strongly negative during the Early Triassic (Fig. 3B). However, this signal is not support by statistically significant extinction peak, likely due to sparse fossil sampling in the latest Permian and earliest Triassic (figs. S17, S19). The absence of a strong statistical signal should therefore be interpreted cautiously, as a limitation of the fossil record rather than evidence of resilience (Jouault et al., 2022). Net diversification rates recovered during the Middle Triassic are positive again (Fig. 3B), driven by increased origination and reduced extinction (Fig. 3A). This signal is particularly strong in Archostemata and Adephaga, suggesting that these groups played a major role in recovery, with Polyphaga contributing more significantly during the Late Triassic (Fig. 3C–H). This rebound is consistent with post-extinction recovery dynamics, in which ecological niches vacated during the crisis are progressively reoccupied (Zhao et al., 2021), although this interpretation remains tentative for the P–Tr. A similar turnover phase is observed during the Jurassic. Genus-level diversification increased markedly during the Middle Jurassic (Fig. 3B), before declining to negative values in the Late Jurassic (Fig. 3B). Importantly, this decline occurred despite continued accumulation of diversity at the family level (Fig. S1), illustrating how extinction-driven turnover can occur without affecting higher-level diversity. These alternating phases of extinction and compensatory origination, as may be the opposite situation, suggest that global diversification is better understood as a dynamic equilibrium rather than a simple continuous accumulation process. In this context, beetle diversification cannot be reduced to a straightforward narrative of progressive radiation.

The drivers of these diversification dynamics are multifactorial, with both abiotic and biotic processes leaving detectable traces in the fossil record from the Jurassic onwards (Fig. 4 and tables S1–S4). During the Jurassic, genus-level origination rates are positively correlated with continental fragmentation and temperature, and negatively with O_2_ and δ^34^S (Fig. 4A, B). The Jurassic was a period of major continental breakup following the fragmentation of Pangaea, which increased provinciality and further habitat heterogeneity (Scotese, 2016, 2021; Zaffos et al., 2017). Such conditions are conducive to speciation through geographic isolation processes. At the same time, fluctuations in atmospheric composition and magmatic activity (captured by O_2_ and δ^34^S proxies) may have imposed physiological or ecological constraints, limiting origination (Fig. 4A, B). Notably, magmatic activity was lower during the Middle Jurassic than during the Early and Late Jurassic, potentially facilitating the observed peak in origination rates during this interval (Rampino et al., 2024). Our results suggest that tectonic processes may have promoted diversification by increasing geographic isolation and ecological opportunity, while other environmental factors may have constrained diversification. Together, these patterns show that early beetle diversification was not only shaped by external environmental changes but also reflects a strong capacity to adapt to shifting ecological conditions. Superimposed on these external drivers is a signal of diversity dependence in Polyphaga. The negative correlation between origination rates and Polyphaga genus diversity during the Jurassic (Fig. 4C, D and tables S3, S4) suggests that as diversity increased, ecological saturation or competition may have limited the establishment of new lineages. This is consistent with diversity-dependent diversification hypotheses in insects, in which speciation slows as niches become filled (Peris and Condamine, 2024).

## Plant turnover restructured beetle diversification in the late Mesozoic

One of the most pronounced shifts in beetle diversification occurred during the Cretaceous– Cenozoic interval, but not as a simple increase in net diversification. Instead, our results point to a major ecological restructuring driven by plant turnover, particularly the rise of angiosperms, and the decline of gymnosperms. During the Early Cretaceous, genus-level diversification remained low (Fig. 3B), despite a marked increase in family-level diversity (Fig. S1). This decoupling suggests that, while higher-level lineages were diversifying, genus-level dynamics were contracting. Notably, this period coincides with the early phase of the angiosperm radiation (125– 90 Ma; Labandeira, 2014). Clearer evidence for plant-driven dynamics emerges during the Late Cretaceous. Genus-level net diversification became positive again (Fig. 3B), with origination negatively correlated with gymnosperm diversity and positively correlated with continental fragmentation (Fig. 4A, B and tables S1, S2). At the same time, extinction rates are positively correlated with gymnosperms and beetle diversity, indicating that lineages associated with declining plant groups were probably progressively lost. Similar patterns are observed within Polyphaga (Fig. 4C, D), reinforcing the idea that vegetation turnover played a central role in shaping beetle evolution, mainly driven by this suborder. Comparable dynamics have been proposed for other insect groups, such as Hemiptera (Boderau et al., 2025) and ants (Jouault et al., 2024).

Interestingly, no significant positive correlation is detected between angiosperm diversity and beetle origination during the Early or Late Cretaceous. This likely reflects the fact that angiosperms, although diversifying, had not yet reached ecological dominance during this interval, which was achieved only in the Cenozoic. Accordingly, significant correlations with angiosperm diversity are detected either across the entire evolutionary history of Coleoptera, or specifically during the Cenozoic, corresponding to the phase of angiosperm dominance (Fig. 4A, B and tables S1, S2). During the Angiosperm Terrestrial Revolution (100–50 Ma), flowering plants transformed terrestrial ecosystems, particularly by progressively replacing gymnosperm-dominated landscapes, creating new ecological niches while simultaneously eliminating others (Benton et al., 2022; Coiro et al., 2019; Condamine et al., 2020; Ding et al., 2025). These results are consistent with a temporal lag between the rise of angiosperms and their detectable impact on beetle diversification. This transition likely produced both winners and losers among beetle lineages. Archostemata, for example, may represent one of the declining group, as indicated by negative diversification rates during the Late Cretaceous (Fig. 3D), driven by elevated extinction (Fig. 3C), consistent with a lineage putatively associated with gymnosperm habitats (Ponomarenko, 2003; Feng et al., 2017). However, this interpretation remains tentative due to the limited fossil record for the group. In contrast, Polyphaga show sustained positive diversification at the genus level (Figs. 2B, D, 3F), highlighting their capacity to exploit emerging ecological opportunities. Overall, these results support the view that beetle diversification during the late Mesozoic was driven by ecological turnover rather than net expansion of diversity.

## Persistence of plant-beetle interactions in the Cenozoic

During the Cenozoic, the association between beetle diversification and plant diversity became even more pronounced, with origination rates showing a positive correlation with angiosperm diversity (Fig. 4A–D and tables S1–S4). Although family-level diversification rates stabilized near zero (Fig. S1), genus-level rates remained positive during the Paleogene, peaking in the Eocene before declining in the Oligocene (Fig. 3B). This peak corresponds to a period of warm global climates, followed by a transition toward cooler conditions and the onset of Antarctic glaciation (Meckler et al., 2022). In addition to angiosperms, beetle diversification also correlates with other plant groups and environmental factors. Origination rates are positively correlated with pteridophytes diversity, and a negatively correlated with continental fragmentation, while extinction rates continue to correlate positively with gymnosperm diversity (Fig. 4A–D and tables S1–S4). The Cenozoic was marked by tectonic reorganization, which altered biogeographic connectivity and habitat structure (Scotese, 2016). Within this environmental context, plant communities followed contrasting trajectories. Angiosperms experienced sustained radiation following the K–Pg extinction (∼66 Ma), with increasing speciation rates toward the present contributing to their ecological dominance (Dimitrov et al., 2023). This expansion closely parallels the continued positive diversification in beetles, supporting a persistent ecological linkage between the two groups (Fig. 2A, B). In contrast, gymnosperms continued their long-term decline, marked by reduced diversification and episodes of elevated extinction during the Oligocene and Miocene, likely linked to climatic cooling and competitive displacement by angiosperms (Condamine et al.,2020; Crisp and Cook, 2011). Together, our results suggest that the ecological restructuring initiated during the Cretaceous and consolidated during the Cenozoic continued to shape beetle diversification over extended timescales. This supports the hypothesis, previously proposed but not formally tested using phylogenetic approaches, that angiosperm radiation acted as a long-term driver of beetle evolution (*e.g*., Zhang et al., 2018). Thus, the influence of the angiosperm radiation appears not as a single, discrete event, but as a long-term ecological driver operating over tens of millions of years. The persistence of gymnosperm-associated extinction signals into the Cenozoic further supports the view of gradual lineage replacements rather than one abrupt turnover (Fig. 4A, C and tables S1–S4).

## Angiosperm diversity as a long-term driver of beetle diversification

Across the entire evolutionary history of Coleoptera, origination rates show consistent positive correlations with angiosperms, gymnosperms, and pteridophytes diversity, among others (Fig. 4A, B and tables S1, S2). In contrast, extinction rates are positively correlated with gymnosperms and pteridophytes, and negatively correlated with angiosperms (Fig. 4A, B and tables S1, S2). Importantly, the positive correlations between origination and angiosperms, gymnosperms, as well as pteridophytes, remain significant when amber occurrences are excluded (fig. S75 and tables S9, S10), indicating that these correlations are robust to potential taphonomic biases linked to amber deposits (Smith and Marcot, 2015; Clapham et al., 2016; Schachat et al., 2019). This results differ from earlier studies that did not detect such an associations for Coleoptera, and rather proposed that other factors may be responsible for their high diversification (Hunt et al., 2007; Smith and Marcot, 2015). However, correlations involving extinction rates are no longer recovered after excluding amber data, suggesting that these signals may be more sensitive to sampling biases. Biologically, these results support the hypothesis that beetle diversification has been tightly linked to the expansion of land plants, which likely provided a sequential arrival of new plant groups and increasingly diverse ecological opportunities through time, ultimately promoting lineage origination (McKenna et al., 2015; Zhang et al., 2018). The role of angiosperms is particularly noteworthy: their diversity fostered origination rates and inhibited extinction rates. This suggests that angiosperm-dominated ecosystems may have both promoted beetle diversification and buffered beetles against extinction, a dynamic also reported in other insect groups, and particularly pollinators (Peris and Condamine, 2024; Jouault et al., 2024).

Within Polyphaga (∼90% of extant beetle species), origination rates are positively correlated with angiosperms, gymnosperms, and pteridophytes (Fig. 4C, D and tables S3, S4). These correlations remain largely unchanged when amber occurrences are excluded, highlighting a strong and persistent link between polyphagan diversification and the evolution of vascular plants (fig. S75C, D and tables S11, S12). In contrast, extinction in Polyphaga is positively correlated with gymnosperm diversity and their own diversity (Fig. 4C, D and tables S3, S4). Taken together, these results suggest that Polyphaga diversification has been influenced by vascular plant evolution. The positive correlation between angiosperms and origination, coupled with the positive correlation between gymnosperms and extinction, support the hypothesis that the progressive expansion and ecological diversification of flowering plants acted as a major driver of lineage accumulation in Polyphaga. This is consistent with repeated host shifts from gymnosperms to angiosperms and the exploitation of newly available ecological niches (Benton et al., 2022; Peris and Condamine, 2024; Seppey et al., 2019; Zuntini et al., 2024). In parallel, the diversity dependent effect on extinction may indicate ecological saturation and/or increased intra-clade competition (Rabosky, 2013). As diversity accumulated, the availability of ecological niches may have become limiting, and/or competitive interactions may have intensified, particularly between lineages associated with different plant groups (*e.g*., angiosperms and gymnosperms).

In contrast, when non-Polyphaga beetles are analyzed all together, no significant correlations between diversification and plant diversity are detected across most of their evolutionary history. The only exception occurred during the Cenozoic, when origination is positively correlated with gymnosperms (fig. S74C, D and tables S7, S8). However, this correlation is lost when amber occurrences are excluded. Moreover, no consistent correlation with angiosperms is recovered in these analyses, regardless of the inclusions of amber occurrences (figs. S74C, D, S74C, D and tables S7, S8, S15, S16). This contrast between Polyphaga and other beetle lineages highlights strong clade-specific heterogeneity in macroevolutionary responses. It suggests that the impact of angiosperm diversification on Coleoptera was not uniform, but instead largely concentrated within Polyphaga, reflecting uneven ecological and evolutionary responses among and across beetle suborders.

## Evolutionary perspectives and key findings

Overall, our findings show that the radiation of Coleoptera began ∼300 Ma, well before the emergence of angiosperm-dominated ecosystems. Beetle diversification has been strongly influenced by complex interactions between biotic and abiotic factors through time. In addition to their overall low extinction rates and remarkable resilience to major extinction events, vascular plant diversity appears as one of the most consistent drivers of beetle diversification. In contrast, abiotic variables such as atmospheric composition, sea level fluctuations, continental fragmentation, and temperature exert more variable and time-dependent effects. The strength and consistency of the correlations between plant diversity and beetle diversification, together with its robustness across analyses accounting for fossil sampling biases, indicate that the expansion and dominance of angiosperms and the concurrent decline of gymnosperms played a key role in shaping the macroevolutionary history of the group. These results emphasize the importance of long-term ecological interactions, particularly plant-beetle associations, in driving the exceptional diversity, resilience, and adaptative versatility of Coleoptera.

