## Supplementary Figures and Tables for "Macroevolutionary dynamics of beetles reveal long-term coupling with vascular plant diversification"

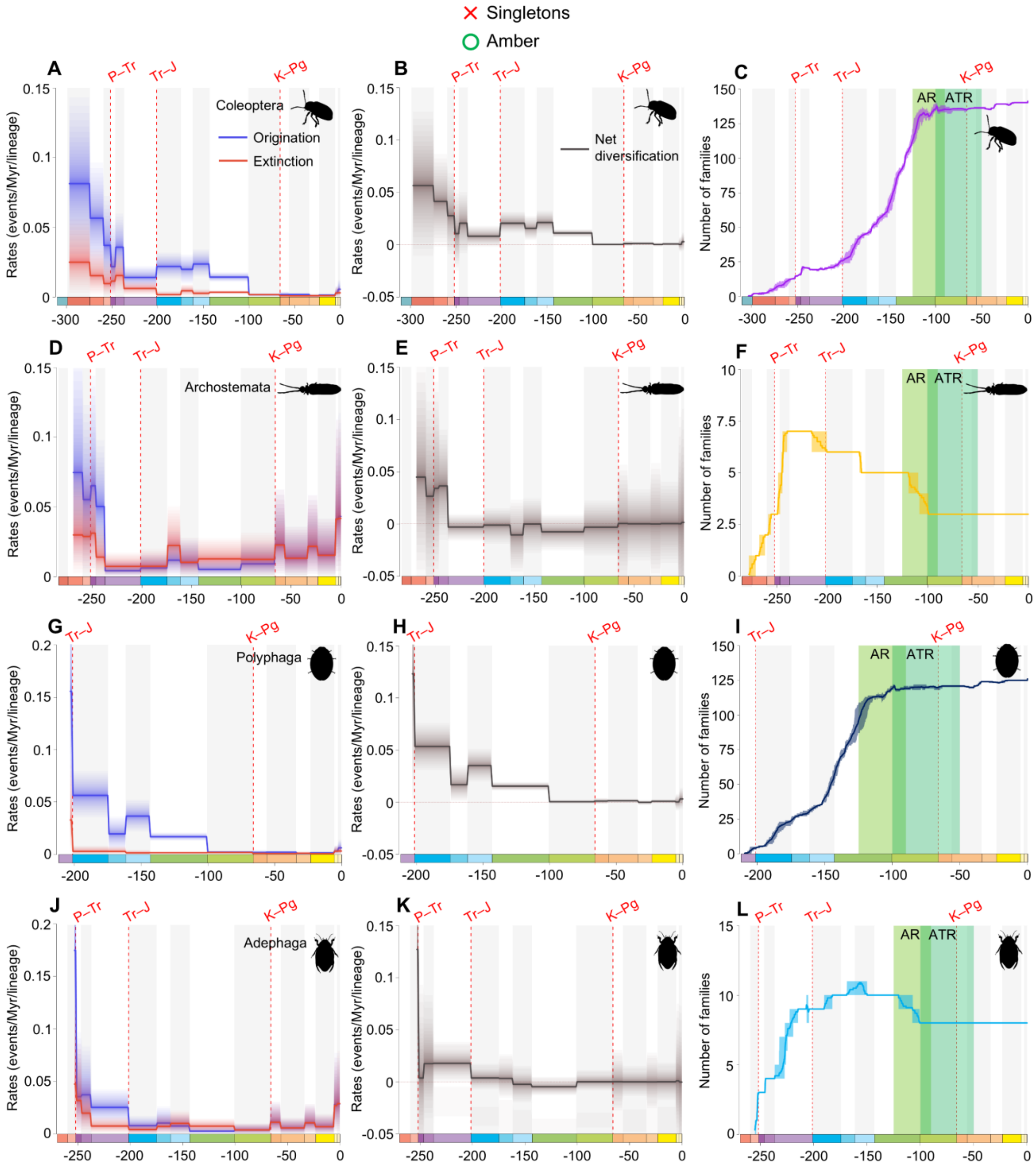


**fig. S1.** Diversification and diversity dynamics of Coleoptera, Archostemata, Polyphaga, and Adephaga families excluding singletons, but considering amber occurrences. Bayesian fossil-based inferences of Coleoptera (**A**), Archostemata (**D**), Polyphaga (**G**), and Adephaga (**J**) origination and extinction rates at the family level under the birth-death model with epochs as constrained shifts (BDCS), without singletons and considering amber occurrences. Net diversification rates for Coleoptera (**B**), Archostemata (**E**), Polyphaga (**H**), and Adephaga (**K**) obtained from the difference between origination and extinction rates (rates above 0 indicate increasing diversity, and rates below 0 indicate declining diversity). Solid lines indicate mean posterior rates and the shaded areas show 95% HPD. Number of families through time computed by summing up the lifespans of all genera for Coleoptera (**C**), Archostemata (**F**), Polyphaga (**I**), and Adephaga (**L**). Light-green area represents the AR, angiosperm radiation, and dark-green area represents the ATR, angiosperm terrestrial revolution. Solid lines indicate mean diversity at each point in time and shaded areas show estimations of different replications that incorporate age uncertainties of fossil occurrences. Red-dashed vertical lines indicate major crises: P–Tr, Permian–Triassic; Tr–J, Triassic–Jurassic; K–Pg, Cretaceous–Paleogene. Time is in millions of years. The color of each geological period in the chronostratigraphic scale follows that of the International Chronostratigraphic Chart (v2024/12). Insect silhouettes are from http://phylopic.org/.

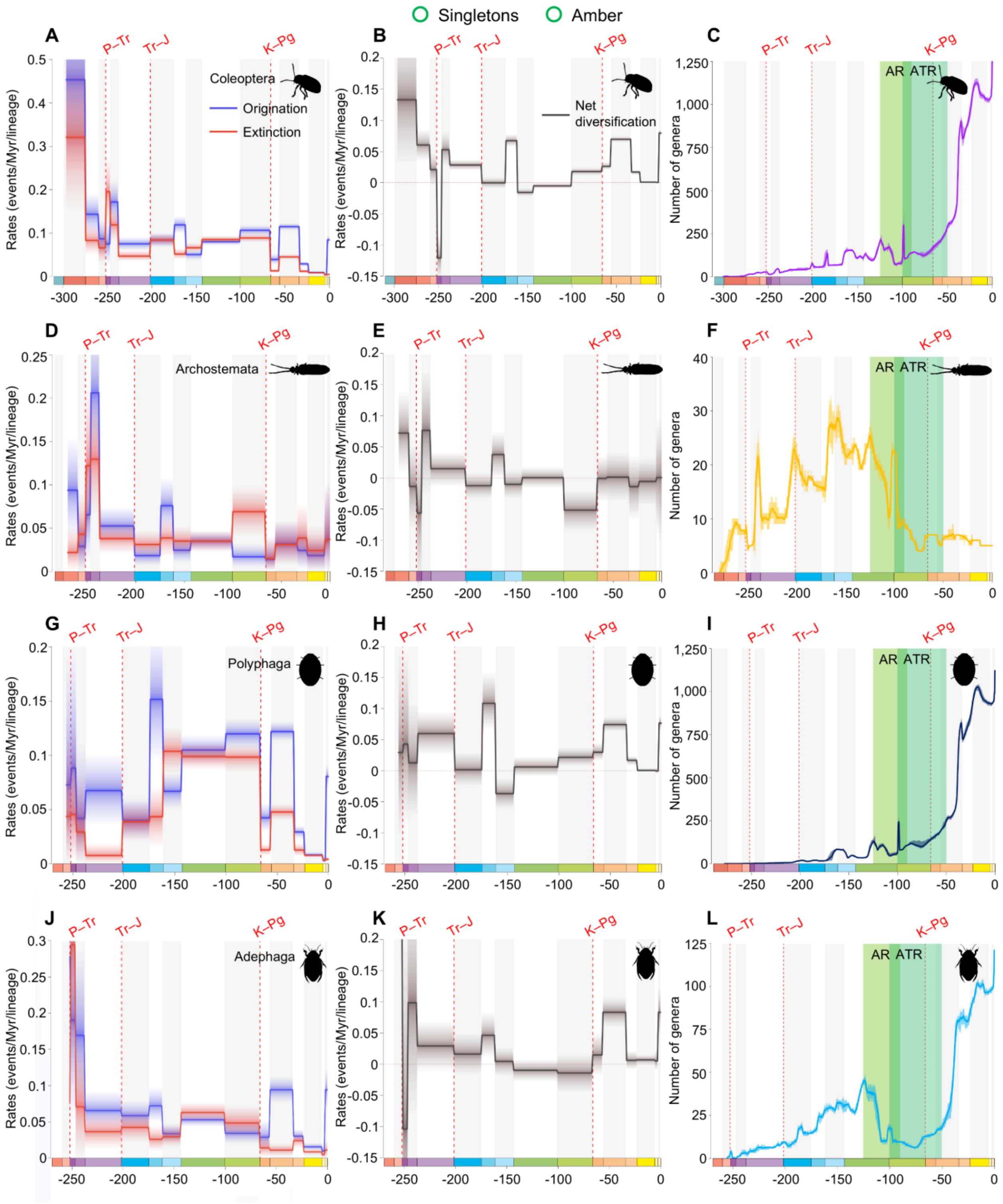


**fig. S2.** Diversification and diversity dynamics of Coleoptera, Archostemata, Polyphaga, and Adephaga genera, considering singletons and amber occurrences. Bayesian fossil-based inferences of Coleoptera (A), Archostemata (D), Polyphaga (G), and Adephaga (J) origination and extinction rates at the genus level under the birth-death model with epochs as constrained shifts (BDCS). Net diversification rates for Coleoptera (B), Archostemata (E), Polyphaga (H), and Adephaga (K) obtained from the difference between origination and extinction rates (rates above 0 indicate increasing diversity, and rates below 0 indicate declining diversity). Solid lines indicate mean posterior rates and the shaded areas show 95% HPD. Number of genera through time computed by summing up the lifespans of all genera for Coleoptera (C), Archostemata (F), Polyphaga (I), and Adephaga (L). Light-green area represents the AR, angiosperm radiation, and dark-green area represents the ATR, angiosperm terrestrial revolution. Solid lines indicate mean diversity at each point in time and shaded areas show estimations of different replications that incorporate age uncertainties of fossil occurrences. Red-dashed vertical lines indicate major crises: P–Tr, Permian–Triassic; Tr–J, Triassic–Jurassic; K–Pg, Cretaceous–Paleogene. Time is in millions of years. The color of each geological period in the chronostratigraphic scale follows that of the International Chronostratigraphic Chart (v2024/12). Insect silhouettes are from http://phylopic.org/.

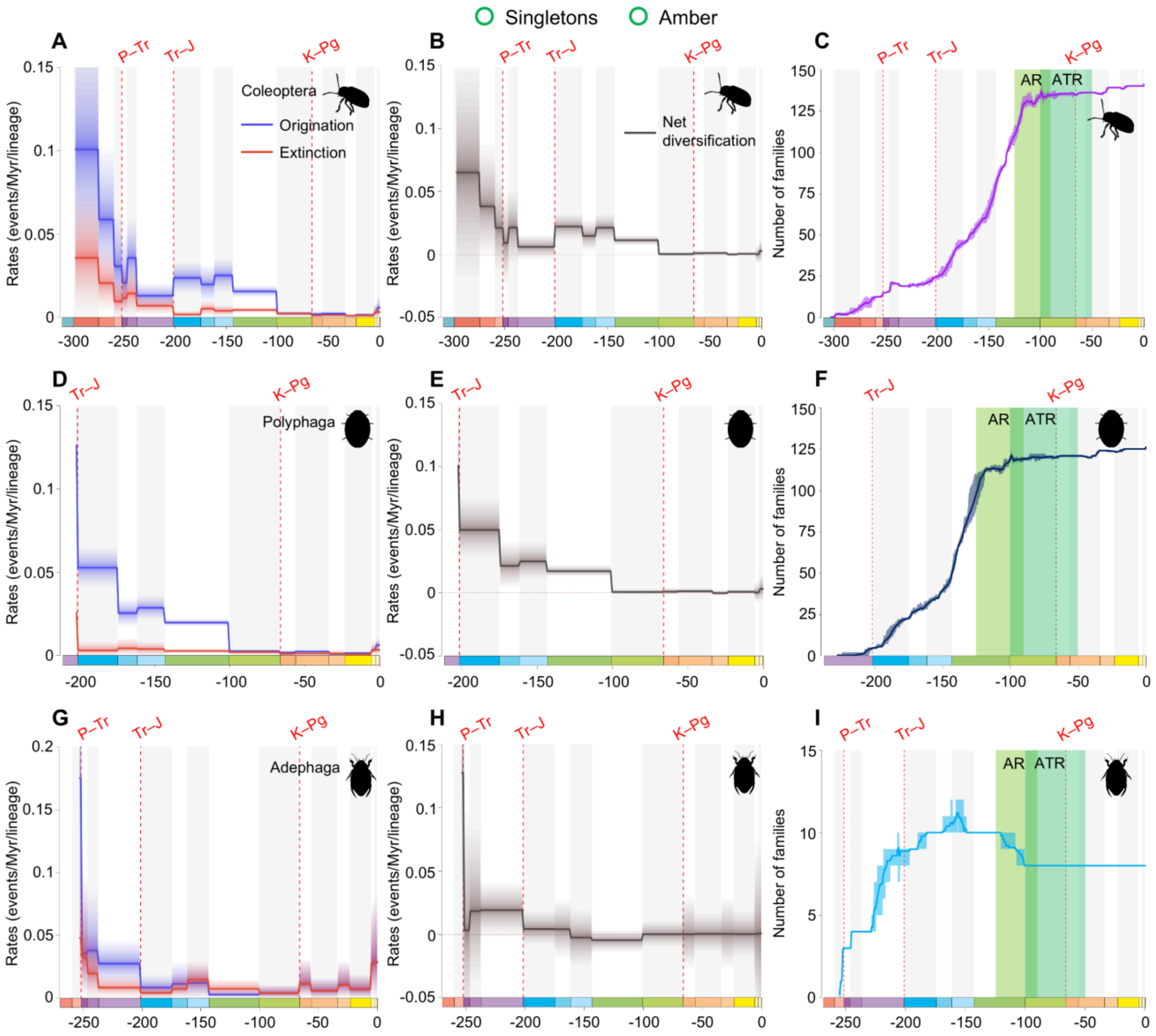


**fig. S3.** Diversification and diversity dynamics of Coleoptera, Polyphaga, and Adephaga families, considering singletons and amber occurrences. There are no singletons at the family-level in Archostemata. Bayesian fossil-based inferences of Coleoptera (A), Polyphaga (D), and Adephaga (G) origination and extinction rates at the family level under the birth-death model with epochs as constrained shifts (BDCS). Net diversification rates for Coleoptera (B), Polyphaga (E), and Adephaga (H) obtained from the difference between origination and extinction rates (rates above 0 indicate increasing diversity, and rates below 0 indicate declining diversity). Solid lines indicate mean posterior rates and the shaded areas show 95% HPD. Number of families through time computed by summing up the lifespans of all families for Coleoptera (C), Polyphaga (F), and Adephaga (I). Light-green area represents the AR, angiosperm radiation, and dark-green area represents the ATR, angiosperm terrestrial revolution. Solid lines indicate mean diversity at each point in time and shaded areas show estimations of different replications that incorporate age uncertainties of fossil occurrences. Red-dashed vertical lines indicate major crises: P–Tr, Permian–Triassic; Tr–J, Triassic–Jurassic; K–Pg, Cretaceous–Paleogene. Time is in millions of years. The color of each geological period in the chronostratigraphic scale follows that of the International Chronostratigraphic Chart (v2024/12). Insect silhouettes are from http://phylopic.org/.

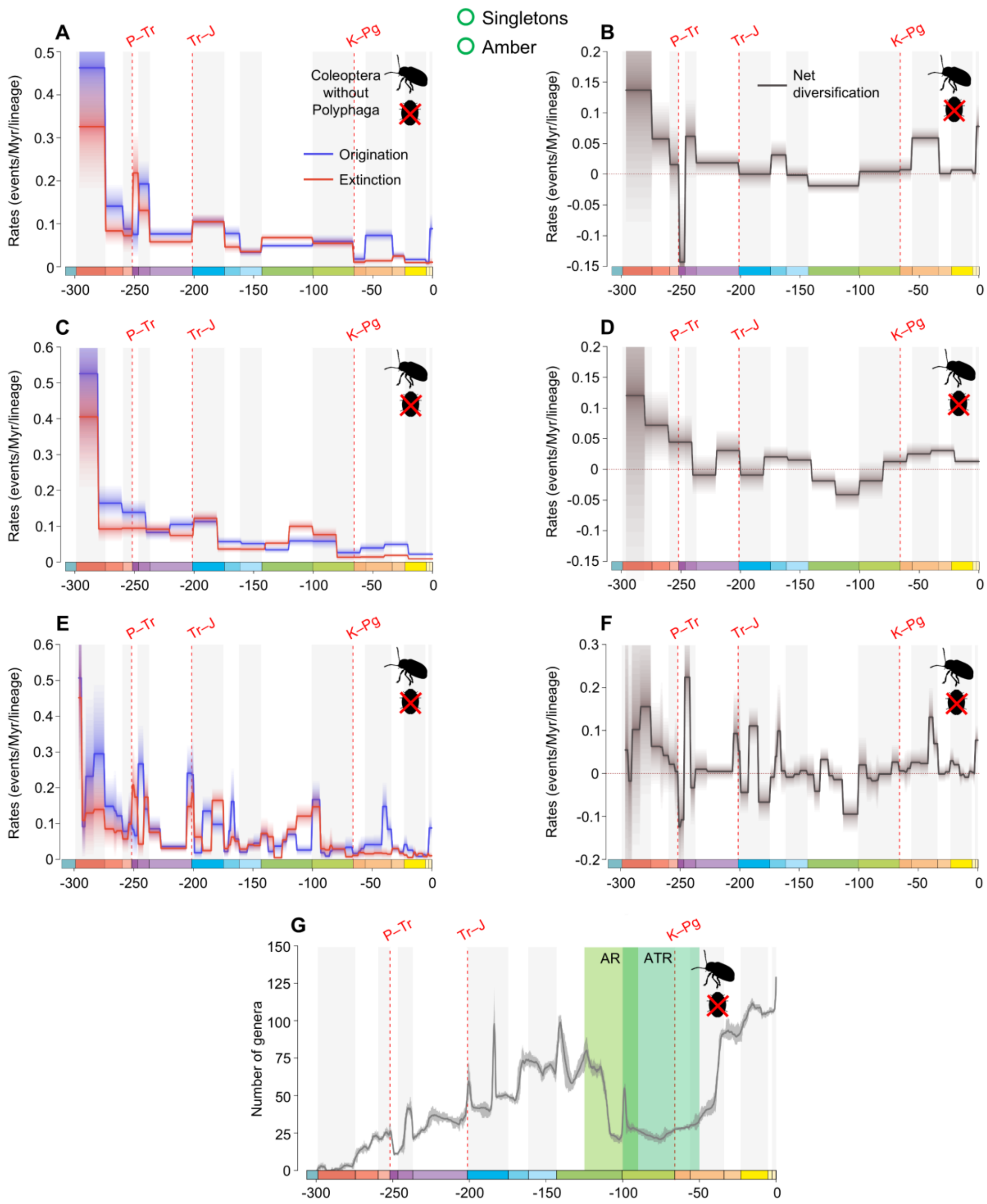


**fig. S4.** Diversification and diversity dynamics of Coleoptera without Polyphaga genera, considering singletons and amber occurrences. Bayesian fossil-based inferences of Coleoptera without Polyphaga origination and extinction rates at the genus level under the birth-death model with epochs (A), 20 Ma bins (C), and stages (E) as constrained shifts (BDCS). Net diversification rates for Coleoptera without Polyphaga obtained from the difference between origination and extinction rates (rates above 0 indicate increasing diversity, and rates below 0 indicate declining diversity) per epochs (B), 20 Ma bins (D), and stages (F). Solid lines indicate mean posterior rates

and the shaded areas show 95% HPD. Number of genera through time computed by summing up the lifespans of all genera for Coleoptera without Polyphaga (G). Solid lines indicate mean diversity at each point in time and shaded areas show estimations of different replications that incorporate age uncertainties of fossil occurrences. Light-green area represents the AR, angiosperm radiation, and dark-green area represents the ATR, angiosperm terrestrial revolution. Red-dashed vertical lines indicate major crises: P–Tr, Permian–Triassic; Tr–J, Triassic–Jurassic; K–Pg, Cretaceous–Paleogene. Time is in millions of years. The color of each geological period in the chronostratigraphic scale follows that of the International Chronostratigraphic Chart (v2024/12). Insect silhouettes are from http://phylopic.org/.

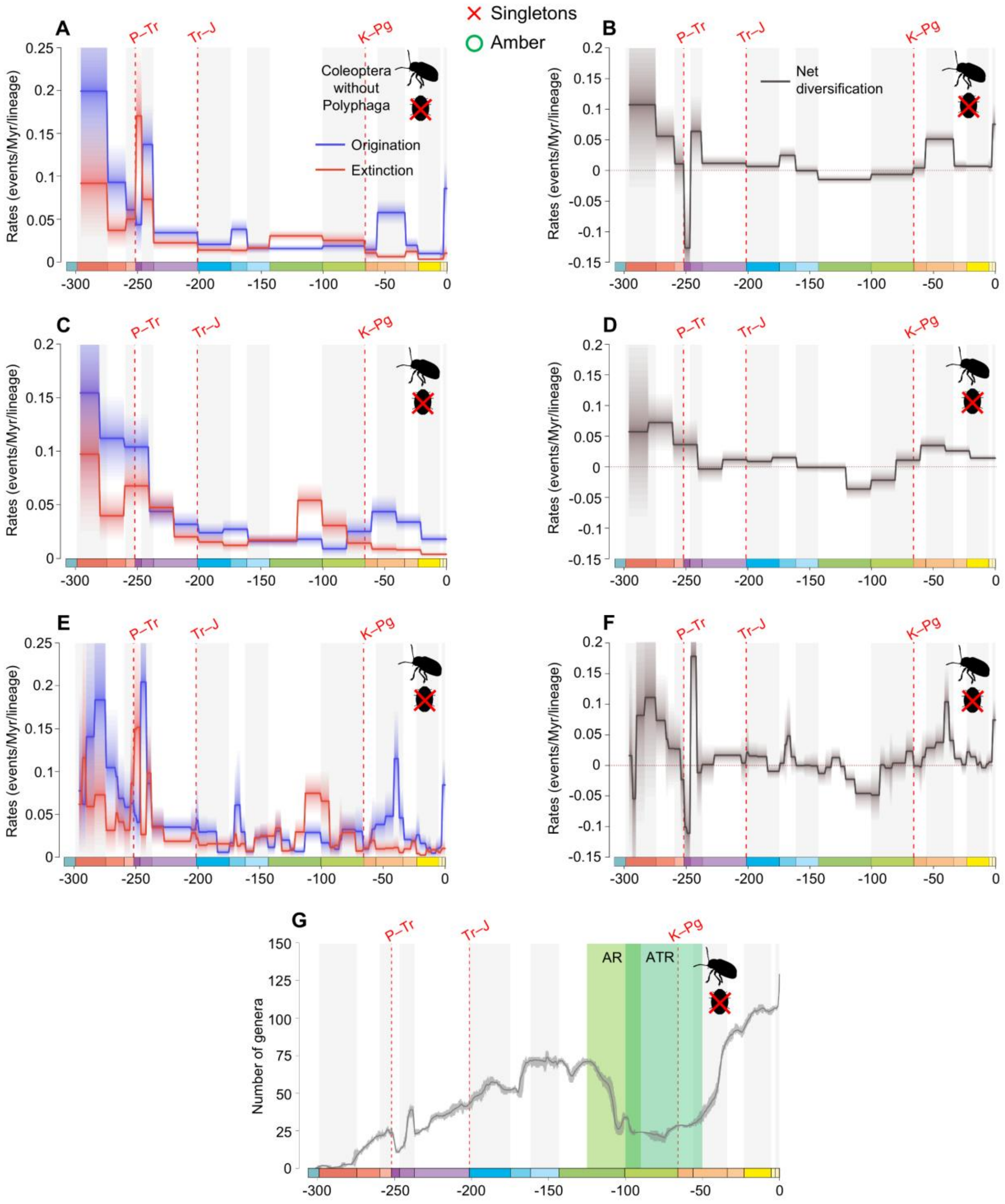


**fig. S5.** Diversification and diversity dynamics of Coleoptera without Polyphaga genera, excluding singletons, but considering amber occurrences. Bayesian fossil-based inferences of Coleoptera without Polyphaga origination and extinction rates at the genus level under the birth-death model with epochs (A), 20 Ma bins (C), and stages (E) as constrained shifts (BDCS). Net diversification rates for Coleoptera without Polyphaga obtained from the difference between origination and extinction rates (rates above 0 indicate increasing diversity, and rates below 0 indicate declining diversity) per epochs (B), 20 Ma bins (D), and stages (F). Solid lines indicate mean posterior rates

and the shaded areas show 95% HPD. Number of genera through time computed by summing up the lifespans of all genera for Coleoptera without Polyphaga (G). Solid lines indicate mean diversity at each point in time and shaded areas show estimations of different replications that incorporate age uncertainties of fossil occurrences. Light-green area represents the AR, angiosperm radiation, and dark-green area represents the ATR, angiosperm terrestrial revolution. Red-dashed vertical lines indicate major crises: P–Tr, Permian–Triassic; Tr–J, Triassic–Jurassic; K–Pg, Cretaceous–Paleogene. Time is in millions of years. The color of each geological period in the chronostratigraphic scale follows that of the International Chronostratigraphic Chart (v2024/12). Insect silhouettes are from http://phylopic.org/.

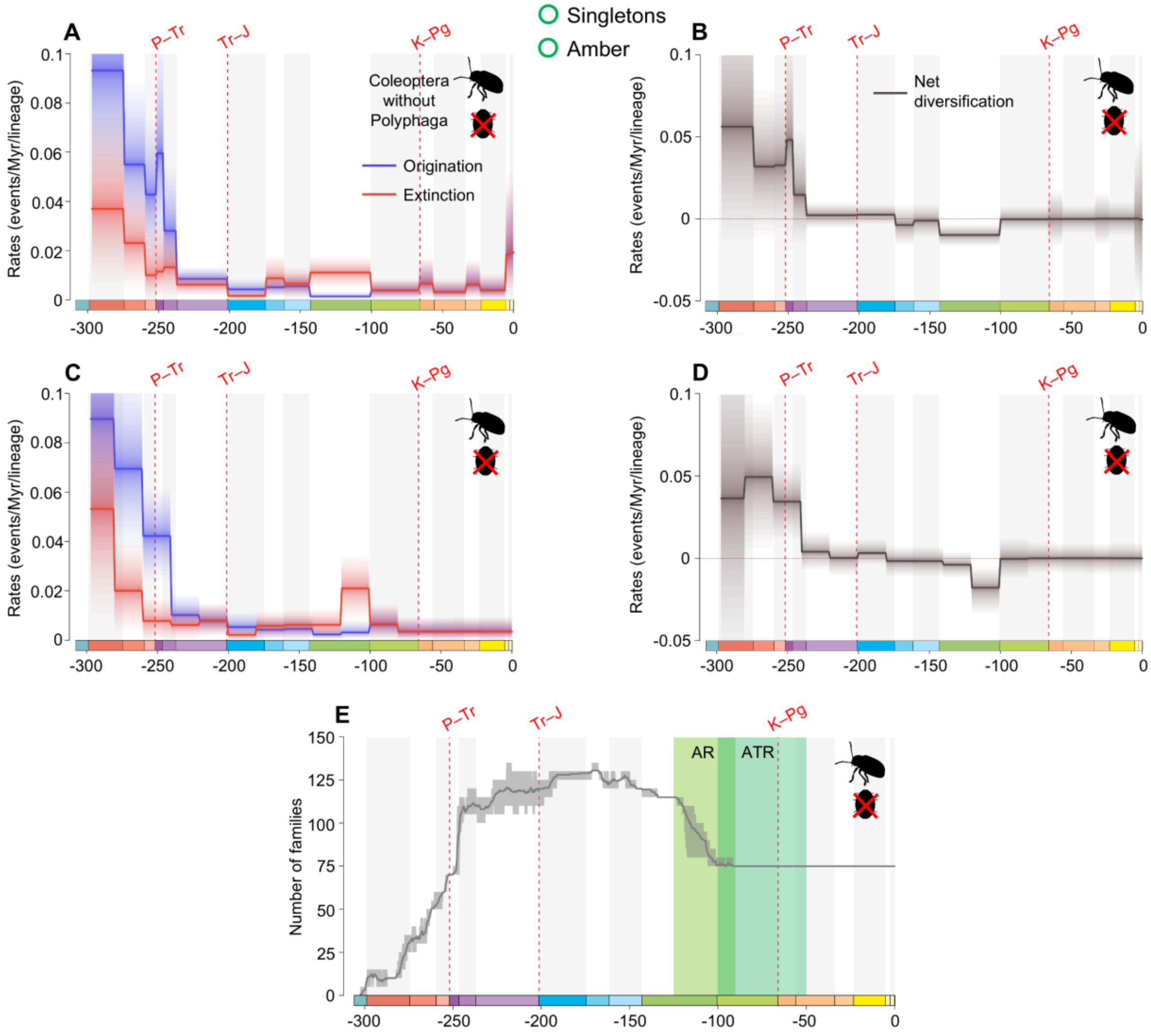


**fig. S6.** Diversification and diversity dynamics of Coleoptera without Polyphaga families, considering singletons and amber occurrences. Bayesian fossil-based inferences of Coleoptera without Polyphaga origination and extinction rates at the family level under the birth-death model with epochs (A), and 20 Ma bins (C) as constrained shifts (BDCS). Net diversification rates for Coleoptera without Polyphaga obtained from the difference between origination and extinction rates (rates above 0 indicate increasing diversity, and rates below 0 indicate declining diversity) per epochs (B), and 20 Ma bins (D). Solid lines indicate mean posterior rates and the shaded areas show 95% HPD. Number of families through time computed by summing up the lifespans of all families for Coleoptera without Polyphaga (E). Solid lines indicate mean diversity at each point in time and shaded areas show estimations of different replications that incorporate age uncertainties of fossil occurrences. Light-green area represents the AR, angiosperm radiation, and dark-green area represents the ATR, angiosperm terrestrial revolution. Red-dashed vertical lines indicate major crises: P–Tr, Permian–Triassic; Tr–J, Triassic–Jurassic; K–Pg, Cretaceous–Paleogene. Time is in millions of years. The color of each geological period in the chronostratigraphic scale follows that of the International Chronostratigraphic Chart (v2024/12). Insect silhouettes are from http://phylopic.org/.

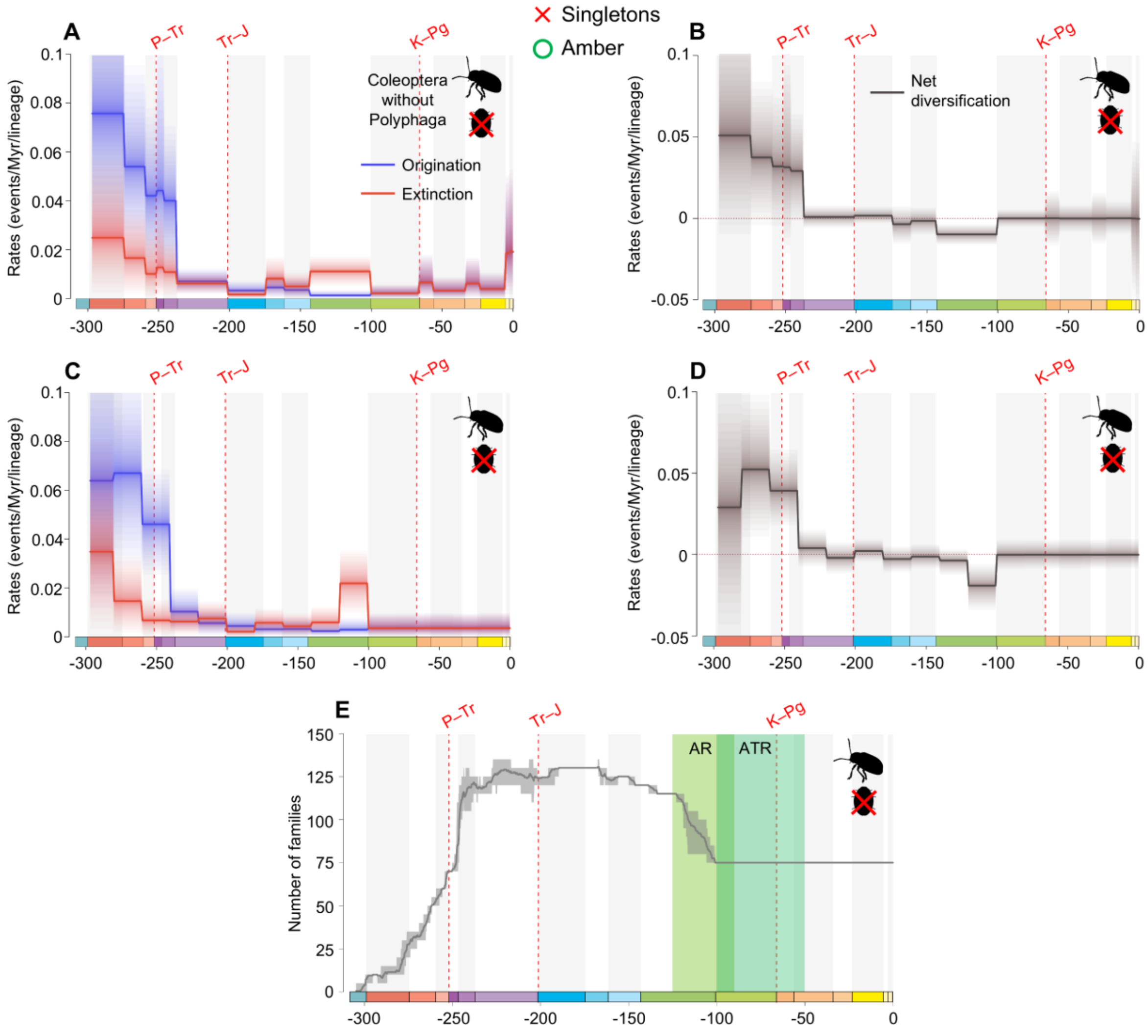


**fig. S7.** Diversification and diversity dynamics of Coleoptera without Polyphaga families, excluding singletons, but considering amber occurrences. Bayesian fossil-based inferences of Coleoptera without Polyphaga origination and extinction rates at the family level under the birth-death model with epochs (A), and 20 Ma bins (C) as constrained shifts (BDCS). Net diversification rates for Coleoptera without Polyphaga obtained from the difference between origination and extinction rates (rates above 0 indicate increasing diversity, and rates below 0 indicate declining diversity) per epochs (B), and 20 Ma bins (D). Solid lines indicate mean posterior rates and the shaded areas show 95% HPD. Number of families through time computed by summing up the lifespans of all families for Coleoptera without Polyphaga (E). Solid lines indicate mean diversity at each point in time and shaded areas show estimations of different replications that incorporate age uncertainties of fossil occurrences. Light-green area represents the AR, angiosperm radiation, and dark-green area represents the ATR, angiosperm terrestrial revolution. Red-dashed vertical lines indicate major crises: P–Tr, Permian–Triassic; Tr–J, Triassic–Jurassic; K–Pg, Cretaceous–Paleogene. Time is in millions of years. The color of each geological period in the chronostratigraphic scale follows that of the International Chronostratigraphic Chart (v2024/12). Insect silhouettes are from http://phylopic.org/.

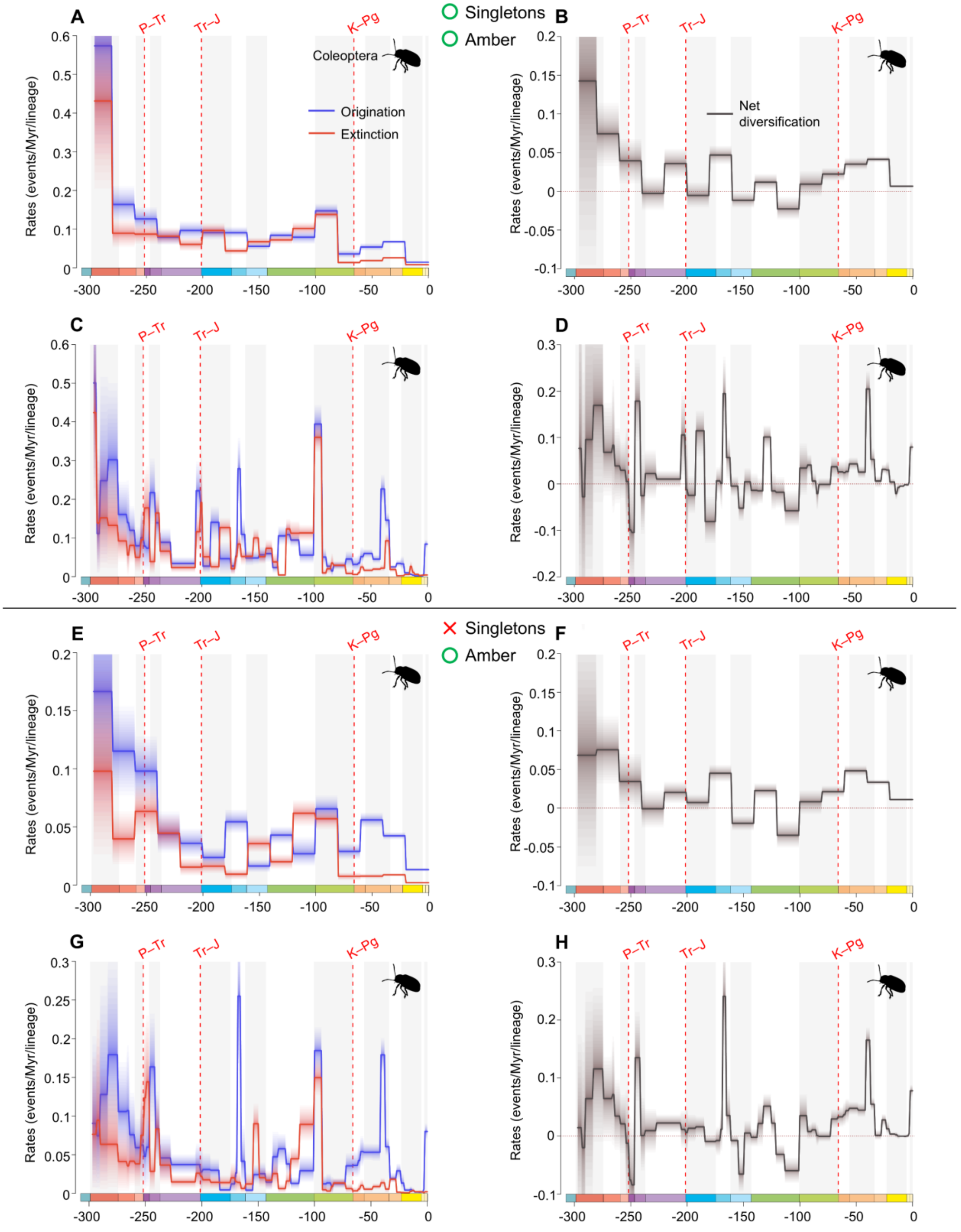


**fig. S8.** Diversification and diversity dynamics of Coleoptera genera, with and without singletons, but considering amber occurrences. Bayesian fossil-based inferences of Coleoptera origination and extinction rates at the genus level under the birth-death model with 20 Ma bins (A: with singletons;

E: without singletons), and stages (C: with singletons; G: without singletons) as constrained shifts (BDCS). Net diversification rates for Coleoptera obtained from the difference between origination and extinction rates (rates above 0 indicate increasing diversity, and rates below 0 indicate declining diversity) per 20 Ma bins (B: with singletons; F: without singletons), and stages (D: with singletons; H: without singletons). Solid lines indicate mean posterior rates and the shaded areas show 95% HPD. Red-dashed vertical lines indicate major crises: P–Tr, Permian–Triassic; Tr–J, Triassic–Jurassic; K–Pg, Cretaceous–Paleogene. Time is in millions of years. The color of each geological period in the chronostratigraphic scale follows that of the International Chronostratigraphic Chart (v2024/12). Insect silhouettes are from http://phylopic.org/.

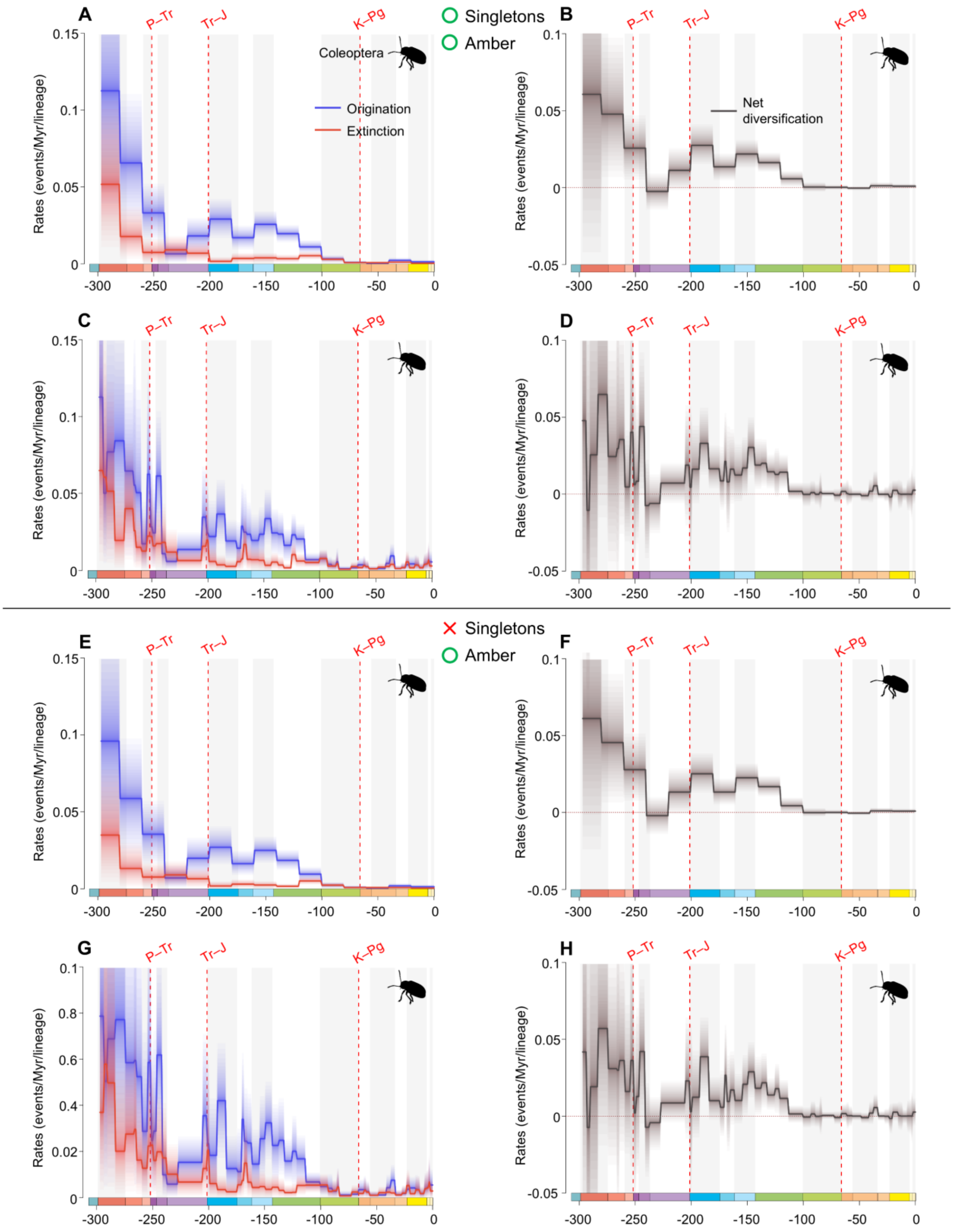


**fig. S9.** Diversification and diversity dynamics of Coleoptera families, with and without singletons, but considering amber occurrences. Bayesian fossil-based inferences of Coleoptera origination and extinction rates at the family level under the birth-death model with 20 Ma bins (A: with singletons;

E: without singletons), and stages (C: with singletons; G: without singletons) as constrained shifts (BDCS). Net diversification rates for Coleoptera obtained from the difference between origination and extinction rates (rates above 0 indicate increasing diversity, and rates below 0 indicate declining diversity) per 20 Ma bins (B: with singletons; F: without singletons), and stages (D: with singletons; H: without singletons). Solid lines indicate mean posterior rates and the shaded areas show 95% HPD. Red-dashed vertical lines indicate major crises: P–Tr, Permian–Triassic; Tr–J, Triassic–Jurassic; K–Pg, Cretaceous–Paleogene. Time is in millions of years. The color of each geological period in the chronostratigraphic scale follows that of the International Chronostratigraphic Chart (v2024/12). Insect silhouettes are from http://phylopic.org/.

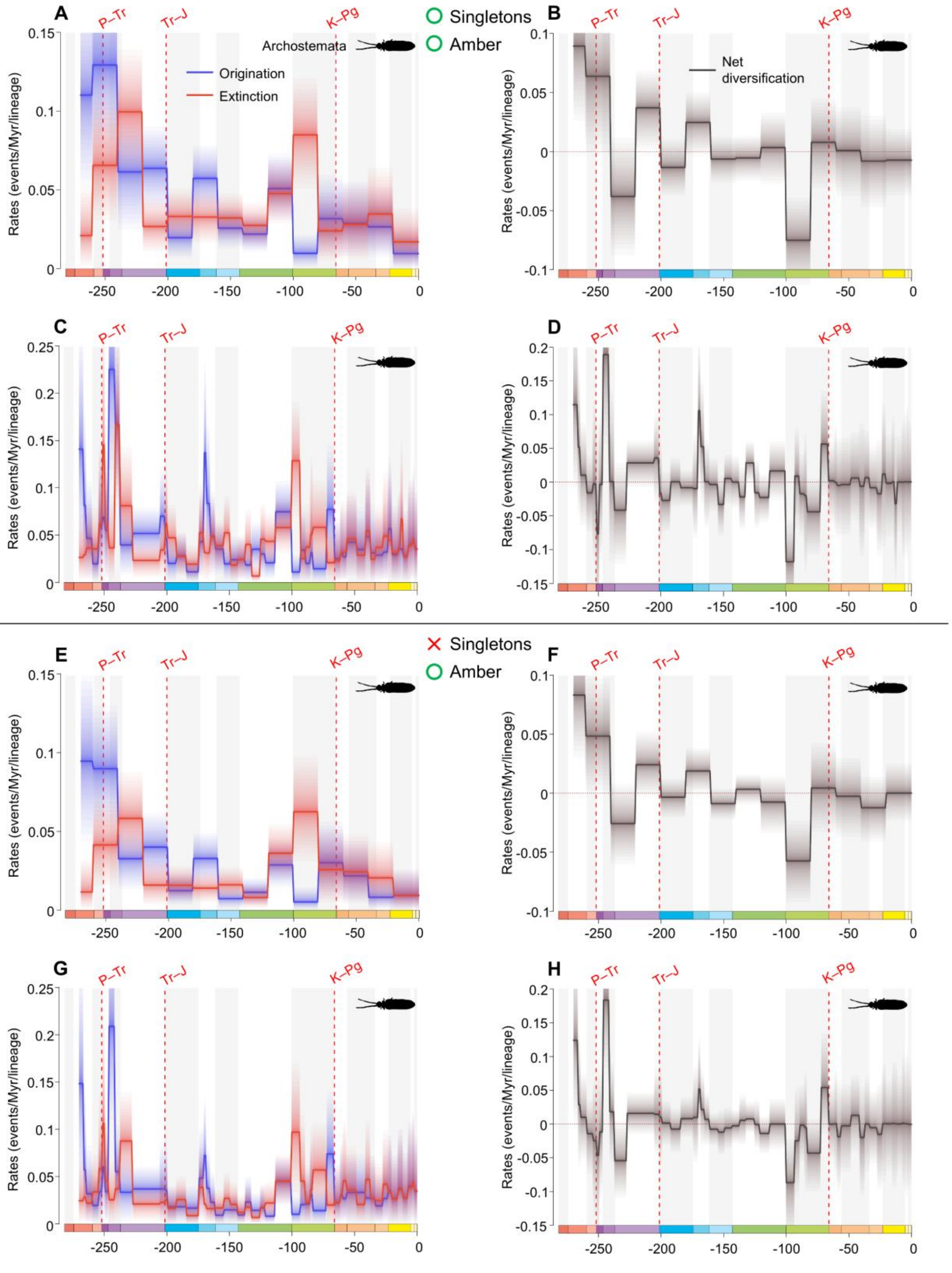


**fig. S10.** Diversification and diversity dynamics of Archostemata genera, with and without singletons, but considering amber occurrences. Bayesian fossil-based inferences of Archostemata origination and extinction rates at the genus level under the birth-death model with 20 Ma bins (A:

with singletons; E: without singletons), and stages (C: with singletons; G: without singletons) as constrained shifts (BDCS). Net diversification rates for Archostemata obtained from the difference between origination and extinction rates (rates above 0 indicate increasing diversity, and rates below 0 indicate declining diversity) per 20 Ma bins (B: with singletons; F: without singletons), and stages (D: with singletons; H: without singletons). Solid lines indicate mean posterior rates and the shaded areas show 95% HPD. Red-dashed vertical lines indicate major crises: P–Tr, Permian–Triassic; Tr–J, Triassic–Jurassic; K–Pg, Cretaceous–Paleogene. Time is in millions of years. The color of each geological period in the chronostratigraphic scale follows that of the International Chronostratigraphic Chart (v2024/12). Insect silhouettes are from http://phylopic.org/.

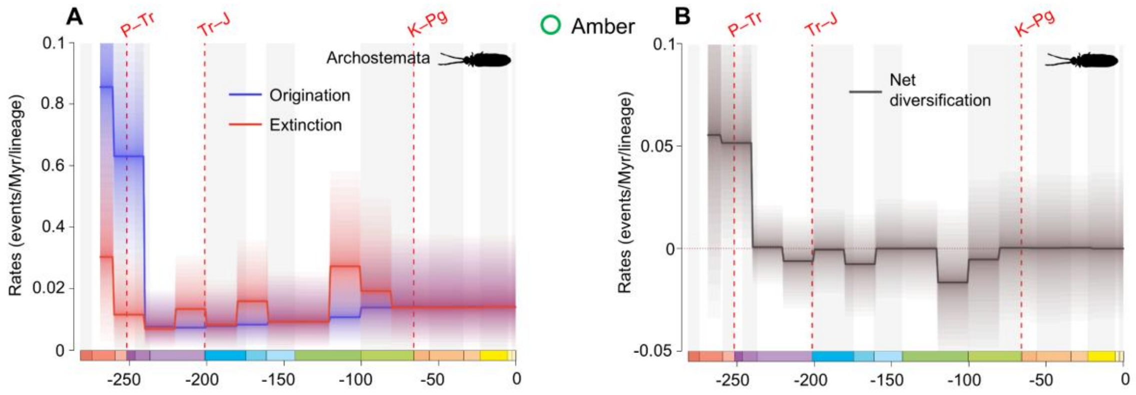


**fig. S11.** Diversification and diversity dynamics of Archostemata families, considering amber occurrences. There are no singletons in Archostemata at the family-level. Bayesian fossil-based inferences of Archostemata origination and extinction rates at the family level under the birth-death model with 20 Ma bins (A) as constrained shifts (BDCS). Net diversification rates for Coleoptera obtained from the difference between origination and extinction rates (rates above 0 indicate increasing diversity, and rates below 0 indicate declining diversity) per 20 Ma bins (B). Solid lines indicate mean posterior rates and the shaded areas show 95% HPD. Red-dashed vertical lines indicate major crises: P–Tr, Permian–Triassic; Tr–J, Triassic–Jurassic; K–Pg, Cretaceous–Paleogene. Time is in millions of years. The color of each geological period in the chronostratigraphic scale follows that of the International Chronostratigraphic Chart (v2024/12). Insect silhouettes are from http://phylopic.org/.

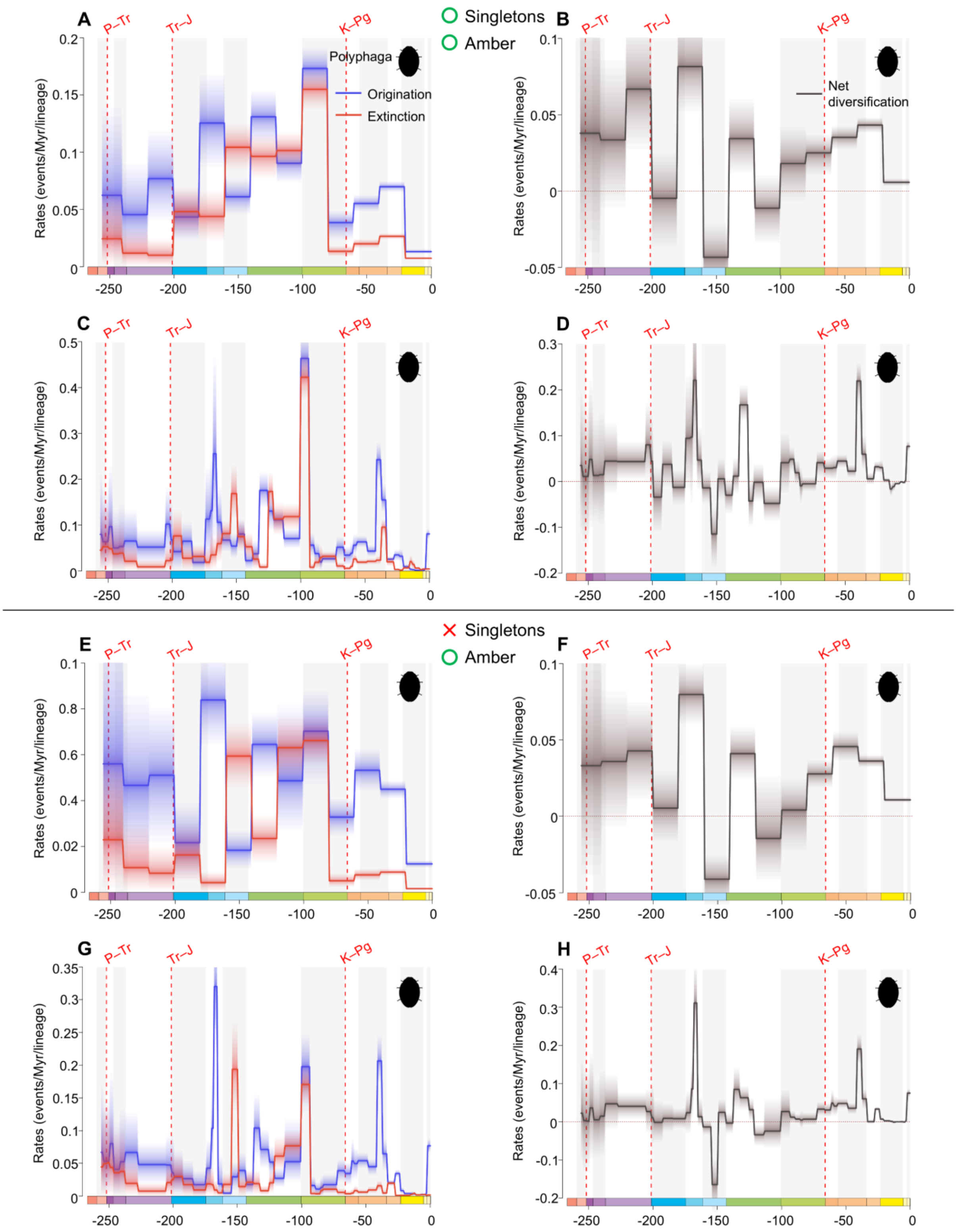


**fig. S12.** Diversification and diversity dynamics of Polyphaga genera, with and without singletons, but considering amber occurrences. Bayesian fossil-based inferences of Polyphaga origination and extinction rates at the genus level under the birth-death model with 20 Ma bins (A: with singletons;

E: without singletons), and stages (C: with singletons; G: without singletons) as constrained shifts (BDCS). Net diversification rates for Polyphaga obtained from the difference between origination and extinction rates (rates above 0 indicate increasing diversity, and rates below 0 indicate declining diversity) per 20 Ma bins (B: with singletons; F: without singletons), and stages (D: with singletons; H: without singletons). Solid lines indicate mean posterior rates and the shaded areas show 95% HPD. Red-dashed vertical lines indicate major crises: P–Tr, Permian–Triassic; Tr–J, Triassic–Jurassic; K–Pg, Cretaceous–Paleogene. Time is in millions of years. The color of each geological period in the chronostratigraphic scale follows that of the International Chronostratigraphic Chart (v2024/12). Insect silhouettes are from http://phylopic.org/.

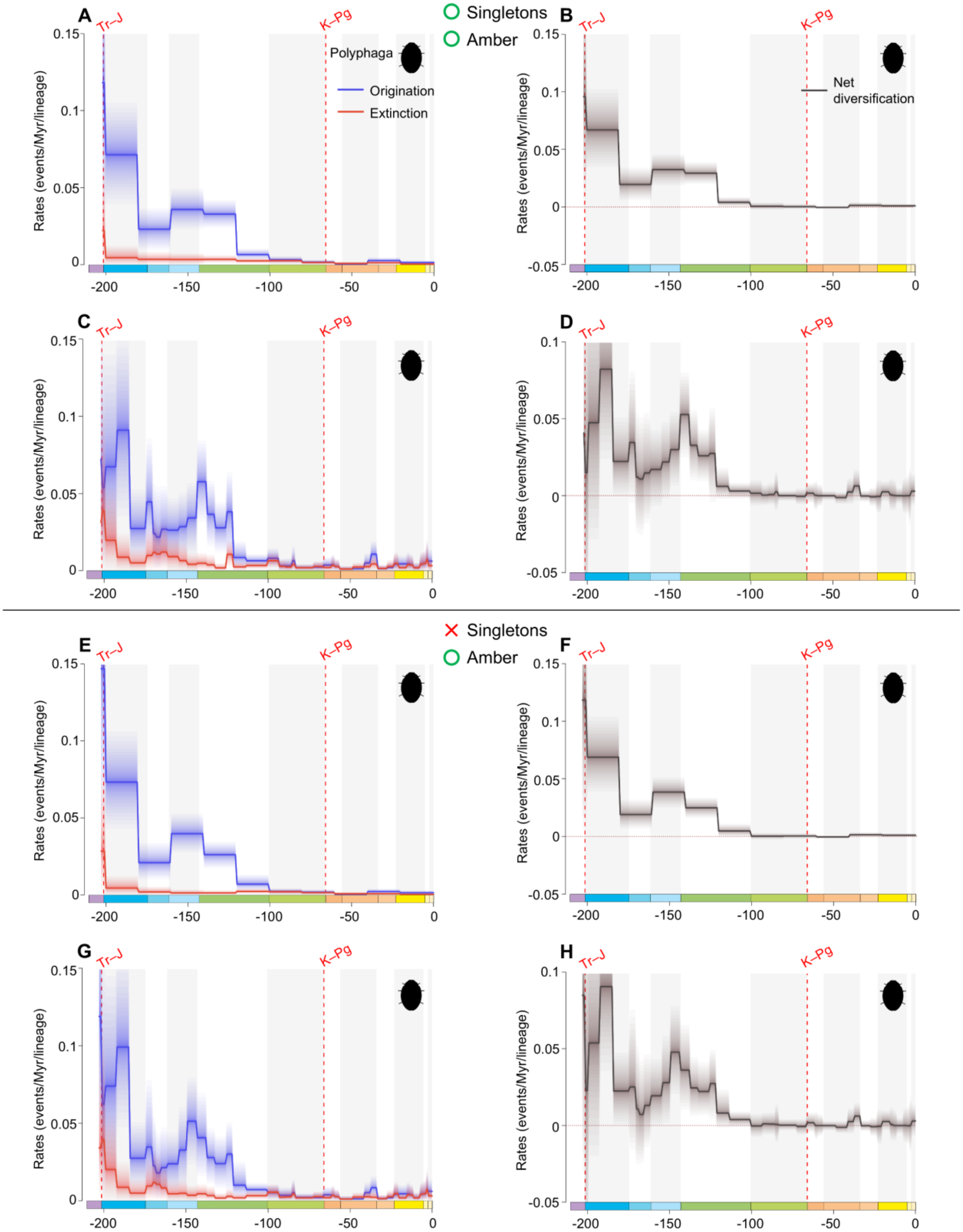


**fig. S13.** Diversification and diversity dynamics of Polyphaga families, with and without singletons, but considering amber occurrences. Bayesian fossil-based inferences of Polyphaga origination and extinction rates at the family level under the birth-death model with 20 Ma bins

(A: with singletons; E: without singletons), and stages (C: with singletons; G: without singletons) as constrained shifts (BDCS). Net diversification rates for Polyphaga obtained from the difference between origination and extinction rates (rates above 0 indicate increasing diversity, and rates below 0 indicate declining diversity) per 20 Ma bins (B: with singletons; F: without singletons), and stages (D: with singletons; H: without singletons). Solid lines indicate mean posterior rates and the shaded areas show 95% HPD. Red-dashed vertical lines indicate major crises: Tr–J, Triassic–Jurassic; K–Pg, Cretaceous–Paleogene. Time is in millions of years. The color of each geological period in the chronostratigraphic scale follows that of the International Chronostratigraphic Chart (v2024/12). Insect silhouettes are from http://phylopic.org/.

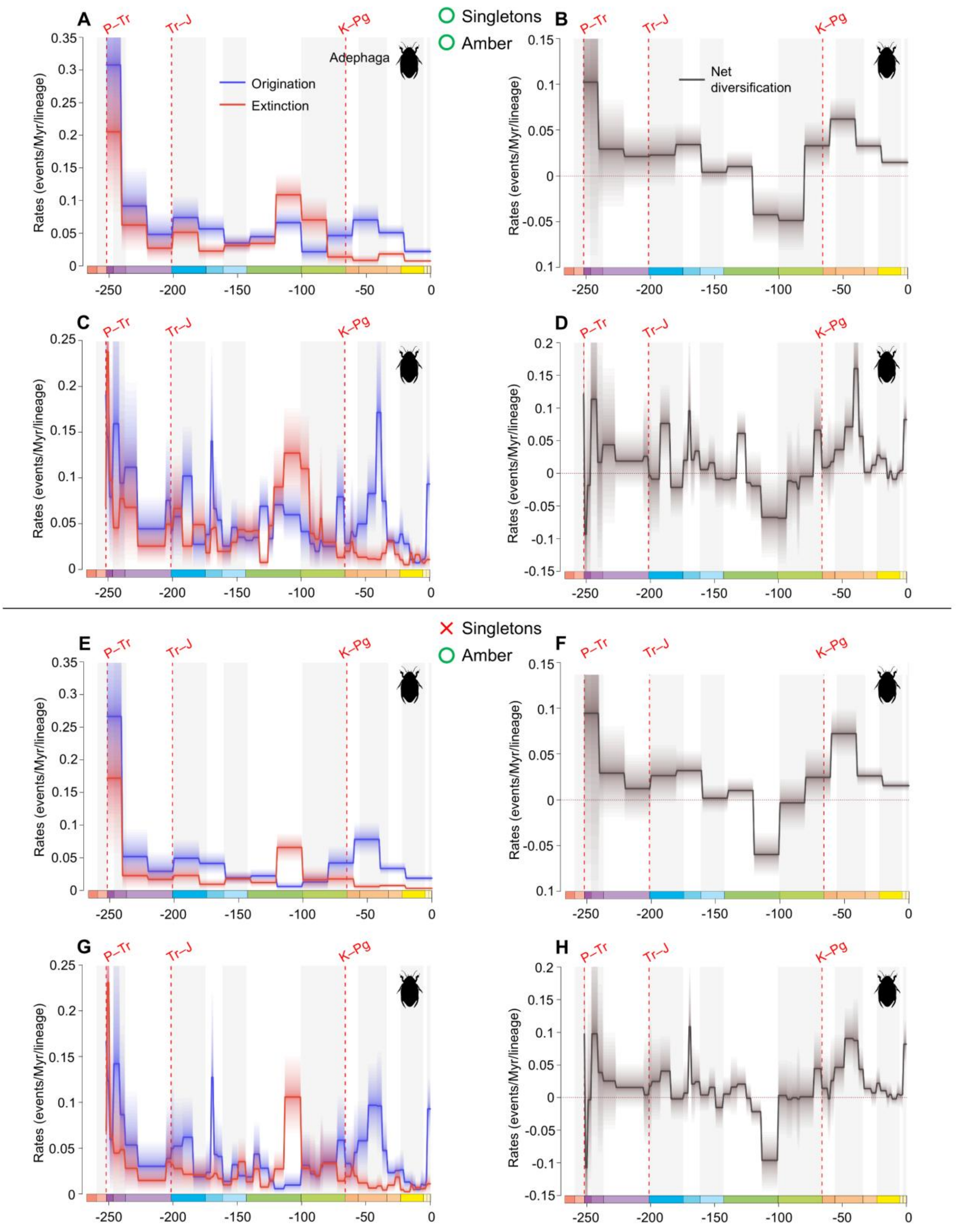


**fig. S14.** Diversification and diversity dynamics of Adephaga genera, with and without singletons, but considering amber occurrences. Bayesian fossil-based inferences of Adephaga origination and extinction rates at the genus level under the birth-death model with 20 Ma bins (A: with singletons;

E: without singletons), and stages (C: with singletons; G: without singletons) as constrained shifts (BDCS). Net diversification rates for Adephaga obtained from the difference between origination and extinction rates (rates above 0 indicate increasing diversity, and rates below 0 indicate declining diversity) per 20 Ma bins (B: with singletons; F: without singletons), and stages (D: with singletons; H: without singletons). Solid lines indicate mean posterior rates and the shaded areas show 95% HPD. Red-dashed vertical lines indicate major crises: P–Tr, Permian–Triassic; Tr–J, Triassic–Jurassic; K–Pg, Cretaceous–Paleogene. Time is in millions of years. The color of each geological period in the chronostratigraphic scale follows that of the International Chronostratigraphic Chart (v2024/12). Insect silhouettes are from http://phylopic.org/.

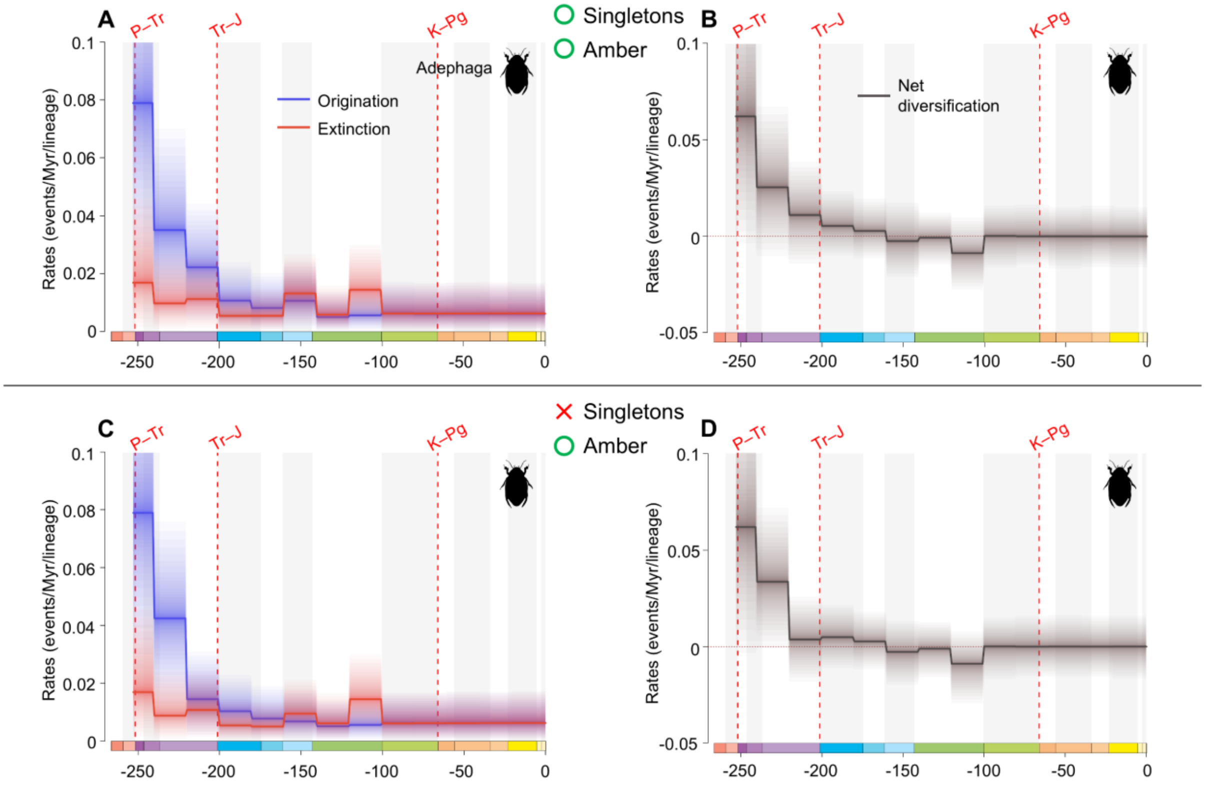


**fig. S15.** Diversification and diversity dynamics of Adephaga families, with and without singletons, but considering amber occurrences. Bayesian fossil-based inferences of Adephaga origination and extinction rates at the family level under the birth-death model with 20 Ma bins (A: with singletons; C: without singletons) as constrained shifts (BDCS). Net diversification rates for Adephaga obtained from the difference between origination and extinction rates (rates above 0 indicate increasing diversity, and rates below 0 indicate declining diversity) per 20 Ma bins (B: with singletons; D: without singletons). Solid lines indicate mean posterior rates and the shaded areas show 95% HPD. Red-dashed vertical lines indicate major crises: P–Tr, Permian–Triassic; Tr–J, Triassic–Jurassic; K–Pg, Cretaceous–Paleogene. Time is in millions of years. The color of each geological period in the chronostratigraphic scale follows that of the International Chronostratigraphic Chart (v2024/12). Insect silhouettes are from http://phylopic.org/.

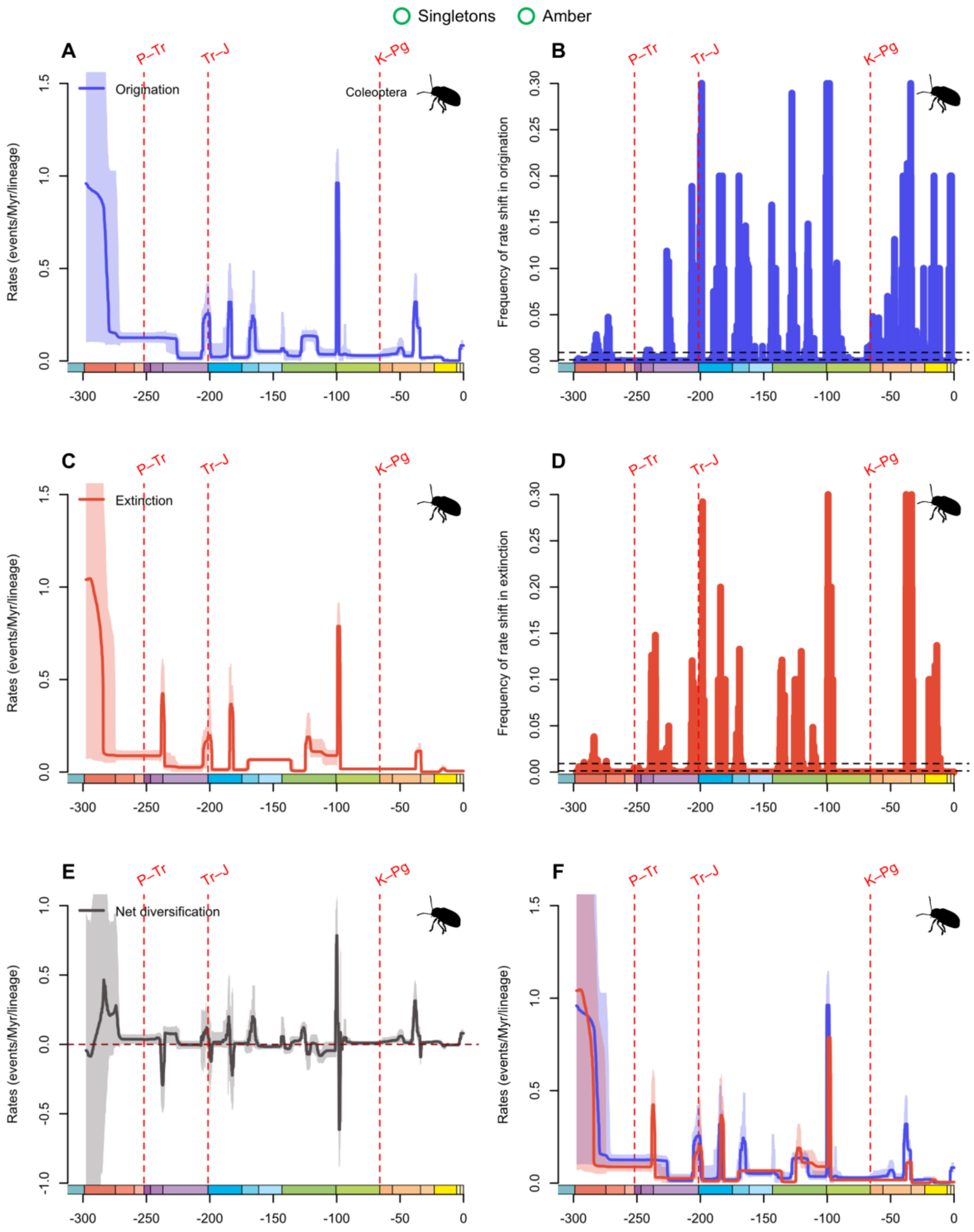


**fig. S16.** Diversification and diversity dynamics of Coleoptera genera, considering singletons and amber occurrences. Bayesian estimations of origination and extinction rates through time as inferred by PyRate using reversible jump Markov Chain Monte Carlo (RJMCMC). Marginal estimates of origination rates (A) and extinction rates (C) through time are shown as mean and 95% HPD (shaded areas). The frequency of a sampled rate shift is computed within small time bins for origination and extinction rates (B and D, respectively), with black-dashed horizontal lines

indicating log-Bayes factors of 2 (bottom) and 6 (top). Sampling frequencies higher than log-Bayes factors = 6 indicate strong statistical support for a rate shift. Panel (E) shows net diversification rates through time (computed as the posterior difference between origination and extinction rates through time). Panel (F) shows both marginal estimates of origination and extinction rates through time on the same plot. Solid lines indicate mean posterior rates and the shaded areas show 95% HPD. Red-dashed vertical lines indicate major crises: P–Tr, Permian–Triassic; Tr–J, Triassic–Jurassic; K–Pg, Cretaceous–Paleogene. Time is in millions of years. The color of each geological period in the chronostratigraphic scale follows that of the International Chronostratigraphic Chart (v2024/12). Insect silhouettes are from http://phylopic.org/.

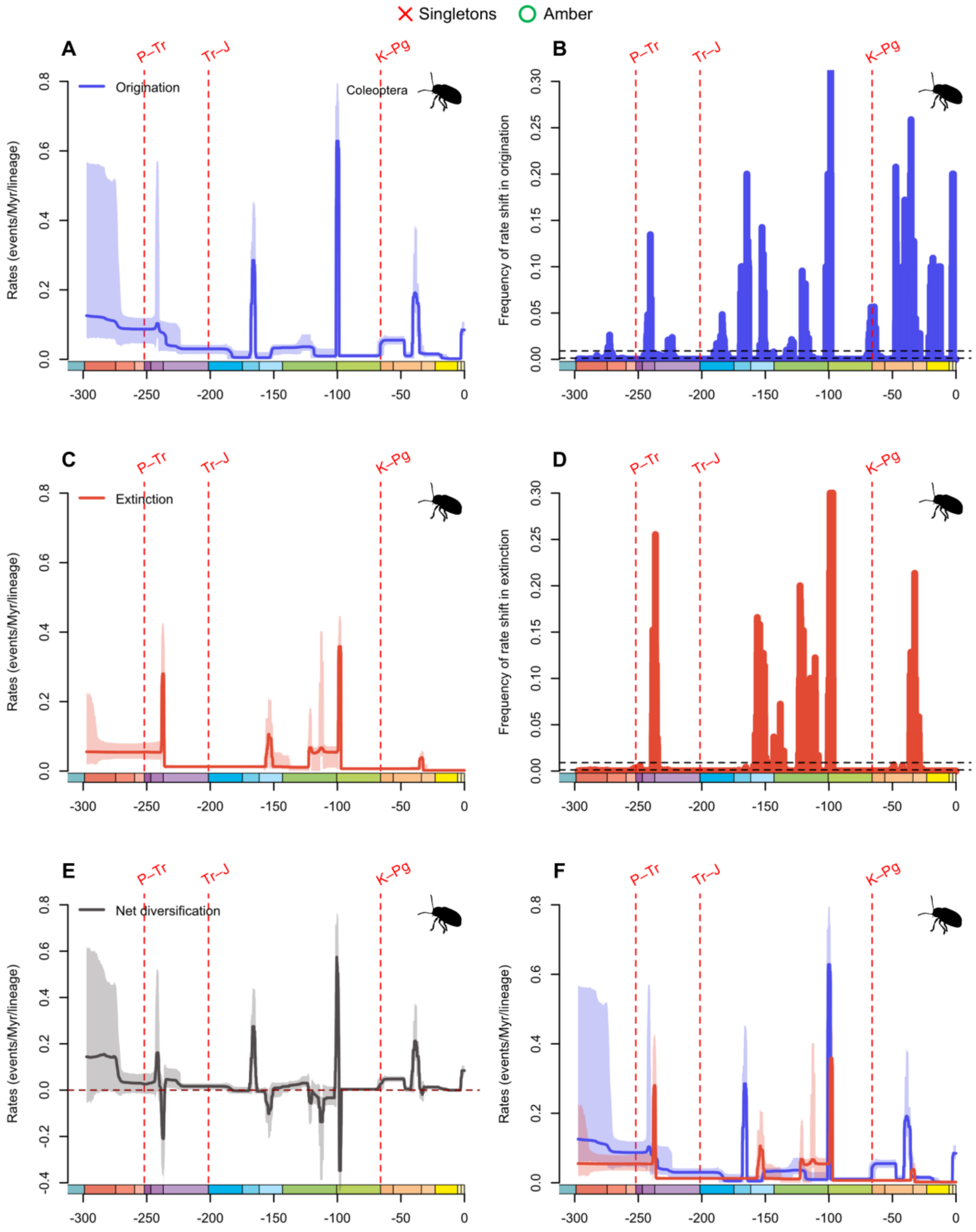


**fig. S17.** Diversification and diversity dynamics of Coleoptera genera, excluding singletons, but considering amber occurrences. Bayesian estimations of origination and extinction rates through time as inferred by PyRate using reversible jump Markov Chain Monte Carlo (RJMCMC). Marginal estimates of origination rates (A) and extinction rates (C) through time are shown as mean and 95% HPD (shaded areas). The frequency of a sampled rate shift is computed within small time bins for origination and extinction rates (B and D, respectively), with black-dashed

horizontal lines indicating log-Bayes factors of 2 (bottom) and 6 (top). Sampling frequencies higher than log-Bayes factors = 6 indicate strong statistical support for a rate shift. Panel (E) shows net diversification rates through time (computed as the posterior difference between origination and extinction rates through time). Panel (F) shows both marginal estimates of origination and extinction rates through time on the same plot. Solid lines indicate mean posterior rates and the shaded areas show 95% HPD. Red-dashed vertical lines indicate major crises: P–Tr, Permian–Triassic; Tr–J, Triassic–Jurassic; K–Pg, Cretaceous–Paleogene. Time is in millions of years. The color of each geological period in the chronostratigraphic scale follows that of the International Chronostratigraphic Chart (v2024/12). Insect silhouettes are from http://phylopic.org/.

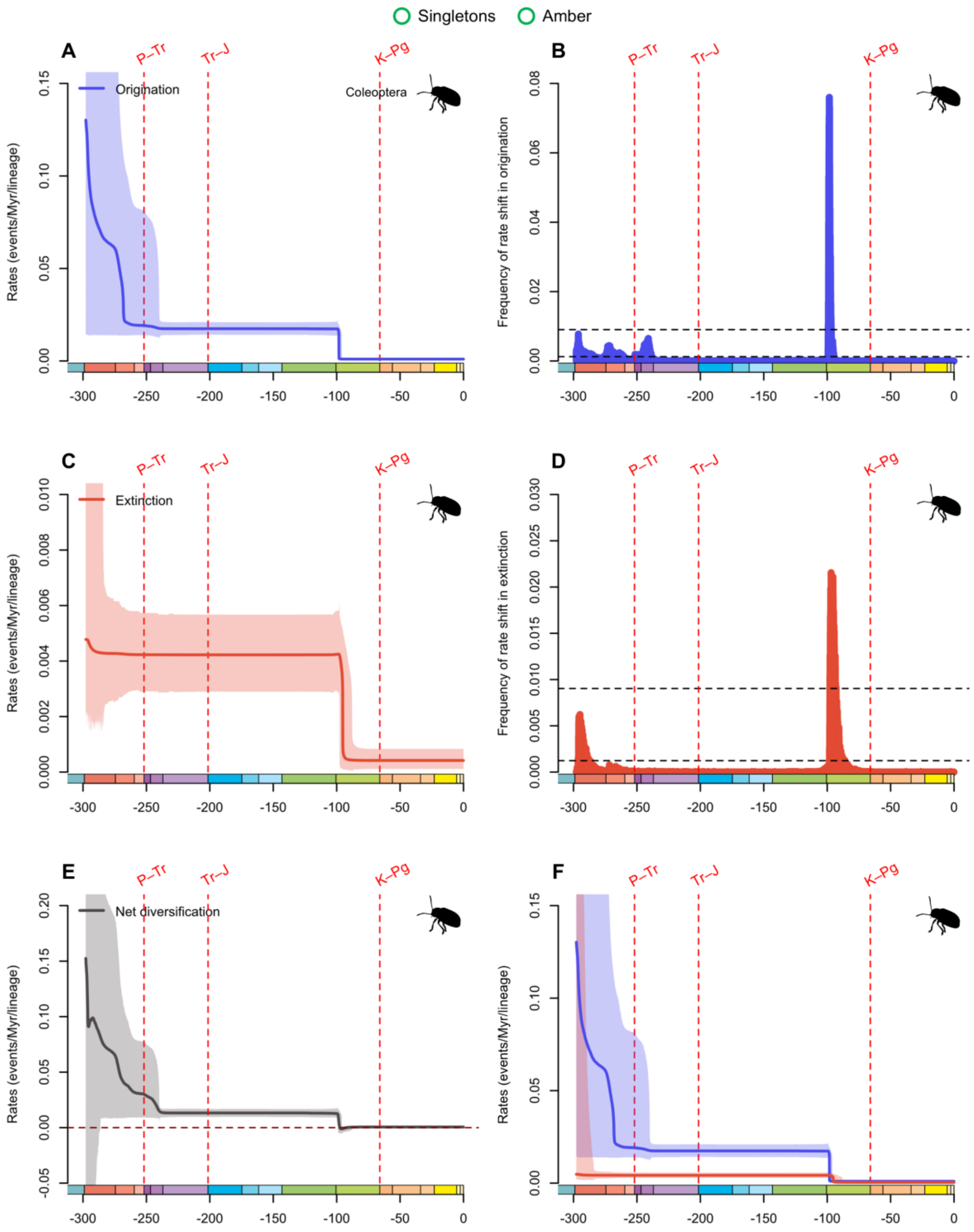


**fig. S18.** Diversification and diversity dynamics of Coleoptera families, considering singletons and amber occurrences. Bayesian estimations of origination and extinction rates through time as inferred by PyRate using reversible jump Markov Chain Monte Carlo (RJMCMC). Marginal estimates of origination rates (A) and extinction rates (C) through time are shown as mean and 95% HPD (shaded areas). The frequency of a sampled rate shift is computed within small time bins for origination and extinction rates (B and D, respectively), with black-dashed horizontal lines

indicating log-Bayes factors of 2 (bottom) and 6 (top). Sampling frequencies higher than log-Bayes factors = 6 indicate strong statistical support for a rate shift. Panel (E) shows net diversification rates through time (computed as the posterior difference between origination and extinction rates through time). Panel (F) shows both marginal estimates of origination and extinction rates through time on the same plot. Solid lines indicate mean posterior rates and the shaded areas show 95% HPD. Red-dashed vertical lines indicate major crises: P–Tr, Permian–Triassic; Tr–J, Triassic–Jurassic; K–Pg, Cretaceous–Paleogene. Time is in millions of years. The color of each geological period in the chronostratigraphic scale follows that of the International Chronostratigraphic Chart (v2024/12). Insect silhouettes are from http://phylopic.org/.

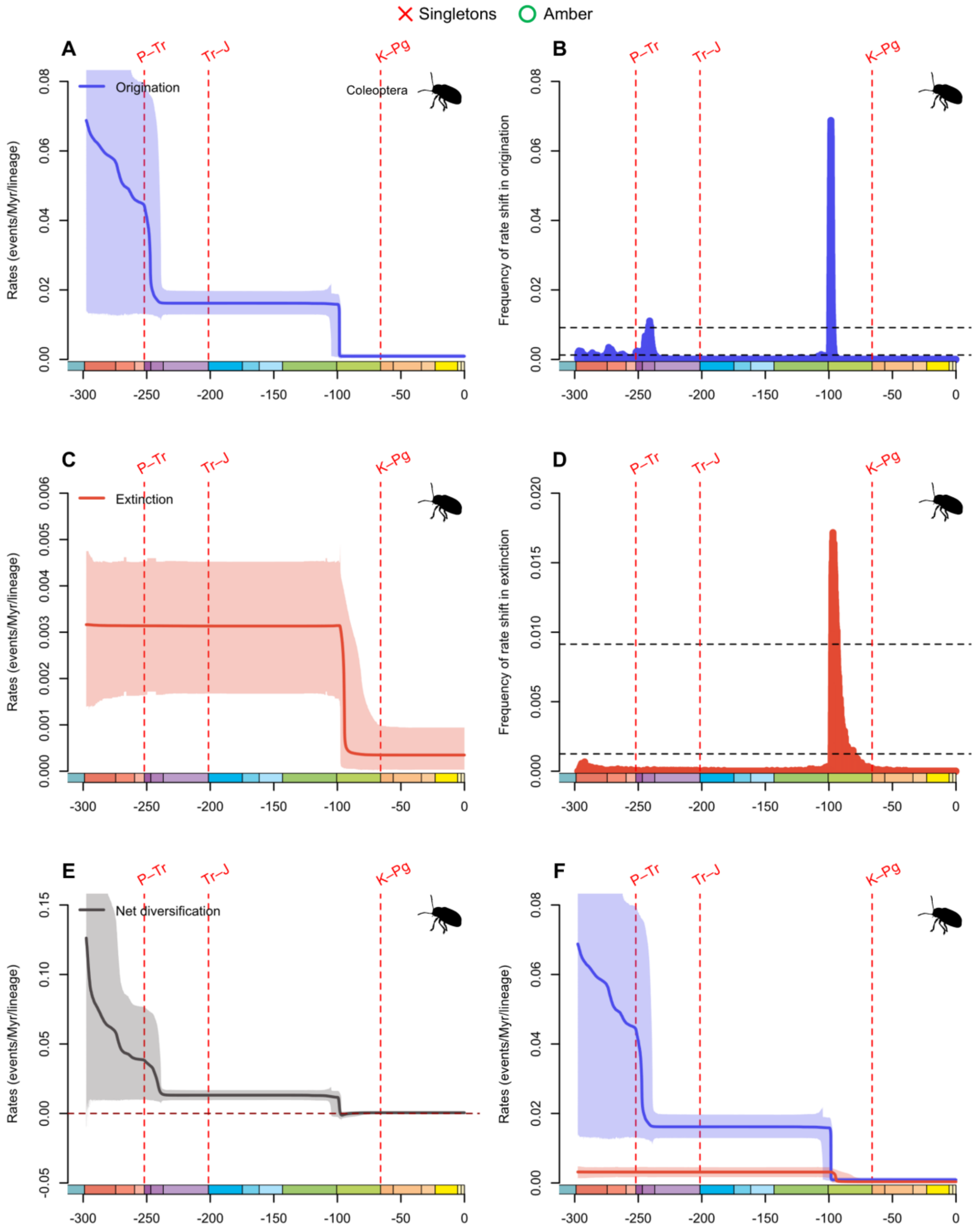


**fig. S19.** Diversification and diversity dynamics of Coleoptera families, excluding singletons, but considering amber occurrences. Bayesian estimations of origination and extinction rates through time as inferred by PyRate using reversible jump Markov Chain Monte Carlo (RJMCMC). Marginal estimates of origination rates (A) and extinction rates (C) through time are shown as mean and 95% HPD (shaded areas). The frequency of a sampled rate shift is computed within small time bins for origination and extinction rates (B and D, respectively), with black-dashed

horizontal lines indicating log-Bayes factors of 2 (bottom) and 6 (top). Sampling frequencies higher than log-Bayes factors = 6 indicate strong statistical support for a rate shift. Panel (E) shows net diversification rates through time (computed as the posterior difference between origination and extinction rates through time). Panel (F) shows both marginal estimates of origination and extinction rates through time on the same plot. Solid lines indicate mean posterior rates and the shaded areas show 95% HPD. Red-dashed vertical lines indicate major crises: P–Tr, Permian–Triassic; Tr–J, Triassic–Jurassic; K–Pg, Cretaceous–Paleogene. Time is in millions of years. The color of each geological period in the chronostratigraphic scale follows that of the International Chronostratigraphic Chart (v2024/12). Insect silhouettes are from http://phylopic.org/.

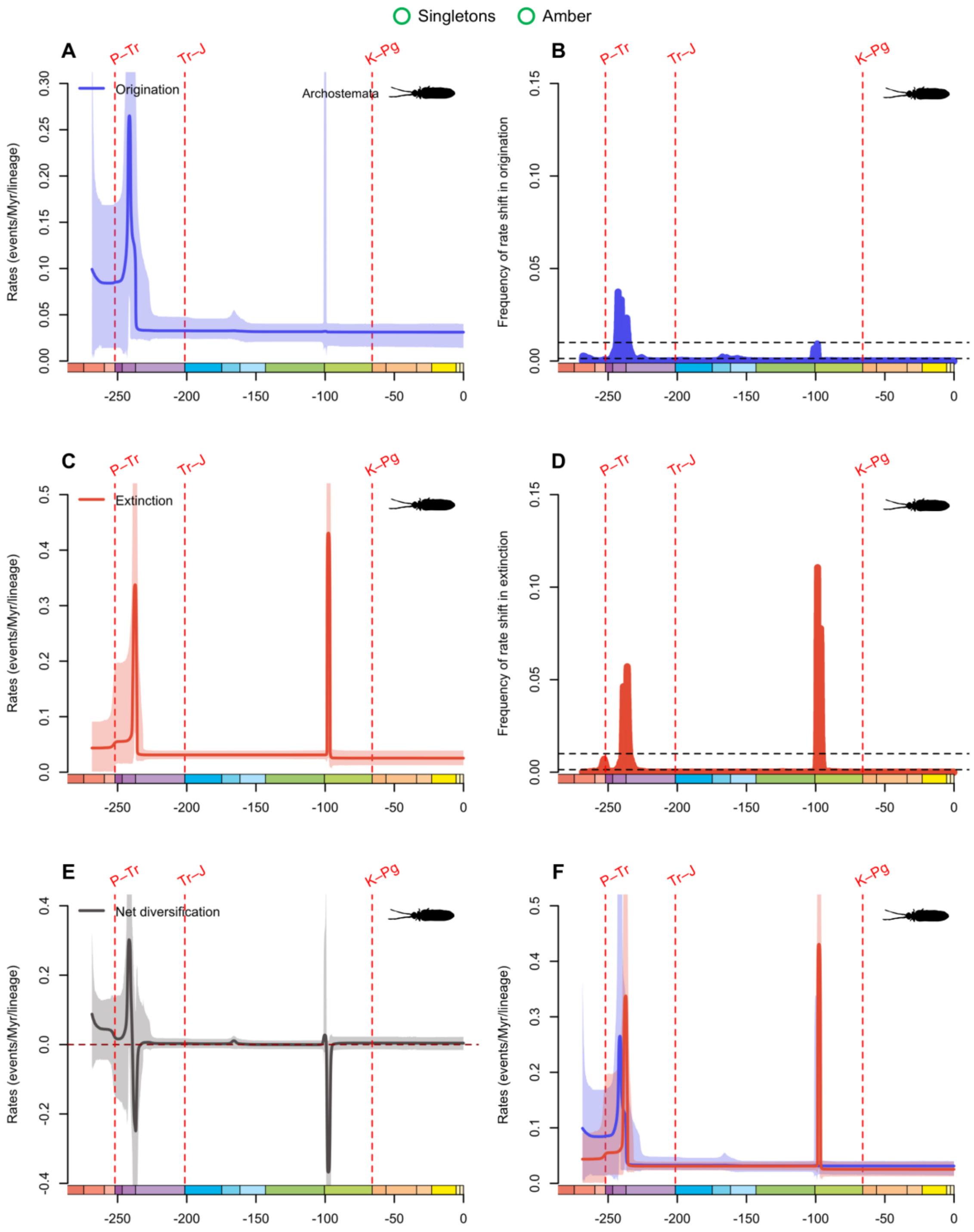


**fig. S20.** Diversification and diversity dynamics of Archostemata genera, considering singletons and amber occurrences. Bayesian estimations of origination and extinction rates through time as inferred by PyRate using reversible jump Markov Chain Monte Carlo (RJMCMC). Marginal estimates of origination rates (A) and extinction rates (C) through time are shown as mean and 95% HPD (shaded areas). The frequency of a sampled rate shift is computed within small time bins for origination and extinction rates (B and D, respectively), with black-dashed horizontal lines

indicating log-Bayes factors of 2 (bottom) and 6 (top). Sampling frequencies higher than log-Bayes factors = 6 indicate strong statistical support for a rate shift. Panel (E) shows net diversification rates through time (computed as the posterior difference between origination and extinction rates through time). Panel (F) shows both marginal estimates of origination and extinction rates through time on the same plot. Solid lines indicate mean posterior rates and the shaded areas show 95% HPD. Red-dashed vertical lines indicate major crises: P–Tr, Permian–Triassic; Tr–J, Triassic–Jurassic; K–Pg, Cretaceous–Paleogene. Time is in millions of years. The color of each geological period in the chronostratigraphic scale follows that of the International Chronostratigraphic Chart (v2024/12). Insect silhouettes are from http://phylopic.org/.

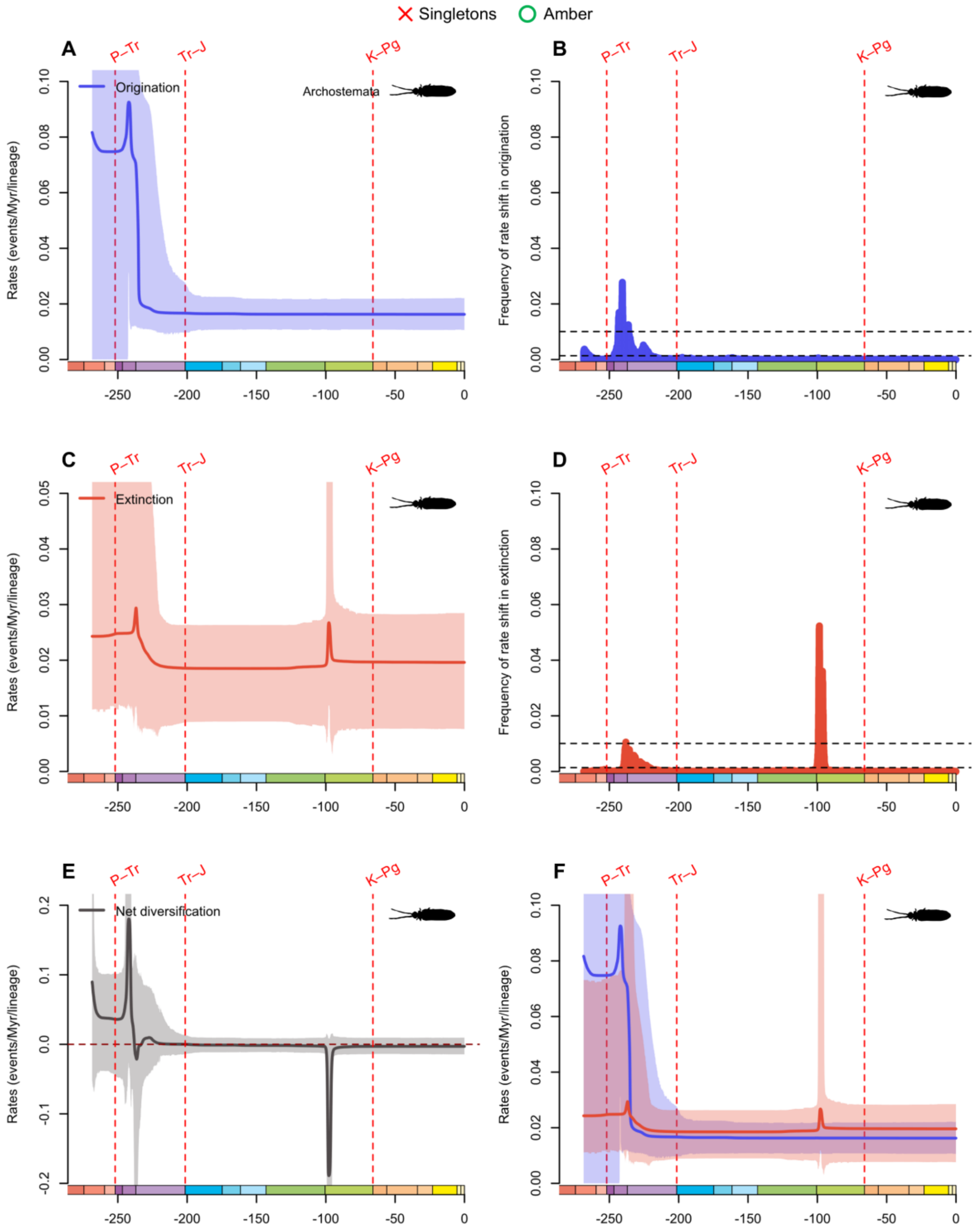


**fig. S21.** Diversification and diversity dynamics of Archostemata genera, excluding singletons, but considering amber occurrences. Bayesian estimations of origination and extinction rates through time as inferred by PyRate using reversible jump Markov Chain Monte Carlo (RJMCMC). Marginal estimates of origination rates (A) and extinction rates (C) through time are shown as mean and 95% HPD (shaded areas). The frequency of a sampled rate shift is computed within small time bins for origination and extinction rates (B and D, respectively), with black-dashed

horizontal lines indicating log-Bayes factors of 2 (bottom) and 6 (top). Sampling frequencies higher than log-Bayes factors = 6 indicate strong statistical support for a rate shift. Panel (E) shows net diversification rates through time (computed as the posterior difference between origination and extinction rates through time). Panel (F) shows both marginal estimates of origination and extinction rates through time on the same plot. Solid lines indicate mean posterior rates and the shaded areas show 95% HPD. Red-dashed vertical lines indicate major crises: P–Tr, Permian–Triassic; Tr–J, Triassic–Jurassic; K–Pg, Cretaceous–Paleogene. Time is in millions of years. The color of each geological period in the chronostratigraphic scale follows that of the International Chronostratigraphic Chart (v2024/12). Insect silhouettes are from http://phylopic.org/.

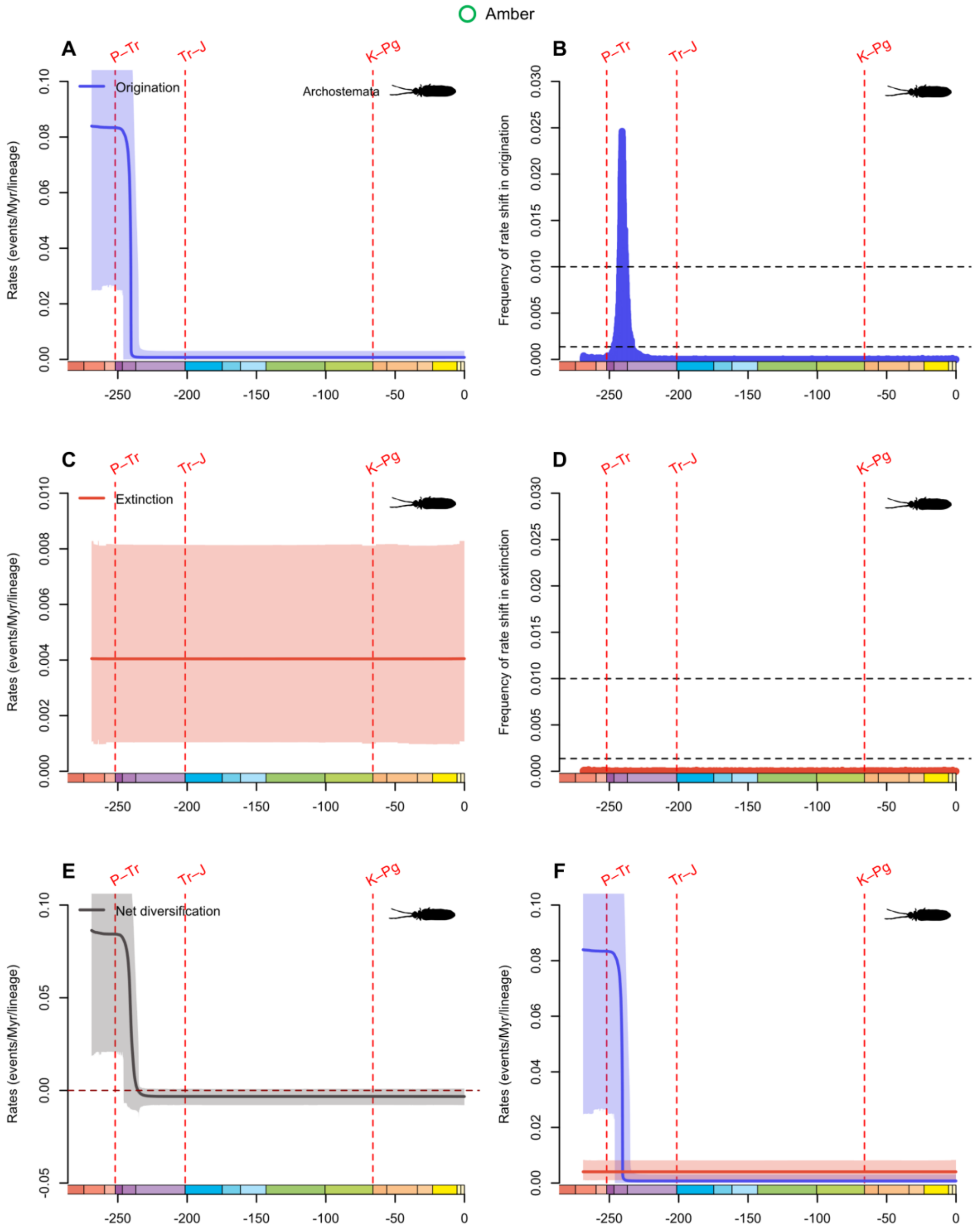


**fig. S22.** Diversification and diversity dynamics of Archostemata families, considering amber occurrences. There are no singletons in Archostemata at the family-level. Bayesian estimations of origination and extinction rates through time as inferred by PyRate using reversible jump Markov Chain Monte Carlo (RJMCMC). Marginal estimates of origination rates (A) and extinction rates (C) through time are shown as mean and 95% HPD (shaded areas). The frequency of a sampled rate shift is computed within small time bins for origination and extinction rates (B and D, respectively), with black-dashed horizontal lines indicating log-Bayes factors of 2 (bottom) and 6 (top). Sampling frequencies higher than log-Bayes factors = 6 indicate strong statistical support for a rate shift. Panel (E) shows net diversification rates through time (computed as the posterior difference between origination and extinction rates through time). Panel (F) shows both marginal estimates of origination and extinction rates through time on the same plot. Solid lines indicate mean posterior rates and the shaded areas show 95% HPD. Red-dashed vertical lines indicate major crises: P–Tr, Permian–Triassic; Tr–J, Triassic–Jurassic; K–Pg, Cretaceous–Paleogene. Time is in millions of years. The color of each geological period in the chronostratigraphic scale follows that of the International Chronostratigraphic Chart (v2024/12). Insect silhouettes are from http://phylopic.org/.

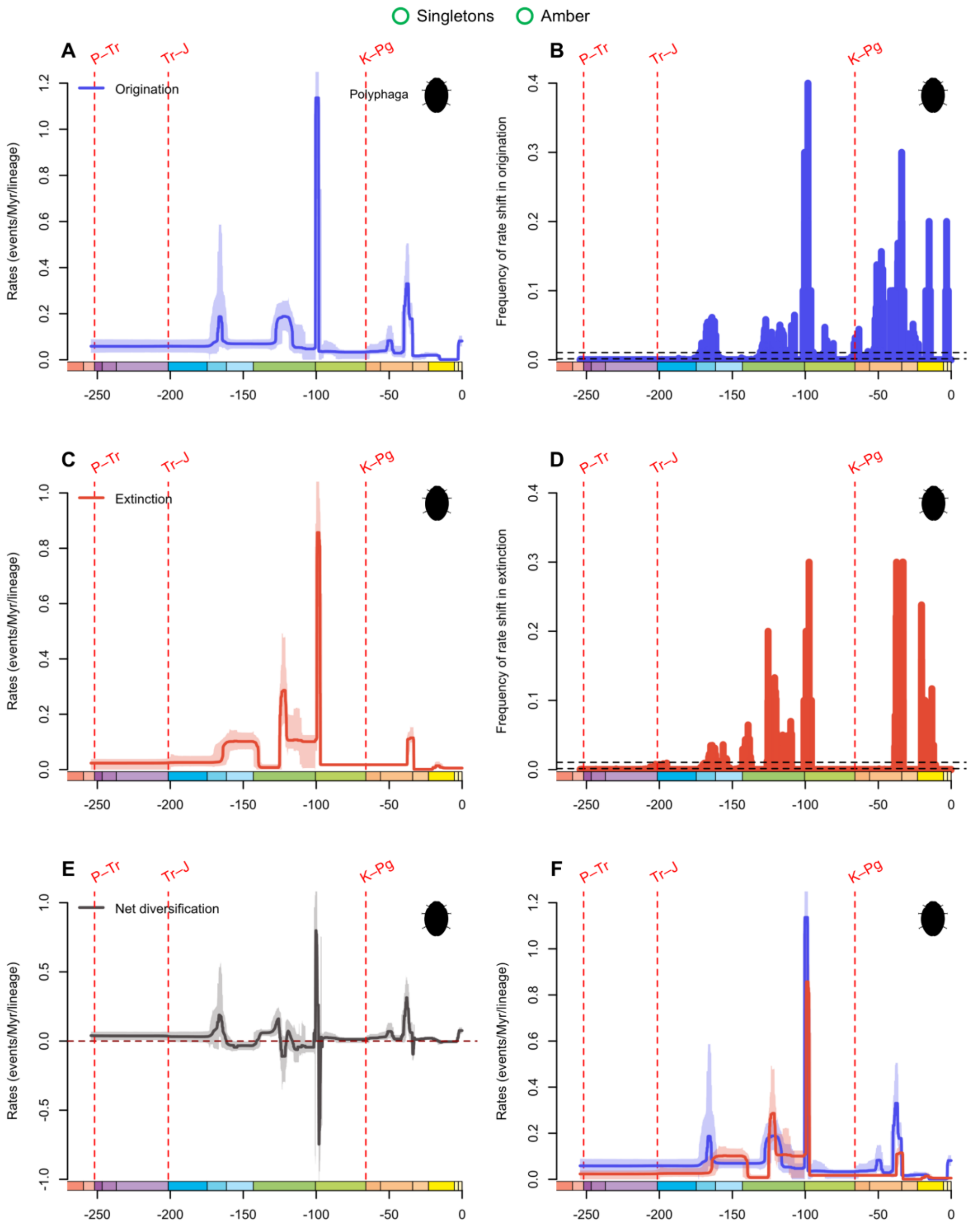


**fig. S23.** Diversification and diversity dynamics of Polyphaga genera, considering singletons and amber occurrences. Bayesian estimations of origination and extinction rates through time as inferred by PyRate using reversible jump Markov Chain Monte Carlo (RJMCMC). Marginal estimates of origination rates (A) and extinction rates (C) through time are shown as mean and 95% HPD (shaded areas). The frequency of a sampled rate shift is computed within small time bins for origination and extinction rates (B and D, respectively), with black-dashed horizontal lines

indicating log-Bayes factors of 2 (bottom) and 6 (top). Sampling frequencies higher than log-Bayes factors = 6 indicate strong statistical support for a rate shift. Panel (E) shows net diversification rates through time (computed as the posterior difference between origination and extinction rates through time). Panel (F) shows both marginal estimates of origination and extinction rates through time on the same plot. Solid lines indicate mean posterior rates and the shaded areas show 95% HPD. Red-dashed vertical lines indicate major crises: P–Tr, Permian–Triassic; Tr–J, Triassic–Jurassic; K–Pg, Cretaceous–Paleogene. Time is in millions of years. The color of each geological period in the chronostratigraphic scale follows that of the International Chronostratigraphic Chart (v2024/12). Insect silhouettes are from http://phylopic.org/.

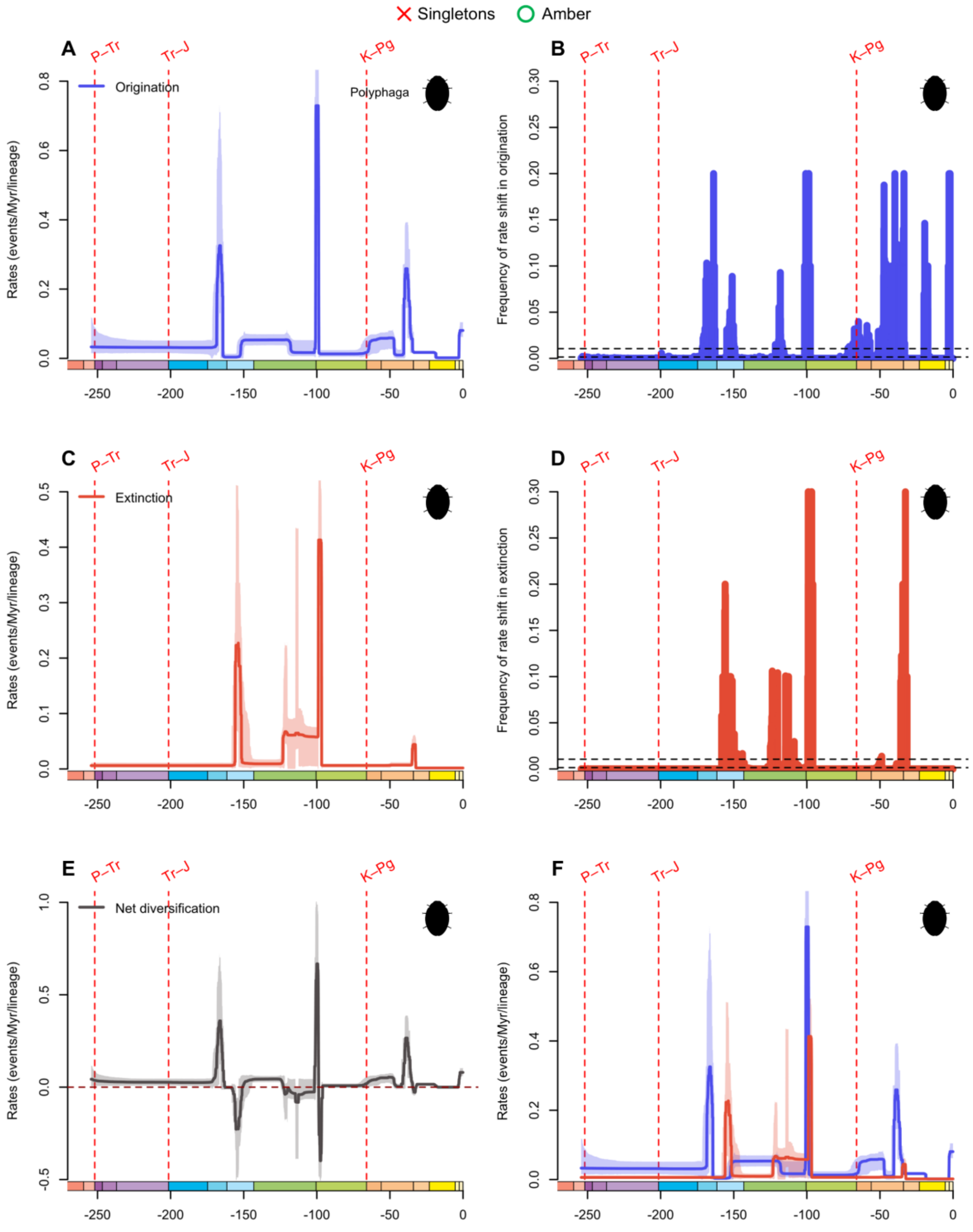


**fig. S24.** Diversification and diversity dynamics of Polyphaga genera, excluding singletons, but considering amber occurrences. Bayesian estimations of origination and extinction rates through time as inferred by PyRate using reversible jump Markov Chain Monte Carlo (RJMCMC). Marginal estimates of origination rates (A) and extinction rates (C) through time are shown as mean and 95% HPD (shaded areas). The frequency of a sampled rate shift is computed within small time bins for origination and extinction rates (B and D, respectively), with black-dashed

horizontal lines indicating log-Bayes factors of 2 (bottom) and 6 (top). Sampling frequencies higher than log-Bayes factors = 6 indicate strong statistical support for a rate shift. Panel (E) shows net diversification rates through time (computed as the posterior difference between origination and extinction rates through time). Panel (F) shows both marginal estimates of origination and extinction rates through time on the same plot. Solid lines indicate mean posterior rates and the shaded areas show 95% HPD. Red-dashed vertical lines indicate major crises: P–Tr, Permian–Triassic; Tr–J, Triassic–Jurassic; K–Pg, Cretaceous–Paleogene. Time is in millions of years. The color of each geological period in the chronostratigraphic scale follows that of the International Chronostratigraphic Chart (v2024/12). Insect silhouettes are from http://phylopic.org/.

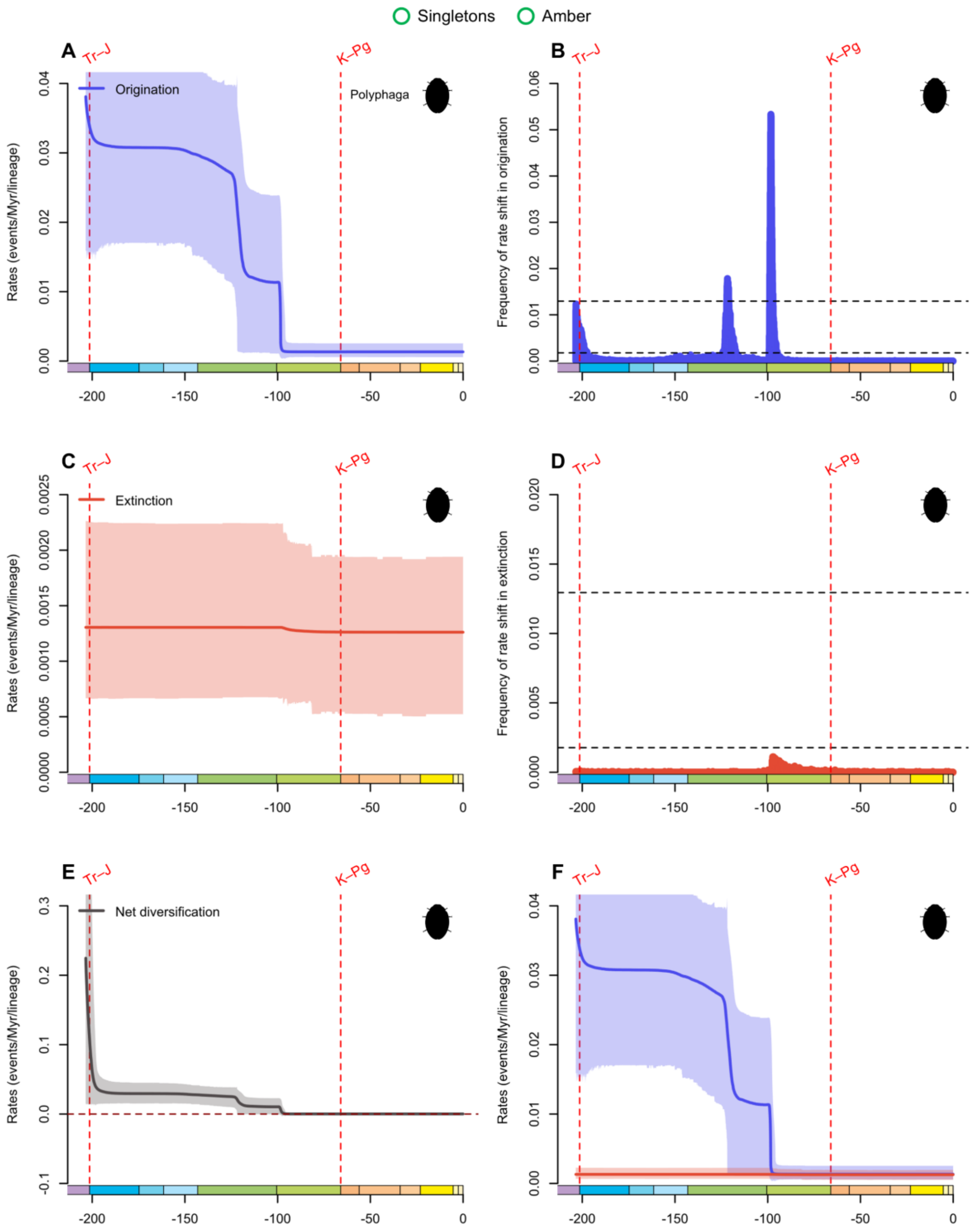


**fig. S25.** Diversification and diversity dynamics of Polyphaga families, considering singletons and amber occurrences. Bayesian estimations of origination and extinction rates through time as inferred by PyRate using reversible jump Markov Chain Monte Carlo (RJMCMC). Marginal estimates of origination rates (A) and extinction rates (C) through time are shown as mean and 95% HPD (shaded areas). The frequency of a sampled rate shift is computed within small time bins for origination and extinction rates (B and D, respectively), with black-dashed horizontal lines indicating log-Bayes factors of 2 (bottom) and 6 (top). Sampling frequencies higher than log-Bayes factors = 6 indicate strong statistical support for a rate shift. Panel (E) shows net diversification rates through time (computed as the posterior difference between origination and extinction rates through time). Panel (F) shows both marginal estimates of origination and extinction rates through time on the same plot. Solid lines indicate mean posterior rates and the shaded areas show 95% HPD. Red-dashed vertical lines indicate major crises: Tr–J, Triassic–Jurassic; K–Pg, Cretaceous–Paleogene. Time is in millions of years. The color of each geological period in the chronostratigraphic scale follows that of the International Chronostratigraphic Chart (v2024/12). Insect silhouettes are from http://phylopic.org/.

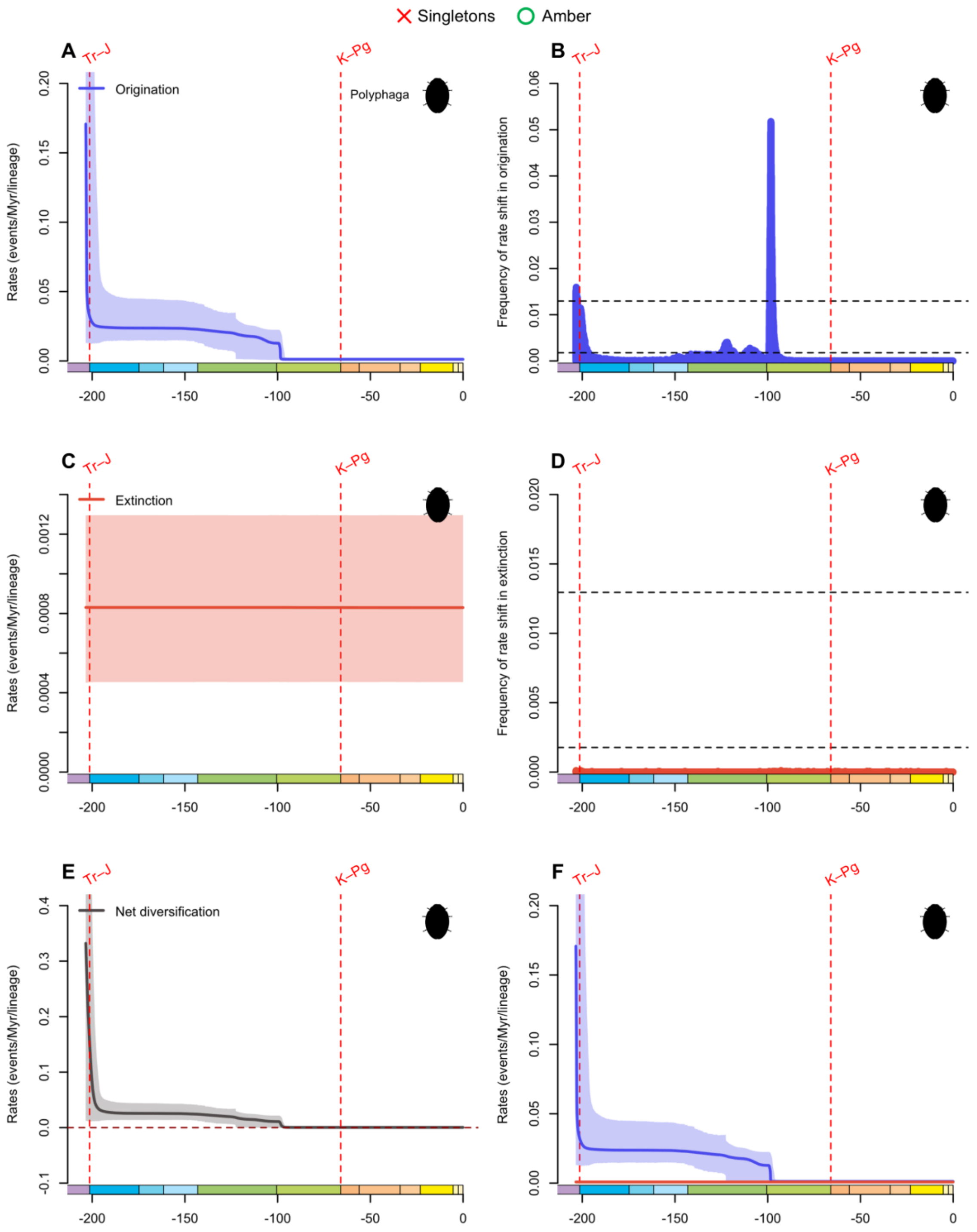


**fig. S26.** Diversification and diversity dynamics of Polyphaga families, excluding singletons, but considering amber occurrences. Bayesian estimations of origination and extinction rates through time as inferred by PyRate using reversible jump Markov Chain Monte Carlo (RJMCMC). Marginal estimates of origination rates (A) and extinction rates (C) through time are shown as mean and 95% HPD (shaded areas). The frequency of a sampled rate shift is computed within small time bins for origination and extinction rates (B and D, respectively), with black-dashed horizontal lines indicating log-Bayes factors of 2 (bottom) and 6 (top). Sampling frequencies higher than log-Bayes factors = 6 indicate strong statistical support for a rate shift. Panel (E) shows net diversification rates through time (computed as the posterior difference between origination and extinction rates through time). Panel (F) shows both marginal estimates of origination and extinction rates through time on the same plot. Solid lines indicate mean posterior rates and the shaded areas show 95% HPD. Red-dashed vertical lines indicate major crises: Tr–J, Triassic–Jurassic; K–Pg, Cretaceous–Paleogene. Time is in millions of years. The color of each geological period in the chronostratigraphic scale follows that of the International Chronostratigraphic Chart (v2024/12). Insect silhouettes are from http://phylopic.org/.

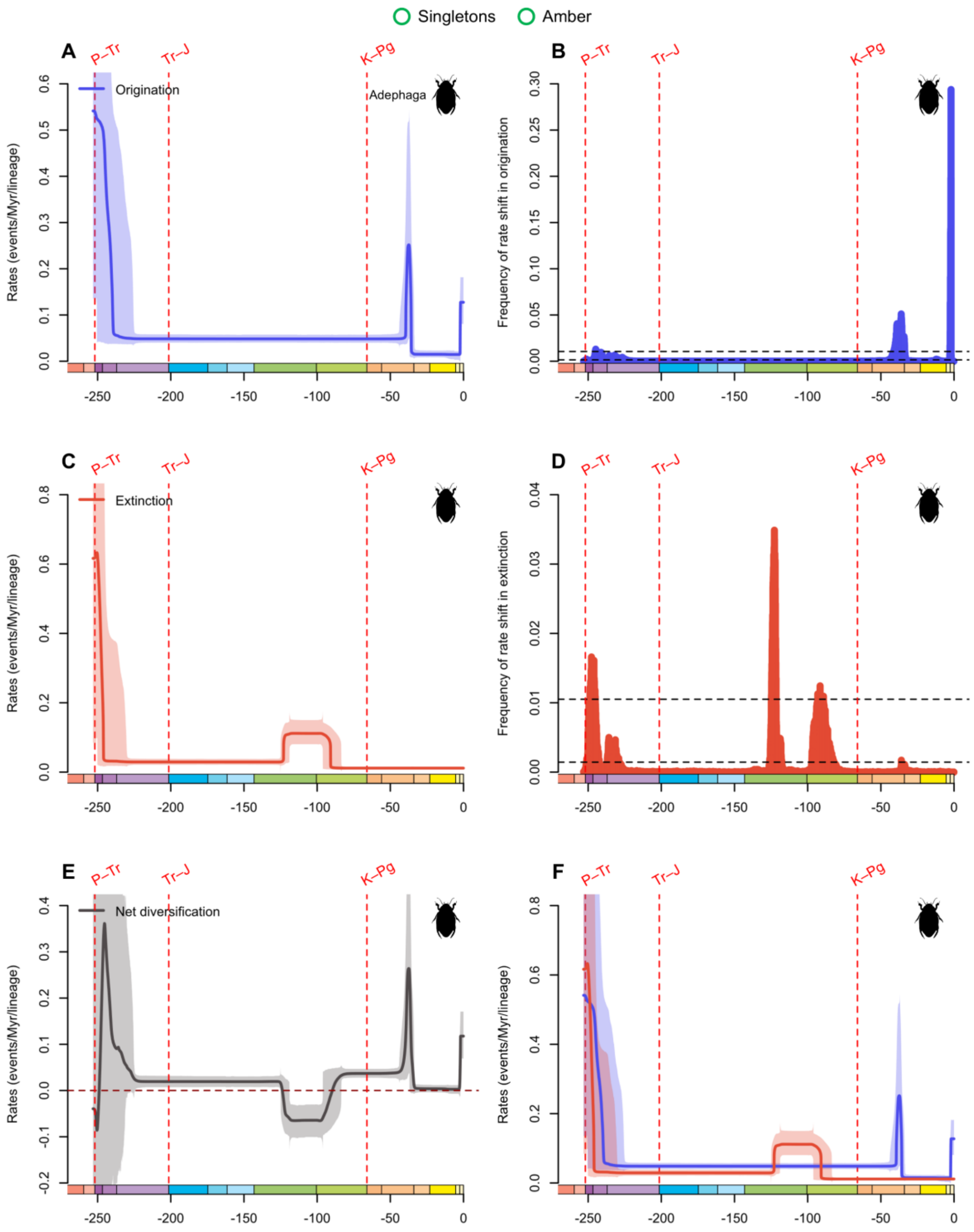


**fig. S27.** Diversification and diversity dynamics of Adephaga genera, considering singletons and amber occurrences. Bayesian estimations of origination and extinction rates through time as inferred by PyRate using reversible jump Markov Chain Monte Carlo (RJMCMC). Marginal estimates of origination rates (A) and extinction rates (C) through time are shown as mean and 95% HPD (shaded areas). The frequency of a sampled rate shift is computed within small time bins for origination and extinction rates (B and D, respectively), with black-dashed horizontal lines

indicating log-Bayes factors of 2 (bottom) and 6 (top). Sampling frequencies higher than log-Bayes factors = 6 indicate strong statistical support for a rate shift. Panel (E) shows net diversification rates through time (computed as the posterior difference between origination and extinction rates through time). Panel (F) shows both marginal estimates of origination and extinction rates through time on the same plot. Solid lines indicate mean posterior rates and the shaded areas show 95% HPD. Red-dashed vertical lines indicate major crises: P–Tr, Permian–Triassic; Tr–J, Triassic–Jurassic; K–Pg, Cretaceous–Paleogene. Time is in millions of years. The color of each geological period in the chronostratigraphic scale follows that of the International Chronostratigraphic Chart (v2024/12). Insect silhouettes are from http://phylopic.org/.

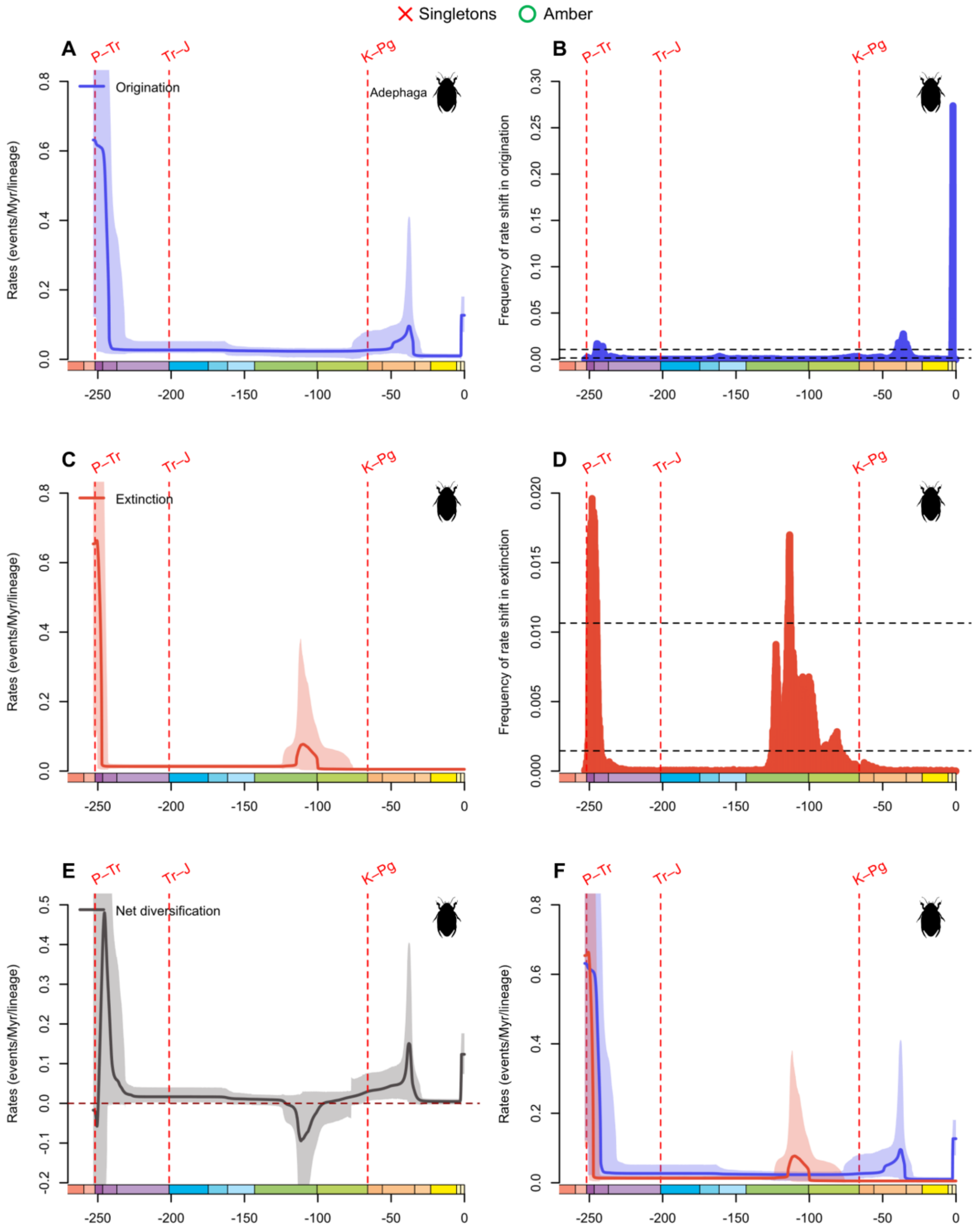


**fig. S28.** Diversification and diversity dynamics of Adephaga genera, excluding singletons, but considering amber occurrences. Bayesian estimations of origination and extinction rates through time as inferred by PyRate using reversible jump Markov Chain Monte Carlo (RJMCMC). Marginal estimates of origination rates (A) and extinction rates (C) through time are shown as mean and 95% HPD (shaded areas). The frequency of a sampled rate shift is computed within small time bins for origination and extinction rates (B and D, respectively), with black-dashed

horizontal lines indicating log-Bayes factors of 2 (bottom) and 6 (top). Sampling frequencies higher than log-Bayes factors = 6 indicate strong statistical support for a rate shift. Panel (E) shows net diversification rates through time (computed as the posterior difference between origination and extinction rates through time). Panel (F) shows both marginal estimates of origination and extinction rates through time on the same plot. Solid lines indicate mean posterior rates and the shaded areas show 95% HPD. Red-dashed vertical lines indicate major crises: P–Tr, Permian–Triassic; Tr–J, Triassic–Jurassic; K–Pg, Cretaceous–Paleogene. Time is in millions of years. The color of each geological period in the chronostratigraphic scale follows that of the International Chronostratigraphic Chart (v2024/12). Insect silhouettes are from http://phylopic.org/.

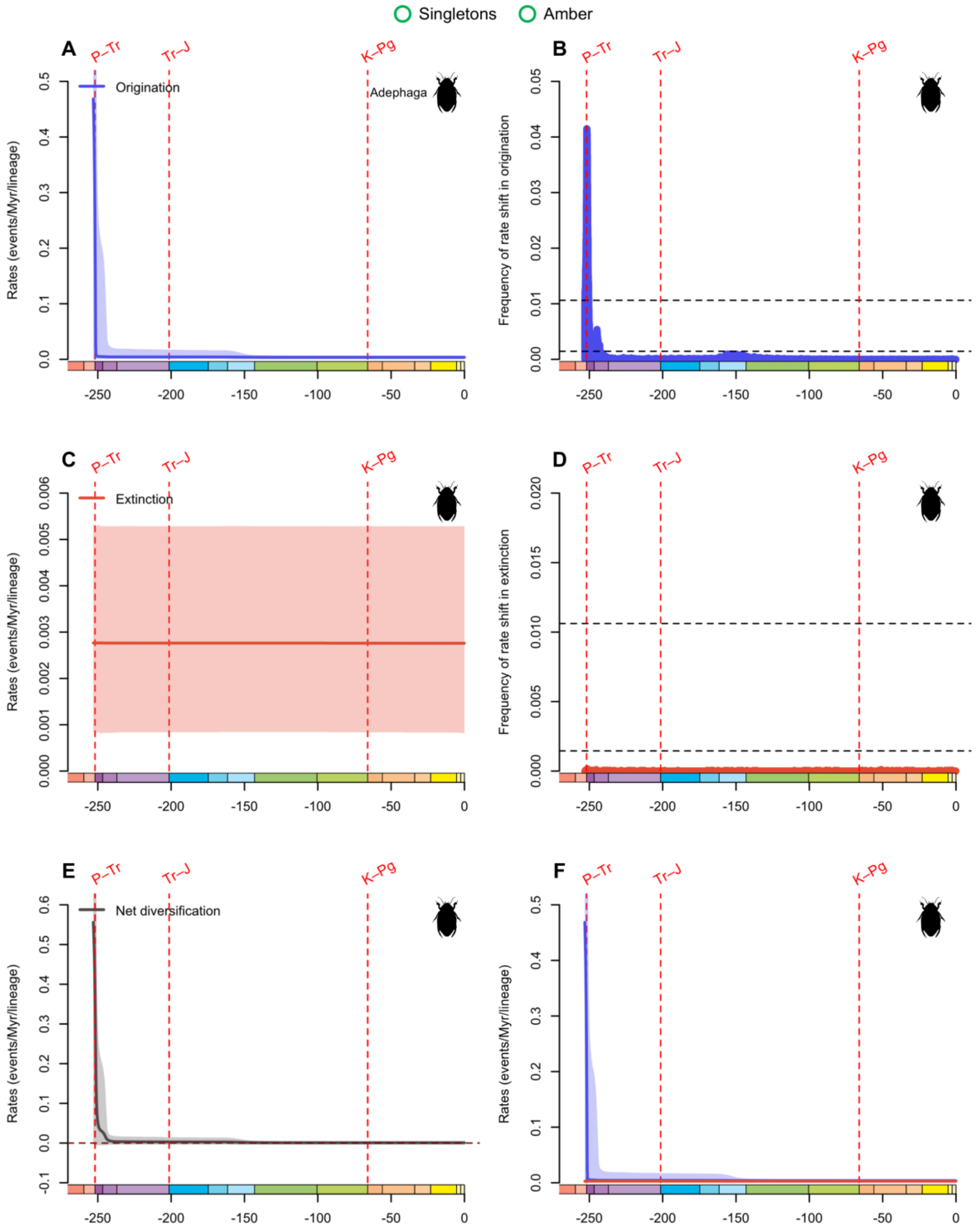


**fig. S29.** Diversification and diversity dynamics of Adephaga families, considering singletons and amber occurrences. Bayesian estimations of origination and extinction rates through time as inferred by PyRate using reversible jump Markov Chain Monte Carlo (RJMCMC). Marginal estimates of origination rates (A) and extinction rates (C) through time are shown as mean and 95% HPD (shaded areas). The frequency of a sampled rate shift is computed within small time bins for origination and extinction rates (B and D, respectively), with black-dashed horizontal lines

indicating log-Bayes factors of 2 (bottom) and 6 (top). Sampling frequencies higher than log-Bayes factors = 6 indicate strong statistical support for a rate shift. Panel (E) shows net diversification rates through time (computed as the posterior difference between origination and extinction rates through time). Panel (F) shows both marginal estimates of origination and extinction rates through time on the same plot. Solid lines indicate mean posterior rates and the shaded areas show 95% HPD. Red-dashed vertical lines indicate major crises: P–Tr, Permian–Triassic; Tr–J, Triassic–Jurassic; K–Pg, Cretaceous–Paleogene. Time is in millions of years. The color of each geological period in the chronostratigraphic scale follows that of the International Chronostratigraphic Chart (v2024/12). Insect silhouettes are from http://phylopic.org/.

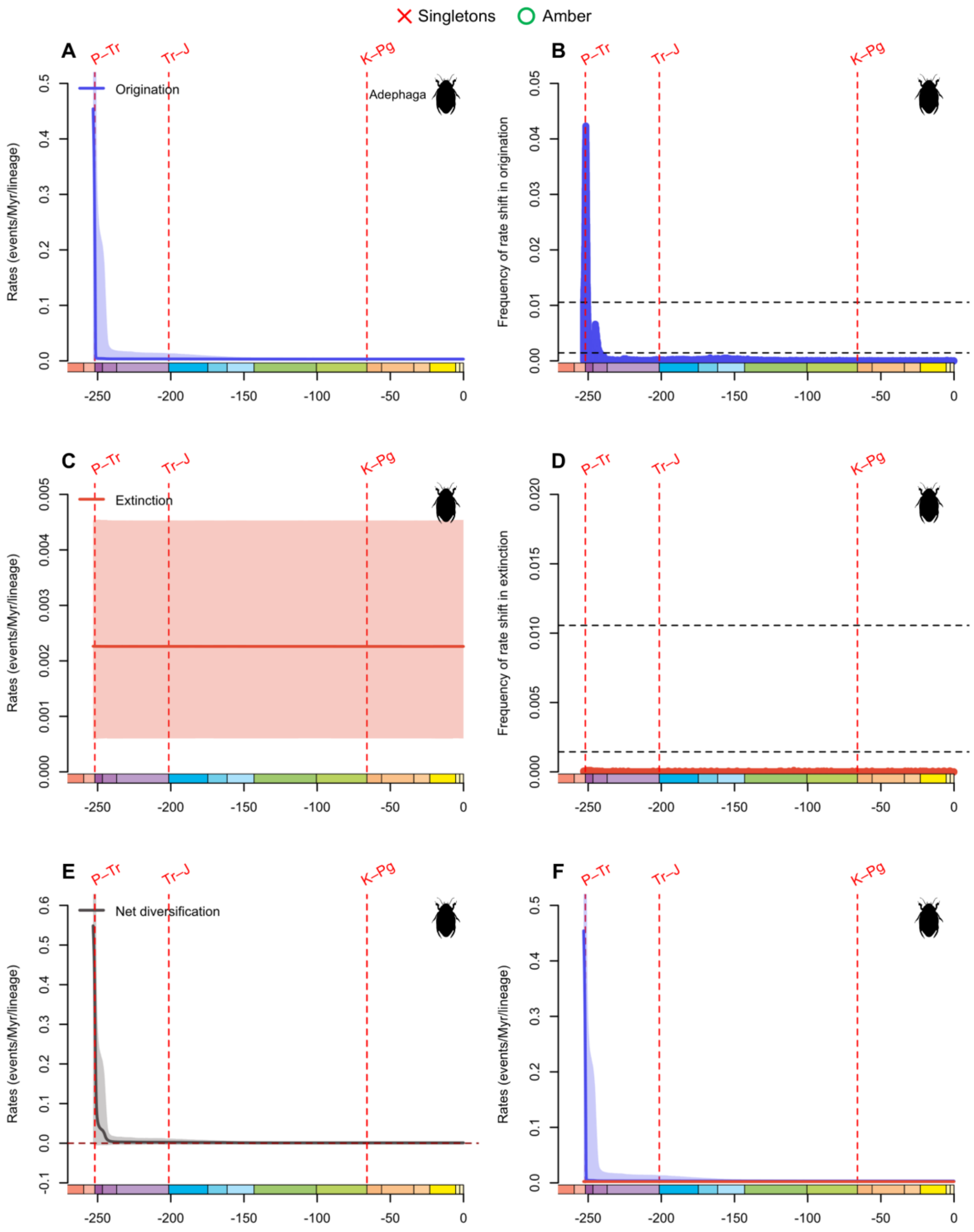


**fig. S30.** Diversification and diversity dynamics of Adephaga families, excluding singletons, but considering amber occurrences. Bayesian estimations of origination and extinction rates through time as inferred by PyRate using reversible jump Markov Chain Monte Carlo (RJMCMC). Marginal estimates of origination rates (A) and extinction rates (C) through time are shown as mean and 95% HPD (shaded areas). The frequency of a sampled rate shift is computed within small time bins for origination and extinction rates (B and D, respectively), with black-dashed

horizontal lines indicating log-Bayes factors of 2 (bottom) and 6 (top). Sampling frequencies higher than log-Bayes factors = 6 indicate strong statistical support for a rate shift. Panel (E) shows net diversification rates through time (computed as the posterior difference between origination and extinction rates through time). Panel (F) shows both marginal estimates of origination and extinction rates through time on the same plot. Solid lines indicate mean posterior rates and the shaded areas show 95% HPD. Red-dashed vertical lines indicate major crises: P–Tr, Permian–Triassic; Tr–J, Triassic–Jurassic; K–Pg, Cretaceous–Paleogene. Time is in millions of years. The color of each geological period in the chronostratigraphic scale follows that of the International Chronostratigraphic Chart (v2024/12). Insect silhouettes are from http://phylopic.org/.

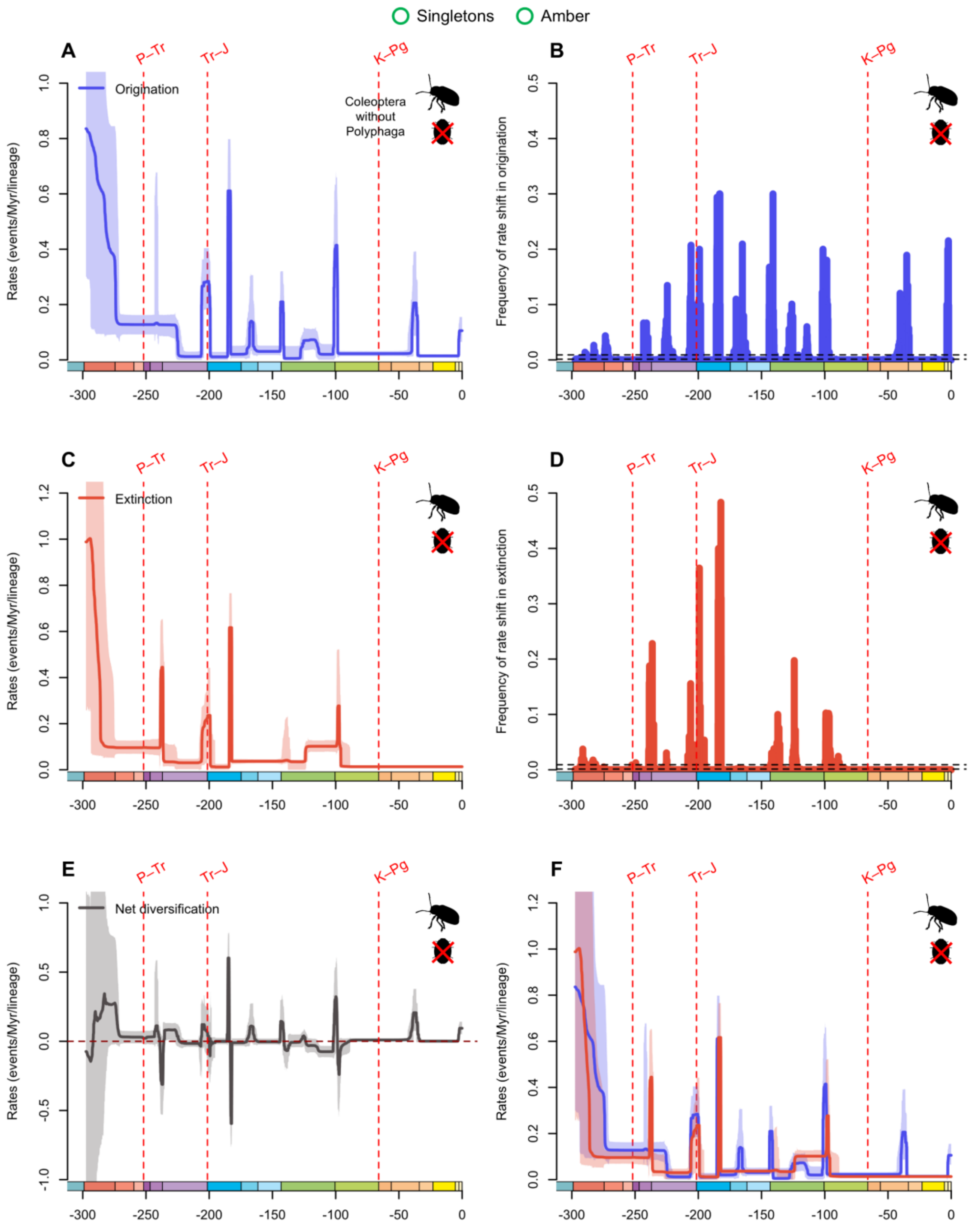


**fig. S31.** Diversification and diversity dynamics of Coleoptera genera without Polyphaga, considering singletons and amber occurrences. Bayesian estimations of origination and extinction rates through time as inferred by PyRate using reversible jump Markov Chain Monte Carlo (RJMCMC). Marginal estimates of origination rates (A) and extinction rates (C) through time are shown as mean and 95% HPD (shaded areas). The frequency of a sampled rate shift is computed within small time bins for origination and extinction rates (B and D, respectively), with black-

dashed horizontal lines indicating log-Bayes factors of 2 (bottom) and 6 (top). Sampling frequencies higher than log-Bayes factors = 6 indicate strong statistical support for a rate shift. Panel (E) shows net diversification rates through time (computed as the posterior difference between origination and extinction rates through time). Panel (F) shows both marginal estimates of origination and extinction rates through time on the same plot. Solid lines indicate mean posterior rates and the shaded areas show 95% HPD. Red-dashed vertical lines indicate major crises: P–Tr, Permian–Triassic; Tr–J, Triassic–Jurassic; K–Pg, Cretaceous–Paleogene. Time is in millions of years. The color of each geological period in the chronostratigraphic scale follows that of the International Chronostratigraphic Chart (v2024/12). Insect silhouettes are from http://phylopic.org/.

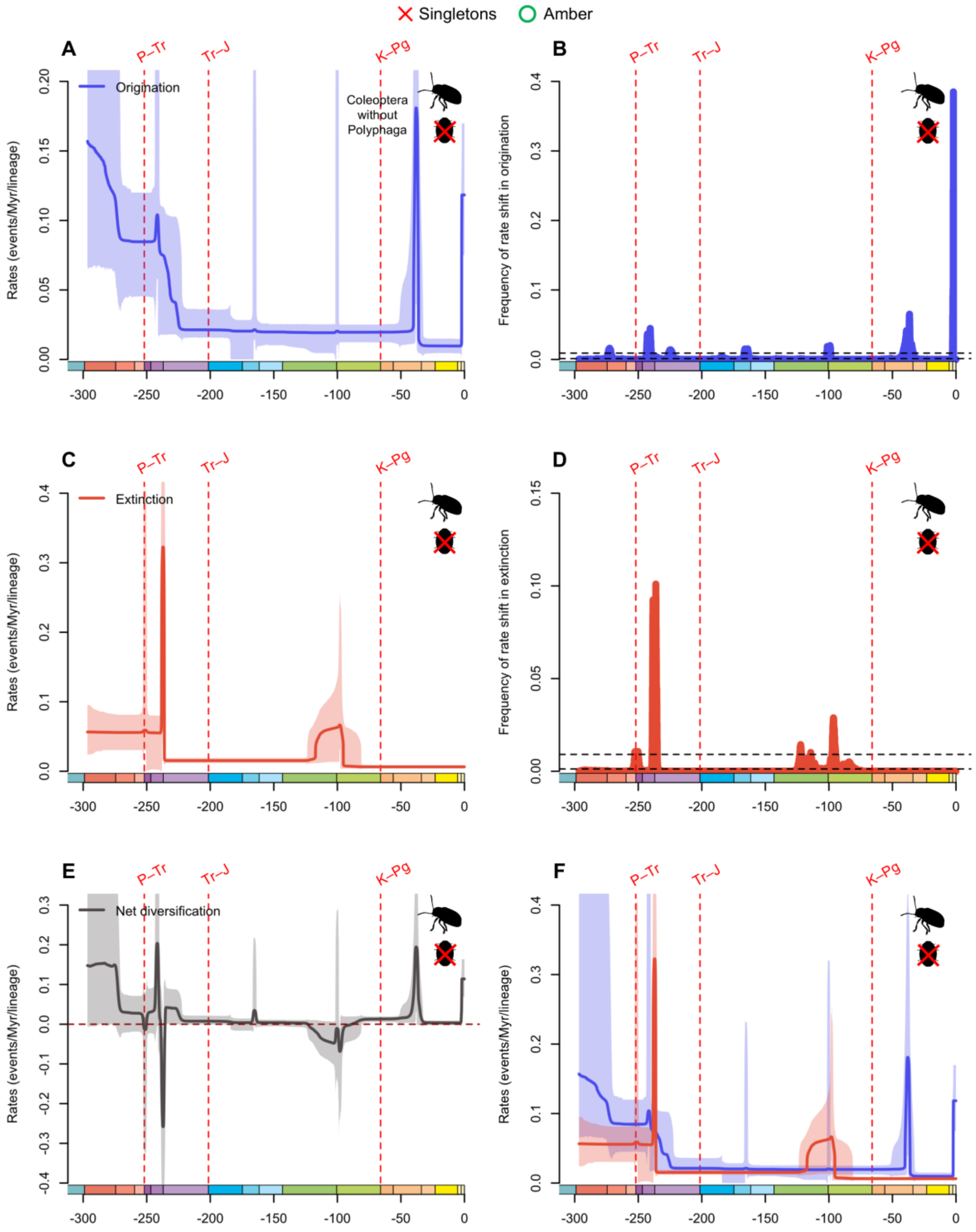


**fig. S32.** Diversification and diversity dynamics of Coleoptera genera without Polyphaga, excluding singletons, but considering amber occurrences. Bayesian estimations of origination and extinction rates through time as inferred by PyRate using reversible jump Markov Chain Monte Carlo (RJMCMC). Marginal estimates of origination rates (A) and extinction rates (C) through time are shown as mean and 95% HPD (shaded areas). The frequency of a sampled rate shift is computed within small time bins for origination and extinction rates (B and D, respectively), with

black-dashed horizontal lines indicating log-Bayes factors of 2 (bottom) and 6 (top). Sampling frequencies higher than log-Bayes factors = 6 indicate strong statistical support for a rate shift. Panel (E) shows net diversification rates through time (computed as the posterior difference between origination and extinction rates through time). Panel (F) shows both marginal estimates of origination and extinction rates through time on the same plot. Solid lines indicate mean posterior rates and the shaded areas show 95% HPD. Red-dashed vertical lines indicate major crises: P–Tr, Permian–Triassic; Tr–J, Triassic–Jurassic; K–Pg, Cretaceous–Paleogene. Time is in millions of years. The color of each geological period in the chronostratigraphic scale follows that of the International Chronostratigraphic Chart (v2024/12). Insect silhouettes are from http://phylopic.org/.

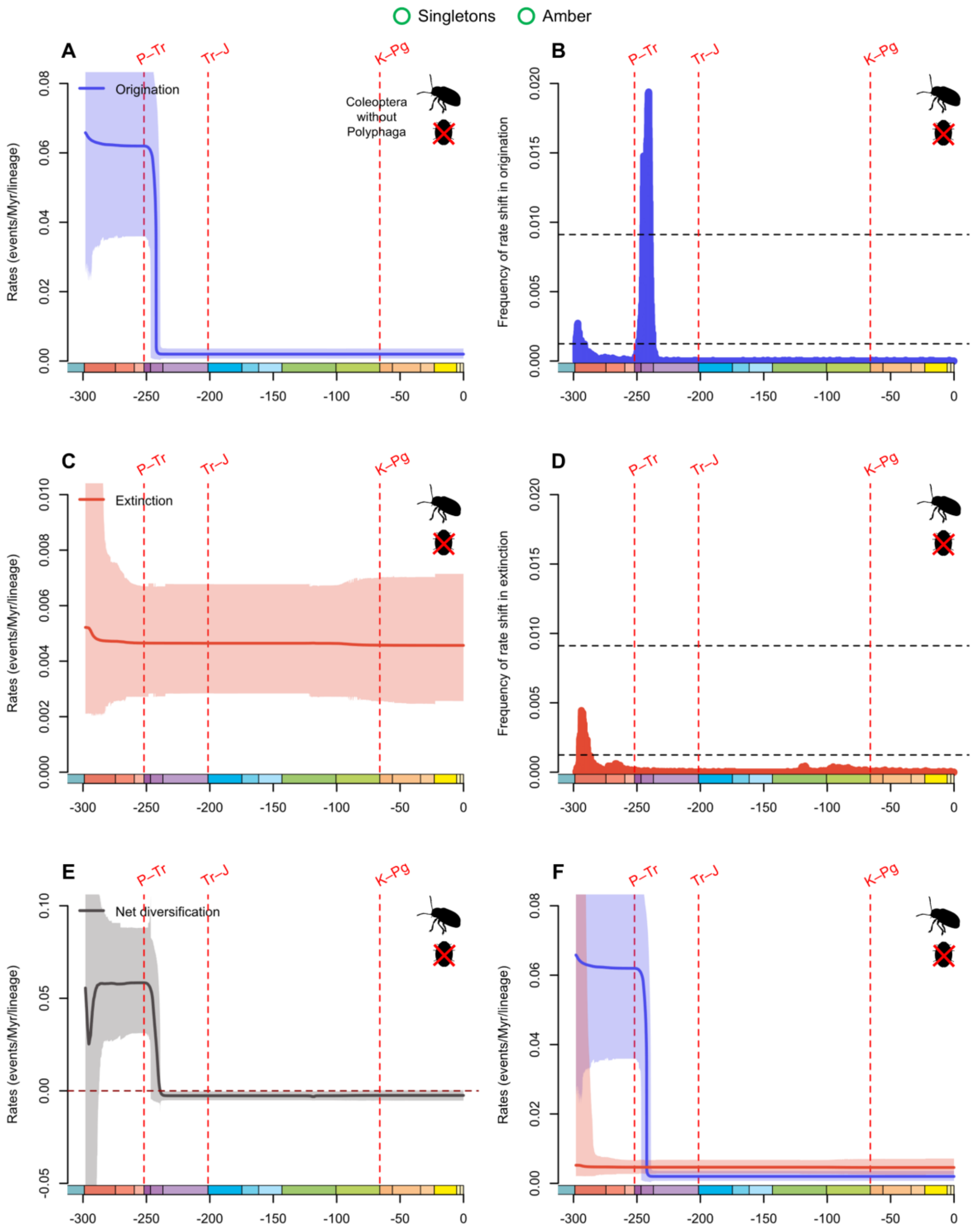


**fig. S33.** Diversification and diversity dynamics of Coleoptera families without Polyphaga, considering singletons and amber occurrences. Bayesian estimations of origination and extinction rates through time as inferred by PyRate using reversible jump Markov Chain Monte Carlo (RJMCMC). Marginal estimates of origination rates (A) and extinction rates (C) through time are shown as mean and 95% HPD (shaded areas). The frequency of a sampled rate shift is computed within small time bins for origination and extinction rates (B and D, respectively), with black-

dashed horizontal lines indicating log-Bayes factors of 2 (bottom) and 6 (top). Sampling frequencies higher than log-Bayes factors = 6 indicate strong statistical support for a rate shift. Panel (E) shows net diversification rates through time (computed as the posterior difference between origination and extinction rates through time). Panel (F) shows both marginal estimates of origination and extinction rates through time on the same plot. Solid lines indicate mean posterior rates and the shaded areas show 95% HPD. Red-dashed vertical lines indicate major crises: P–Tr, Permian–Triassic; Tr–J, Triassic–Jurassic; K–Pg, Cretaceous–Paleogene. Time is in millions of years. The color of each geological period in the chronostratigraphic scale follows that of the International Chronostratigraphic Chart (v2024/12). Insect silhouettes are from http://phylopic.org/.

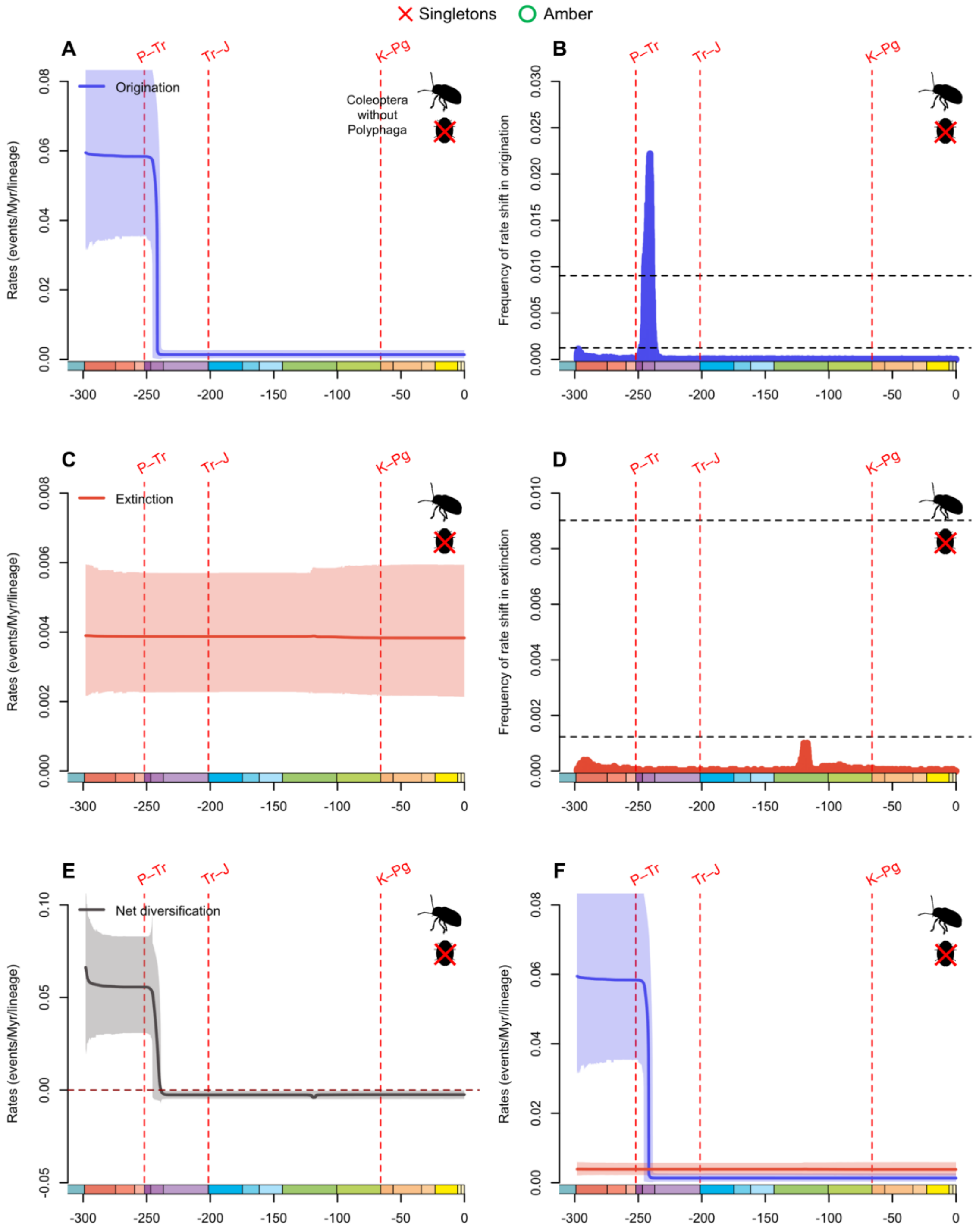


**fig. S34.** Diversification and diversity dynamics of Coleoptera families without Polyphaga, excluding singletons, but considering amber occurrences. Bayesian estimations of origination and extinction rates through time as inferred by PyRate using reversible jump Markov Chain Monte Carlo (RJMCMC). Marginal estimates of origination rates (A) and extinction rates (C) through time are shown as mean and 95% HPD (shaded areas). The frequency of a sampled rate shift is computed within small time bins for origination and extinction rates (B and D, respectively), with

black-dashed horizontal lines indicating log-Bayes factors of 2 (bottom) and 6 (top). Sampling frequencies higher than log-Bayes factors = 6 indicate strong statistical support for a rate shift. Panel (E) shows net diversification rates through time (computed as the posterior difference between origination and extinction rates through time). Panel (F) shows both marginal estimates of origination and extinction rates through time on the same plot. Solid lines indicate mean posterior rates and the shaded areas show 95% HPD. Red-dashed vertical lines indicate major crises: P–Tr, Permian–Triassic; Tr–J, Triassic–Jurassic; K–Pg, Cretaceous–Paleogene. Time is in millions of years. The color of each geological period in the chronostratigraphic scale follows that of the International Chronostratigraphic Chart (v2024/12). Insect silhouettes are from http://phylopic.org/.

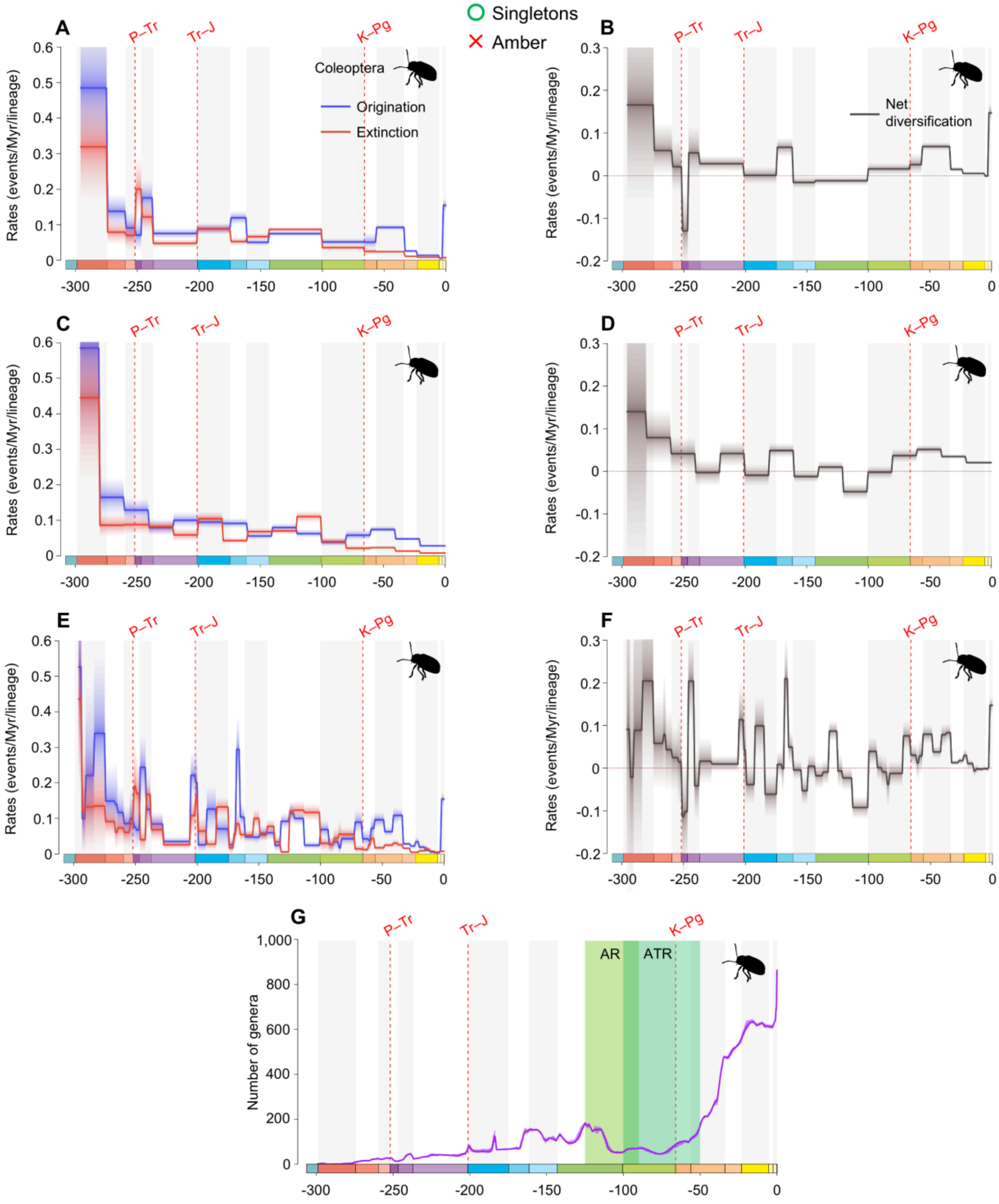


**fig. S35.** Diversification and diversity dynamics of Coleoptera genera, considering singletons and excluding amber occurrences. Bayesian fossil-based inferences of Coleoptera origination and extinction rates at the genus level under the birth-death model with epochs (A), 20 Ma bins (C), and stages (E) as constrained shifts (BDCS). Net diversification rates for Coleoptera obtained from the difference between origination and extinction rates (rates above 0 indicate increasing diversity, and rates below 0 indicate declining diversity) per epochs (B), 20 Ma bins (D), and stages (F). Solid lines indicate mean posterior rates and the shaded areas show 95% HPD. Number of genera through time computed by summing up the lifespans of all genera for Coleoptera (G). Solid lines indicate mean diversity at each point in time and shaded areas show estimations of different replications that incorporate age uncertainties of fossil occurrences. Light-green area represents the AR, angiosperm radiation, and dark-green area represents the ATR, angiosperm terrestrial revolution. Red-dashed vertical lines indicate major crises: P–Tr, Permian–Triassic; Tr–J, Triassic–Jurassic; K–Pg, Cretaceous–Paleogene. Time is in millions of years. The color of each geological period in the chronostratigraphic scale follows that of the International Chronostratigraphic Chart (v2024/12). Insect silhouettes are from http://phylopic.org/.

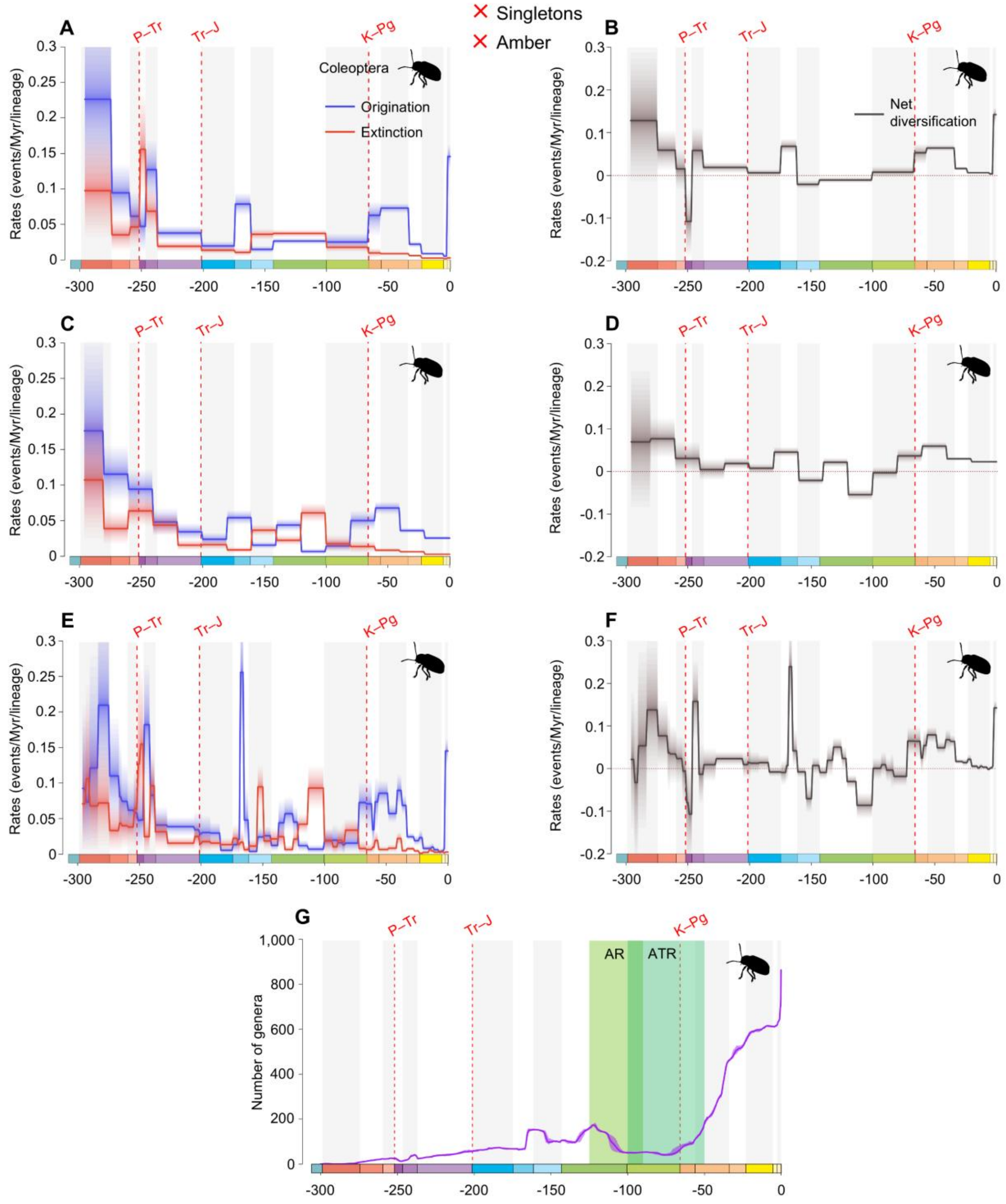


**fig. S36.** Diversification and diversity dynamics of Coleoptera genera, excluding singletons and amber occurrences. Bayesian fossil-based inferences of Coleoptera origination and extinction rates at the genus level under the birth-death model with epochs (A), 20 Ma bins (C), and stages (E) as constrained shifts (BDCS). Net diversification rates for Coleoptera obtained from the difference between origination and extinction rates (rates above 0 indicate increasing diversity, and rates below 0 indicate declining diversity) per epochs (B), 20 Ma bins (D), and stages (F). Solid lines indicate mean posterior rates and the shaded areas show 95% HPD. Number of genera through time computed by summing up the lifespans of all genera for Coleoptera (G). Solid lines indicate mean diversity at each point in time and shaded areas show estimations of different replications that incorporate age uncertainties of fossil occurrences. Light-green area represents the AR, angiosperm radiation, and dark-green area represents the ATR, angiosperm terrestrial revolution. Red-dashed vertical lines indicate major crises: P–Tr, Permian–Triassic; Tr–J, Triassic–Jurassic; K–Pg, Cretaceous–Paleogene. Time is in millions of years. The color of each geological period in the chronostratigraphic scale follows that of the International Chronostratigraphic Chart (v2024/12). Insect silhouettes are from http://phylopic.org/.

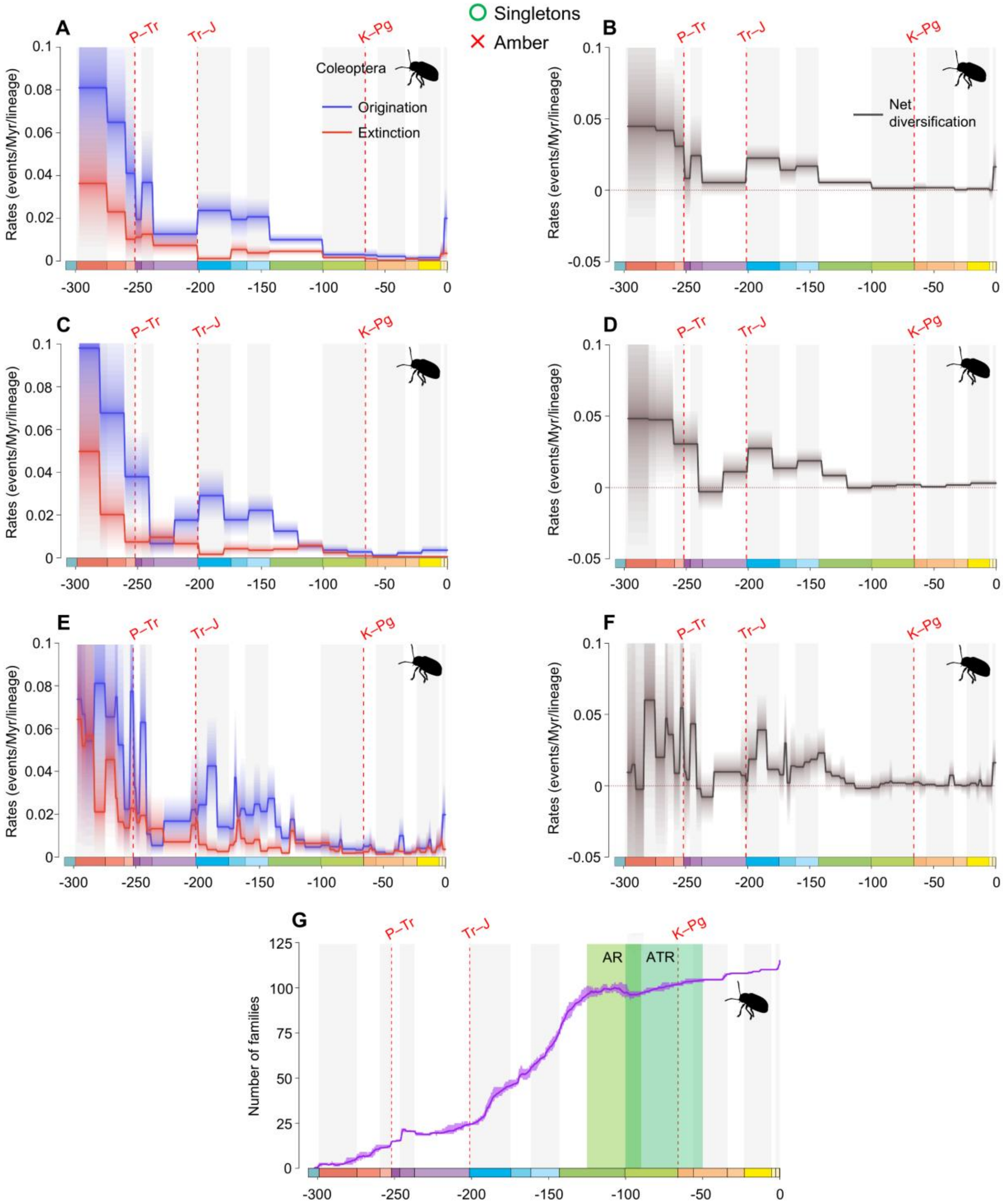


**fig. S37.** Diversification and diversity dynamics of Coleoptera families, considering singletons and excluding amber occurrences. Bayesian fossil-based inferences of Coleoptera origination and extinction rates at the family level under the birth-death model with epochs (A), 20 Ma bins (C), and stages (E) as constrained shifts (BDCS). Net diversification rates for Coleoptera obtained from the difference between origination and extinction rates (rates above 0 indicate increasing diversity, and rates below 0 indicate declining diversity) per epochs (B), 20 Ma bins (D), and stages (F). Solid lines indicate mean posterior rates and the shaded areas show 95% HPD. Number of families through time computed by summing up the lifespans of all families for Coleoptera (G). Solid lines indicate mean diversity at each point in time and shaded areas show estimations of different replications that incorporate age uncertainties of fossil occurrences. Light-green area represents the AR, angiosperm radiation, and dark-green area represents the ATR, angiosperm terrestrial revolution. Red-dashed vertical lines indicate major crises: P–Tr, Permian–Triassic; Tr–J, Triassic–Jurassic; K–Pg, Cretaceous–Paleogene. Time is in millions of years. The color of each geological period in the chronostratigraphic scale follows that of the International Chronostratigraphic Chart (v2024/12). Insect silhouettes are from http://phylopic.org/.

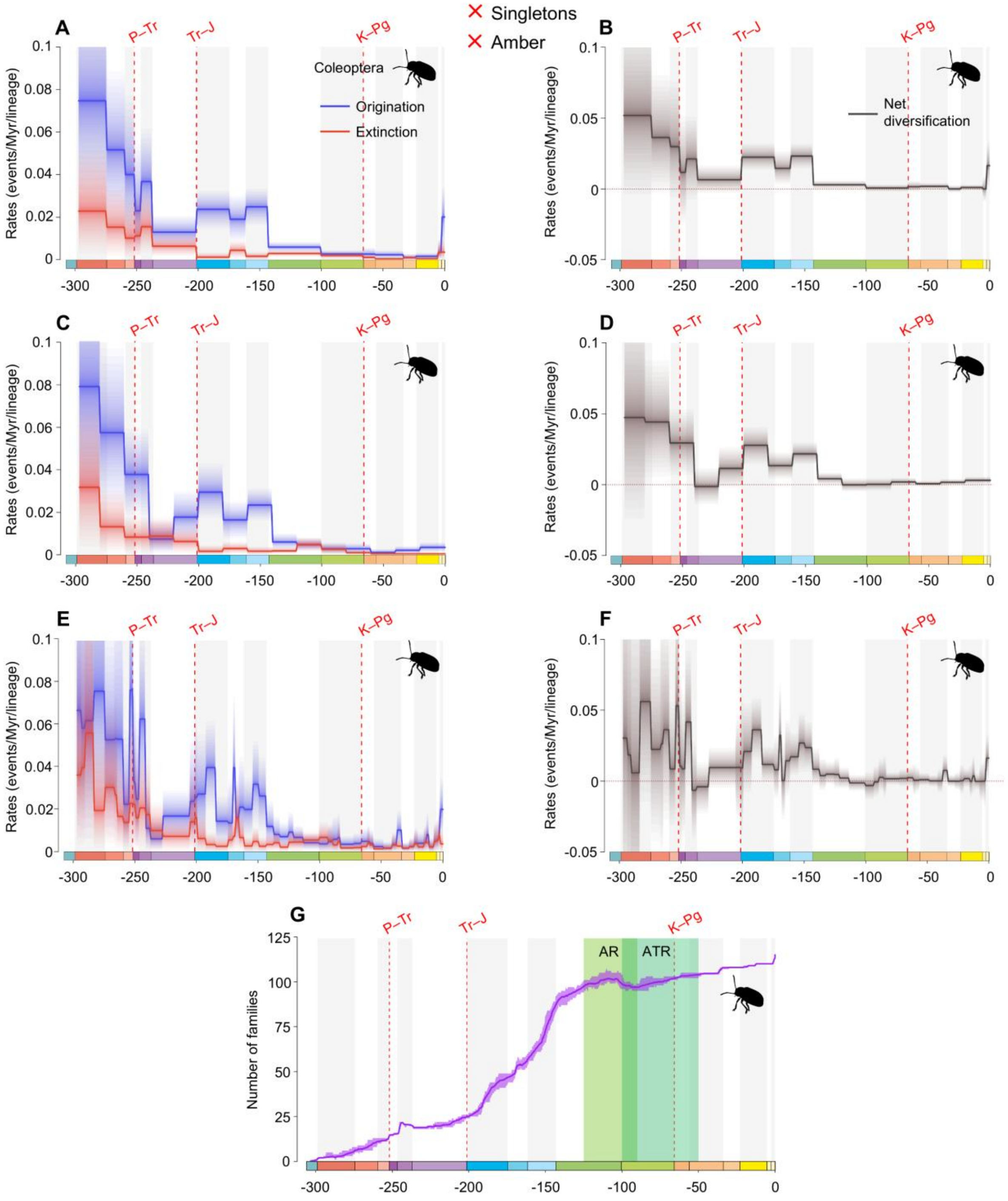


**fig. S38.** Diversification and diversity dynamics of Coleoptera families, excluding singletons and amber occurrences. Bayesian fossil-based inferences of Coleoptera origination and extinction rates at the family level under the birth-death model with epochs (A), 20 Ma bins (C), and stages (E) as constrained shifts (BDCS). Net diversification rates for Coleoptera obtained from the difference between origination and extinction rates (rates above 0 indicate increasing diversity, and rates below 0 indicate declining diversity) per epochs (B), 20 Ma bins (D), and stages (F). Solid lines indicate mean posterior rates and the shaded areas show 95% HPD. Number of families through time computed by summing up the lifespans of all families for Coleoptera (G). Solid lines indicate mean diversity at each point in time and shaded areas show estimations of different replications that incorporate age uncertainties of fossil occurrences. Light-green area represents the AR, angiosperm radiation, and dark-green area represents the ATR, angiosperm terrestrial revolution. Red-dashed vertical lines indicate major crises: P–Tr, Permian–Triassic; Tr–J, Triassic–Jurassic; K–Pg, Cretaceous–Paleogene. Time is in millions of years. The color of each geological period in the chronostratigraphic scale follows that of the International Chronostratigraphic Chart (v2024/12). Insect silhouettes are from http://phylopic.org/.

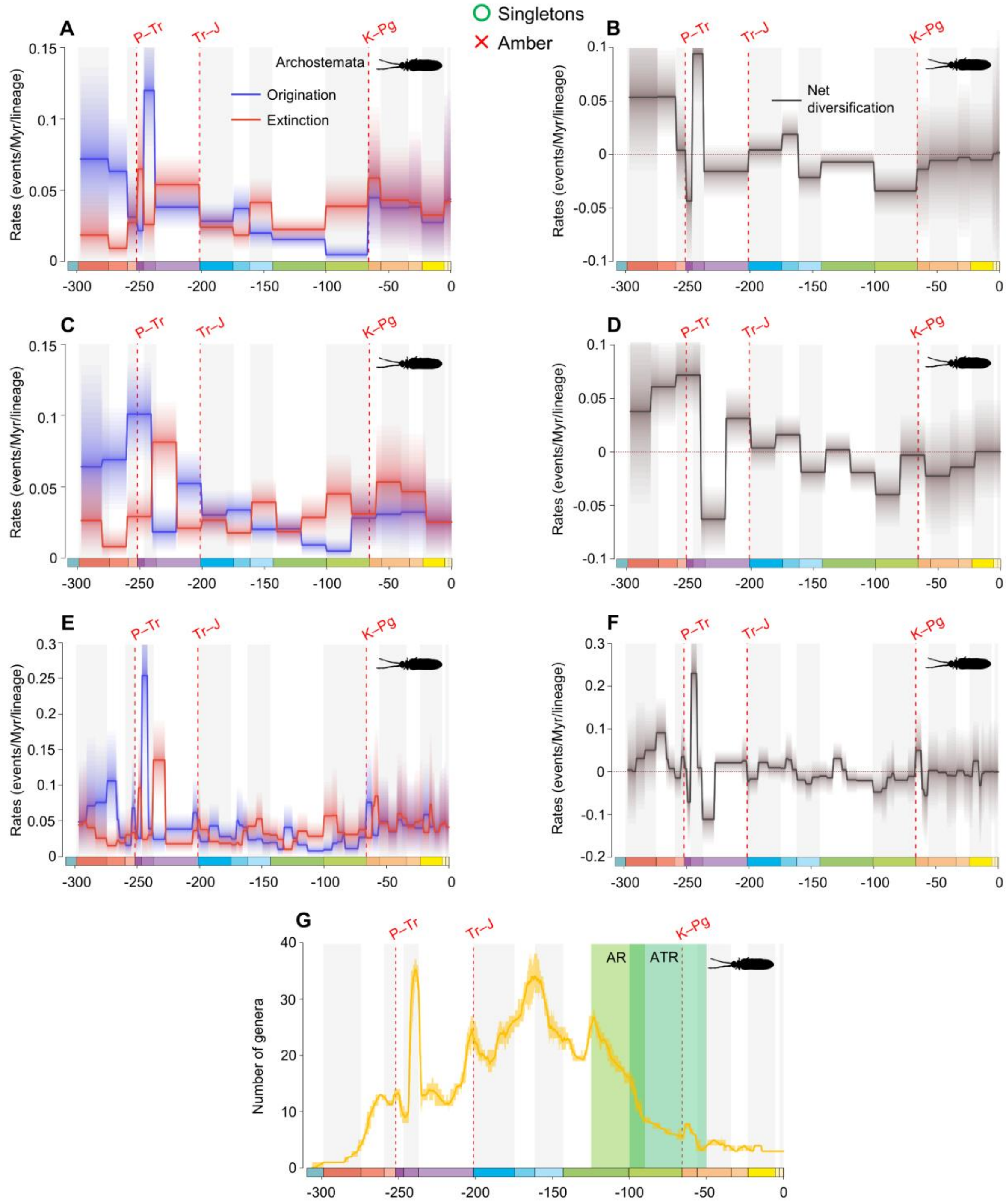


**fig. S39.** Diversification and diversity dynamics of Archostemata genera, considering singletons and excluding amber occurrences. Bayesian fossil-based inferences of Archostemata origination and extinction rates at the genus level under the birth-death model with epochs (A), 20 Ma bins (C), and stages (E) as constrained shifts (BDCS). Net diversification rates for Archostemata obtained from the difference between origination and extinction rates (rates above 0 indicate increasing diversity, and rates below 0 indicate declining diversity) per epochs (B), 20 Ma bins (D), and stages (F). Solid lines indicate mean posterior rates and the shaded areas show 95% HPD.

Number of genera through time computed by summing up the lifespans of all genera for Archostemata (G). Solid lines indicate mean diversity at each point in time and shaded areas show estimations of different replications that incorporate age uncertainties of fossil occurrences. Light-green area represents the AR, angiosperm radiation, and dark-green area represents the ATR, angiosperm terrestrial revolution. Red-dashed vertical lines indicate major crises: P–Tr, Permian–Triassic; Tr–J, Triassic–Jurassic; K–Pg, Cretaceous–Paleogene. Time is in millions of years. The color of each geological period in the chronostratigraphic scale follows that of the International Chronostratigraphic Chart (v2024/12). Insect silhouettes are from http://phylopic.org/.

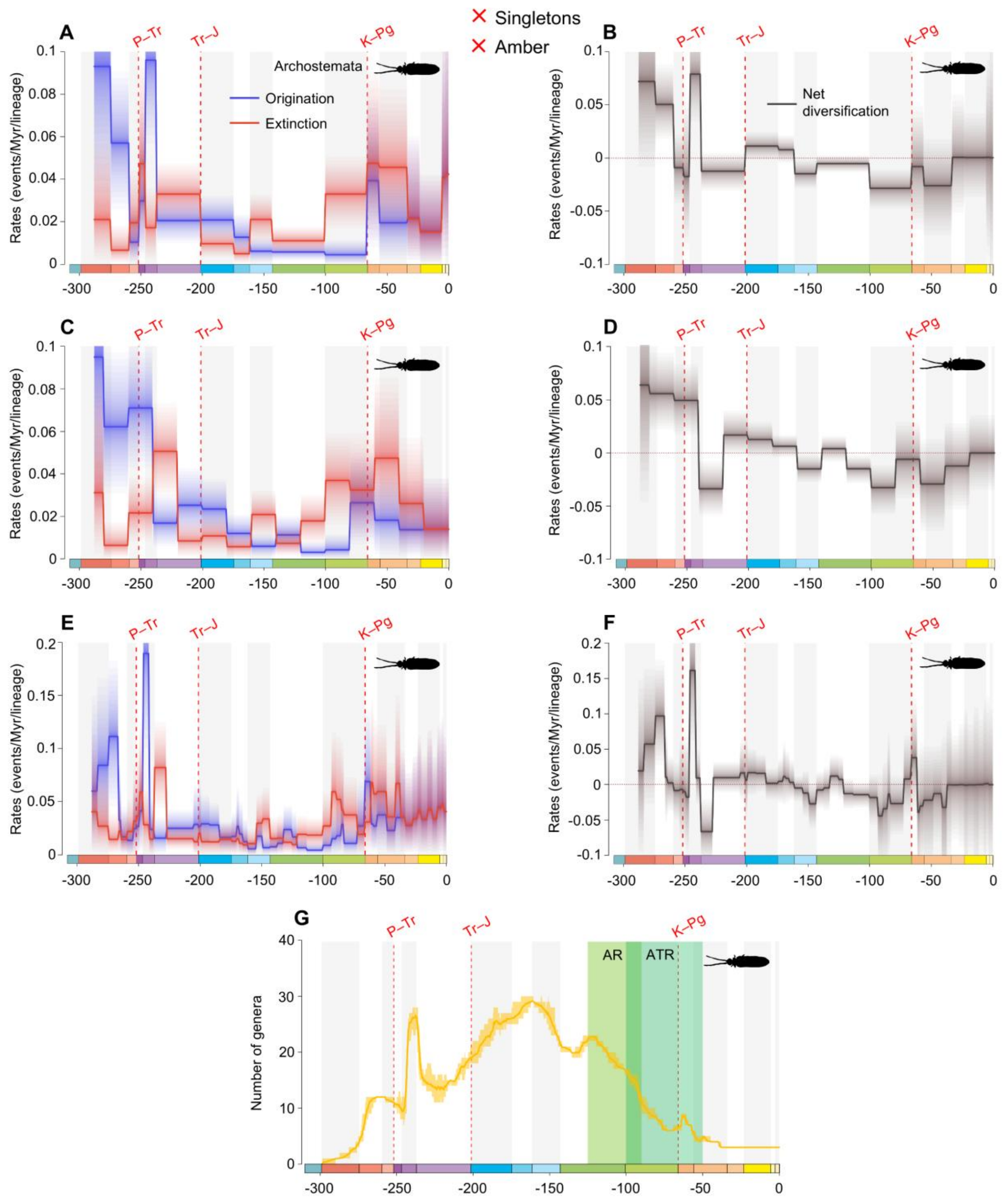


**fig. S40.** Diversification and diversity dynamics of Archostemata genera, excluding singletons and amber occurrences. Bayesian fossil-based inferences of Archostemata origination and extinction rates at the genus level under the birth-death model with epochs (A), 20 Ma bins (C), and stages (E) as constrained shifts (BDCS). Net diversification rates for Archostemata obtained from the difference between origination and extinction rates (rates above 0 indicate increasing diversity, and rates below 0 indicate declining diversity) per epochs (B), 20 Ma bins (D), and stages (F). Solid lines indicate mean posterior rates and the shaded areas show 95% HPD. Number of genera through time computed by summing up the lifespans of all genera for Archostemata (G). Solid lines indicate mean diversity at each point in time and shaded areas show estimations of different replications that incorporate age uncertainties of fossil occurrences. Light-green area represents the AR, angiosperm radiation, and dark-green area represents the ATR, angiosperm terrestrial revolution. Red-dashed vertical lines indicate major crises: P–Tr, Permian–Triassic; Tr–J, Triassic–Jurassic; K–Pg, Cretaceous–Paleogene. Time is in millions of years. The color of each geological period in the chronostratigraphic scale follows that of the International Chronostratigraphic Chart (v2024/12). Insect silhouettes are from http://phylopic.org/.

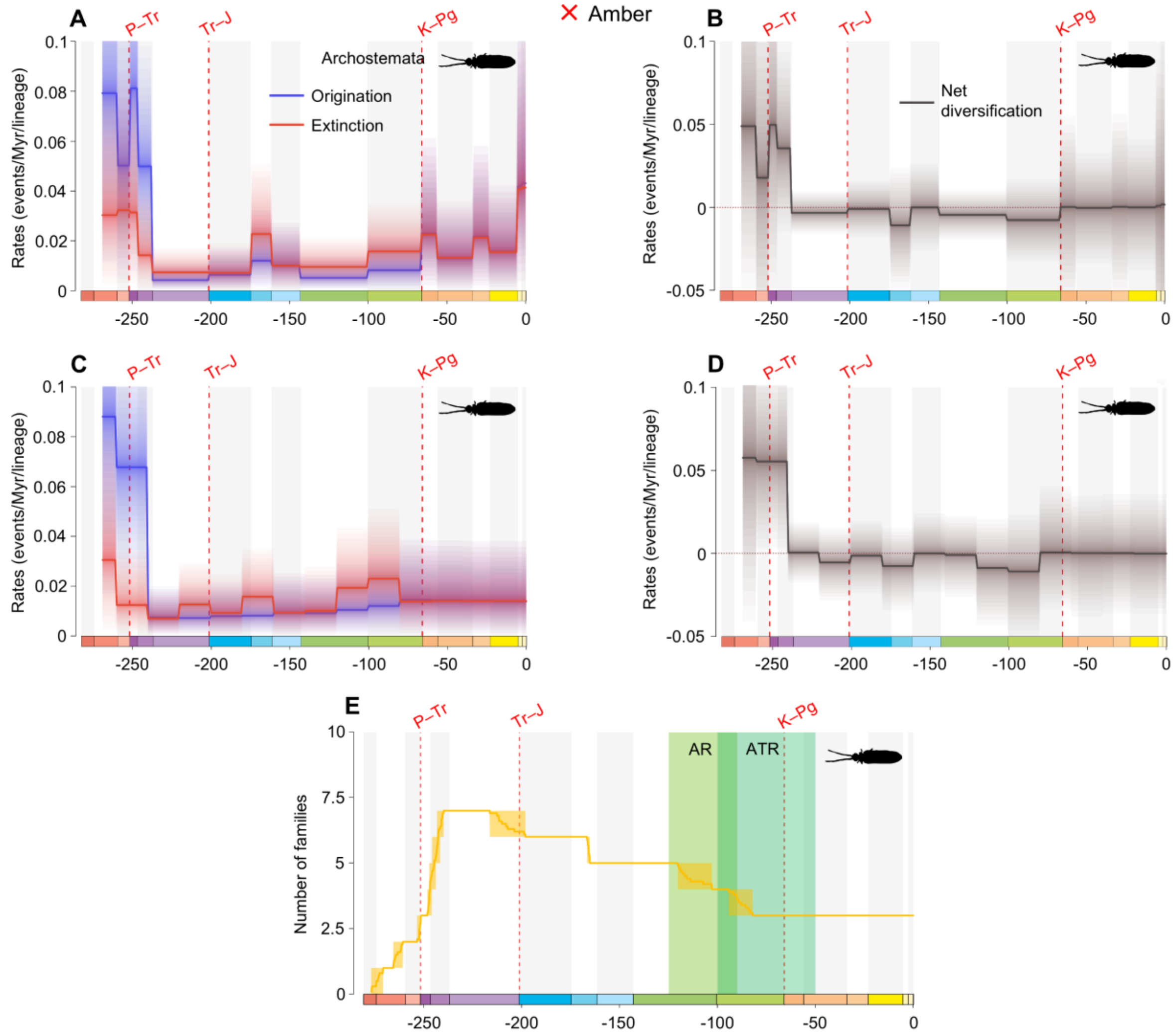


**fig. S41.** Diversification and diversity dynamics of Archostemata families excluding amber occurrences. There are no singletons at the family-level in Archostemata. Bayesian fossil-based inferences of Archostemata origination and extinction rates at the family level under the birth-death model with epochs (A), and 20 Ma bins (C) constrained shifts (BDCS). Net diversification rates for Archostemata obtained from the difference between origination and extinction rates (rates above 0 indicate increasing diversity, and rates below 0 indicate declining diversity) per epochs (B), and 20 Ma bins (D). Solid lines indicate mean posterior rates and the shaded areas show 95% HPD. Number of families through time computed by summing up the lifespans of all families for Archostemata (E). Solid lines indicate mean diversity at each point in time and shaded areas show estimations of different replications that incorporate age uncertainties of fossil occurrences. Light-green area represents the AR, angiosperm radiation, and dark-green area represents the ATR, angiosperm terrestrial revolution. Red-dashed vertical lines indicate major crises: P–Tr, Permian–Triassic; Tr–J, Triassic–Jurassic; K–Pg, Cretaceous–Paleogene. Time is in millions of years. The color of each geological period in the chronostratigraphic scale follows that of the International Chronostratigraphic Chart (v2024/12). Insect silhouettes are from http://phylopic.org/.

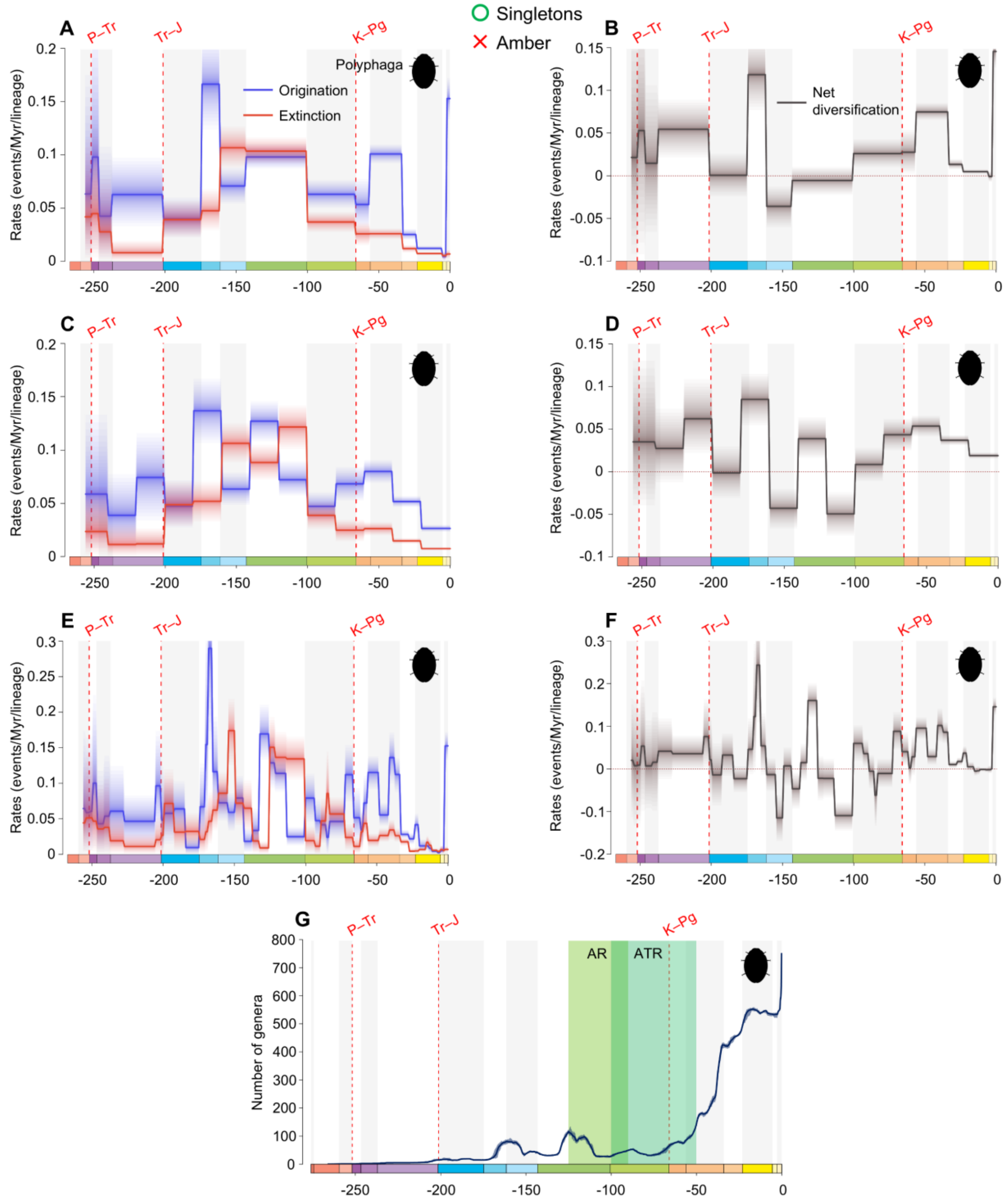


**fig. S42.** Diversification and diversity dynamics of Polyphaga genera, considering singletons and excluding amber occurrences. Bayesian fossil-based inferences of Polyphaga origination and extinction rates at the genus level under the birth-death model with epochs (A), 20 Ma bins (C), and stages (E) as constrained shifts (BDCS). Net diversification rates for Polyphaga obtained from the difference between origination and extinction rates (rates above 0 indicate increasing diversity, and rates below 0 indicate declining diversity) per epochs (B), 20 Ma bins (D), and stages (F). Solid lines indicate mean posterior rates and the shaded areas show 95% HPD. Number of genera through time computed by summing up the lifespans of all genera for Polyphaga (G). Solid lines indicate mean diversity at each point in time and shaded areas show estimations of different replications that incorporate age uncertainties of fossil occurrences. Light-green area represents the AR, angiosperm radiation, and dark-green area represents the ATR, angiosperm terrestrial revolution. Red-dashed vertical lines indicate major crises: P–Tr, Permian–Triassic; Tr–J, Triassic–Jurassic; K–Pg, Cretaceous–Paleogene. Time is in millions of years. The color of each geological period in the chronostratigraphic scale follows that of the International Chronostratigraphic Chart (v2024/12). Insect silhouettes are from http://phylopic.org/.

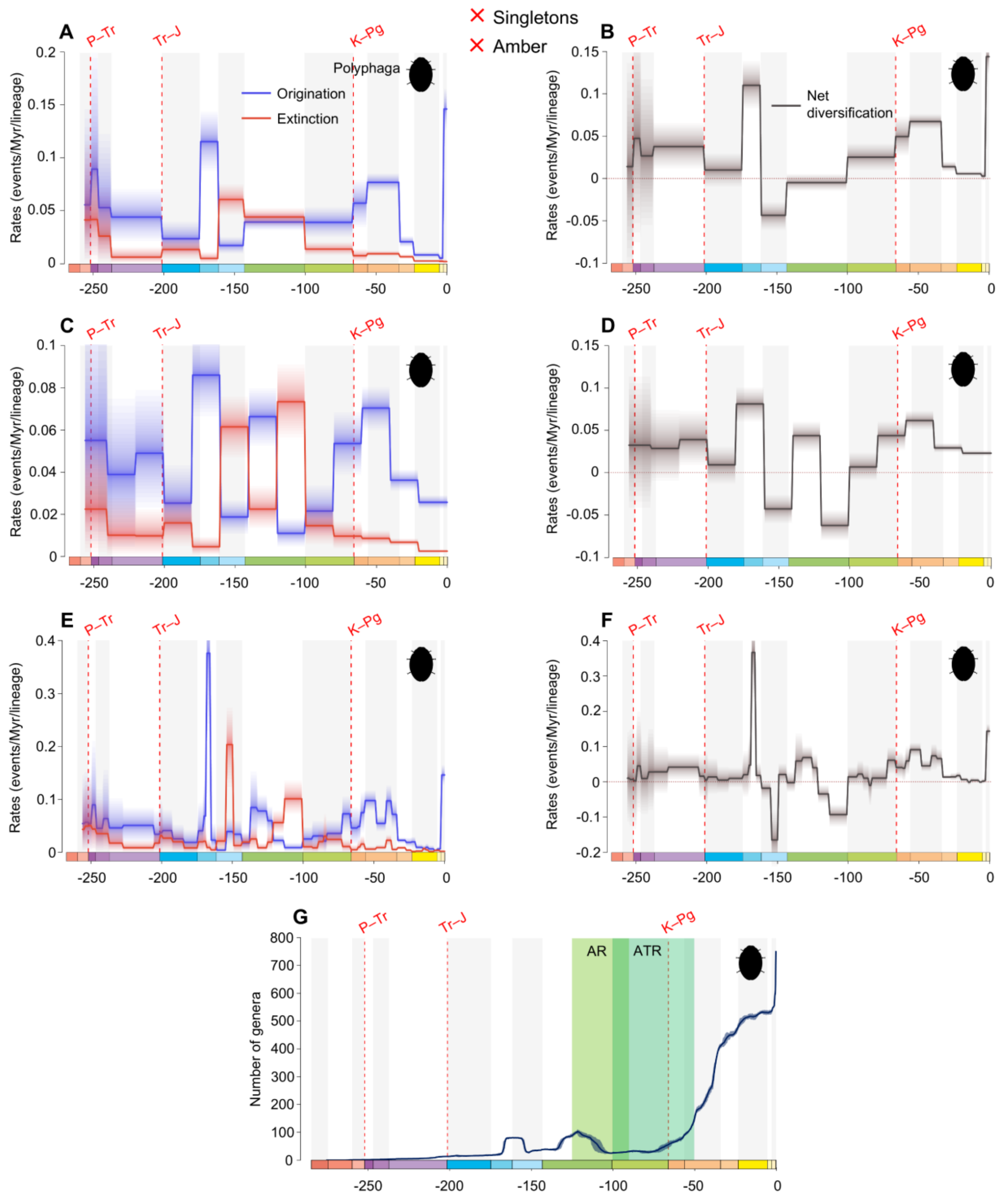


**fig. S43.** Diversification and diversity dynamics of Polyphaga genera, excluding singletons and amber occurrences. Bayesian fossil-based inferences of Polyphaga origination and extinction rates at the genus level under the birth-death model with epochs (A), 20 Ma bins (C), and stages (E) as constrained shifts (BDCS). Net diversification rates for Polyphaga obtained from the difference between origination and extinction rates (rates above 0 indicate increasing diversity, and rates below 0 indicate declining diversity) per epochs (B), 20 Ma bins (D), and stages (F). Solid lines indicate mean posterior rates and the shaded areas show 95% HPD. Number of genera through time computed by summing up the lifespans of all genera for Polyphaga (G). Solid lines indicate mean diversity at each point in time and shaded areas show estimations of different replications that incorporate age uncertainties of fossil occurrences. Light-green area represents the AR, angiosperm radiation, and dark-green area represents the ATR, angiosperm terrestrial revolution. Red-dashed vertical lines indicate major crises: P–Tr, Permian–Triassic; Tr–J, Triassic–Jurassic; K–Pg, Cretaceous–Paleogene. Time is in millions of years. The color of each geological period in the chronostratigraphic scale follows that of the International Chronostratigraphic Chart (v2024/12). Insect silhouettes are from http://phylopic.org/.

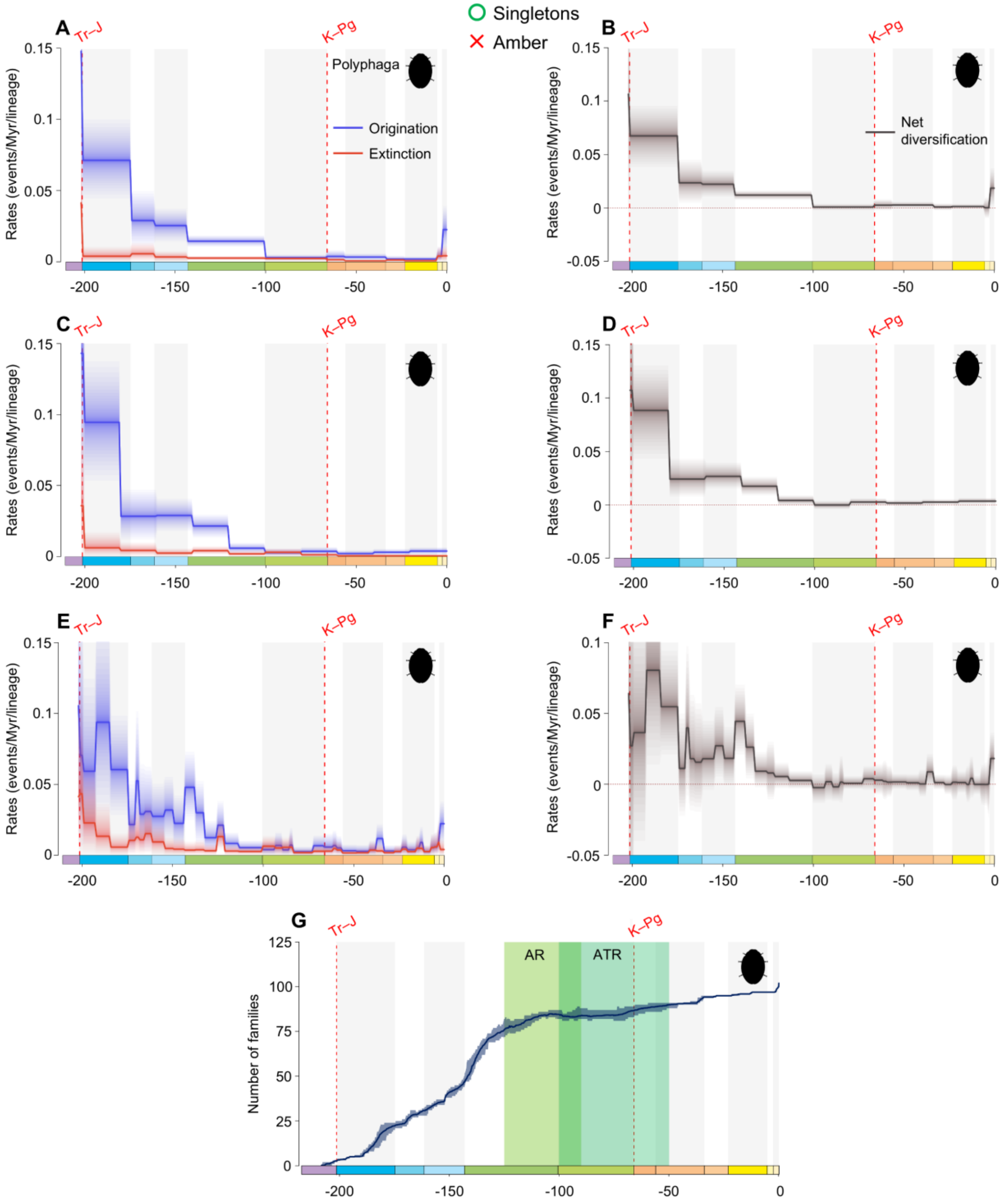


**fig. S44.** Diversification and diversity dynamics of Polyphaga families, considering singletons and excluding amber occurrences. Bayesian fossil-based inferences of Polyphaga origination and extinction rates at the family level under the birth-death model with epochs (A), 20 Ma bins (C), and stages (E) as constrained shifts (BDCS). Net diversification rates for Polyphaga obtained from the difference between origination and extinction rates (rates above 0 indicate increasing diversity, and rates below 0 indicate declining diversity) per epochs (B), 20 Ma bins (D), and stages (F). Solid lines indicate mean posterior rates and the shaded areas show 95% HPD. Number of families through time computed by summing up the lifespans of all families for Polyphaga (G). Solid lines indicate mean diversity at each point in time and shaded areas show estimations of different replications that incorporate age uncertainties of fossil occurrences. Light-green area represents the AR, angiosperm radiation, and dark-green area represents the ATR, angiosperm terrestrial revolution. Red-dashed vertical lines indicate major crises: Tr–J, Triassic–Jurassic; K–Pg, Cretaceous–Paleogene. Time is in millions of years. The color of each geological period in the chronostratigraphic scale follows that of the International Chronostratigraphic Chart (v2024/12). Insect silhouettes are from http://phylopic.org/.

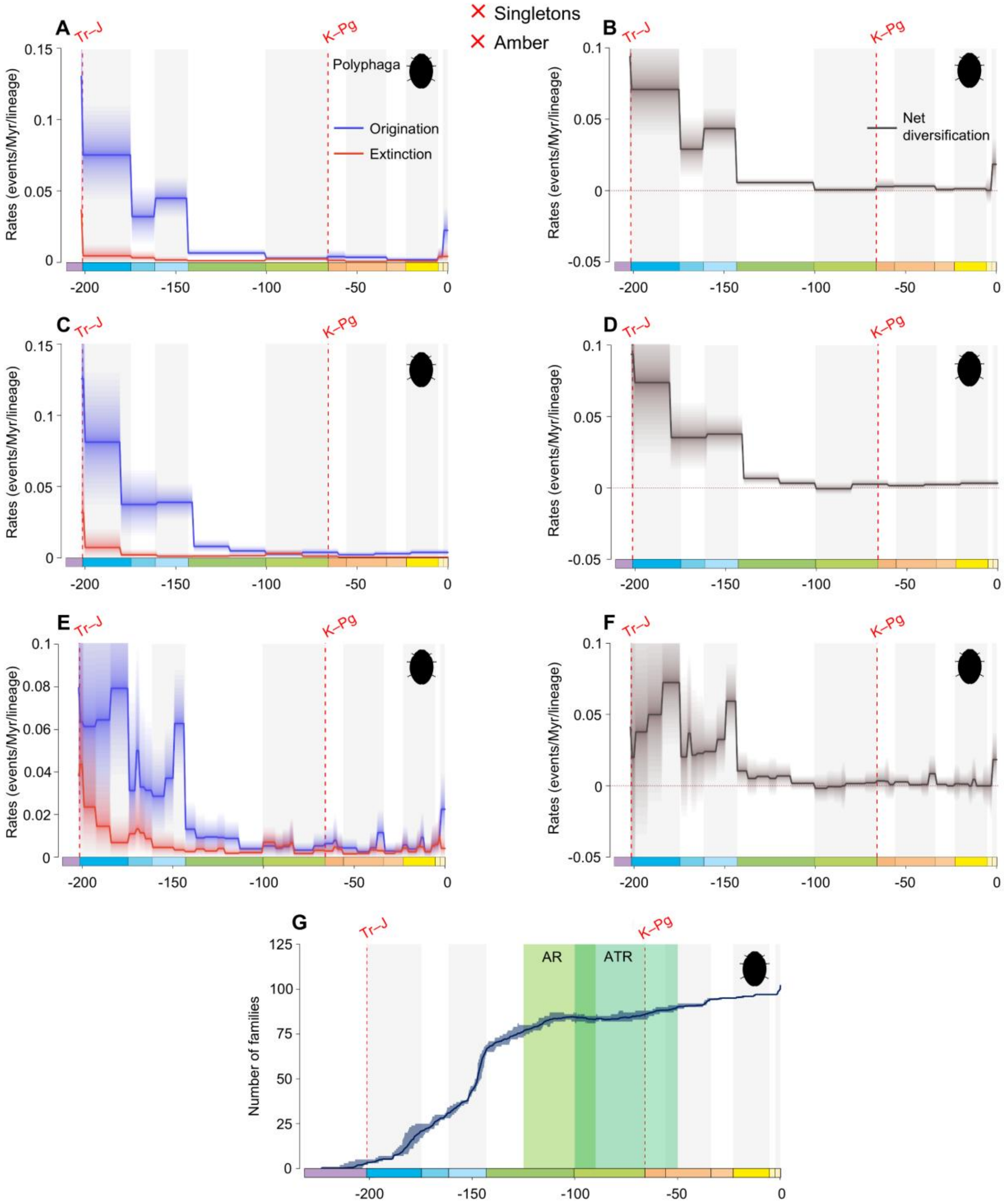


**fig. S45.** Diversification and diversity dynamics of Polyphaga families, excluding singletons and amber occurrences. Bayesian fossil-based inferences of Polyphaga origination and extinction rates at the family level under the birth-death model with epochs (A), 20 Ma bins (C), and stages (E) as constrained shifts (BDCS). Net diversification rates for Polyphaga obtained from the difference between origination and extinction rates (rates above 0 indicate increasing diversity, and rates below 0 indicate declining diversity) per epochs (B), 20 Ma bins (D), and stages (F). Solid lines indicate mean posterior rates and the shaded areas show 95% HPD. Number of families through time computed by summing up the lifespans of all families for Polyphaga (G). Solid lines indicate mean diversity at each point in time and shaded areas show estimations of different replications that incorporate age uncertainties of fossil occurrences. Light-green area represents the AR, angiosperm radiation, and dark-green area represents the ATR, angiosperm terrestrial revolution. Red-dashed vertical lines indicate major crises: Tr–J, Triassic–Jurassic; K–Pg, Cretaceous–Paleogene. Time is in millions of years. The color of each geological period in the chronostratigraphic scale follows that of the International Chronostratigraphic Chart (v2024/12). Insect silhouettes are from http://phylopic.org/.

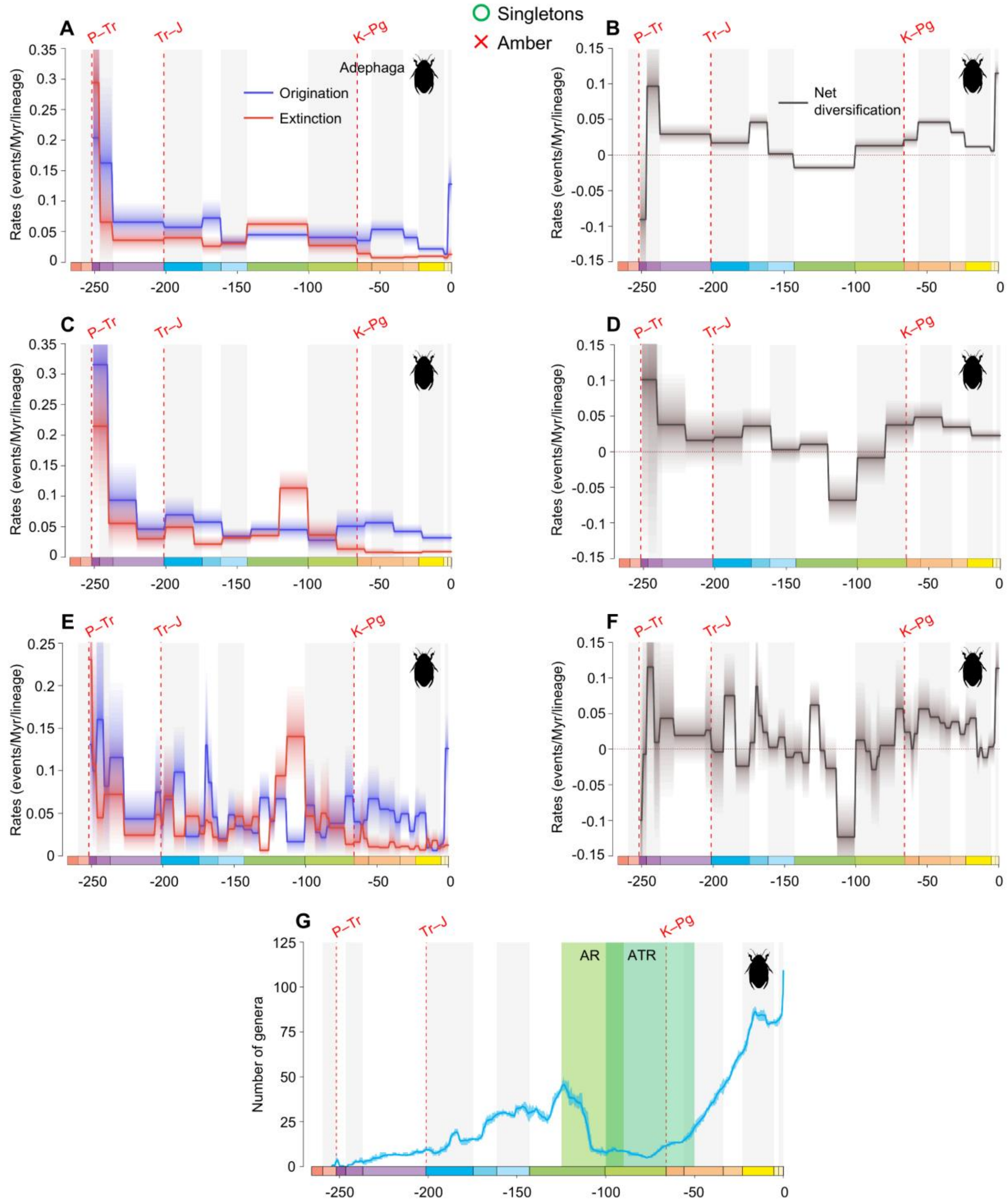


**fig. S46.** Diversification and diversity dynamics of Adephaga genera, considering singletons and excluding amber occurrences. Bayesian fossil-based inferences of Adephaga origination and extinction rates at the genus level under the birth-death model with epochs (A), 20 Ma bins (C), and stages (E) as constrained shifts (BDCS). Net diversification rates for Adephaga obtained from the difference between origination and extinction rates (rates above 0 indicate increasing diversity, and rates below 0 indicate declining diversity) per epochs (B), 20 Ma bins (D), and stages (F). Solid lines indicate mean posterior rates and the shaded areas show 95% HPD. Number of genera through time computed by summing up the lifespans of all genera for Adephaga (G). Solid lines indicate mean diversity at each point in time and shaded areas show estimations of different replications that incorporate age uncertainties of fossil occurrences. Light-green area represents the AR, angiosperm radiation, and dark-green area represents the ATR, angiosperm terrestrial revolution. Red-dashed vertical lines indicate major crises: P–Tr, Permian–Triassic; Tr–J, Triassic–Jurassic; K–Pg, Cretaceous–Paleogene. Time is in millions of years. The color of each geological period in the chronostratigraphic scale follows that of the International Chronostratigraphic Chart (v2024/12). Insect silhouettes are from http://phylopic.org/.

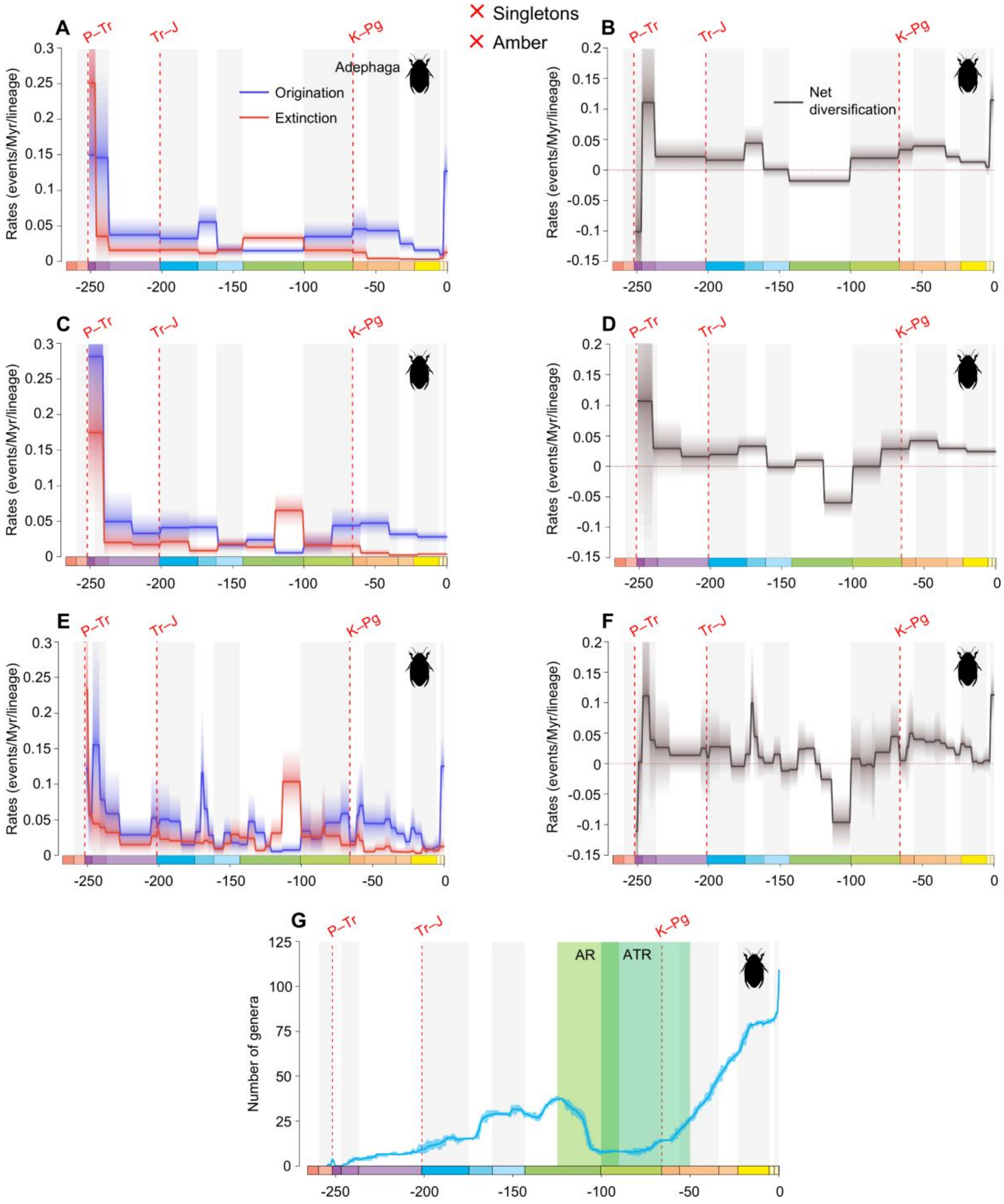


**fig. S47.** Diversification and diversity dynamics of Adephaga genera, excluding singletons and amber occurrences. Bayesian fossil-based inferences of Adephaga origination and extinction rates at the genus level under the birth-death model with epochs (A), 20 Ma bins (C), and stages (E) as constrained shifts (BDCS). Net diversification rates for Adephaga obtained from the difference between origination and extinction rates (rates above 0 indicate increasing diversity, and rates below 0 indicate declining diversity) per epochs (B), 20 Ma bins (D), and stages (F). Solid lines indicate mean posterior rates and the shaded areas show 95% HPD. Number of genera through time computed by summing up the lifespans of all genera for Adephaga (G). Solid lines indicate mean diversity at each point in time and shaded areas show estimations of different replications that incorporate age uncertainties of fossil occurrences. Light-green area represents the AR, angiosperm radiation, and dark-green area represents the ATR, angiosperm terrestrial revolution. Red-dashed vertical lines indicate major crises: P–Tr, Permian–Triassic; Tr–J, Triassic–Jurassic; K–Pg, Cretaceous–Paleogene. Time is in millions of years. The color of each geological period in the chronostratigraphic scale follows that of the International Chronostratigraphic Chart (v2024/12). Insect silhouettes are from http://phylopic.org/.

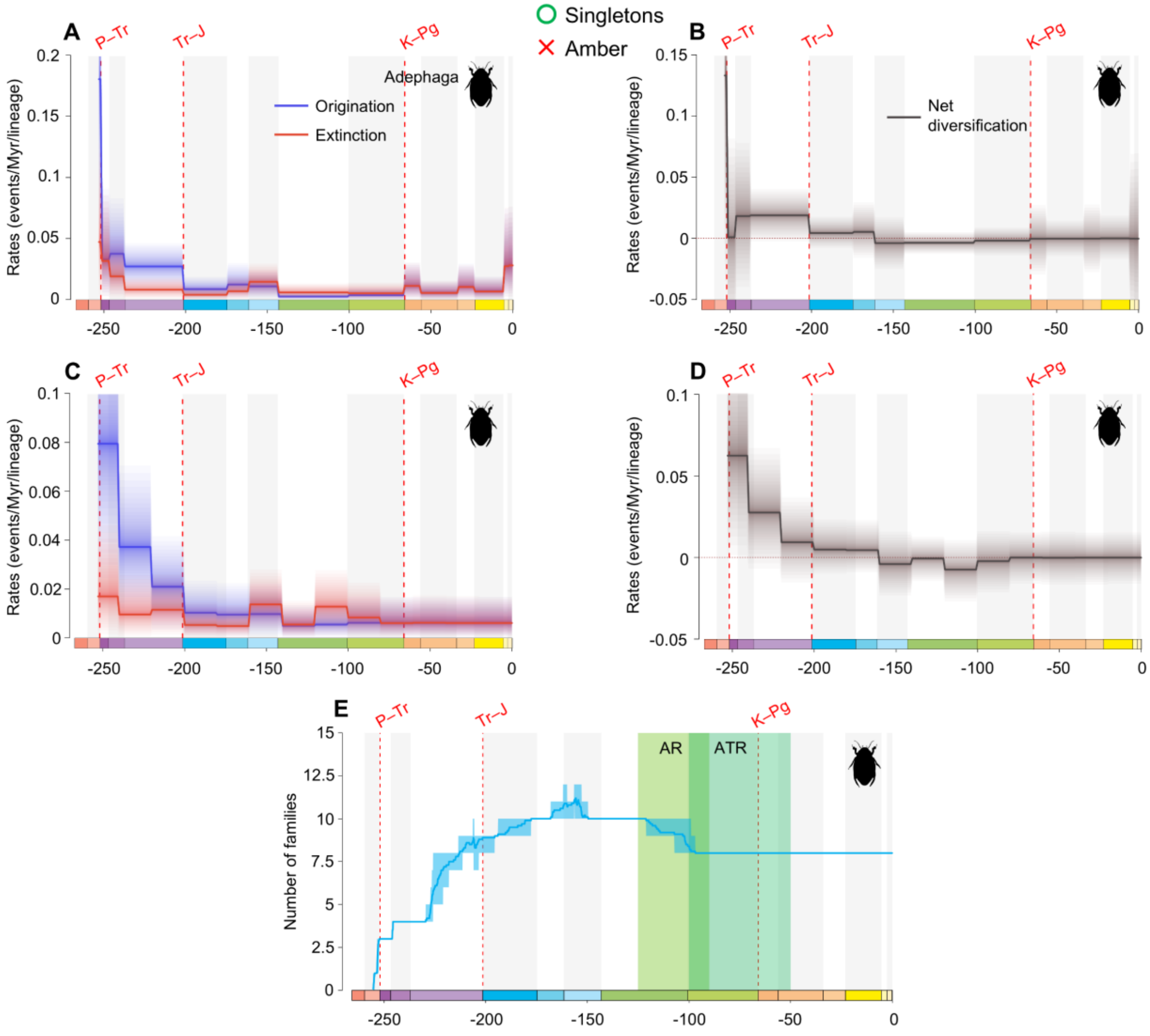


**fig. S48.** Diversification and diversity dynamics of Adephaga families, considering singletons and excluding amber occurrences. Bayesian fossil-based inferences of Adephaga origination and extinction rates at the family level under the birth-death model with epochs (A), and 20 Ma bins (C) as constrained shifts (BDCS). Net diversification rates for Adephaga obtained from the difference between origination and extinction rates (rates above 0 indicate increasing diversity, and rates below 0 indicate declining diversity) per epochs (B), 20 Ma bins (D). Solid lines indicate mean posterior rates and the shaded areas show 95% HPD. Number of families through time computed by summing up the lifespans of all families for Adephaga (E). Solid lines indicate mean diversity at each point in time and shaded areas show estimations of different replications that incorporate age uncertainties of fossil occurrences. Light-green area represents the AR, angiosperm radiation, and dark-green area represents the ATR, angiosperm terrestrial revolution. Red-dashed vertical lines indicate major crises: P–Tr, Permian–Triassic; Tr–J, Triassic–Jurassic; K–Pg, Cretaceous–Paleogene. Time is in millions of years. The color of each geological period in the chronostratigraphic scale follows that of the International Chronostratigraphic Chart (v2024/12). Insect silhouettes are from http://phylopic.org/.

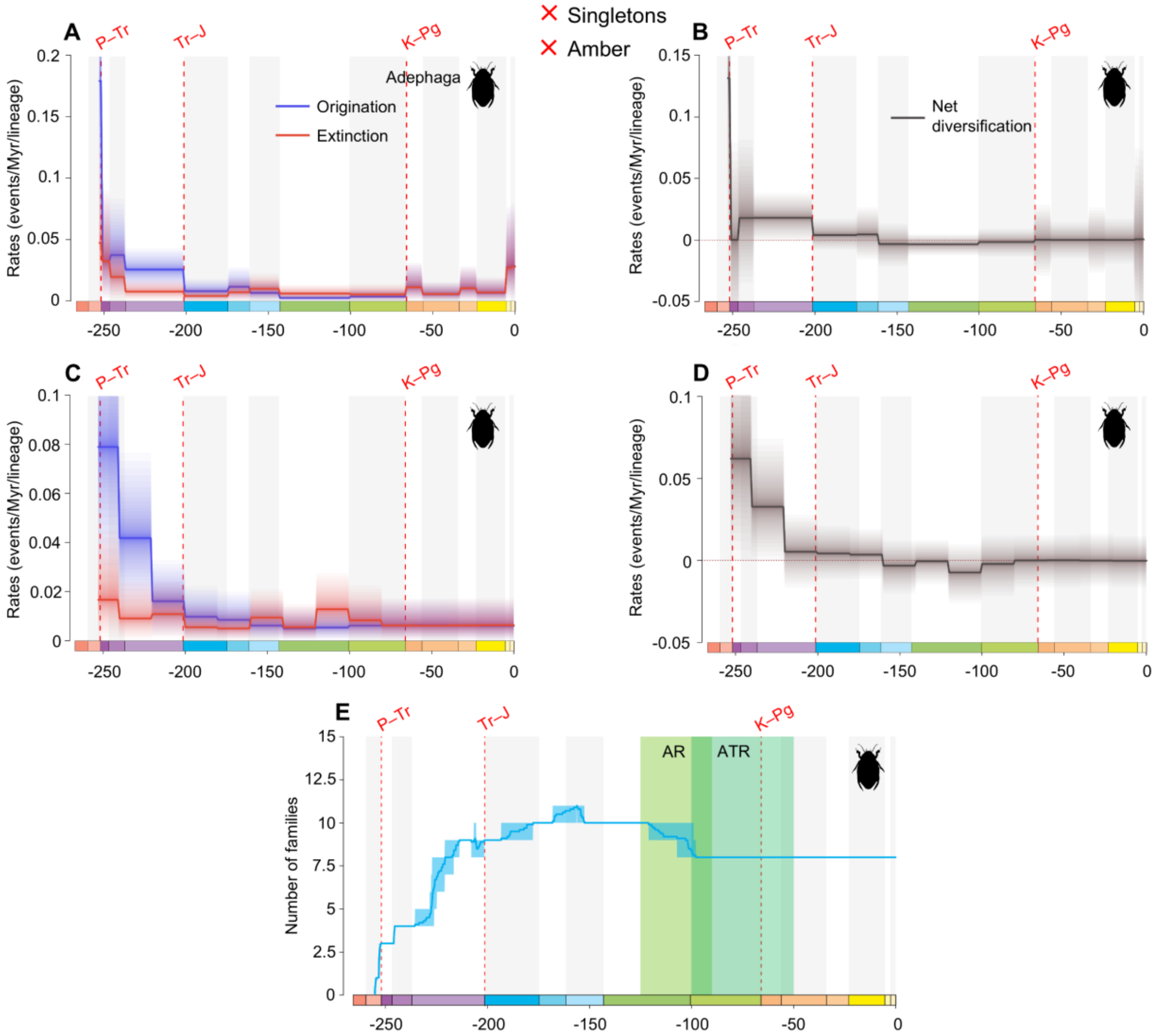


**fig. S49.** Diversification and diversity dynamics of Adephaga families, excluding singletons and amber occurrences. Bayesian fossil-based inferences of Adephaga origination and extinction rates at the family level under the birth-death model with epochs (A), and 20 Ma bins (C) as constrained shifts (BDCS). Net diversification rates for Adephaga obtained from the difference between origination and extinction rates (rates above 0 indicate increasing diversity, and rates below 0 indicate declining diversity) per epochs (B), 20 Ma bins (D). Solid lines indicate mean posterior rates and the shaded areas show 95% HPD. Number of families through time computed by summing up the lifespans of all families for Adephaga (E). Solid lines indicate mean diversity at each point in time and shaded areas show estimations of different replications that incorporate age uncertainties of fossil occurrences. Light-green area represents the AR, angiosperm radiation, and dark-green area represents the ATR, angiosperm terrestrial revolution. Red-dashed vertical lines indicate major crises: P–Tr, Permian–Triassic; Tr–J, Triassic–Jurassic; K–Pg, Cretaceous–Paleogene. Time is in millions of years. The color of each geological period in the chronostratigraphic scale follows that of the International Chronostratigraphic Chart (v2024/12). Insect silhouettes are from http://phylopic.org/.

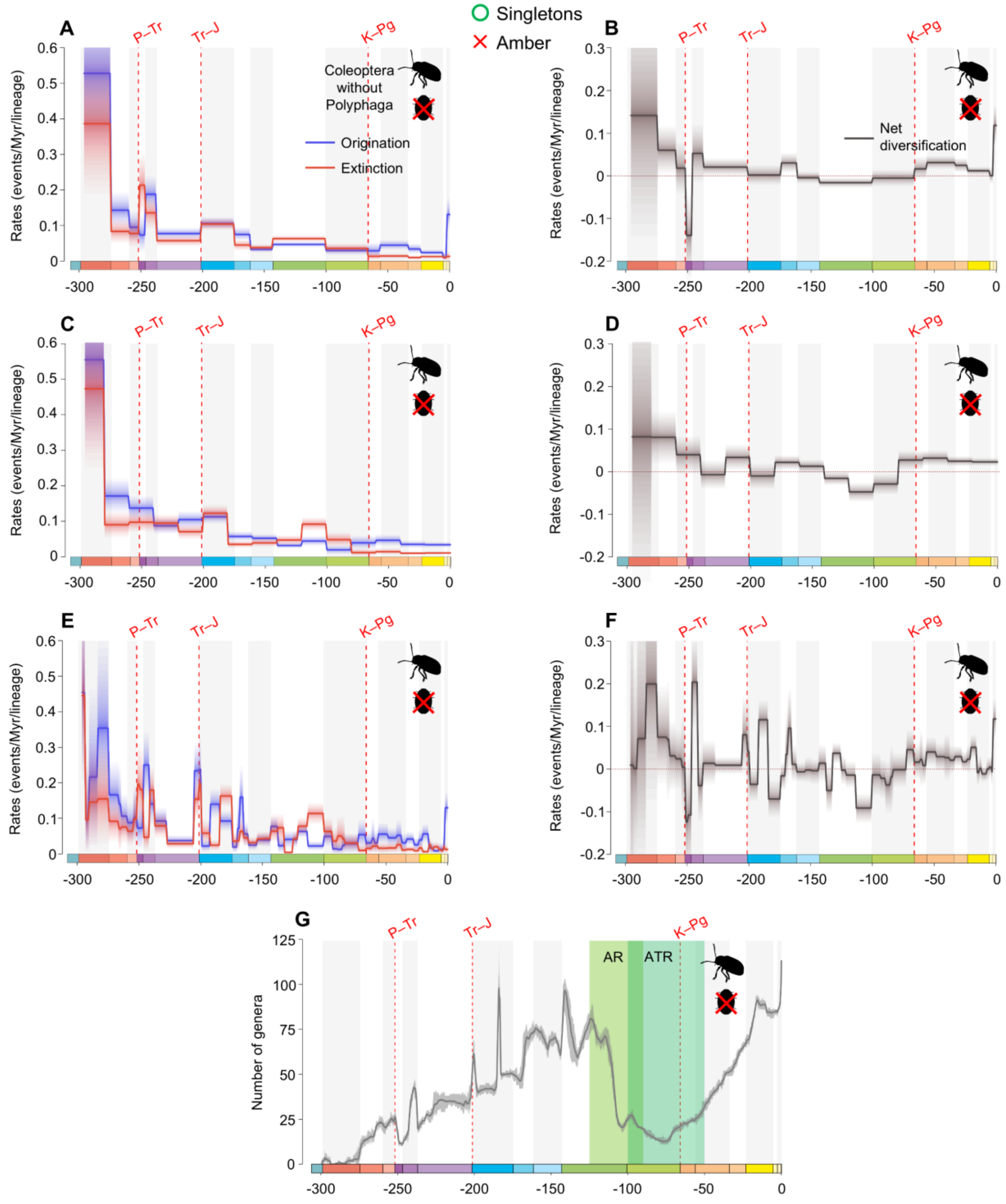


**fig. S50.** Diversification and diversity dynamics of Coleoptera genera without Polyphaga, considering singletons and excluding amber occurrences. Bayesian fossil-based inferences of Coleoptera without Polyphaga origination and extinction rates at the genus level under the birth-death model with epochs (A), 20 Ma bins (C), and stages (E) as constrained shifts (BDCS). Net diversification rates for Coleoptera without Polyphaga obtained from the difference between origination and extinction rates (rates above 0 indicate increasing diversity, and rates below 0 indicate declining diversity) per epochs (B), 20 Ma bins (D), and stages (F). Solid lines indicate mean posterior rates and the shaded areas show 95% HPD. Number of genera through time computed by summing up the lifespans of all genera for Coleoptera without Polyphaga (G). Solid lines indicate mean diversity at each point in time and shaded areas show estimations of different replications that incorporate age uncertainties of fossil occurrences. Light-green area represents the AR, angiosperm radiation, and dark-green area represents the ATR, angiosperm terrestrial revolution. Red-dashed vertical lines indicate major crises: P–Tr, Permian–Triassic; Tr–J, Triassic–Jurassic; K–Pg, Cretaceous–Paleogene. Time is in millions of years. The color of each geological period in the chronostratigraphic scale follows that of the International Chronostratigraphic Chart (v2024/12). Insect silhouettes are from http://phylopic.org/.

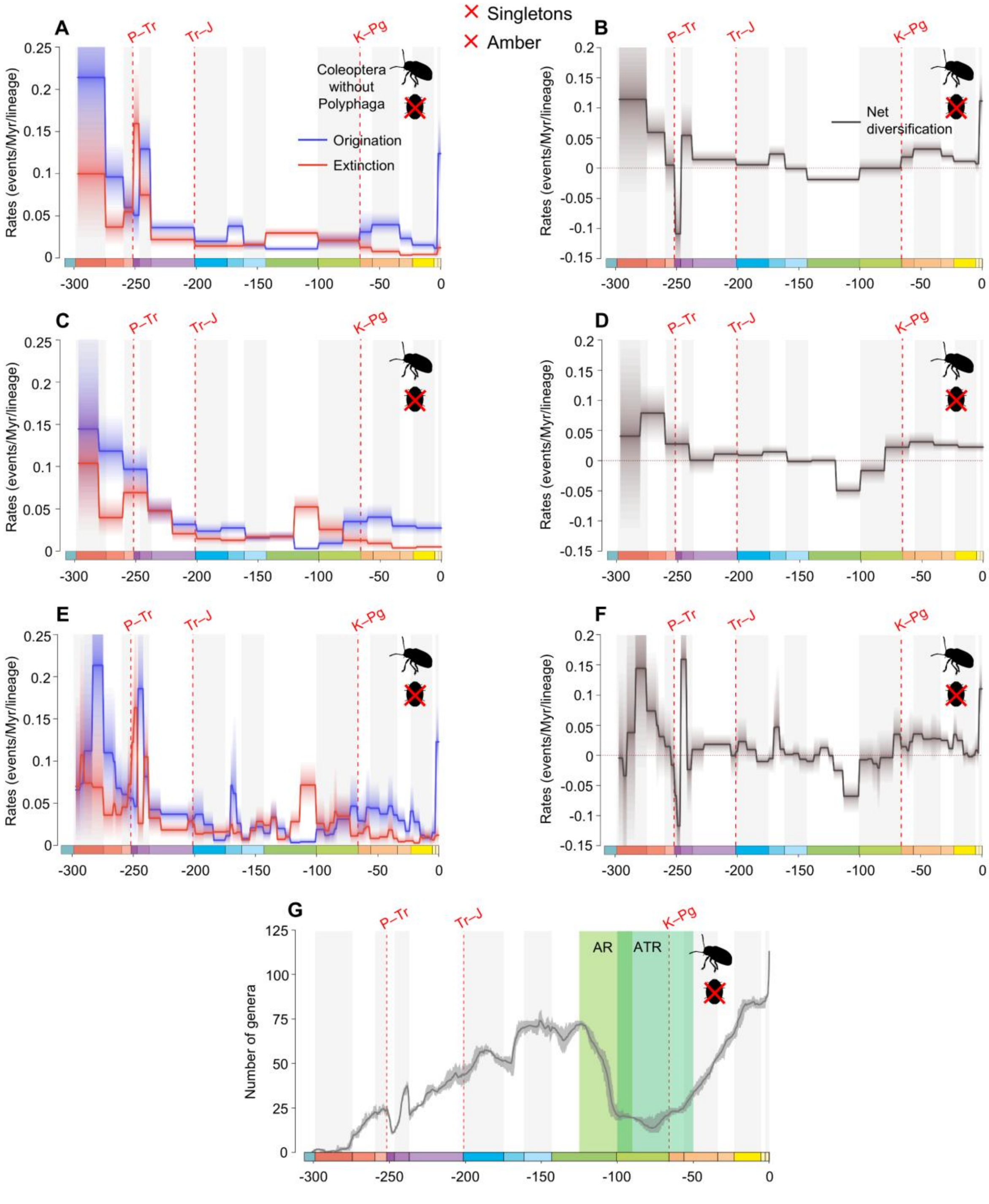


**fig. S51.** Diversification and diversity dynamics of Coleoptera genera without Polyphaga, excluding singletons and amber occurrences. Bayesian fossil-based inferences of Coleoptera without Polyphaga origination and extinction rates at the genus level under the birth-death model with epochs (A), 20 Ma bins (C), and stages (E) as constrained shifts (BDCS). Net diversification rates for Coleoptera without Polyphaga obtained from the difference between origination and extinction rates (rates above 0 indicate increasing diversity, and rates below 0 indicate declining diversity) per epochs (B), 20 Ma bins (D), and stages (F). Solid lines indicate mean posterior rates

and the shaded areas show 95% HPD. Number of genera through time computed by summing up the lifespans of all genera for Coleoptera without Polyphaga (G). Solid lines indicate mean diversity at each point in time and shaded areas show estimations of different replications that incorporate age uncertainties of fossil occurrences. Light-green area represents the AR, angiosperm radiation, and dark-green area represents the ATR, angiosperm terrestrial revolution. Red-dashed vertical lines indicate major crises: P–Tr, Permian–Triassic; Tr–J, Triassic–Jurassic; K–Pg, Cretaceous–Paleogene. Time is in millions of years. The color of each geological period in the chronostratigraphic scale follows that of the International Chronostratigraphic Chart (v2024/12). Insect silhouettes are from http://phylopic.org/.

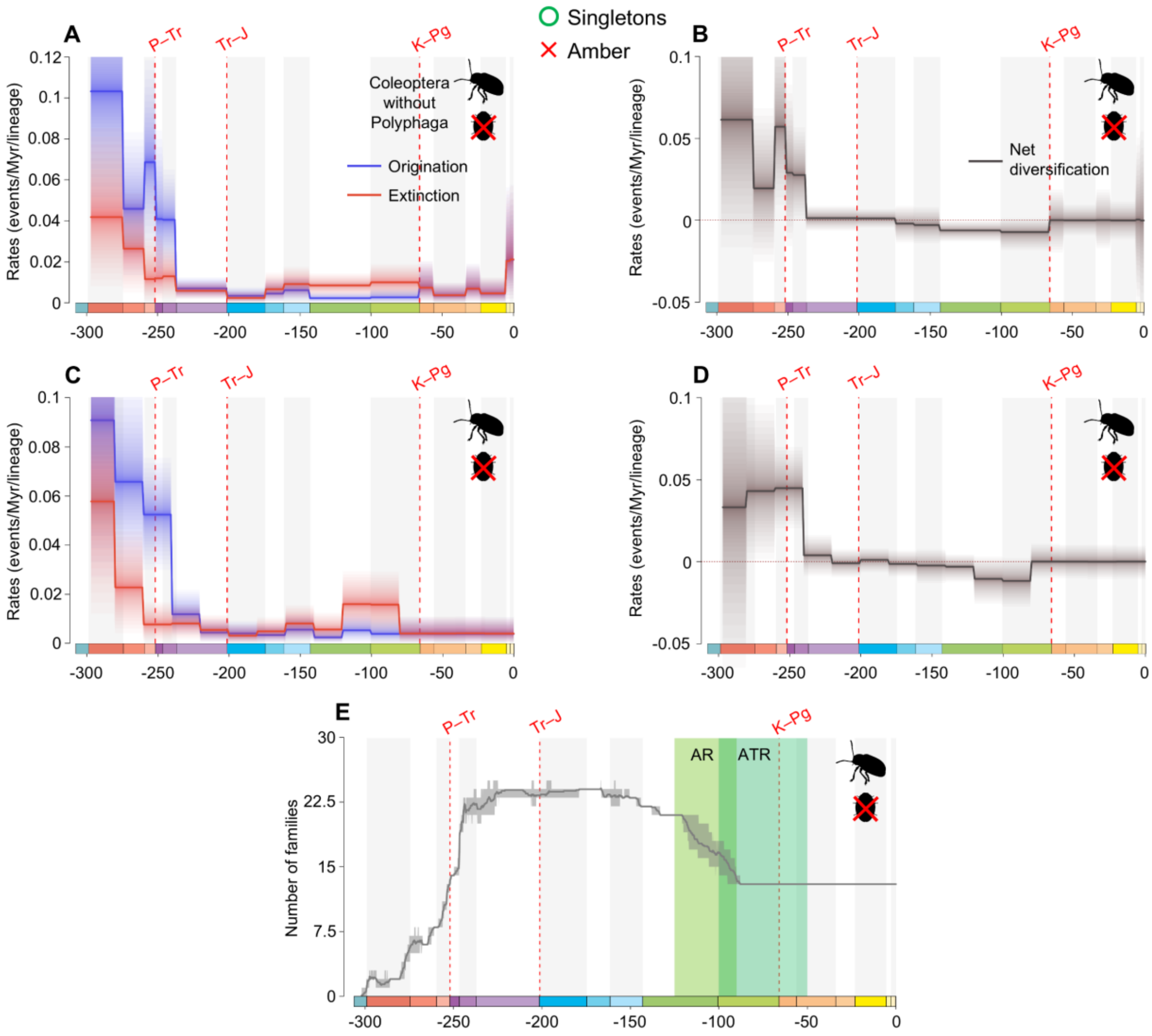


**fig. S52.** Diversification and diversity dynamics of Coleoptera families without Polyphaga, considering singletons and excluding amber occurrences. Bayesian fossil-based inferences of Coleoptera without Polyphaga origination and extinction rates at the family level under the birth-death model with epochs (A), 20 Ma bins (C) as constrained shifts (BDCS). Net diversification rates for Coleoptera without Polyphaga obtained from the difference between origination and extinction rates (rates above 0 indicate increasing diversity, and rates below 0 indicate declining diversity) per epochs (B), 20 Ma bins (D). Solid lines indicate mean posterior rates and the shaded areas show 95% HPD. Number of families through time computed by summing up the lifespans of all families for Coleoptera without Polyphaga (E). Solid lines indicate mean diversity at each point in time and shaded areas show estimations of different replications that incorporate age uncertainties of fossil occurrences. Light-green area represents the AR, angiosperm radiation, and dark-green area represents the ATR, angiosperm terrestrial revolution. Red-dashed vertical lines indicate major crises: P–Tr, Permian–Triassic; Tr–J, Triassic–Jurassic; K–Pg, Cretaceous–Paleogene. Time is in millions of years. The color of each geological period in the chronostratigraphic scale follows that of the International Chronostratigraphic Chart (v2024/12). Insect silhouettes are from http://phylopic.org/.

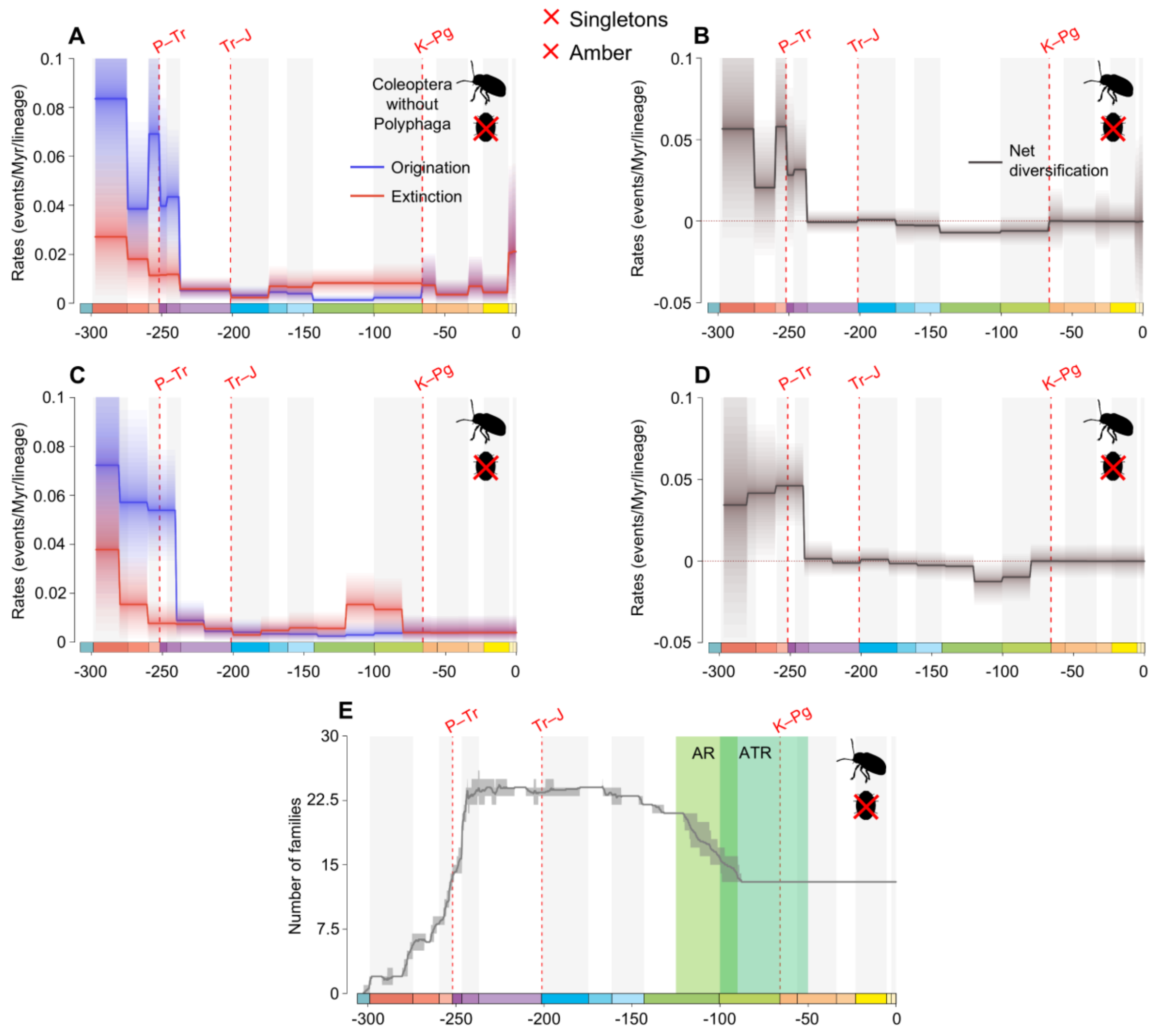


**fig. S53.** Diversification and diversity dynamics of Coleoptera families without Polyphaga, excluding singletons and amber occurrences. Bayesian fossil-based inferences of Coleoptera without Polyphaga origination and extinction rates at the family level under the birth-death model with epochs (A), 20 Ma bins (C) as constrained shifts (BDCS). Net diversification rates for Coleoptera without Polyphaga obtained from the difference between origination and extinction rates (rates above 0 indicate increasing diversity, and rates below 0 indicate declining diversity) per epochs (B), 20 Ma bins (D). Solid lines indicate mean posterior rates and the shaded areas show 95% HPD. Number of families through time computed by summing up the lifespans of all families for Coleoptera without Polyphaga (E). Solid lines indicate mean diversity at each point in time and shaded areas show estimations of different replications that incorporate age uncertainties of fossil occurrences. Light-green area represents the AR, angiosperm radiation, and dark-green area represents the ATR, angiosperm terrestrial revolution. Red-dashed vertical lines indicate major crises: P–Tr, Permian–Triassic; Tr–J, Triassic–Jurassic; K–Pg, Cretaceous–Paleogene. Time is in millions of years. The color of each geological period in the chronostratigraphic scale follows that of the International Chronostratigraphic Chart (v2024/12). Insect silhouettes are from http://phylopic.org/.

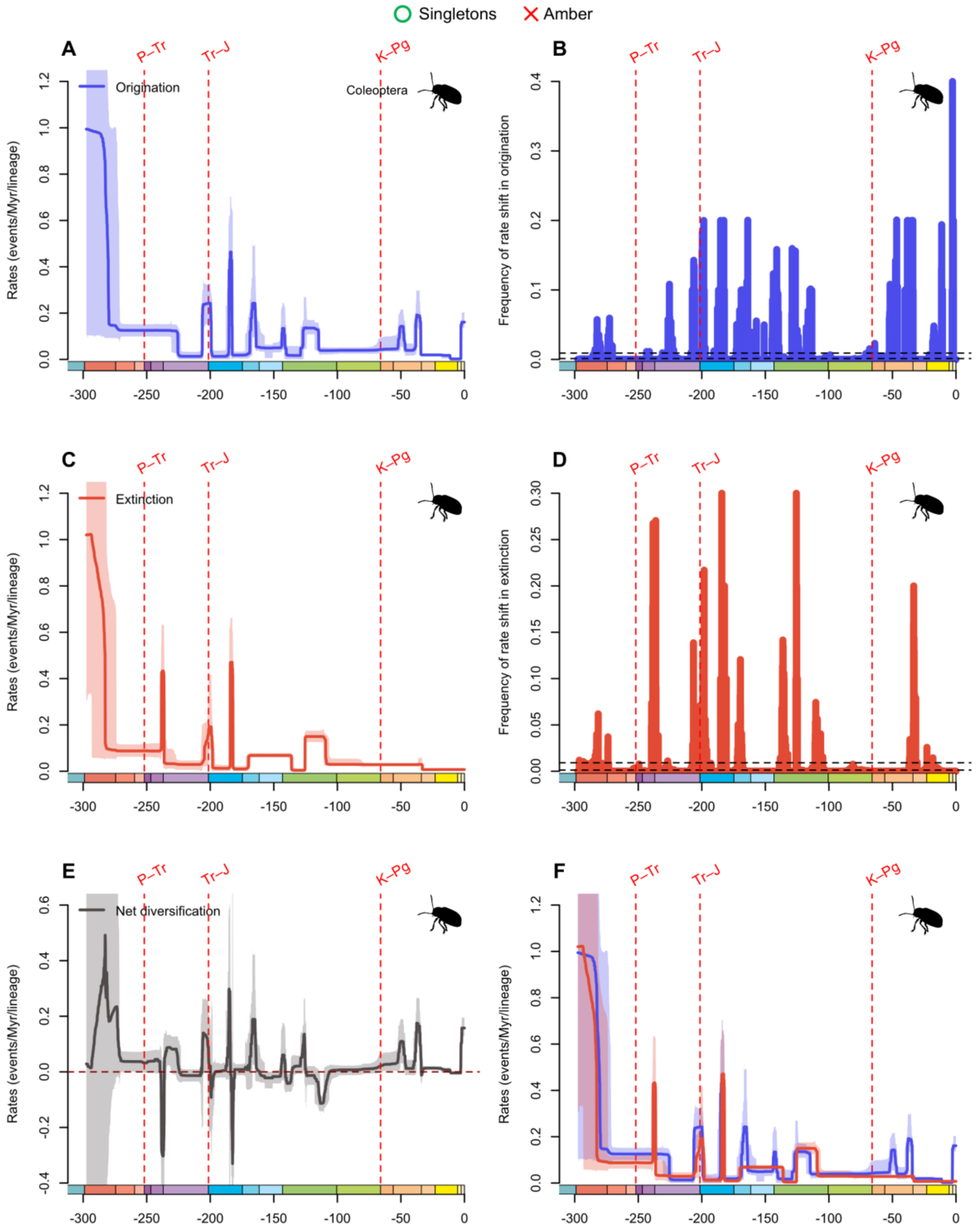


**fig. S54.** Diversification and diversity dynamics of Coleoptera genera, considering singletons and excluding amber occurrences. Bayesian estimations of origination and extinction rates through time as inferred by PyRate using reversible jump Markov Chain Monte Carlo (RJMCMC). Marginal estimates of origination rates (A) and extinction rates (C) through time are shown as mean and 95% HPD (shaded areas). The frequency of a sampled rate shift is computed within small time bins for origination and extinction rates (B and D, respectively), with black-dashed

horizontal lines indicating log-Bayes factors of 2 (bottom) and 6 (top). Sampling frequencies higher than log-Bayes factors = 6 indicate strong statistical support for a rate shift. Panel (E) shows net diversification rates through time (computed as the posterior difference between origination and extinction rates through time). Panel (F) shows both marginal estimates of origination and extinction rates through time on the same plot. Solid lines indicate mean posterior rates and the shaded areas show 95% HPD. Red-dashed vertical lines indicate major crises: P–Tr, Permian–Triassic; Tr–J, Triassic–Jurassic; K–Pg, Cretaceous–Paleogene. Time is in millions of years. The color of each geological period in the chronostratigraphic scale follows that of the International Chronostratigraphic Chart (v2024/12). Insect silhouettes are from http://phylopic.org/.

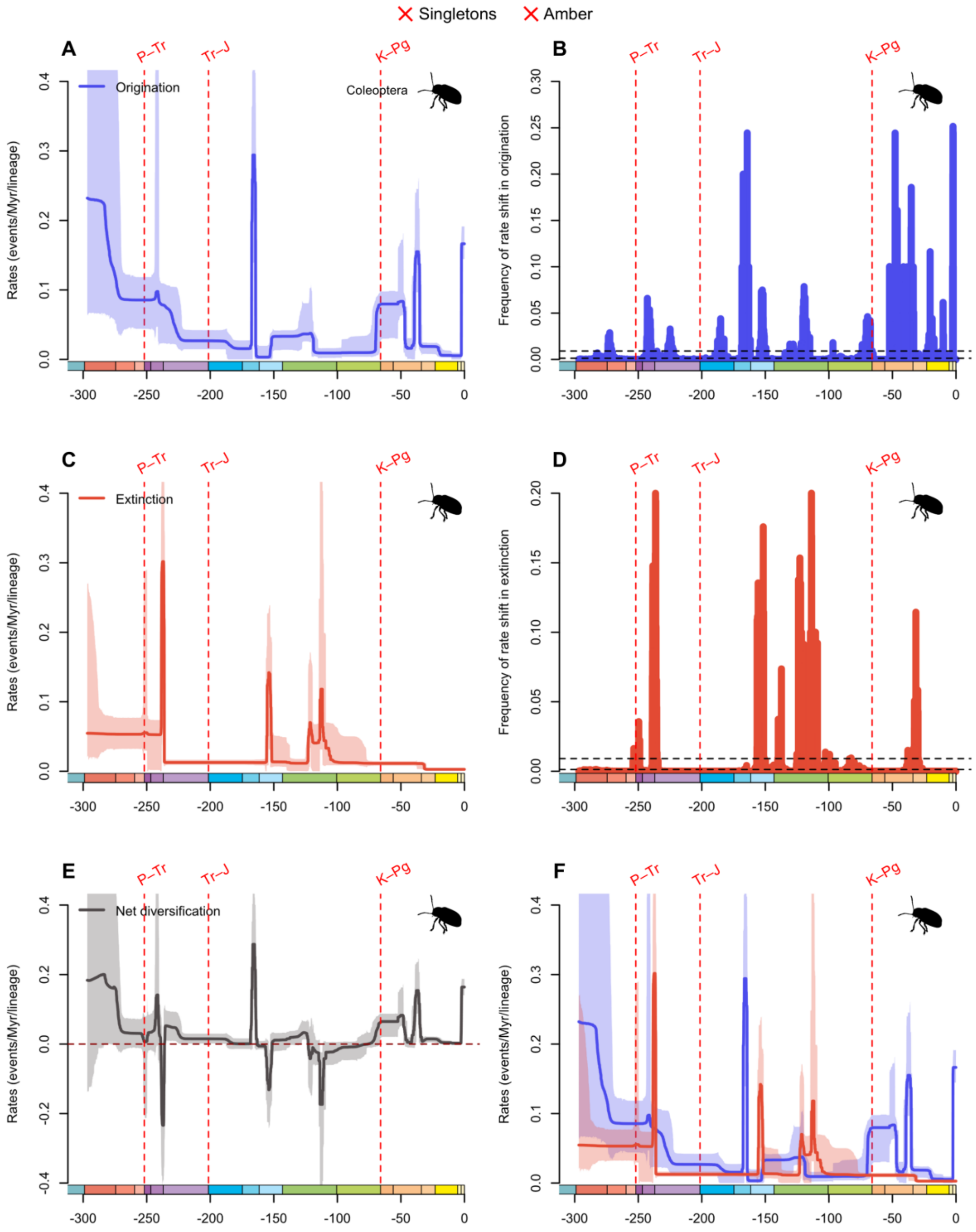


**fig. S55.** Diversification and diversity dynamics of Coleoptera genera, excluding singletons and amber occurrences. Bayesian estimations of origination and extinction rates through time as inferred by PyRate using reversible jump Markov Chain Monte Carlo (RJMCMC). Marginal estimates of origination rates (A) and extinction rates (C) through time are shown as mean and 95% HPD (shaded areas). The frequency of a sampled rate shift is computed within small time bins for origination and extinction rates (B and D, respectively), with black-dashed horizontal lines

indicating log-Bayes factors of 2 (bottom) and 6 (top). Sampling frequencies higher than log-Bayes factors = 6 indicate strong statistical support for a rate shift. Panel (E) shows net diversification rates through time (computed as the posterior difference between origination and extinction rates through time). Panel (F) shows both marginal estimates of origination and extinction rates through time on the same plot. Solid lines indicate mean posterior rates and the shaded areas show 95% HPD. Red-dashed vertical lines indicate major crises: P–Tr, Permian–Triassic; Tr–J, Triassic–Jurassic; K–Pg, Cretaceous–Paleogene. Time is in millions of years. The color of each geological period in the chronostratigraphic scale follows that of the International Chronostratigraphic Chart (v2024/12). Insect silhouettes are from http://phylopic.org/.

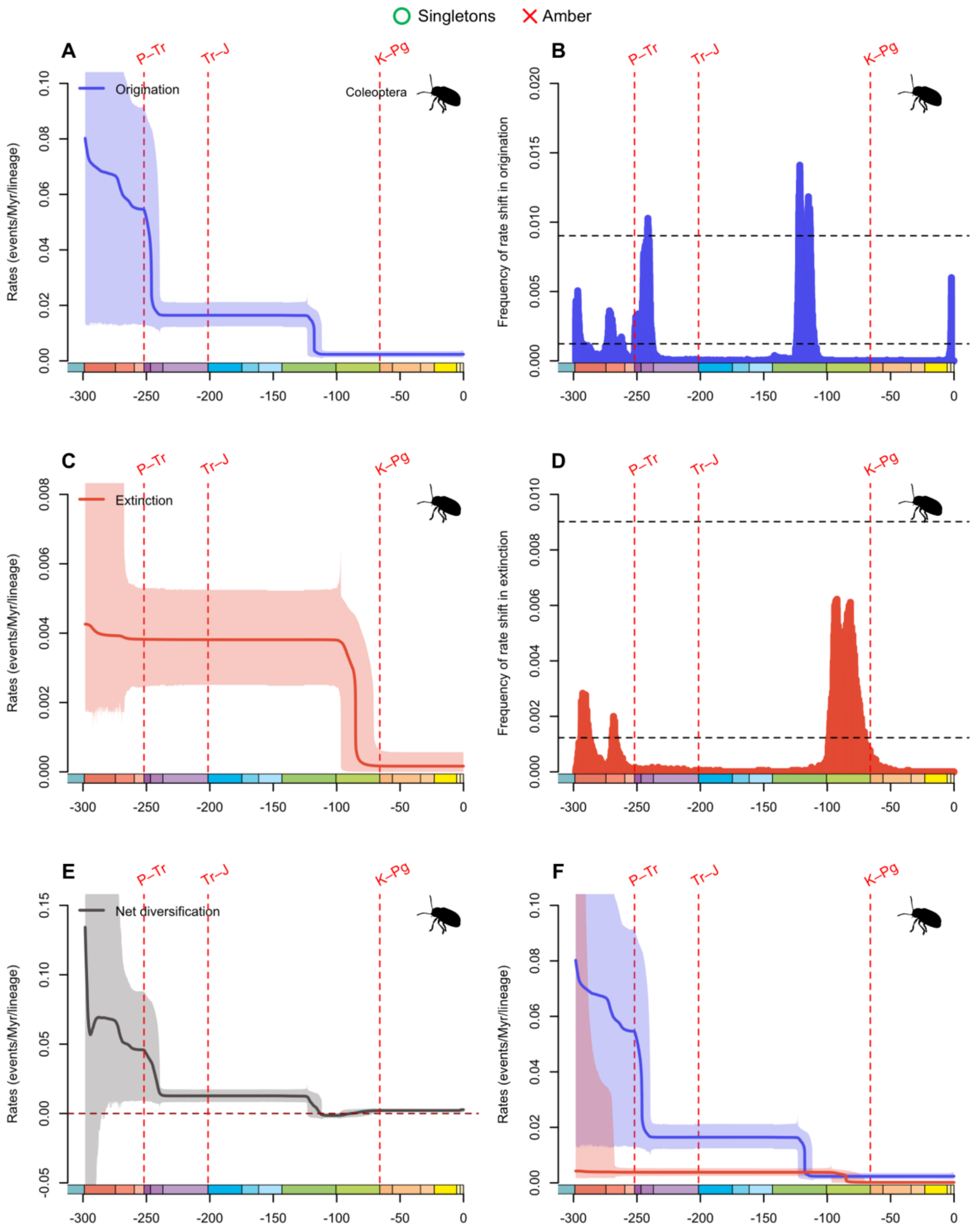


**fig. S56.** Diversification and diversity dynamics of Coleoptera families, considering singletons and excluding amber occurrences. Bayesian estimations of origination and extinction rates through time as inferred by PyRate using reversible jump Markov Chain Monte Carlo (RJMCMC). Marginal estimates of origination rates (A) and extinction rates (C) through time are shown as mean and 95% HPD (shaded areas). The frequency of a sampled rate shift is computed within small time bins for origination and extinction rates (B and D, respectively), with black-dashed

horizontal lines indicating log-Bayes factors of 2 (bottom) and 6 (top). Sampling frequencies higher than log-Bayes factors = 6 indicate strong statistical support for a rate shift. Panel (E) shows net diversification rates through time (computed as the posterior difference between origination and extinction rates through time). Panel (F) shows both marginal estimates of origination and extinction rates through time on the same plot. Solid lines indicate mean posterior rates and the shaded areas show 95% HPD. Red-dashed vertical lines indicate major crises: P–Tr, Permian–Triassic; Tr–J, Triassic–Jurassic; K–Pg, Cretaceous–Paleogene. Time is in millions of years. The color of each geological period in the chronostratigraphic scale follows that of the International Chronostratigraphic Chart (v2024/12). Insect silhouettes are from http://phylopic.org/.

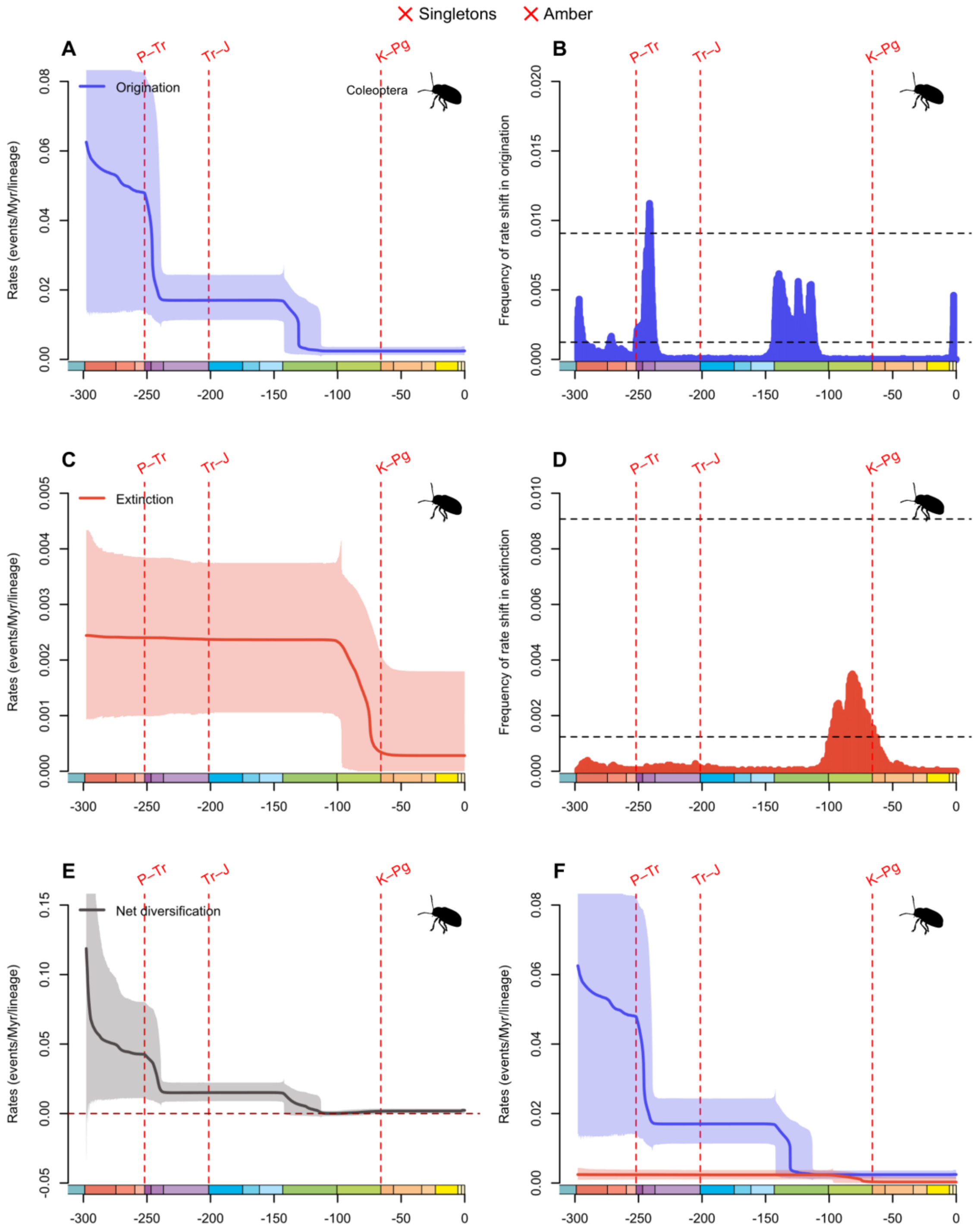


**fig. S57.** Diversification and diversity dynamics of Coleoptera families, excluding singletons and amber occurrences. Bayesian estimations of origination and extinction rates through time as inferred by PyRate using reversible jump Markov Chain Monte Carlo (RJMCMC). Marginal estimates of origination rates (A) and extinction rates (C) through time are shown as mean and 95% HPD (shaded areas). The frequency of a sampled rate shift is computed within small time bins for origination and extinction rates (B and D, respectively), with black-dashed horizontal lines

indicating log-Bayes factors of 2 (bottom) and 6 (top). Sampling frequencies higher than log-Bayes factors = 6 indicate strong statistical support for a rate shift. Panel (E) shows net diversification rates through time (computed as the posterior difference between origination and extinction rates through time). Panel (F) shows both marginal estimates of origination and extinction rates through time on the same plot. Solid lines indicate mean posterior rates and the shaded areas show 95% HPD. Red-dashed vertical lines indicate major crises: P–Tr, Permian–Triassic; Tr–J, Triassic–Jurassic; K–Pg, Cretaceous–Paleogene. Time is in millions of years. The color of each geological period in the chronostratigraphic scale follows that of the International Chronostratigraphic Chart (v2024/12). Insect silhouettes are from http://phylopic.org/.

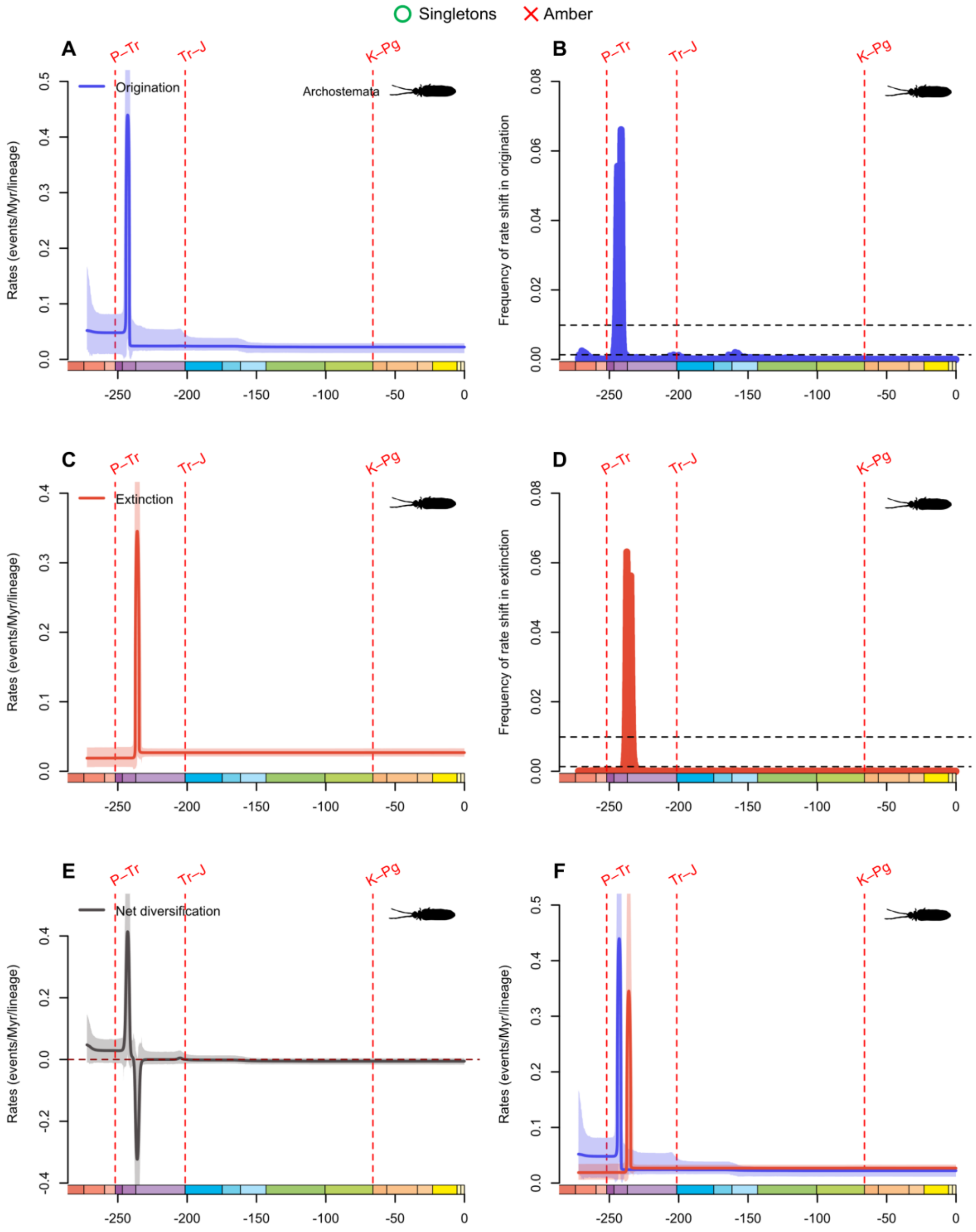


**fig. S58.** Diversification and diversity dynamics of Archostemata genera, considering singletons and excluding amber occurrences. Bayesian estimations of origination and extinction rates through time as inferred by PyRate using reversible jump Markov Chain Monte Carlo (RJMCMC). Marginal estimates of origination rates (A) and extinction rates (C) through time are shown as mean and 95% HPD (shaded areas). The frequency of a sampled rate shift is computed within small time bins for origination and extinction rates (B and D, respectively), with black-dashed

horizontal lines indicating log-Bayes factors of 2 (bottom) and 6 (top). Sampling frequencies higher than log-Bayes factors = 6 indicate strong statistical support for a rate shift. Panel (E) shows net diversification rates through time (computed as the posterior difference between origination and extinction rates through time). Panel (F) shows both marginal estimates of origination and extinction rates through time on the same plot. Solid lines indicate mean posterior rates and the shaded areas show 95% HPD. Red-dashed vertical lines indicate major crises: P–Tr, Permian–Triassic; Tr–J, Triassic–Jurassic; K–Pg, Cretaceous–Paleogene. Time is in millions of years. The color of each geological period in the chronostratigraphic scale follows that of the International Chronostratigraphic Chart (v2024/12). Insect silhouettes are from http://phylopic.org/.

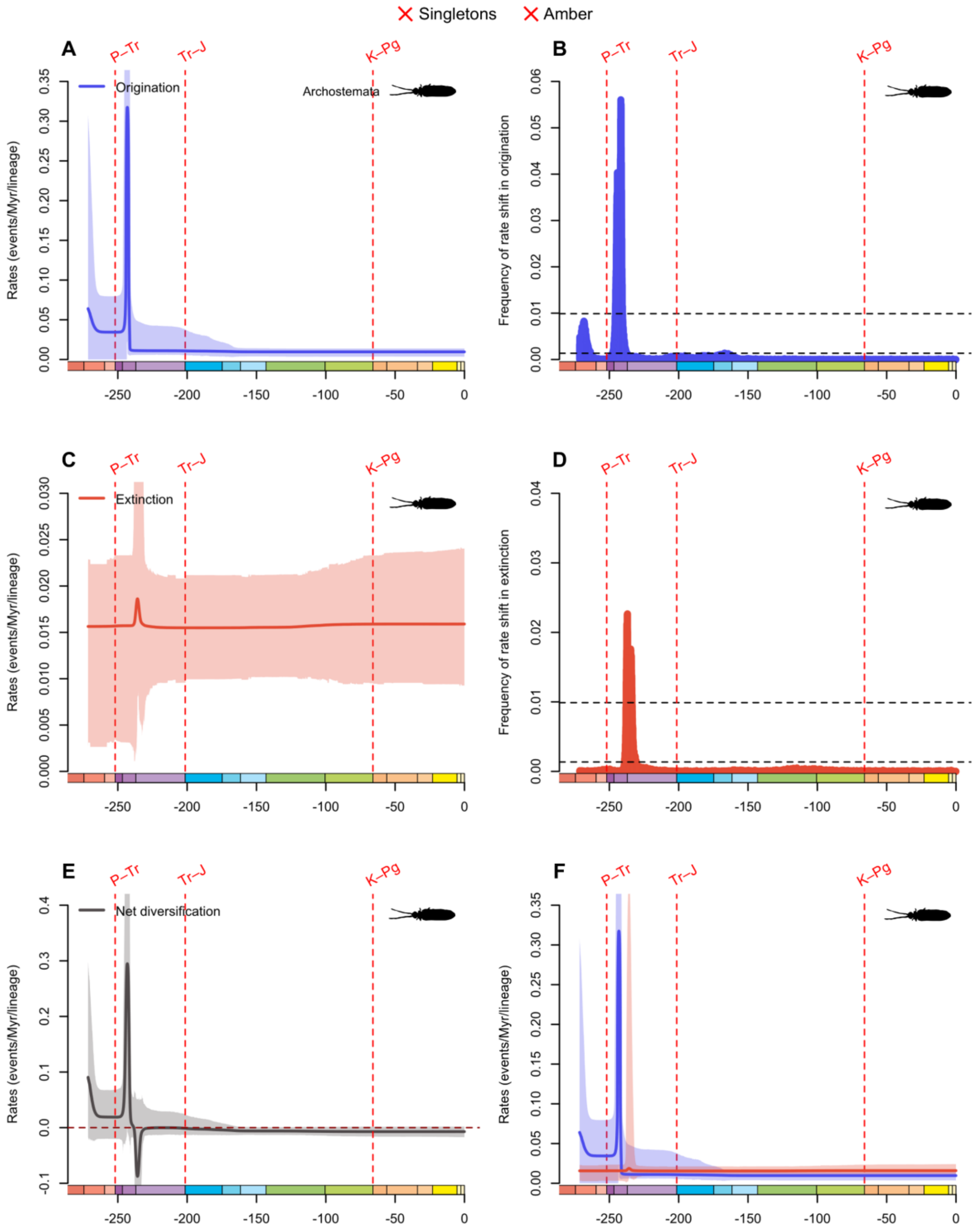


**fig. S59.** Diversification and diversity dynamics of Archostemata genera, excluding singletons and amber occurrences. Bayesian estimations of origination and extinction rates through time as inferred by PyRate using reversible jump Markov Chain Monte Carlo (RJMCMC). Marginal estimates of origination rates (A) and extinction rates (C) through time are shown as mean and 95% HPD (shaded areas). The frequency of a sampled rate shift is computed within small time bins for origination and extinction rates (B and D, respectively), with black-dashed horizontal lines

indicating log-Bayes factors of 2 (bottom) and 6 (top). Sampling frequencies higher than log-Bayes factors = 6 indicate strong statistical support for a rate shift. Panel (E) shows net diversification rates through time (computed as the posterior difference between origination and extinction rates through time). Panel (F) shows both marginal estimates of origination and extinction rates through time on the same plot. Solid lines indicate mean posterior rates and the shaded areas show 95% HPD. Red-dashed vertical lines indicate major crises: P–Tr, Permian–Triassic; Tr–J, Triassic–Jurassic; K–Pg, Cretaceous–Paleogene. Time is in millions of years. The color of each geological period in the chronostratigraphic scale follows that of the International Chronostratigraphic Chart (v2024/12). Insect silhouettes are from http://phylopic.org/.

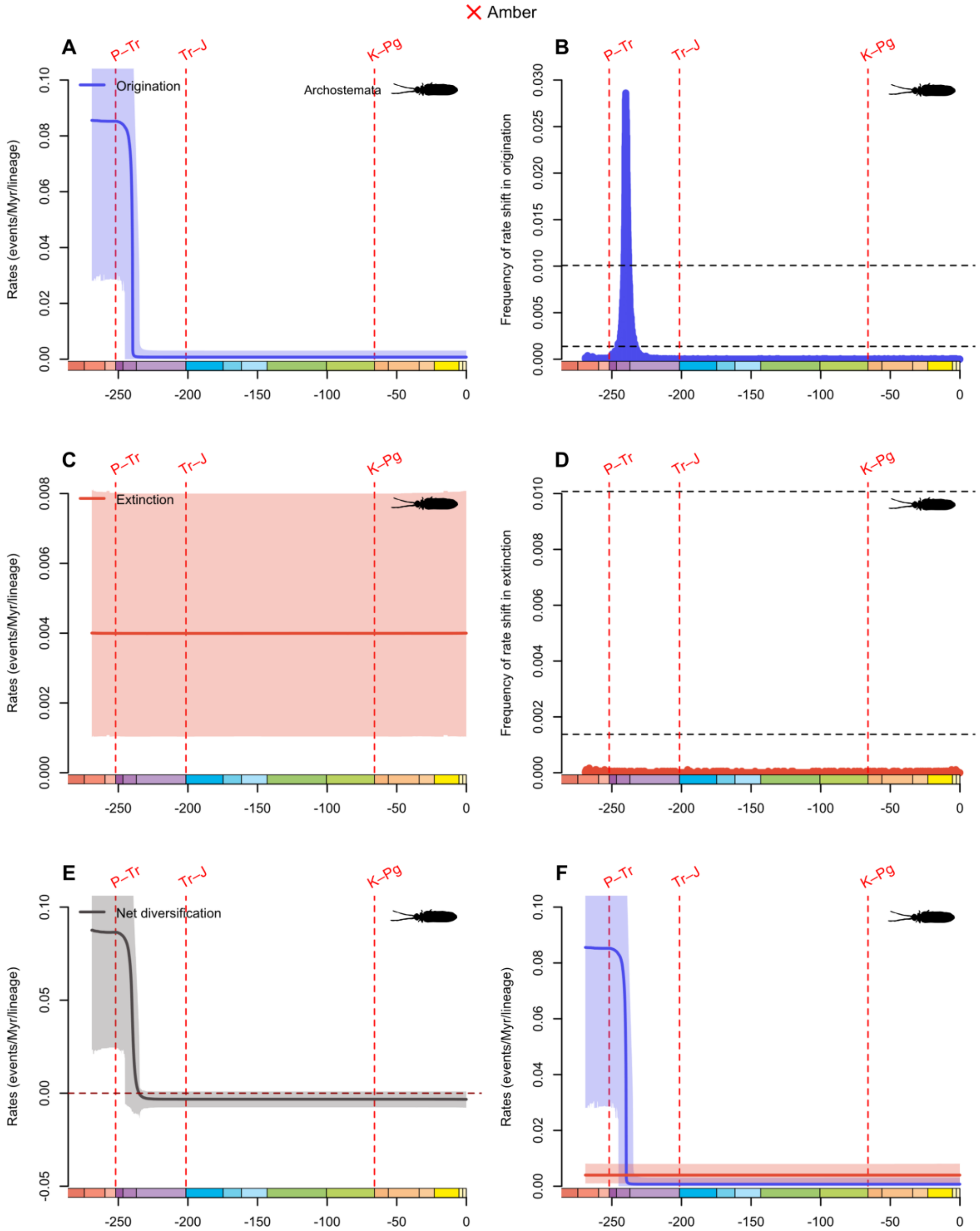


**fig. S60.** Diversification and diversity dynamics of Archostemata families, excluding amber occurrences. There are no singletons at the family-level in Archostemata. Bayesian estimations of origination and extinction rates through time as inferred by PyRate using reversible jump Markov Chain Monte Carlo (RJMCMC). Marginal estimates of origination rates (A) and extinction rates (C) through time are shown as mean and 95% HPD (shaded areas). The frequency of a sampled rate shift is computed within small time bins for origination and extinction rates (B and D, respectively), with black-dashed horizontal lines indicating log-Bayes factors of 2 (bottom) and 6 (top). Sampling frequencies higher than log-Bayes factors = 6 indicate strong statistical support for a rate shift. Panel (E) shows net diversification rates through time (computed as the posterior difference between origination and extinction rates through time). Panel (F) shows both marginal estimates of origination and extinction rates through time on the same plot. Solid lines indicate mean posterior rates and the shaded areas show 95% HPD. Red-dashed vertical lines indicate major crises: P–Tr, Permian–Triassic; Tr–J, Triassic–Jurassic; K–Pg, Cretaceous–Paleogene. Time is in millions of years. The color of each geological period in the chronostratigraphic scale follows that of the International Chronostratigraphic Chart (v2024/12). Insect silhouettes are from http://phylopic.org/.

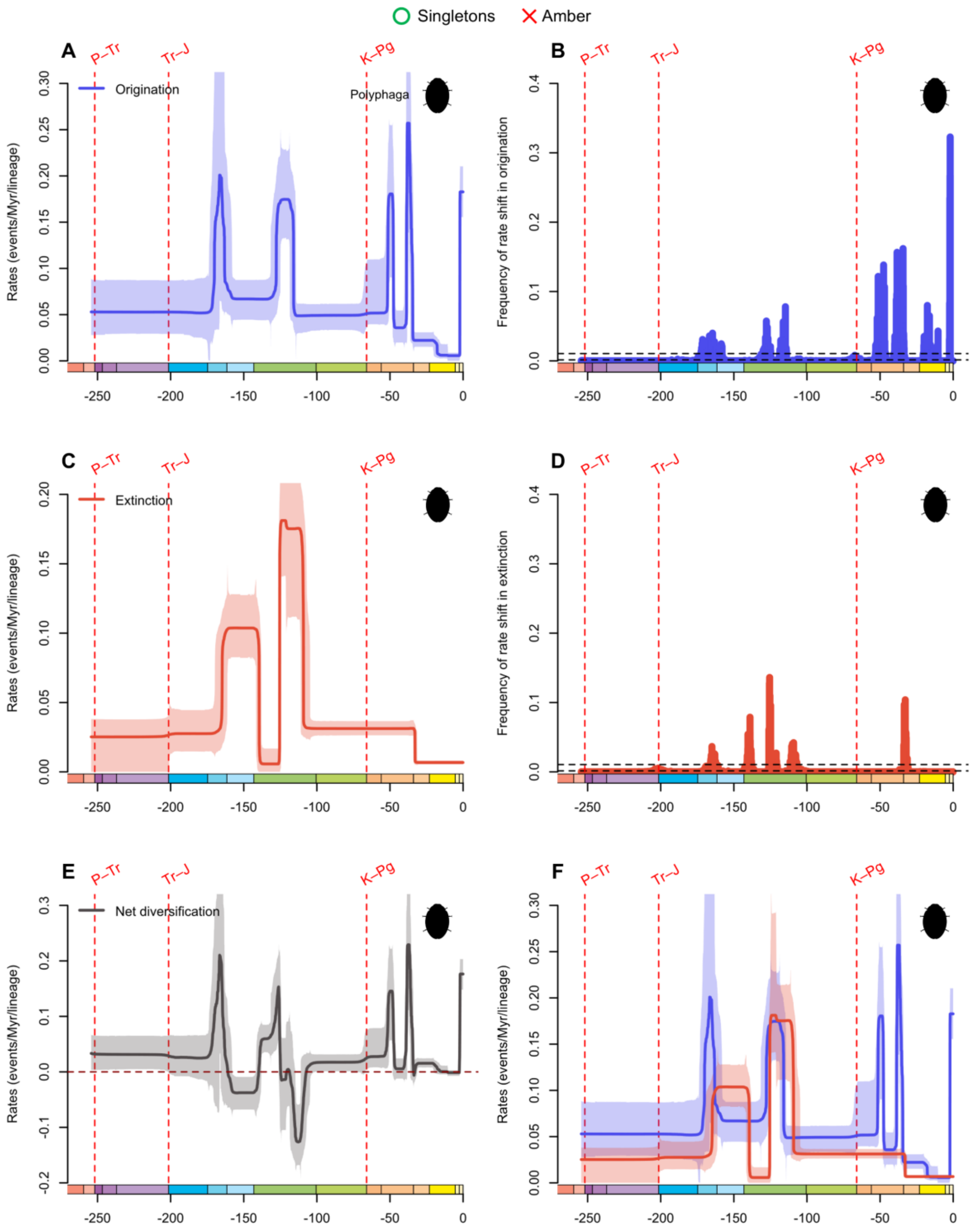


**fig. S61.** Diversification and diversity dynamics of Polyphaga genera, considering singletons and excluding amber occurrences. Bayesian estimations of origination and extinction rates through time as inferred by PyRate using reversible jump Markov Chain Monte Carlo (RJMCMC). Marginal estimates of origination rates (A) and extinction rates (C) through time are shown as mean and 95% HPD (shaded areas). The frequency of a sampled rate shift is computed within small time bins for origination and extinction rates (B and D, respectively), with black-dashed

horizontal lines indicating log-Bayes factors of 2 (bottom) and 6 (top). Sampling frequencies higher than log-Bayes factors = 6 indicate strong statistical support for a rate shift. Panel (E) shows net diversification rates through time (computed as the posterior difference between origination and extinction rates through time). Panel (F) shows both marginal estimates of origination and extinction rates through time on the same plot. Solid lines indicate mean posterior rates and the shaded areas show 95% HPD. Red-dashed vertical lines indicate major crises: P–Tr, Permian–Triassic; Tr–J, Triassic–Jurassic; K–Pg, Cretaceous–Paleogene. Time is in millions of years. The color of each geological period in the chronostratigraphic scale follows that of the International Chronostratigraphic Chart (v2024/12). Insect silhouettes are from http://phylopic.org/.

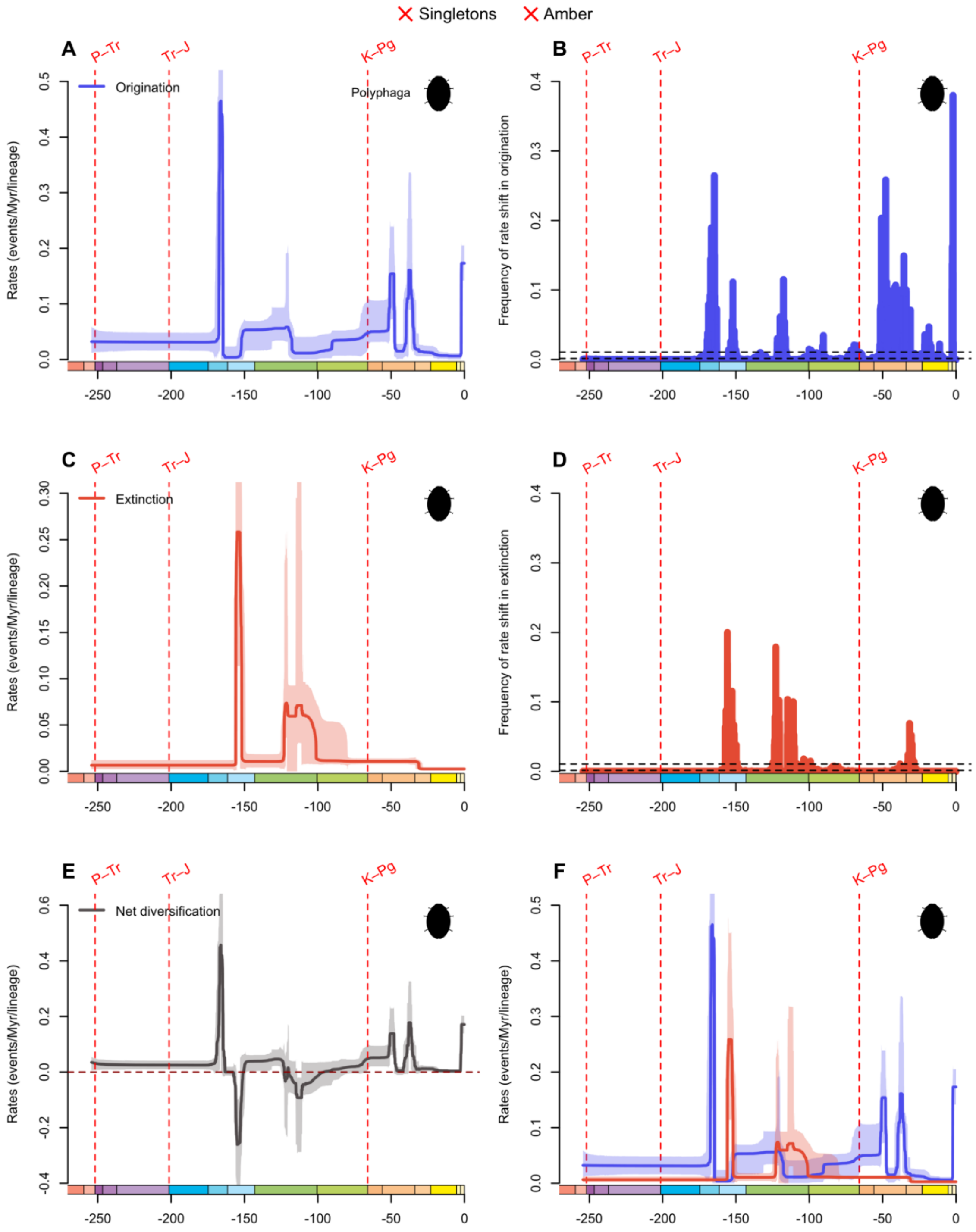


**fig. S62.** Diversification and diversity dynamics of Polyphaga genera, excluding singletons and amber occurrences. Bayesian estimations of origination and extinction rates through time as inferred by PyRate using reversible jump Markov Chain Monte Carlo (RJMCMC). Marginal estimates of origination rates (A) and extinction rates (C) through time are shown as mean and 95% HPD (shaded areas). The frequency of a sampled rate shift is computed within small time bins for origination and extinction rates (B and D, respectively), with black-dashed horizontal lines

indicating log-Bayes factors of 2 (bottom) and 6 (top). Sampling frequencies higher than log-Bayes factors = 6 indicate strong statistical support for a rate shift. Panel (E) shows net diversification rates through time (computed as the posterior difference between origination and extinction rates through time). Panel (F) shows both marginal estimates of origination and extinction rates through time on the same plot. Solid lines indicate mean posterior rates and the shaded areas show 95% HPD. Red-dashed vertical lines indicate major crises: P–Tr, Permian–Triassic; Tr–J, Triassic–Jurassic; K–Pg, Cretaceous–Paleogene. Time is in millions of years. The color of each geological period in the chronostratigraphic scale follows that of the International Chronostratigraphic Chart (v2024/12). Insect silhouettes are from http://phylopic.org/.

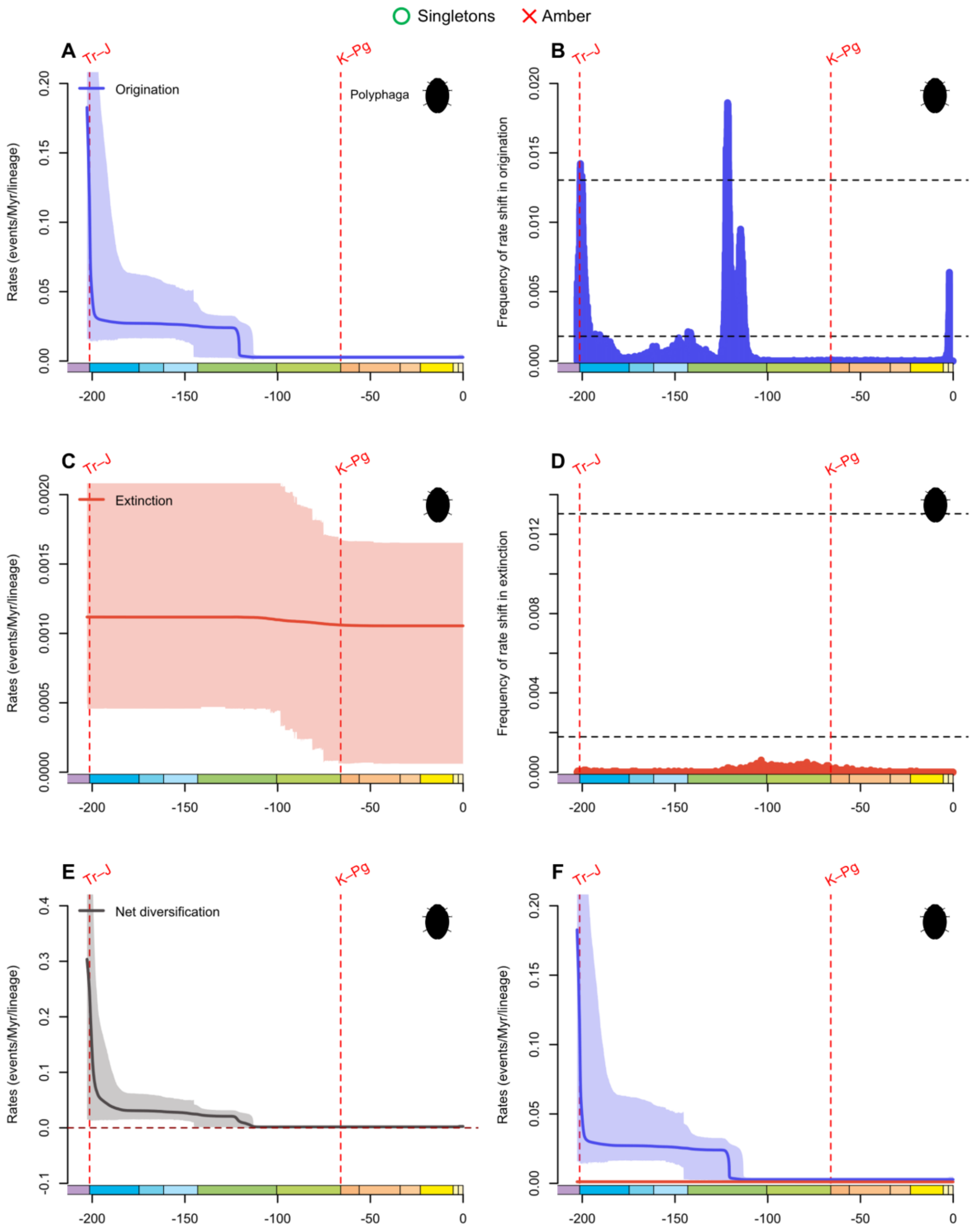


**fig. S63.** Diversification and diversity dynamics of Polyphaga families, considering singletons and excluding amber occurrences. Bayesian estimations of origination and extinction rates through time as inferred by PyRate using reversible jump Markov Chain Monte Carlo (RJMCMC). Marginal estimates of origination rates (A) and extinction rates (C) through time are shown as mean and 95% HPD (shaded areas). The frequency of a sampled rate shift is computed within small time bins for origination and extinction rates (B and D, respectively), with black-dashed horizontal lines indicating log-Bayes factors of 2 (bottom) and 6 (top). Sampling frequencies higher than log-Bayes factors = 6 indicate strong statistical support for a rate shift. Panel (E) shows net diversification rates through time (computed as the posterior difference between origination and extinction rates through time). Panel (F) shows both marginal estimates of origination and extinction rates through time on the same plot. Solid lines indicate mean posterior rates and the shaded areas show 95% HPD. Red-dashed vertical lines indicate major crises: Tr–J, Triassic–Jurassic; K–Pg, Cretaceous–Paleogene. Time is in millions of years. The color of each geological period in the chronostratigraphic scale follows that of the International Chronostratigraphic Chart (v2024/12). Insect silhouettes are from http://phylopic.org/.

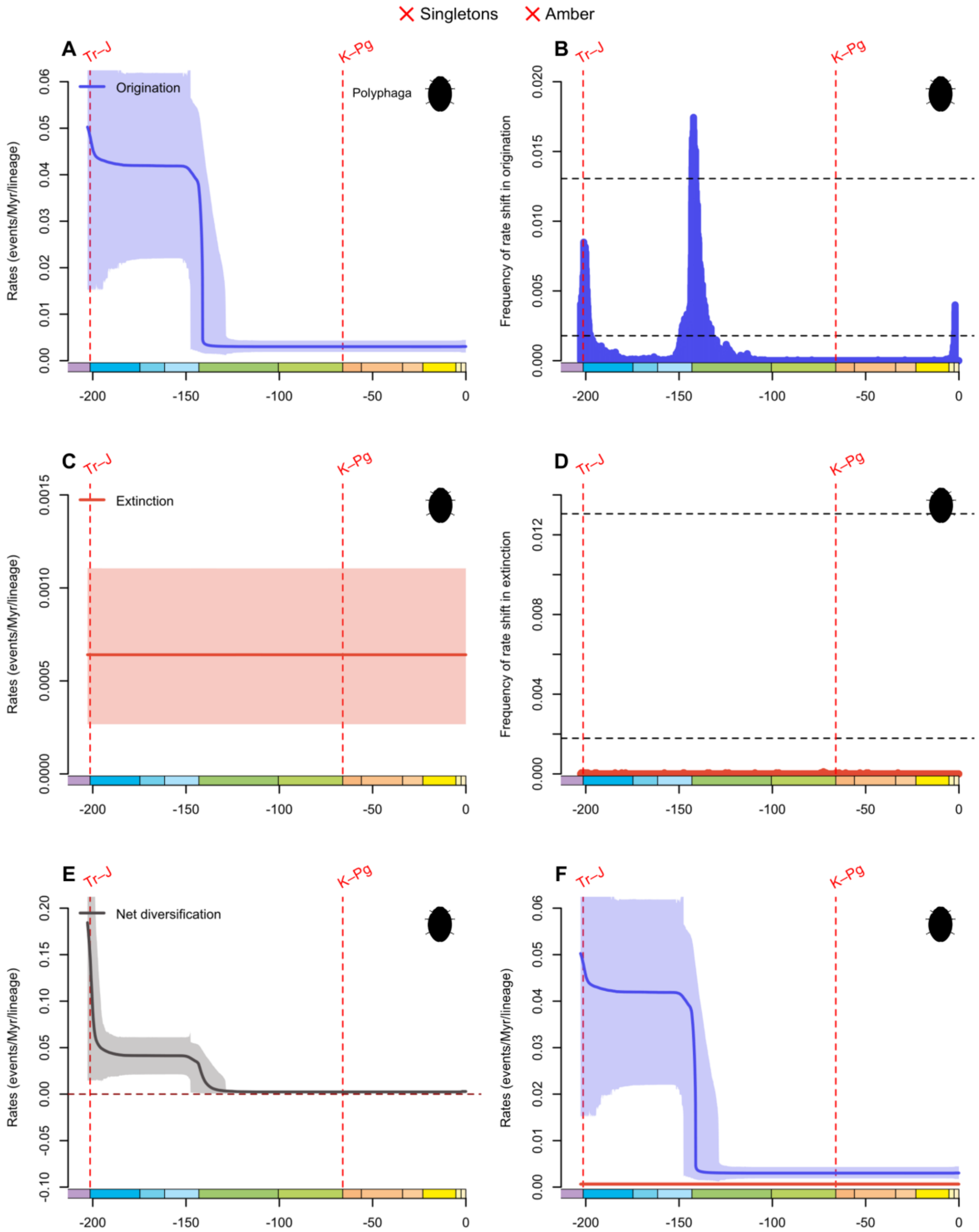


**fig. S64.** Diversification and diversity dynamics of Polyphaga families, excluding singletons and amber occurrences. Bayesian estimations of origination and extinction rates through time as inferred by PyRate using reversible jump Markov Chain Monte Carlo (RJMCMC). Marginal estimates of origination rates (A) and extinction rates (C) through time are shown as mean and 95% HPD (shaded areas). The frequency of a sampled rate shift is computed within small time bins for origination and extinction rates (B and D, respectively), with black-dashed horizontal lines indicating log-Bayes factors of 2 (bottom) and 6 (top). Sampling frequencies higher than log-Bayes factors = 6 indicate strong statistical support for a rate shift. Panel (E) shows net diversification rates through time (computed as the posterior difference between origination and extinction rates through time). Panel (F) shows both marginal estimates of origination and extinction rates through time on the same plot. Solid lines indicate mean posterior rates and the shaded areas show 95% HPD. Red-dashed vertical lines indicate major crises: Tr–J, Triassic–Jurassic; K–Pg, Cretaceous–Paleogene. Time is in millions of years. The color of each geological period in the chronostratigraphic scale follows that of the International Chronostratigraphic Chart (v2024/12). Insect silhouettes are from http://phylopic.org/.

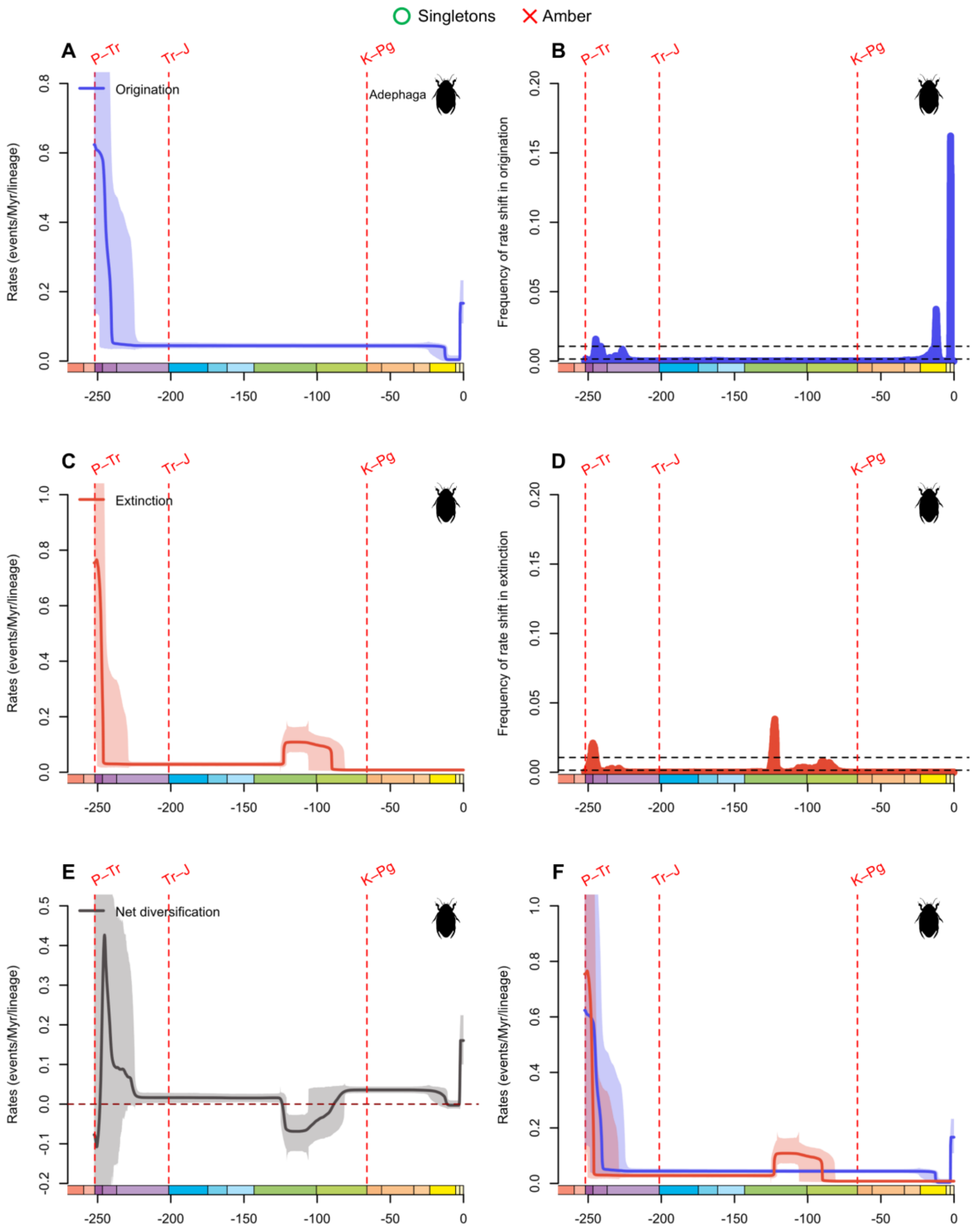


**fig. S65.** Diversification and diversity dynamics of Adephaga genera, considering singletons and excluding amber occurrences. Bayesian estimations of origination and extinction rates through time as inferred by PyRate using reversible jump Markov Chain Monte Carlo (RJMCMC). Marginal estimates of origination rates (A) and extinction rates (C) through time are shown as mean and 95% HPD (shaded areas). The frequency of a sampled rate shift is computed within small time bins for origination and extinction rates (B and D, respectively), with black-dashed

horizontal lines indicating log-Bayes factors of 2 (bottom) and 6 (top). Sampling frequencies higher than log-Bayes factors = 6 indicate strong statistical support for a rate shift. Panel (E) shows net diversification rates through time (computed as the posterior difference between origination and extinction rates through time). Panel (F) shows both marginal estimates of origination and extinction rates through time on the same plot. Solid lines indicate mean posterior rates and the shaded areas show 95% HPD. Red-dashed vertical lines indicate major crises: P–Tr, Permian–Triassic; Tr–J, Triassic–Jurassic; K–Pg, Cretaceous–Paleogene. Time is in millions of years. The color of each geological period in the chronostratigraphic scale follows that of the International Chronostratigraphic Chart (v2024/12). Insect silhouettes are from http://phylopic.org/.

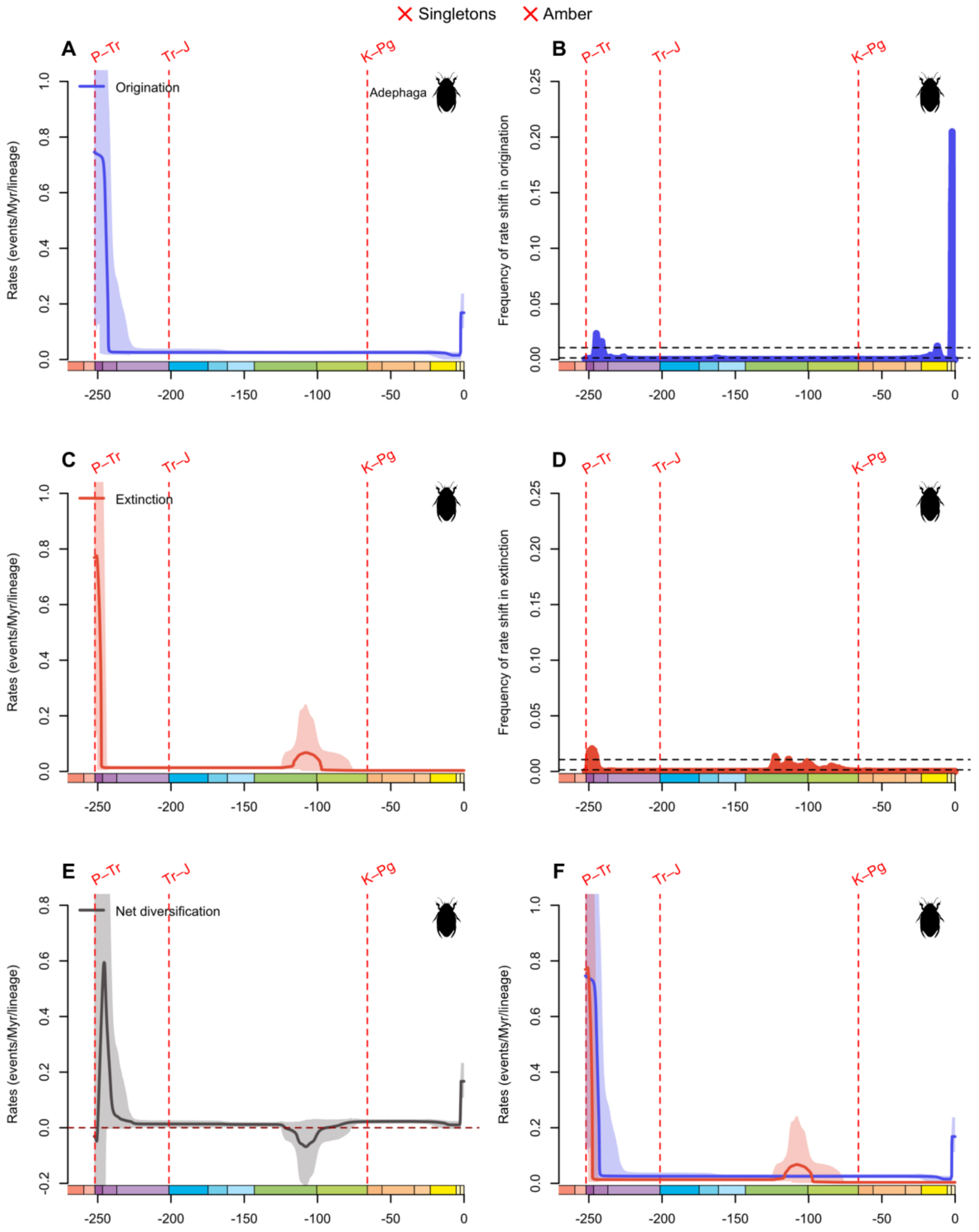


**fig. S66.** Diversification and diversity dynamics of Adephaga genera, excluding singletons and amber occurrences. Bayesian estimations of origination and extinction rates through time as inferred by PyRate using reversible jump Markov Chain Monte Carlo (RJMCMC). Marginal estimates of origination rates (A) and extinction rates (C) through time are shown as mean and 95% HPD (shaded areas). The frequency of a sampled rate shift is computed within small time bins for origination and extinction rates (B and D, respectively), with black-dashed horizontal lines

indicating log-Bayes factors of 2 (bottom) and 6 (top). Sampling frequencies higher than log-Bayes factors = 6 indicate strong statistical support for a rate shift. Panel (E) shows net diversification rates through time (computed as the posterior difference between origination and extinction rates through time). Panel (F) shows both marginal estimates of origination and extinction rates through time on the same plot. Solid lines indicate mean posterior rates and the shaded areas show 95% HPD. Red-dashed vertical lines indicate major crises: P–Tr, Permian–Triassic; Tr–J, Triassic–Jurassic; K–Pg, Cretaceous–Paleogene. Time is in millions of years. The color of each geological period in the chronostratigraphic scale follows that of the International Chronostratigraphic Chart (v2024/12). Insect silhouettes are from http://phylopic.org/.

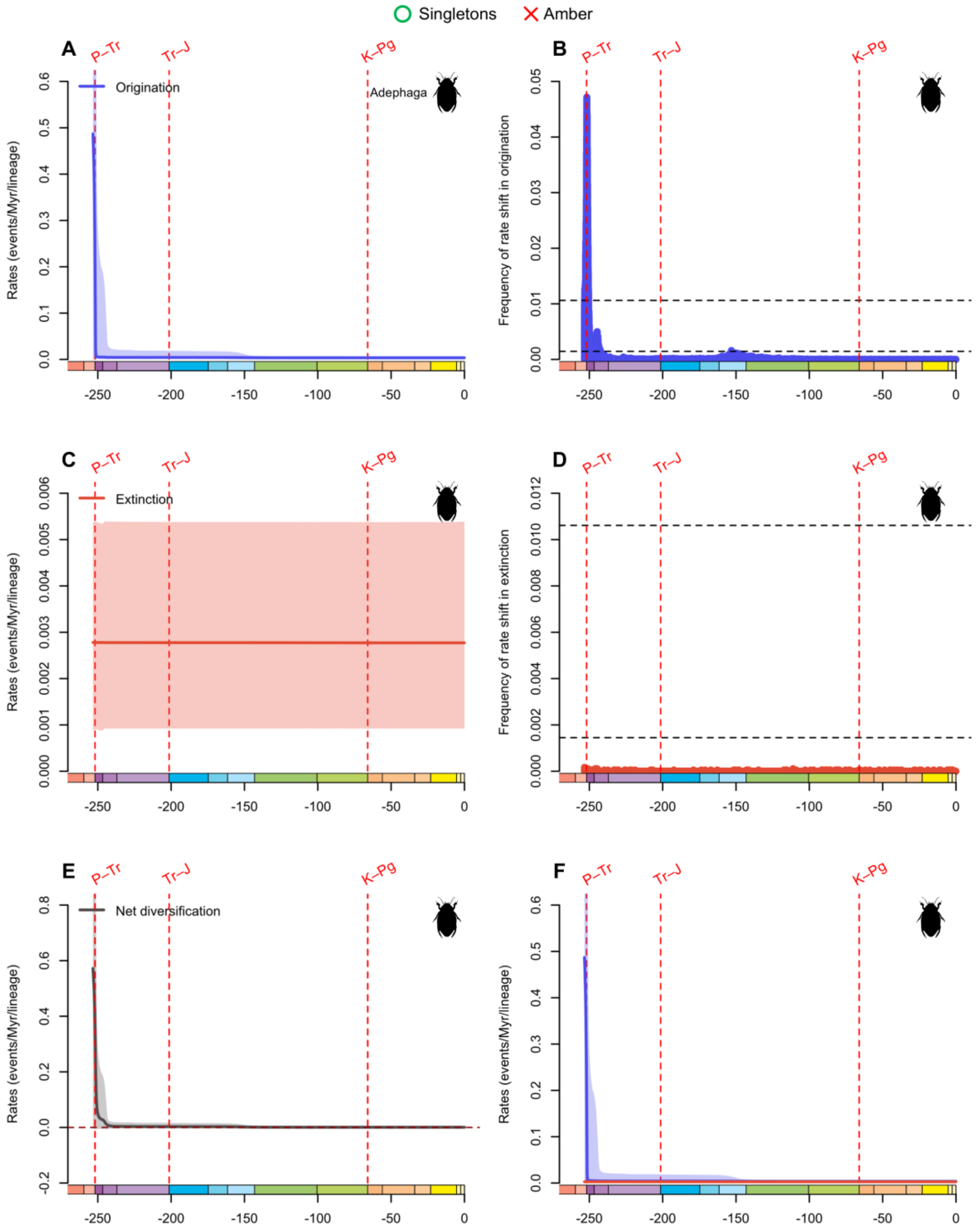


**fig. S67.** Diversification and diversity dynamics of Adephaga families, considering singletons and excluding amber occurrences. Bayesian estimations of origination and extinction rates through time as inferred by PyRate using reversible jump Markov Chain Monte Carlo (RJMCMC). Marginal estimates of origination rates (A) and extinction rates (C) through time are shown as mean and 95% HPD (shaded areas). The frequency of a sampled rate shift is computed within small time bins for origination and extinction rates (B and D, respectively), with black-dashed

horizontal lines indicating log-Bayes factors of 2 (bottom) and 6 (top). Sampling frequencies higher than log-Bayes factors = 6 indicate strong statistical support for a rate shift. Panel (E) shows net diversification rates through time (computed as the posterior difference between origination and extinction rates through time). Panel (F) shows both marginal estimates of origination and extinction rates through time on the same plot. Solid lines indicate mean posterior rates and the shaded areas show 95% HPD. Red-dashed vertical lines indicate major crises: P–Tr, Permian–Triassic; Tr–J, Triassic–Jurassic; K–Pg, Cretaceous–Paleogene. Time is in millions of years. The color of each geological period in the chronostratigraphic scale follows that of the International Chronostratigraphic Chart (v2024/12). Insect silhouettes are from http://phylopic.org/.

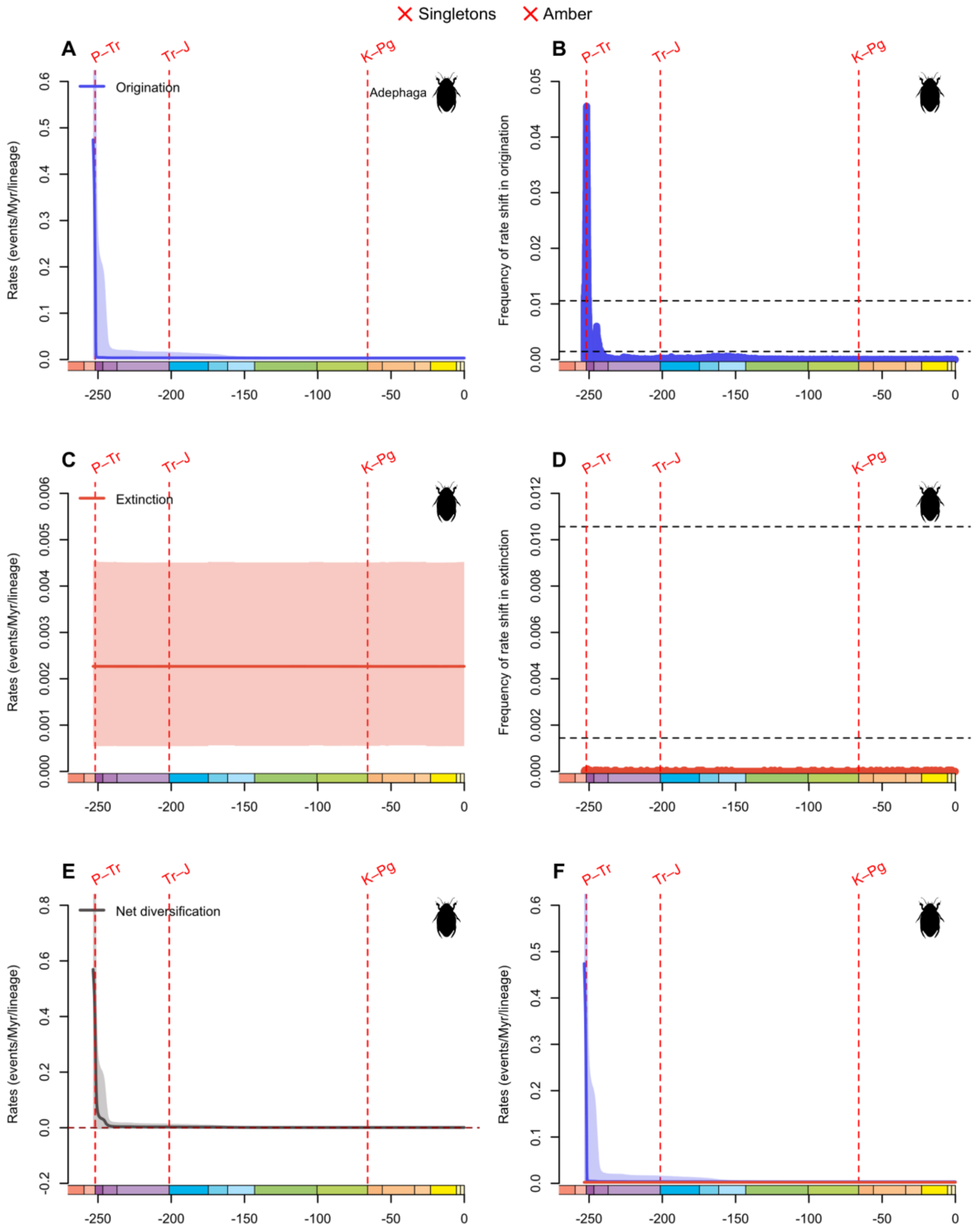


**fig. S68.** Diversification and diversity dynamics of Adephaga families, excluding singletons and amber occurrences. Bayesian estimations of origination and extinction rates through time as inferred by PyRate using reversible jump Markov Chain Monte Carlo (RJMCMC). Marginal estimates of origination rates (A) and extinction rates (C) through time are shown as mean and 95% HPD (shaded areas). The frequency of a sampled rate shift is computed within small time bins for origination and extinction rates (B and D, respectively), with black-dashed horizontal lines

indicating log-Bayes factors of 2 (bottom) and 6 (top). Sampling frequencies higher than log-Bayes factors = 6 indicate strong statistical support for a rate shift. Panel (E) shows net diversification rates through time (computed as the posterior difference between origination and extinction rates through time). Panel (F) shows both marginal estimates of origination and extinction rates through time on the same plot. Solid lines indicate mean posterior rates and the shaded areas show 95% HPD. Red-dashed vertical lines indicate major crises: P–Tr, Permian–Triassic; Tr–J, Triassic–Jurassic; K–Pg, Cretaceous–Paleogene. Time is in millions of years. The color of each geological period in the chronostratigraphic scale follows that of the International Chronostratigraphic Chart (v2024/12). Insect silhouettes are from http://phylopic.org/.

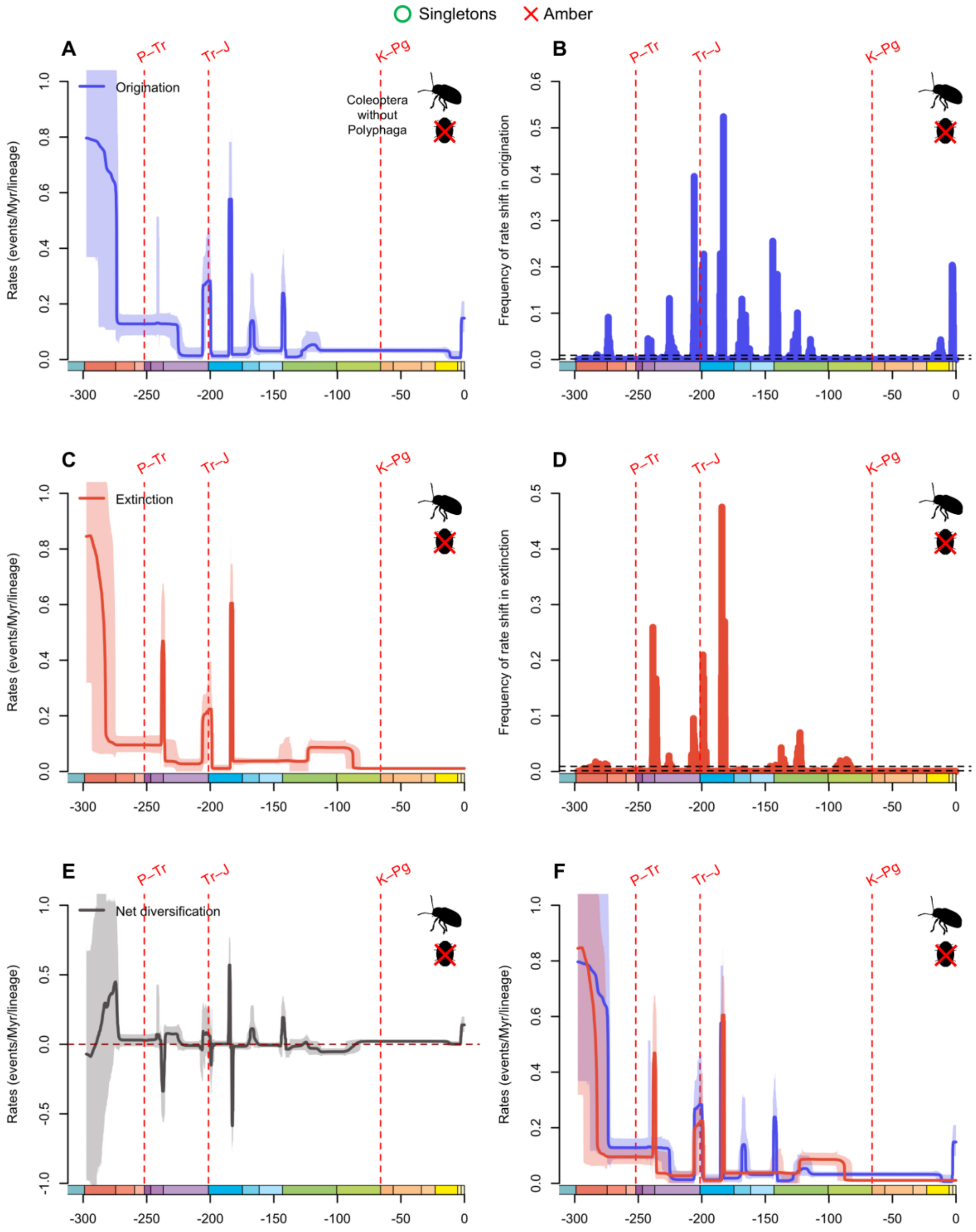


**fig. S69.** Diversification and diversity dynamics of Coleoptera genera without Polyphaga, considering singletons and excluding amber occurrences. Bayesian estimations of origination and extinction rates through time as inferred by PyRate using reversible jump Markov Chain Monte Carlo (RJMCMC). Marginal estimates of origination rates (A) and extinction rates (C) through time are shown as mean and 95% HPD (shaded areas). The frequency of a sampled rate shift is computed within small time bins for origination and extinction rates (B and D, respectively), with

black-dashed horizontal lines indicating log-Bayes factors of 2 (bottom) and 6 (top). Sampling frequencies higher than log-Bayes factors = 6 indicate strong statistical support for a rate shift. Panel (E) shows net diversification rates through time (computed as the posterior difference between origination and extinction rates through time). Panel (F) shows both marginal estimates of origination and extinction rates through time on the same plot. Solid lines indicate mean posterior rates and the shaded areas show 95% HPD. Red-dashed vertical lines indicate major crises: P–Tr, Permian–Triassic; Tr–J, Triassic–Jurassic; K–Pg, Cretaceous–Paleogene. Time is in millions of years. The color of each geological period in the chronostratigraphic scale follows that of the International Chronostratigraphic Chart (v2024/12). Insect silhouettes are from http://phylopic.org/.

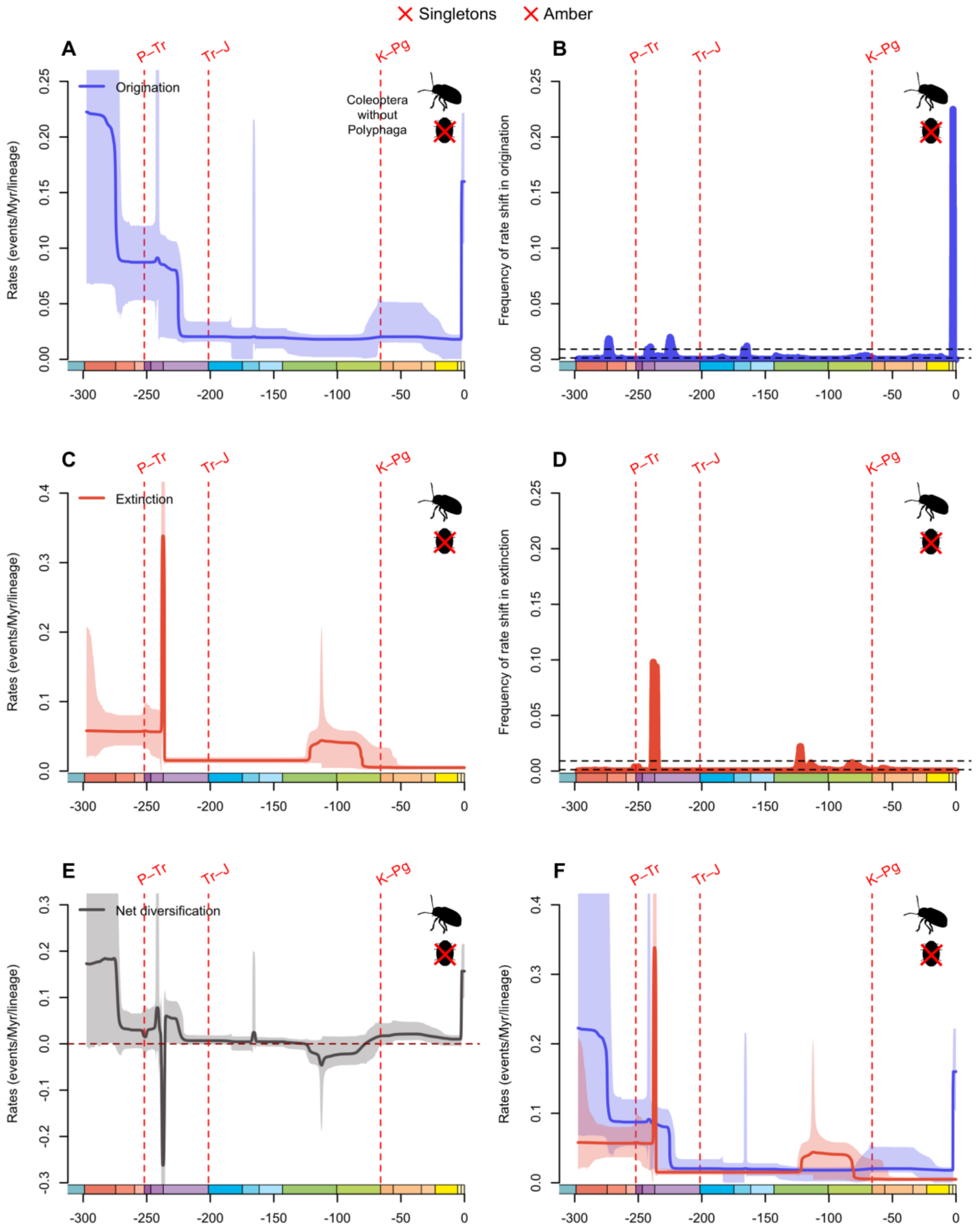


**fig. S70.** Diversification and diversity dynamics of Coleoptera genera without Polyphaga, excluding singletons and amber occurrences. Bayesian estimations of origination and extinction rates through time as inferred by PyRate using reversible jump Markov Chain Monte Carlo (RJMCMC). Marginal estimates of origination rates (A) and extinction rates (C) through time are shown as mean and 95% HPD (shaded areas). The frequency of a sampled rate shift is computed within small time bins for origination and extinction rates (B and D, respectively), with black-

dashed horizontal lines indicating log-Bayes factors of 2 (bottom) and 6 (top). Sampling frequencies higher than log-Bayes factors = 6 indicate strong statistical support for a rate shift. Panel (E) shows net diversification rates through time (computed as the posterior difference between origination and extinction rates through time). Panel (F) shows both marginal estimates of origination and extinction rates through time on the same plot. Solid lines indicate mean posterior rates and the shaded areas show 95% HPD. Red-dashed vertical lines indicate major crises: P–Tr, Permian–Triassic; Tr–J, Triassic–Jurassic; K–Pg, Cretaceous–Paleogene. Time is in millions of years. The color of each geological period in the chronostratigraphic scale follows that of the International Chronostratigraphic Chart (v2024/12). Insect silhouettes are from http://phylopic.org/.

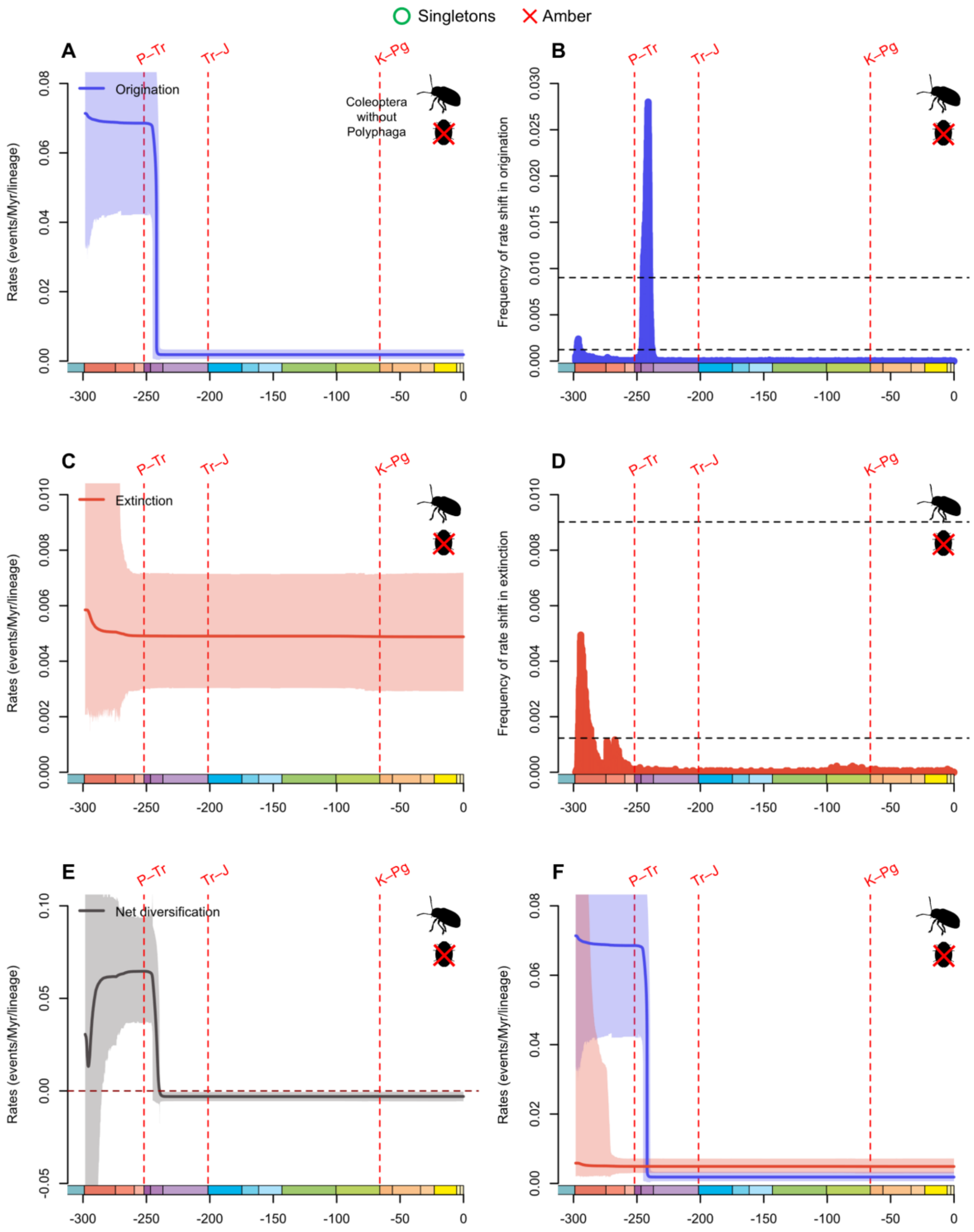


**fig. S71.** Diversification and diversity dynamics of Coleoptera families without Polyphaga, considering singletons and excluding amber occurrences. Bayesian estimations of origination and extinction rates through time as inferred by PyRate using reversible jump Markov Chain Monte Carlo (RJMCMC). Marginal estimates of origination rates (A) and extinction rates (C) through time are shown as mean and 95% HPD (shaded areas). The frequency of a sampled rate shift is computed within small time bins for origination and extinction rates (B and D, respectively), with

black-dashed horizontal lines indicating log-Bayes factors of 2 (bottom) and 6 (top). Sampling frequencies higher than log-Bayes factors = 6 indicate strong statistical support for a rate shift. Panel (E) shows net diversification rates through time (computed as the posterior difference between origination and extinction rates through time). Panel (F) shows both marginal estimates of origination and extinction rates through time on the same plot. Solid lines indicate mean posterior rates and the shaded areas show 95% HPD. Red-dashed vertical lines indicate major crises: P–Tr, Permian–Triassic; Tr–J, Triassic–Jurassic; K–Pg, Cretaceous–Paleogene. Time is in millions of years. The color of each geological period in the chronostratigraphic scale follows that of the International Chronostratigraphic Chart (v2024/12). Insect silhouettes are from http://phylopic.org/.

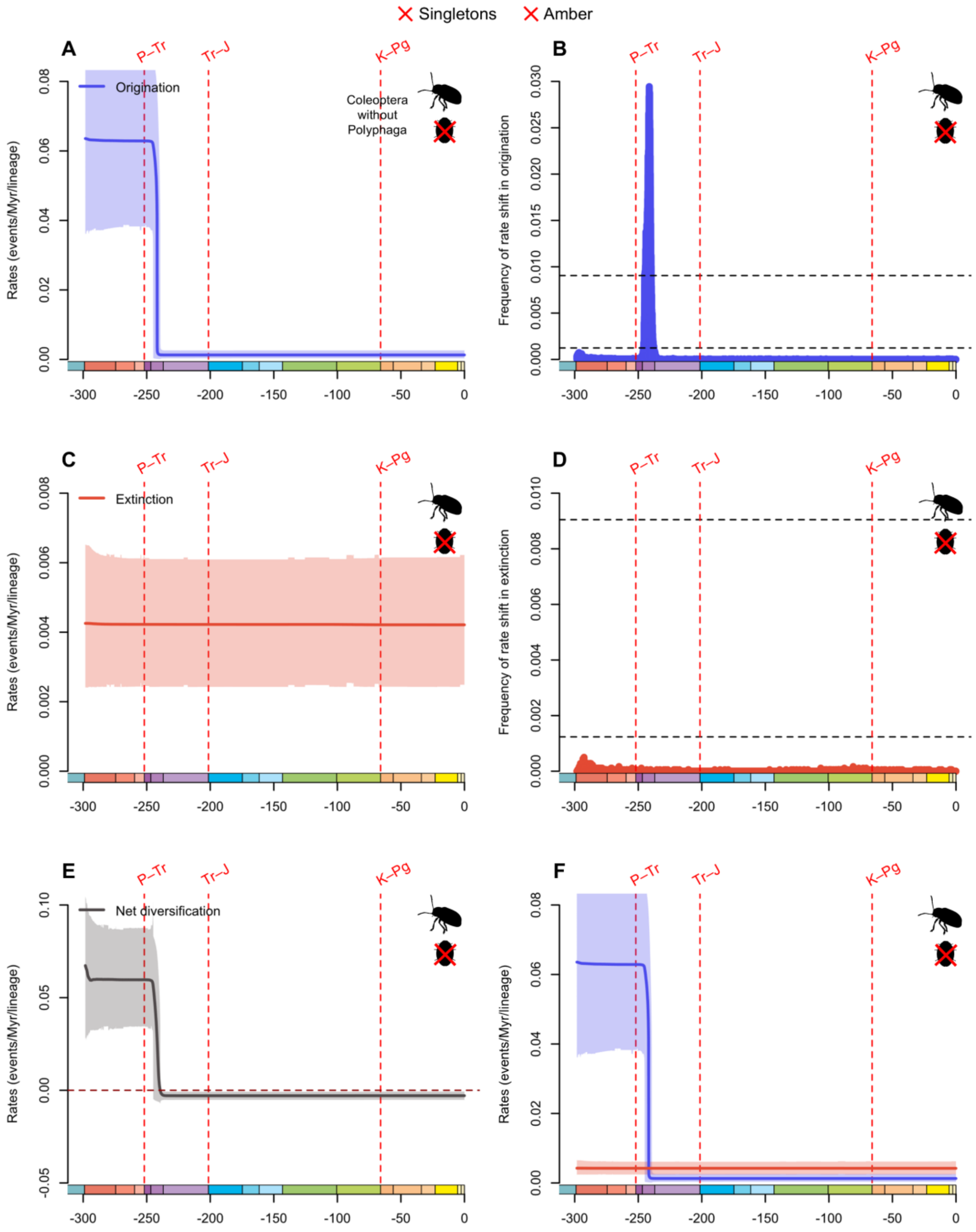


**fig. S72.** Diversification and diversity dynamics of Coleoptera families without Polyphaga, excluding singletons and amber occurrences. Bayesian estimations of origination and extinction rates through time as inferred by PyRate using reversible jump Markov Chain Monte Carlo (RJMCMC). Marginal estimates of origination rates (A) and extinction rates (C) through time are shown as mean and 95% HPD (shaded areas). The frequency of a sampled rate shift is computed within small time bins for origination and extinction rates (B and D, respectively), with black-

dashed horizontal lines indicating log-Bayes factors of 2 (bottom) and 6 (top). Sampling frequencies higher than log-Bayes factors = 6 indicate strong statistical support for a rate shift. Panel (E) shows net diversification rates through time (computed as the posterior difference between origination and extinction rates through time). Panel (F) shows both marginal estimates of origination and extinction rates through time on the same plot. Solid lines indicate mean posterior rates and the shaded areas show 95% HPD. Red-dashed vertical lines indicate major crises: P–Tr, Permian–Triassic; Tr–J, Triassic–Jurassic; K–Pg, Cretaceous–Paleogene. Time is in millions of years. The color of each geological period in the chronostratigraphic scale follows that of the International Chronostratigraphic Chart (v2024/12). Insect silhouettes are from http://phylopic.org/.

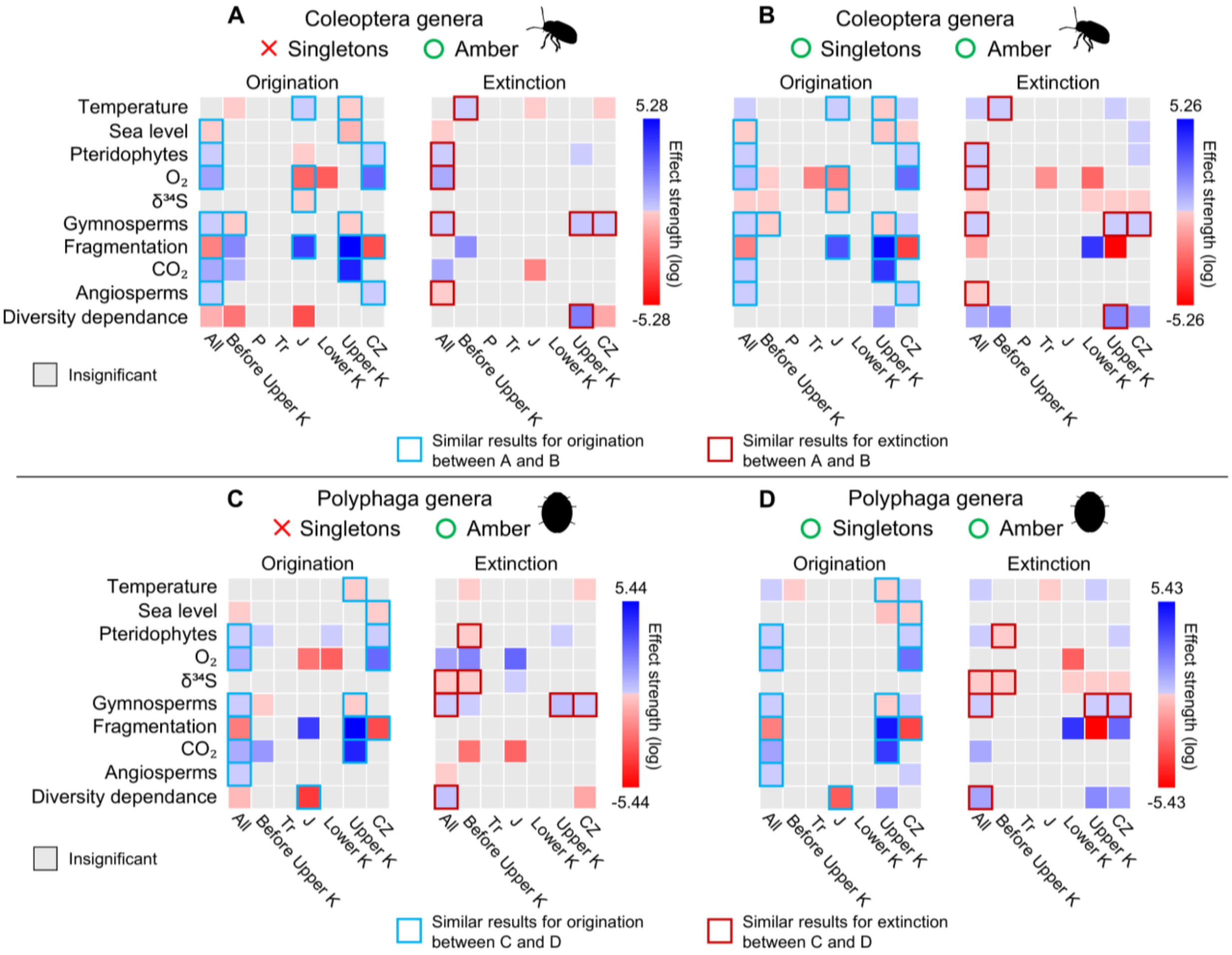


**fig. S73.** Bayesian estimates of correlation parameters for origination and extinction inferred under the MBD model for Coleoptera genera (A, B) and Polyphaga genera (C, D), with and without singletons, including amber occurrences, across multiple temporal windows. The strength effect corresponds to the median value of the correlation parameter of the origination/extinction rate. A variable was considered to have a significant effect when its shrinkage weight exceeded 0.5 and when the 95% HPD interval of the corresponding correlation parameter did not overlap with zero. When a significant correlation is positive, it is represented by a filled blue square; when a significant correlation is negative, it is represented by a filled red square; according to the sign of the median value of the correlation parameter. When a correlation is considered insignificant, it is represented by a light-grey filled square. If a correlation for a given clade is congruent between the MBD analyses with and without singletons, it is highlighted by a light-blue open square for origination, and by a dark-red open square for extinction. "All" corresponds to the time window encompassing the entire evolutionary history of the clade. "Before Upper K" corresponds to the time window spanning from the start of the clade's evolutionary history up to the beginning of the Upper Cretaceous (100.5 Ma). The other time intervals are defined as follows: P, Permian (298.9–251.902 Ma); Tr, Triassic (251.902–201.4 Ma); J, Jurassic (201.4–143.1 Ma); Lower K, Lower

Cretaceous (143.1–100.5 Ma); Upper K, Upper Cretaceous (100.5–66 Ma); CZ, Cenozoic (66 Ma to the present). Insect silhouettes are from http://phylopic.org/.

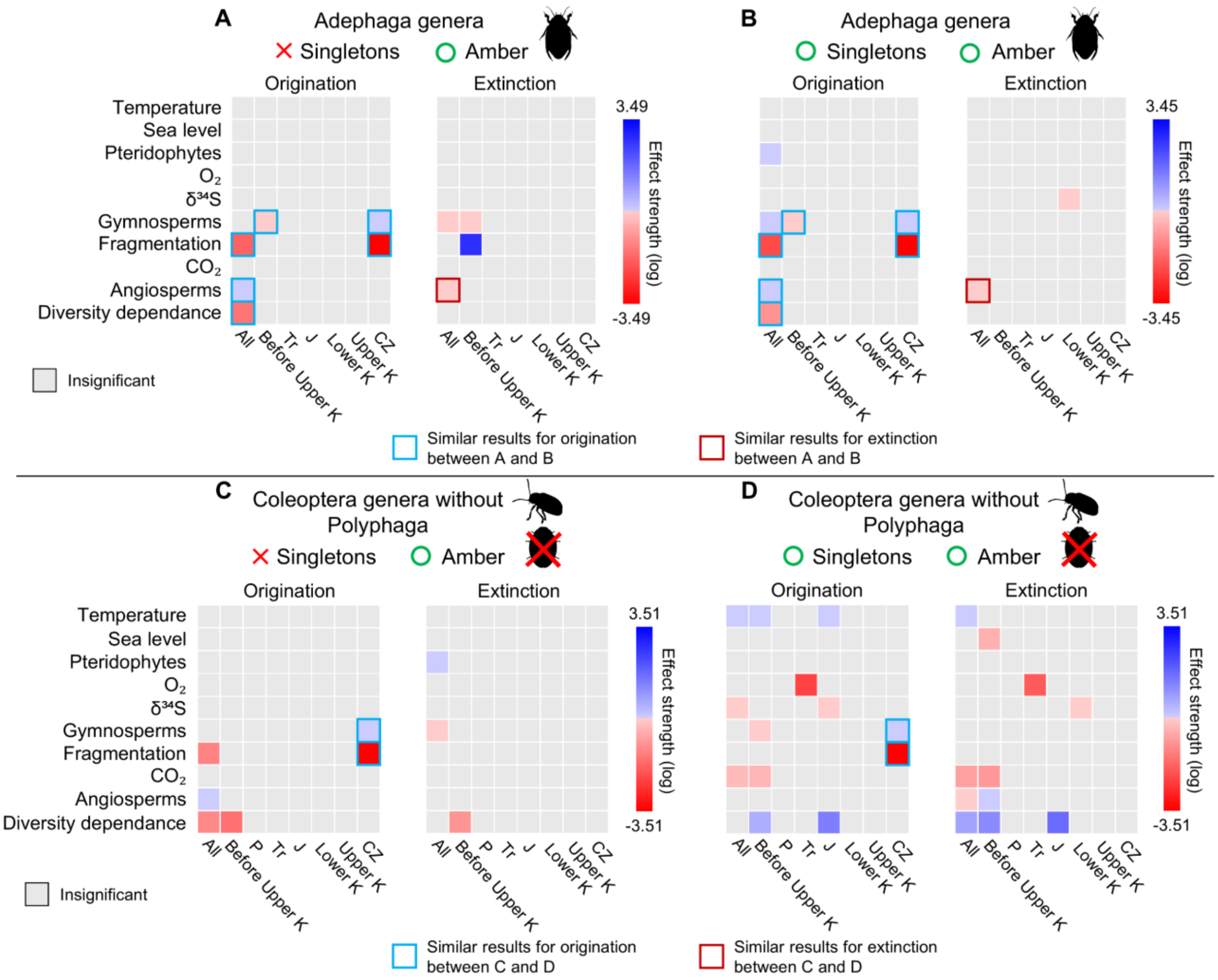


**fig. S74.** Bayesian estimates of correlation parameters for origination and extinction inferred under the MBD model for Adephaga genera (A, B) and Coleoptera genera excluding Polyphaga (C, D), with and without singletons, including amber occurrences, across multiple temporal windows. The strength effect corresponds to the median value of the correlation parameter of the origination/extinction rate. A variable was considered to have a significant effect when its shrinkage weight exceeded 0.5 and when the 95% HPD interval of the corresponding correlation parameter did not overlap with zero. When a significant correlation is positive, it is represented by a filled blue square; when a significant correlation is negative, it is represented by a filled red square; according to the sign of the median value of the correlation parameter. When a correlation is considered insignificant, it is represented by a light-grey filled square. If a correlation for a given clade is congruent between the MBD analyses with and without singletons, it is highlighted by a light-blue open square for origination, and by a dark-red open square for extinction. "All" corresponds to the time window encompassing the entire evolutionary history of the clade. "Before Upper K" corresponds to the time window spanning from the start of the clade's evolutionary history up to the beginning of the Upper Cretaceous (100.5 Ma). The other time intervals are defined as follows: P, Permian (298.9–251.902 Ma); Tr, Triassic (251.902–201.4 Ma); J, Jurassic (201.4–143.1 Ma); Lower K, Lower Cretaceous (143.1–100.5 Ma); Upper K, Upper Cretaceous

(100.5–66 Ma); CZ, Cenozoic (66 Ma to the present). Insect silhouettes are from http://phylopic.org/.

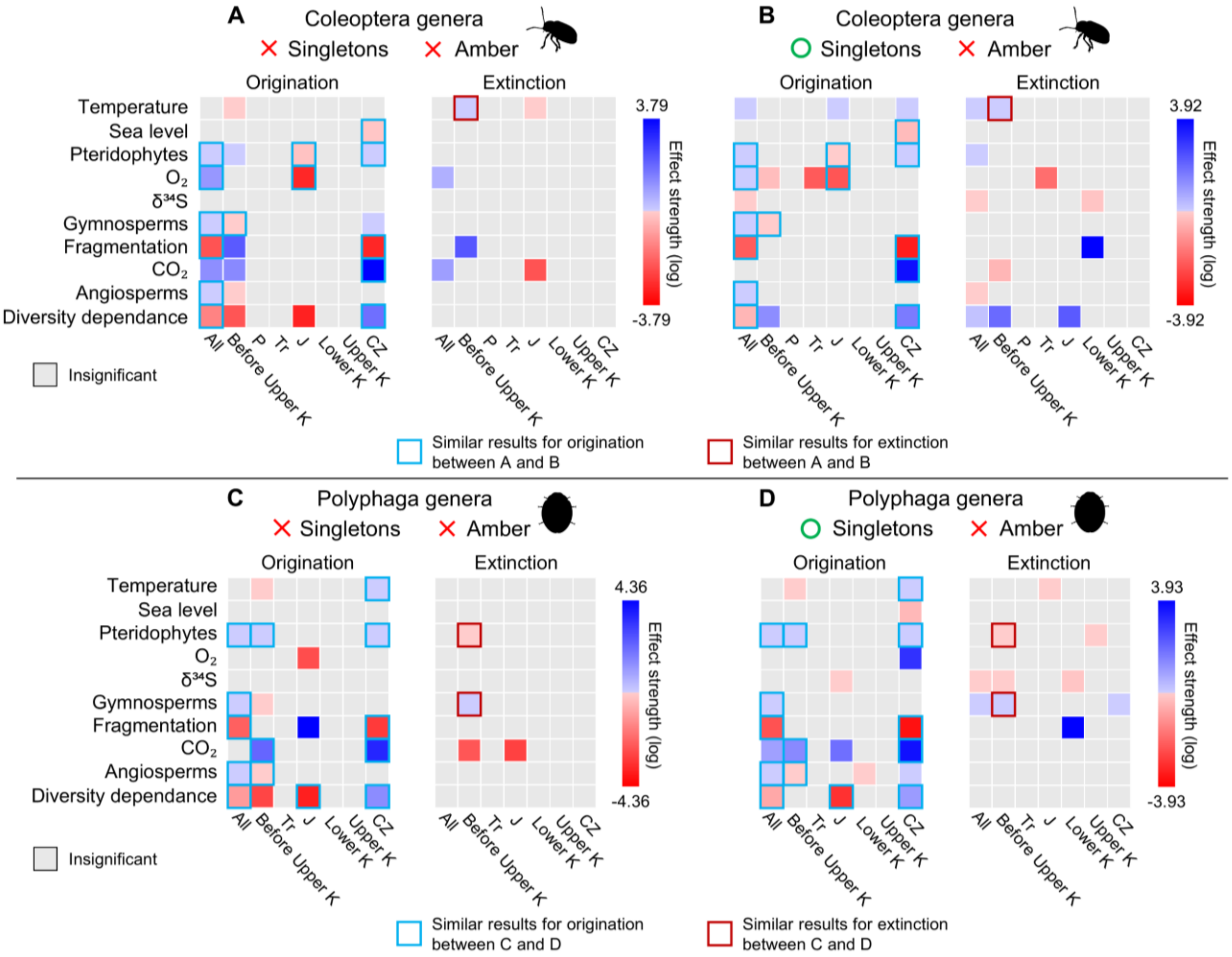


**fig. S75.** Bayesian estimates of correlation parameters for origination and extinction inferred under the MBD model for Coleoptera genera (A, B) and Polyphaga genera (C, D), with and without singletons, and excluding amber occurrences, across multiple temporal windows. The strength effect corresponds to the median value of the correlation parameter of the origination/extinction rate. A variable was considered to have a significant effect when its shrinkage weight exceeded 0.5 and when the 95% HPD interval of the corresponding correlation parameter did not overlap with zero. When a significant correlation is positive, it is represented by a filled blue square; when a significant correlation is negative, it is represented by a filled red square; according to the sign of the median value of the correlation parameter. When a correlation is considered insignificant, it is represented by a light-grey filled square. If a correlation for a given clade is congruent between the MBD analyses with and without singletons, it is highlighted by a light-blue open square for origination, and by a dark-red open square for extinction. "All" corresponds to the time window encompassing the entire evolutionary history of the clade. "Before Upper K" corresponds to the time window spanning from the start of the clade's evolutionary history up to the beginning of the Upper Cretaceous (100.5 Ma). The other time intervals are defined as follows: P, Permian (298.9–251.902 Ma); Tr, Triassic (251.902–201.4 Ma); J, Jurassic (201.4–143.1 Ma); Lower K, Lower Cretaceous (143.1–100.5 Ma); Upper K, Upper Cretaceous (100.5–66 Ma); CZ, Cenozoic (66 Ma to the present). Insect silhouettes are from http://phylopic.org/.

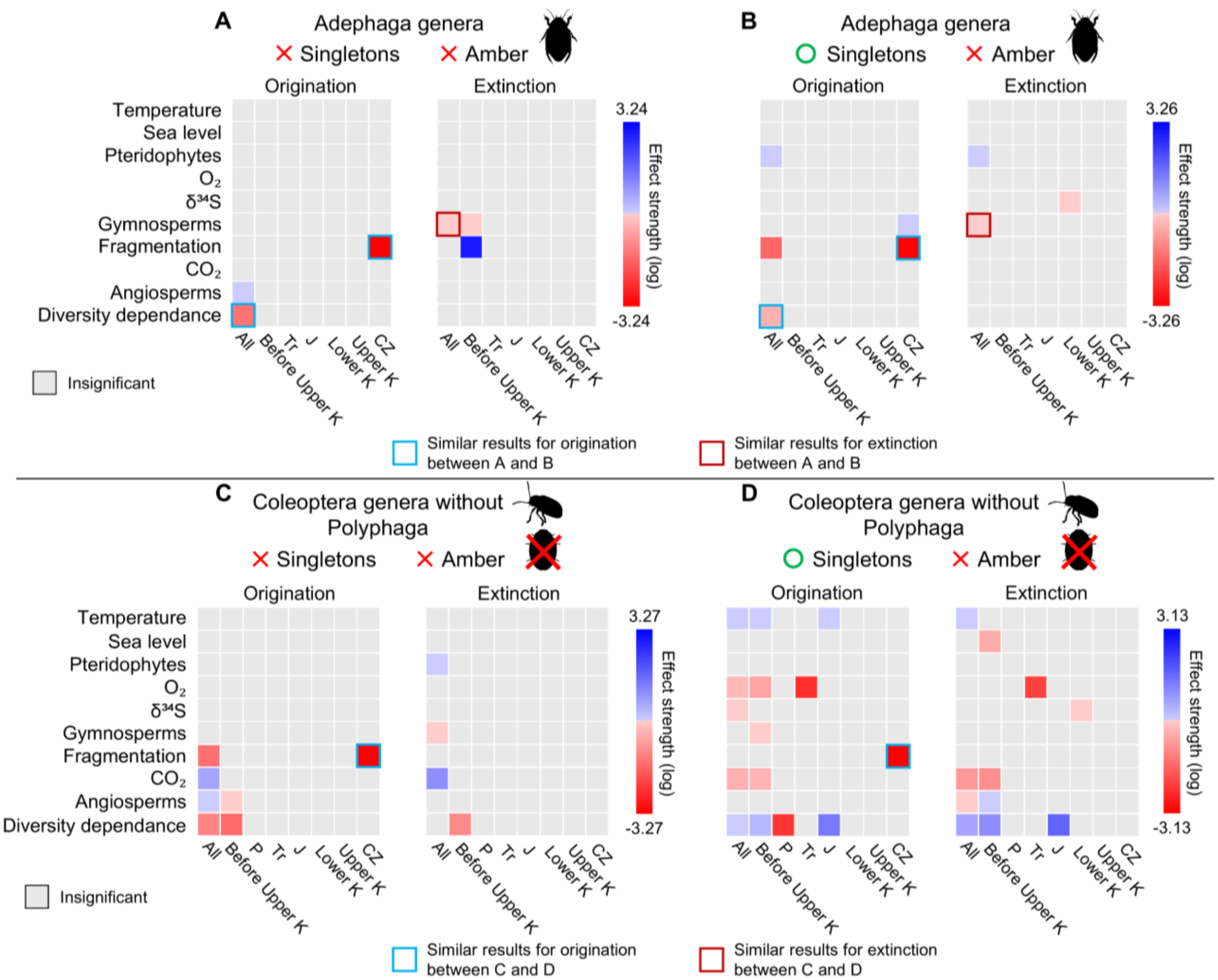


**fig. S76.** Bayesian estimates of correlation parameters for origination and extinction inferred under the MBD model for Adephaga genera (A, B) and Coleoptera genera excluding Polyphaga (C, D), with and without singletons, and excluding amber occurrences, across multiple temporal windows. The strength effect corresponds to the median value of the correlation parameter of the origination/extinction rate. A variable was considered to have a significant effect when its shrinkage weight exceeded 0.5 and when the 95% HPD interval of the corresponding correlation parameter did not overlap with zero. When a significant correlation is positive, it is represented by a filled blue square; when a significant correlation is negative, it is represented by a filled red square; according to the sign of the median value of the correlation parameter. When a correlation is considered insignificant, it is represented by a light-grey filled square. If a correlation for a given clade is congruent between the MBD analyses with and without singletons, it is highlighted by a light-blue open square for origination, and by a dark-red open square for extinction. "All" corresponds to the time window encompassing the entire evolutionary history of the clade. "Before Upper K" corresponds to the time window spanning from the start of the clade's evolutionary history up to the beginning of the Upper Cretaceous (100.5 Ma). The other time intervals are defined as follows: P, Permian (298.9–251.902 Ma); Tr, Triassic (251.902–201.4 Ma); J, Jurassic (201.4–143.1 Ma); Lower K, Lower Cretaceous (143.1–100.5 Ma); Upper K, Upper Cretaceous

(100.5–66 Ma); CZ, Cenozoic (66 Ma to the present). Insect silhouettes are from http://phylopic.org/.

**table S1.**

Posterior parameter estimates for the MBD model applied to Coleoptera genera, considering singletons and amber occurrences, across multiple temporal windows. The MBD model estimates the baseline origination and extinction rates ($\lambda 0$ and $\mu 0$), the correlation parameters ($G\lambda$ and $G\mu$) for each variable, and the shrinkage weights ($\omega$) of the correlation parameters. A variable was considered to have a significant effect (positive or negative depending on the sign of $G\lambda$ or $G\mu$) when its shrinkage weight exceeded 0.5 and when the 95% HPD interval of the corresponding correlation parameter did not overlap with zero (values highlighted in bold). The drivers are numbered as follows: (0) diversity of Coleoptera genera through time, (1) angiosperms diversity through time, (2) global variation of atmospheric $CO_2$ through time, (3) continental fragmentation through time, (4) gymnosperms diversity through time, (5) global variation in $\delta^{34}S$ through time (used here as an inverted proxy for global magmatic activity), (6) global variation of atmospheric $O_2$ through time, (7) Pteridophytes diversity through time, (8) Sea level fluctuations through time, and (9) variation of the global mean temperature through time. "All" corresponds to the time window encompassing the entire evolutionary history of Coleoptera genera, around 300 Ma to the present. "Before Upper Cretaceous" spans from around 300 to 100.5 Ma. The other time intervals are defined as follows: Permian (298.9 to 251.902 Ma), Triassic (251.902 to 201.4 Ma), Jurassic (201.4 to 143.1 Ma), Lower Cretaceous (143.1 to 100.5 Ma), Upper Cretaceous (100.5 to 66 Ma), and Cenozoic (66 Ma to the present).

| Parameters | | All | | Before Upper Cretaceous | |
|---|---|---|---|---|---|
| | | Median | 95% HPD | Median | 95% HPD |
| Baseline rates | λ0 | 2.48E-05 | [7.6576E-6, 5.9847E-5] | 0.7516 | [0.1801, 1.6154] |
| | μ0 | 2.27E-05 | [1.4482E-6, 8.0443E-5] | 0.0188 | [6.1864E-5, 0.0905] |
| Correlation parameters to origination | Gλ0_0 | -0.4326 | [-0.879, 0.0154] | 4.167 | [-0.5744, 8.2575] |
| | Gλ0_1 | **4.25E-03** | **[3.3278E-3, 5.0907E-3]** | -0.0013 | [-0.0368, 5.5503E-3] |
| | Gλ0_2 | **1.9771** | **[0.8851, 3.1246]** | -0.0137 | [-1.1509, 0.9926] |
| | Gλ0_3 | **-12.3365** | **[-14.2116, -10.4606]** | 0.7039 | [-2.0612, 4.6466] |
| | Gλ0_4 | **0.0787** | **[0.0693, 0.0884]** | **-0.0509** | **[-0.0754, -0.0303]** |
| | Gλ0_5 | **-0.0743** | **[-0.1094, -0.0394]** | **-0.0505** | **[-0.0946, -0.0051]** |
| | Gλ0_6 | **2.8613** | **[2.1303, 3.5451]** | **-1.8899** | **[-3.1955, -0.671]** |
| | Gλ0_7 | **0.0526** | **[0.0454, 0.0598]** | 0.0134 | [-0.0012, 0.0323] |
| | Gλ0_8 | **-0.7075** | **[-1.2089, -0.2076]** | 0.318 | [-0.4306, 1.5006] |
| | Gλ0_9 | **0.1174** | **[0.0999, 0.138]** | 0.0134 | [-0.0097, 0.0508] |
| Correlation parameters to extinction | Gμ0_0 | **4.1896** | **[3.3288, 5.1188]** | **8.5787** | **[1.4522, 14.5349]** |
| | Gμ0_1 | **-0.0028** | **[-0.0051, -0.0006]** | 0.0107 | [-0.0021, 0.0287] |
| | Gμ0_2 | 0.6539 | [-0.4511, 2.1855] | -1.3274 | [-2.7308, 0.0552] |
| | Gμ0_3 | **-4.8679** | **[-8.775, -2.0017]** | 2.749 | [-1.1278, 7.3889] |
| | Gμ0_4 | **0.0788** | **[0.0617, 0.099]** | 7.41E-03 | [-0.0099, 0.0484] |
| | Gμ0_5 | **-0.1509** | **[-0.206, -0.0925]** | -0.0059 | [-0.0544, 0.0272] |
| | Gμ0_6 | **1.9246** | **[0.9101, 2.887]** | -0.1112 | [-1.3907, 0.74] |
| | Gμ0_7 | **0.0276** | **[0.0148, 0.0411]** | -0.0024 | [-0.0217, 8.8947E-3] |
| | Gμ0_8 | -0.1448 | [-0.8039, 0.425] | -1.0935 | [-2.1755, 0.0514] |
| | Gμ0_9 | **0.1515** | **[0.1247, 0.193]** | **0.0638** | **[8.4503E-3, 0.1335]** |
| Shrinkage weights (origination) | ωλ0_0 | 0.5295 | [0.0305, 0.9986] | 0.9303 | [0.0681, 1] |
| | ωλ0_1 | **0.9348** | **[0.7267, 1]** | 0.8912 | [0.0206, 1] |
| | ωλ0_2 | **0.7702** | **[0.2447, 1]** | 0.1796 | [1.6311E-8, 0.9019] |
| | ωλ0_3 | **0.9182** | **[0.6763, 1]** | 0.264 | [2.0704E-8, 0.9284] |
| | ωλ0_4 | **0.9658** | **[0.8525, 1]** | **0.9111** | **[0.6483, 1]** |
| | ωλ0_5 | **0.8379** | **[0.3968, 0.9999]** | **0.6422** | **[0.1157, 0.9999]** |
| | ωλ0_6 | **0.9166** | **[0.6642, 1]** | **0.8055** | **[0.3266, 1]** |
| | ωλ0_7 | **0.9254** | **[0.7064, 1]** | 0.5542 | [1.1882E-6, 0.9631] |
| | ωλ0_8 | **0.6591** | **[0.1184, 0.9997]** | 0.351 | [1.6708E-7, 0.9436] |
| | ωλ0_9 | **0.959** | **[0.8317, 1]** | 0.4505 | [2.4916E-8, 0.9599] |
| Shrinkage weights (extinction) | ωμ0_0 | **0.9531** | **[0.7912, 1]** | **0.981** | **[0.7818, 1]** |
| | ωμ0_1 | **0.8901** | **[0.4151, 1]** | 0.9794 | [0.0474, 1] |
| | ωμ0_2 | 0.5126 | [6.413E-9, 0.9737] | 0.5292 | [0.0452, 0.9993] |
| | ωμ0_3 | **0.7833** | **[0.2643, 1]** | 0.4856 | [2.8207E-8, 0.9594] |
| | ωμ0_4 | **0.9653** | **[0.8492, 1]** | 0.488 | [3.2522E-8, 0.973] |
| | ωμ0_5 | **0.9291** | **[0.6944, 1]** | 0.2652 | [2.171E-7, 0.9336] |
| | ωμ0_6 | **0.8644** | **[0.4554, 1]** | 0.301 | [5.2685E-8, 0.9379] |
| | ωμ0_7 | **0.8458** | **[0.4186, 1]** | 0.2815 | [4.1632E-10, 0.9291] |
| | ωμ0_8 | 0.3695 | [5.2776E-9, 0.968] | 0.6511 | [0.0851, 1] |
| | ωμ0_9 | **0.9734** | **[0.8828, 1]** | **0.865** | **[0.3223, 1]** |

**table S1.** Continued.

| Parameters | | Permian | | Triassic | |
|---|---|---|---|---|---|
| | | Median | 95% HPD | Median | 95% HPD |
| Baseline rates | λ0 | 0.5734 | [0.0734, 1.466] | 0.5473 | [0.0353, 1.5457] |
| | μ0 | 0.553 | [0.0442, 1.483] | 0.454 | [0.0102, 1.342] |
| Correlation parameters to origination | Gλ0_0 | -86.5632 | [-159.8162, 5.6853] | -0.0291 | [-14.7555, 13.5051] |
| | Gλ0_1 | 7.89E-06 | [-0.0212, 0.0205] | 2.95E-06 | [-0.0284, 0.0295] |
| | Gλ0_2 | 0.3926 | [-3.1682, 6.0582] | 5.8311 | [-0.8196, 12.9913] |
| | Gλ0_3 | 0.6537 | [-21.9923, 62.5797] | 29.2095 | [-10.5062, 90.3841] |
| | Gλ0_4 | -0.0006 | [-0.136, 0.0921] | 0.0242 | [-0.0445, 0.194] |
| | Gλ0_5 | 0.0329 | [-0.061, 0.2318] | 0.0128 | [-0.1425, 0.2755] |
| | Gλ0_6 | 0.1096 | [-1.6881, 3.5186] | **-11.9291** | **[-16.3828, -7.1972]** |
| | Gλ0_7 | -0.0001 | [-0.1366, 0.1283] | -0.0962 | [-0.4648, 0.0512] |
| | Gλ0_8 | 0.2064 | [-1.515, 4.218] | -0.0129 | [-3.009, 3.0593] |
| | Gλ0_9 | -0.0139 | [-0.1242, 0.0324] | 2.12E-04 | [-0.1103, 0.1261] |
| Correlation parameters to extinction | Gμ0_0 | -0.0322 | [-19.3378, 15.471] | 3.7645 | [-6.5573, 68.3961] |
| | Gμ0_1 | 0 | [-0.285, 0.0699] | 6.68E-06 | [-0.0292, 0.0307] |
| | Gμ0_2 | 0.1292 | [-3.4832, 5.6133] | -0.9682 | [-15.2506, 4.0784] |
| | Gμ0_3 | 1.4121 | [-19.3685, 67.2625] | 10.3657 | [-11.8739, 77.7285] |
| | Gμ0_4 | 6.26E-04 | [-0.1138, 0.1393] | 1.02E-03 | [-0.0658, 0.0864] |
| | Gμ0_5 | 0.0678 | [-0.0536, 0.2829] | 0.0737 | [-0.0757, 0.2646] |
| | Gμ0_6 | 3.7966 | [-0.6987, 8.8458] | **-8.6934** | **[-16.6254, -2.7929]** |
| | Gμ0_7 | -0.0005 | [-0.1928, 0.1494] | -0.0327 | [-0.3974, 0.0835] |
| | Gμ0_8 | -0.0307 | [-3.3305, 2.9693] | -0.2042 | [-3.6514, 2.3808] |
| | Gμ0_9 | -0.1483 | [-0.273, 4.0001E-3] | 0.0463 | [-0.0394, 0.2918] |
| Shrinkage weights (origination) | ωλ0_0 | 0.9997 | [0.1438, 1] | 0.8327 | [0.0214, 1] |
| | ωλ0_1 | 0.6808 | [8.4066E-3, 1] | 0.8425 | [0.0234, 1] |
| | ωλ0_2 | 0.5181 | [2.3991E-9, 0.9763] | 0.9256 | [0.2249, 1] |
| | ωλ0_3 | 0.6802 | [8.3192E-3, 1] | 0.9769 | [0.1154, 1] |
| | ωλ0_4 | 0.6442 | [1.8461E-9, 0.9924] | 0.8586 | [0.0288, 1] |
| | ωλ0_5 | 0.6541 | [1.055E-8, 0.9846] | 0.6762 | [1.6502E-10, 0.9877] |
| | ωλ0_6 | 0.4545 | [5.5673E-11, 0.9763] | **0.9758** | **[0.8855, 1]** |
| | ωλ0_7 | 0.6332 | [3.9302E-14, 0.9931] | 0.96 | [0.0655, 1] |
| | ωλ0_8 | 0.516 | [7.5426E-10, 0.9813] | 0.6141 | [1.1747E-8, 0.9823] |
| | ωλ0_9 | 0.6261 | [1.1396E-8, 0.987] | 0.7134 | [0.0141, 1] |
| Shrinkage weights (extinction) | ωμ0_0 | 0.6812 | [8.3294E-3, 1] | 0.9716 | [0.0387, 1] |
| | ωμ0_1 | 0.7153 | [8.6443E-3, 1] | 0.8531 | [0.0291, 1] |
| | ωμ0_2 | 0.4715 | [6.3007E-8, 0.9747] | 0.7599 | [0.0205, 1] |
| | ωμ0_3 | 0.7347 | [8.7863E-3, 1] | 0.9304 | [0.0478, 1] |
| | ωμ0_4 | 0.6551 | [7.9426E-3, 1] | 0.6726 | [0.0149, 1] |
| | ωμ0_5 | 0.7865 | [0.0171, 1] | 0.7874 | [0.0407, 1] |
| | ωμ0_6 | 0.9211 | [0.0493, 1] | **0.9618** | **[0.746, 1]** |
| | ωμ0_7 | 0.6948 | [8.1466E-3, 0.9999] | 0.8924 | [0.031, 1] |
| | ωμ0_8 | 0.5111 | [7.288E-8, 0.9791] | 0.6657 | [9.6354E-7, 0.9858] |
| | ωμ0_9 | 0.9638 | [0.4739, 1] | 0.8509 | [0.0305, 1] |

**table S1.** Continued.

| Parameters | | Jurassic | | Lower Cretaceous | |
|---|---|---|---|---|---|
| | | Median | 95% HPD | Median | 95% HPD |
| Baseline rates | λ0 | 1.22E-03 | [1.1653E-7, 0.0283] | 2.10E-01 | [9.3895E-4, 0.9266] |
| | μ0 | 0.0218 | [3.7949E-6, 0.4974] | 0.6352 | [0.0721, 1.75] |
| Correlation parameters to origination | Gλ0_0 | -0.634 | [-15.7245, 6.9024] | 1.4998 | [-4.3641, 13.2978] |
| | Gλ0_1 | 8.26E-06 | [-0.0498, 0.0524] | -4.27E-02 | [-0.2013, 9.9776E-3] |
| | Gλ0_2 | 3.7875 | [-5.6758, 15.793] | -28.1234 | [-62.0186, 0.5389] |
| | Gλ0_3 | **36.1182** | **[7.7383, 60.4873]** | -0.0879 | [-15.7002, 19.5099] |
| | Gλ0_4 | -0.1101 | [-0.4749, 0.2379] | 0.1505 | [-0.0292, 0.4283] |
| | Gλ0_5 | **-0.4427** | **[-1.075, -0.0578]** | 0.4019 | [-0.0864, 1.2528] |
| | Gλ0_6 | **-12.7249** | **[-19.9422, -7.2416]** | 3.16E-03 | [-18.906, 22.2388] |
| | Gλ0_7 | -0.5964 | [-1.2798, 0.0581] | -0.0257 | [-0.2326, 0.0934] |
| | Gλ0_8 | 1.6002 | [-0.6002, 4.2554] | 0.8442 | [-0.5806, 3.1174] |
| | Gλ0_9 | **3.00E-01** | **[0.1765, 0.4257]** | 3.92E-01 | [-0.0708, 0.9731] |
| Correlation parameters to extinction | Gμ0_0 | 15.286 | [-0.4039, 25.582] | 0.2704 | [-6.6197, 9.1621] |
| | Gμ0_1 | 0.00E+00 | [-0.0477, 0.0454] | -2.00E-04 | [-0.0416, 0.0211] |
| | Gμ0_2 | -5.3623 | [-10.701, 0.2918] | -0.0352 | [-6.4039, 7.151] |
| | Gμ0_3 | 15.2559 | [-4.3122, 40.3194] | **64.161** | **[39.3876, 91.399]** |
| | Gμ0_4 | 1.04E-03 | [-0.1441, 0.1851] | -4.20E-03 | [-0.132, 0.1087] |
| | Gμ0_5 | 0.0954 | [-0.0908, 0.366] | **-1.7021** | **[-2.271, -1.1312]** |
| | Gμ0_6 | 0.9793 | [-2.1612, 5.5665] | **-22.249** | **[-36.5175, -8.7776]** |
| | Gμ0_7 | -0.229 | [-0.6735, 0.0665] | 0.0185 | [-0.0383, 0.0944] |
| | Gμ0_8 | 0.2046 | [-1.4616, 2.6715] | 0.8296 | [-0.7394, 2.9093] |
| | Gμ0_9 | 0.0844 | [-0.0282, 0.2356] | -0.009 | [-0.178, 0.1251] |
| Shrinkage weights (origination) | ωλ0_0 | 0.9307 | [0.0751, 1] | 0.9184 | [0.0398, 1] |
| | ωλ0_1 | 0.9408 | [0.0661, 1] | 0.9988 | [0.0999, 1] |
| | ωλ0_2 | 0.9355 | [0.0961, 1] | 0.9934 | [0.6197, 1] |
| | ωλ0_3 | **0.9848** | **[0.8624, 1]** | 0.6957 | [0.0166, 0.9999] |
| | ωλ0_4 | 0.9519 | [0.1351, 1] | 0.9459 | [0.0994, 1] |
| | ωλ0_5 | **0.9462** | **[0.566, 1]** | 0.9307 | [0.0556, 1] |
| | ωλ0_6 | **0.9761** | **[0.8787, 1]** | 0.867 | [0.0261, 1] |
| | ωλ0_7 | 0.9974 | [0.7073, 1] | 0.843 | [0.024, 1] |
| | ωλ0_8 | 0.8534 | [0.0594, 1] | 0.735 | [0.0284, 1] |
| | ωλ0_9 | **0.9847** | **[0.9222, 1]** | 0.9873 | [0.0922, 1] |
| Shrinkage weights (extinction) | ωμ0_0 | 0.9935 | [0.7681, 1] | 0.8861 | [0.0417, 1] |
| | ωμ0_1 | 0.9343 | [0.0622, 1] | 0.9058 | [0.0306, 1] |
| | ωμ0_2 | 0.9362 | [0.3218, 1] | 0.6946 | [0.0135, 1] |
| | ωμ0_3 | 0.9475 | [0.1244, 1] | **0.9886** | **[0.9411, 1]** |
| | ωμ0_4 | 0.8005 | [0.03, 1] | 0.6735 | [0.0124, 0.9999] |
| | ωμ0_5 | 0.7505 | [0.018, 1] | **0.9919** | **[0.96, 1]** |
| | ωμ0_6 | 0.7136 | [0.0192, 1] | **0.9877** | **[0.9191, 1]** |
| | ωμ0_7 | 0.9859 | [0.2482, 1] | 0.7218 | [0.0227, 1] |
| | ωμ0_8 | 0.6757 | [0.0133, 1] | 0.7204 | [0.0215, 1] |
| | ωμ0_9 | 0.9147 | [0.1058, 1] | 0.7587 | [0.0184, 1] |

**table S1.** Continued.

| Parameters | | Upper Cretaceous | | Cenozoic | |
|---|---|---|---|---|---|
| | | Median | 95% HPD | Median | 95% HPD |
| Baseline rates | λ0 | 3.84E-01 | [9.0021E-3, 1.3518] | 2.49E-04 | [1.0782E-6, 4.3733E-3] |
| | μ0 | 0.3883 | [6.0082E-3, 1.372] | 2.84E-04 | [1.8403E-8, 0.0343] |
| Correlation parameters to origination | Gλ0_0 | **6.1254** | **[3.2578, 8.8176]** | 0.3902 | [-1.4531, 2.0286] |
| | Gλ0_1 | -1.88E-02 | [-0.0857, 0.0441] | **7.88E-03** | **[1.4532E-3, 0.0133]** |
| | Gλ0_2 | **66.5281** | **[3.5209, 118.9271]** | 11.7602 | [-4.9033, 35.7644] |
| | Gλ0_3 | **149.5282** | **[44.0212, 218.2266]** | **-49.8682** | **[-69.8463, -24.8501]** |
| | Gλ0_4 | **-0.6242** | **[-1.0269, -0.1849]** | **0.0654** | **[0.0149, 0.1141]** |
| | Gλ0_5 | -0.0798 | [-0.3855, 0.3566] | -0.0128 | [-0.3469, 0.3751] |
| | Gλ0_6 | -1.42E+02 | [-404.2607, 77.0944] | **2.09E+01** | **[2.9272, 43.6929]** |
| | Gλ0_7 | 0.0349 | [-0.3035, 0.4187] | **0.2859** | **[0.1885, 0.3823]** |
| | Gλ0_8 | **-2.3645** | **[-4.0198, -0.7859]** | **-1.7354** | **[-2.7234, -0.7974]** |
| | Gλ0_9 | **-4.61E-01** | **[-0.7344, -0.2561]** | **3.18E-01** | **[0.1368, 0.4763]** |
| Correlation parameters to extinction | Gμ0_0 | **11.3803** | **[7.6858, 15.4623]** | **5.585** | **[3.4159, 7.7005]** |
| | Gμ0_1 | 6.88E-02 | [-0.0113, 0.1598] | 1.18E-03 | [-0.003, 9.298E-3] |
| | Gμ0_2 | 3.6061 | [-79.3607, 76.5] | -31.1563 | [-98.4623, 8.7035] |
| | Gμ0_3 | **-191.6877** | **[-297.8711, -95.9107]** | 31.2653 | [-1.106, 48.0266] |
| | Gμ0_4 | **1.20E+00** | **[0.7171, 1.761]** | **2.20E-01** | **[0.0893, 0.3351]** |
| | Gμ0_5 | **-1.0626** | **[-1.9327, -0.5441]** | **-1.2496** | **[-2.4609, -0.2245]** |
| | Gμ0_6 | -152.7812 | [-456.3672, 58.7132] | -29.2902 | [-49.5063, 1.1377] |
| | Gμ0_7 | 0.2357 | [-0.1088, 0.874] | **0.1978** | **[0.0924, 0.3543]** |
| | Gμ0_8 | 1.5118 | [-1.0729, 4.0609] | **1.7843** | **[0.3014, 4.2689]** |
| | Gμ0_9 | **0.6438** | **[0.306, 1.0755]** | 2.49E-03 | [-0.1656, 0.1639] |
| Shrinkage weights (origination) | ωλ0_0 | **0.9891** | **[0.9076, 1]** | 0.7304 | [0.0447, 1] |
| | ωλ0_1 | 0.9979 | [0.6723, 1] | **0.9616** | **[0.6632, 1]** |
| | ωλ0_2 | **0.9971** | **[0.9446, 1]** | 0.8766 | [0.08, 1] |
| | ωλ0_3 | **0.998** | **[0.9838, 1]** | **0.9801** | **[0.8908, 1]** |
| | ωλ0_4 | **0.9962** | **[0.9617, 1]** | **0.8514** | **[0.2948, 1]** |
| | ωλ0_5 | 0.8895 | [0.0403, 1] | 0.6519 | [0.0142, 1] |
| | ωλ0_6 | 0.9978 | [0.6205, 1] | **0.9409** | **[0.5227, 1]** |
| | ωλ0_7 | 0.9649 | [0.1612, 1] | **0.9506** | **[0.7716, 1]** |
| | ωλ0_8 | **0.9623** | **[0.585, 1]** | **0.831** | **[0.3406, 1]** |
| | ωλ0_9 | **0.9955** | **[0.9705, 1]** | **0.9676** | **[0.8149, 1]** |
| Shrinkage weights (extinction) | ωμ0_0 | **0.995** | **[0.9698, 1]** | **0.971** | **[0.8578, 1]** |
| | ωμ0_1 | 0.9995 | [0.8842, 1] | 0.8235 | [0.0326, 1] |
| | ωμ0_2 | 0.9928 | [0.5596, 1] | 0.9612 | [0.1732, 1] |
| | ωμ0_3 | **0.9988** | **[0.9932, 1]** | 0.9586 | [0.447, 1] |
| | ωμ0_4 | **0.9987** | **[0.9931, 1]** | **0.9672** | **[0.8098, 1]** |
| | ωμ0_5 | **0.9887** | **[0.9083, 1]** | **0.9701** | **[0.7369, 1]** |
| | ωμ0_6 | 0.9981 | [0.7586, 1] | 0.9544 | [0.337, 1] |
| | ωμ0_7 | 0.9825 | [0.3524, 1] | **0.9256** | **[0.62, 1]** |
| | ωμ0_8 | 0.9354 | [0.1183, 1] | **0.8488** | **[0.2646, 1]** |
| | ωμ0_9 | **0.9972** | **[0.9811, 1]** | 0.7551 | [0.0246, 1] |

**table S2.**

Posterior parameter estimates for the MBD model applied to Coleoptera genera, excluding singletons, but considering amber occurrences, across multiple temporal windows. The MBD model estimates the baseline origination and extinction rates ($\lambda 0$ and $\mu 0$), the correlation parameters ($G\lambda$ and $G\mu$) for each variable, and the shrinkage weights ($\omega$) of the correlation parameters. A variable was considered to have a significant effect (positive or negative depending on the sign of $G\lambda$ or $G\mu$) when its shrinkage weight exceeded 0.5 and when the 95% HPD interval of the corresponding correlation parameter did not overlap with zero (values highlighted in bold). The drivers are numbered as follows: (0) diversity of Coleoptera genera through time, (1) angiosperms diversity through time, (2) global variation of atmospheric $CO_2$ through time, (3) continental fragmentation through time, (4) gymnosperms diversity through time, (5) global variation in $\delta^{34}S$ through time (used here as an inverted proxy for global magmatic activity), (6) global variation of atmospheric $O_2$ through time, (7) Pteridophytes diversity through time, (8) Sea level fluctuations through time, and (9) variation of the global mean temperature through time. "All" corresponds to the time window encompassing the entire evolutionary history of Coleoptera genera, around 300 Ma to the present. "Before Upper Cretaceous" spans 300–100.5 Ma. The other time intervals are defined as follows: Permian (298.9–251.902 Ma), Triassic (251.902– 201.4 Ma), Jurassic (201.4–143.1 Ma), Lower Cretaceous (143.1–100.5 Ma), Upper Cretaceous (100.5–66 Ma), and Cenozoic (66 Ma to the present).

| Parameters | | All | | Before Upper Cretaceous | |
|---|---|---|---|---|---|
| | | Median | 95% HPD | Median | 95% HPD |
| Baseline rates | λ0 | 4.50E-05 | [5.1447E-6, 1.3034E-4] | 7.26E-01 | [0.1319, 1.6742] |
| | μ0 | 7.50E-05 | [4.474E-7, 4.6968E-4] | 2.26E-04 | [1.412E-8, 5.2485E-3] |
| Correlation parameters to origination | Gλ0_0 | **-4.1848** | **[-4.7244, -3.5979]** | **-16.6151** | **[-22.9865, -6.2609]** |
| | Gλ0_1 | **6.94E-03** | **[5.8219E-3, 8.2826E-3]** | -5.45E-02 | [-0.084, 1.4516E-3] |
| | Gλ0_2 | **5.0156** | **[3.3079, 6.683]** | **4.1692** | **[2.2234, 6.0291]** |
| | Gλ0_3 | **-12.4292** | **[-14.5721, -10.2194]** | **11.1085** | **[4.6761, 17.2229]** |
| | Gλ0_4 | **0.0589** | **[0.0494, 0.0686]** | **-0.0594** | **[-0.0915, -0.0322]** |
| | Gλ0_5 | 0.0209 | [-0.0106, 0.057] | -0.0244 | [-0.0716, 0.0135] |
| | Gλ0_6 | **5.6002** | **[4.5616, 6.5517]** | -0.1501 | [-1.8222, 1.026] |
| | Gλ0_7 | **0.0518** | **[0.0416, 0.061]** | 0.0159 | [-0.0078, 0.045] |
| | Gλ0_8 | **-1.449** | **[-2.0848, -0.722]** | 0.0652 | [-1.232, 2.0603] |
| | Gλ0_9 | 0.0279 | [-0.0029, 0.0583] | **-0.0474** | **[-0.0816, -0.0079]** |
| Correlation parameters to extinction | Gμ0_0 | 0.9138 | [-0.1781, 2.1349] | -0.7943 | [-5.9397, 1.7904] |
| | Gμ0_1 | **-0.0032** | **[-0.0057, -0.001]** | -0.0001 | [-0.0148, 0.0104] |
| | Gμ0_2 | **5.0493** | **[2.7426, 7.4798]** | 1.3563 | [-0.5762, 3.7523] |
| | Gμ0_3 | -0.6213 | [-4.3107, 2.889] | **9.5957** | **[3.3049, 16.4711]** |
| | Gμ0_4 | **0.0362** | **[0.0121, 0.066]** | 0.0299 | [-0.0108, 0.1023] |
| | Gμ0_5 | 2.03E-03 | [-0.0469, 0.0583] | 3.62E-02 | [-0.0109, 0.0937] |
| | Gμ0_6 | **4.7974** | **[3.2281, 6.3983]** | 1.4914 | [-0.5183, 4.2636] |
| | Gμ0_7 | **0.0341** | **[0.018, 0.0513]** | 8.06E-04 | [-0.0207, 0.0214] |
| | Gμ0_8 | **-1.7562** | **[-3.3304, -0.6871]** | -1.2156 | [-3.4606, 0.3768] |
| | Gμ0_9 | 0.0464 | [-0.0006, 0.0889] | **0.1016** | **[0.016, 0.1954]** |
| Shrinkage weights (origination) | ωλ0_0 | **0.9511** | **[0.7971, 1]** | **0.9947** | **[0.9668, 1]** |
| | ωλ0_1 | **0.9683** | **[0.862, 1]** | 0.9992 | [0.721, 1] |
| | ωλ0_2 | **0.9189** | **[0.6528, 1]** | **0.8786** | **[0.5199, 1]** |
| | ωλ0_3 | **0.9192** | **[0.6733, 1]** | **0.8894** | **[0.5241, 1]** |
| | ωλ0_4 | **0.9448** | **[0.7748, 1]** | **0.9407** | **[0.7353, 1]** |
| | ωλ0_5 | 0.5309 | [1.1738E-10, 0.9766] | 0.5357 | [9.5064E-8, 0.9698] |
| | ωλ0_6 | **0.9681** | **[0.8615, 1]** | 0.4695 | [9.3997E-8, 0.9678] |
| | ωλ0_7 | **0.9228** | **[0.6875, 1]** | 0.6912 | [0.0332, 0.9998] |
| | ωλ0_8 | **0.8196** | **[0.3492, 1]** | 0.486 | [1.1306E-7, 0.97] |
| | ωλ0_9 | 0.7655 | [0.0887, 1] | **0.8365** | **[0.3187, 1]** |
| Shrinkage weights (extinction) | ωμ0_0 | 0.7182 | [0.0602, 1] | 0.7591 | [0.0193, 1] |
| | ωμ0_1 | **0.8989** | **[0.4948, 1]** | 0.7797 | [0.0213, 1] |
| | ωμ0_2 | **0.9206** | **[0.6401, 1]** | 0.6409 | [0.0254, 0.9999] |
| | ωμ0_3 | 0.4213 | [9.0882E-9, 0.9717] | **0.8737** | **[0.4683, 1]** |
| | ωμ0_4 | **0.9031** | **[0.497, 1]** | 0.8678 | [0.0395, 1] |
| | ωμ0_5 | 0.4651 | [1.4513E-7, 0.9731] | 0.6142 | [0.0223, 0.9991] |
| | ωμ0_6 | **0.9601** | **[0.8188, 1]** | 0.7963 | [0.0365, 1] |
| | ωμ0_7 | **0.8723** | **[0.4915, 1]** | 0.4035 | [3.2629E-10, 0.9663] |
| | ωμ0_8 | **0.8536** | **[0.3976, 1]** | 0.7569 | [0.0341, 1] |
| | ωμ0_9 | 0.8551 | [0.2183, 0.9999] | **0.9407** | **[0.5622, 1]** |

**table S2.** Continued.

| Parameters | | Permian | | Triassic | |
|---|---|---|---|---|---|
| | | Median | 95% HPD | Median | 95% HPD |
| Baseline rates | λ0 | 2.23E-01 | [5.2516E-3, 0.8289] | 3.01E-01 | [1.6738E-3, 1.0562] |
| | μ0 | 2.07E-01 | [0.0132, 0.9401] | 2.78E-01 | [4.3774E-3, 1.0124] |
| Correlation parameters to origination | Gλ0_0 | -1.0419 | [-133.8386, 5.5547] | -0.3309 | [-44.6057, 5.6057] |
| | Gλ0_1 | 1.34E-06 | [-0.0107, 0.0103] | 0.00E+00 | [-0.0178, 0.0158] |
| | Gλ0_2 | -0.2612 | [-5.9119, 2.1089] | 1.3596 | [-2.3187, 12.1941] |
| | Gλ0_3 | 0.1967 | [-14.3361, 32.5336] | -10.4493 | [-80.6439, 8.5532] |
| | Gλ0_4 | -0.0003 | [-0.0964, 0.0641] | -0.0067 | [-0.0734, 0.0297] |
| | Gλ0_5 | 4.79E-03 | [-0.0626, 0.1673] | 5.54E-03 | [-0.0993, 0.1347] |
| | Gλ0_6 | -0.0219 | [-2.0101, 1.3054] | -2.4889 | [-7.7868, 0.9722] |
| | Gλ0_7 | -0.0003 | [-0.1098, 0.0635] | -0.0003 | [-0.1044, 0.0826] |
| | Gλ0_8 | 9.40E-03 | [-1.7393, 2.0958] | -4.67E-01 | [-4.3157, 1.4946] |
| | Gλ0_9 | -0.0007 | [-0.0523, 0.033] | -0.0047 | [-0.1356, 0.0459] |
| Correlation parameters to extinction | Gμ0_0 | -0.0056 | [-9.197, 5.9217] | -0.0107 | [-9.1621, 7.7274] |
| | Gμ0_1 | 4.30E-07 | [-0.0092, 8.4283E-3] | 0.00E+00 | [-0.0159, 0.0168] |
| | Gμ0_2 | -0.0181 | [-3.7366, 3.0861] | -0.2468 | [-7.1148, 3.6832] |
| | Gμ0_3 | 0.0566 | [-16.9184, 20.958] | -0.1216 | [-26.5366, 18.8179] |
| | Gμ0_4 | -0.0001 | [-0.069, 0.0567] | -0.0067 | [-0.0877, 0.0294] |
| | Gμ0_5 | 4.86E-03 | [-0.0703, 0.2092] | 8.00E-02 | [-0.0347, 0.2406] |
| | Gμ0_6 | -0.0829 | [-3.0244, 1.2883] | -4.2499 | [-9.9572, 0.9152] |
| | Gμ0_7 | -2.00E-04 | [-0.0926, 0.0674] | -6.00E-04 | [-0.1246, 0.0814] |
| | Gμ0_8 | -0.0198 | [-2.8456, 2.3221] | -0.7728 | [-6.5603, 1.7974] |
| | Gμ0_9 | -0.0195 | [-0.1041, 0.0165] | 5.40E-03 | [-0.0402, 0.1036] |
| Shrinkage weights (origination) | ωλ0_0 | 0.8619 | [1.9481E-3, 1] | 0.6878 | [7.2173E-3, 1] |
| | ωλ0_1 | 0.249 | [1.6976E-9, 0.9921] | 0.5365 | [5.1249E-3, 1] |
| | ωλ0_2 | 0.2718 | [1.2731E-10, 0.9568] | 0.6132 | [5.2882E-8, 0.9894] |
| | ωλ0_3 | 0.2513 | [1.3221E-9, 0.9809] | 0.8951 | [0.0139, 1] |
| | ωλ0_4 | 0.2509 | [1.5785E-9, 0.9798] | 0.4585 | [1.2407E-9, 0.9709] |
| | ωλ0_5 | 0.2299 | [3.4747E-10, 0.9521] | 0.3474 | [1.7962E-8, 0.9494] |
| | ωλ0_6 | 0.149 | [3.7935E-9, 0.9101] | 0.7225 | [0.017, 0.9995] |
| | ωλ0_7 | 0.2501 | [3.1321E-10, 0.9784] | 0.4419 | [4.7322E-8, 0.979] |
| | ωλ0_8 | 0.1736 | [7.6372E-10, 0.9179] | 0.5433 | [4.6179E-8, 0.9717] |
| | ωλ0_9 | 0.1553 | [4.1319E-8, 0.9088] | 0.3957 | [6.8089E-11, 0.9779] |
| Shrinkage weights (extinction) | ωμ0_0 | 0.2456 | [5.6661E-10, 0.9898] | 0.5044 | [1.7838E-8, 0.9916] |
| | ωμ0_1 | 0.2279 | [7.951E-12, 0.9878] | 0.5203 | [3.7562E-3, 1] |
| | ωμ0_2 | 0.158 | [7.7605E-9, 0.9218] | 0.405 | [1.2219E-10, 0.9717] |
| | ωμ0_3 | 0.227 | [1.1204E-8, 0.9769] | 0.4826 | [7.1202E-8, 0.9854] |
| | ωμ0_4 | 0.2278 | [3.3068E-9, 0.9734] | 0.4944 | [2.1863E-9, 0.9723] |
| | ωμ0_5 | 0.253 | [7.2182E-10, 0.9674] | 0.6916 | [4.9304E-8, 0.9818] |
| | ωμ0_6 | 0.2232 | [2.5807E-11, 0.9446] | 0.8338 | [0.0191, 1] |
| | ωμ0_7 | 0.2272 | [3.12E-9, 0.9777] | 0.4499 | [2.0704E-10, 0.9835] |
| | ωμ0_8 | 0.1834 | [1.5407E-9, 0.9432] | 0.6921 | [1.7472E-7, 0.9879] |
| | ωμ0_9 | 0.5051 | [9.4482E-10, 0.9746] | 0.367 | [2.7471E-7, 0.9687] |

**table S2.** Continued.

| Parameters | | Jurassic | | Lower Cretaceous | |
|---|---|---|---|---|---|
| | | Median | 95% HPD | Median | 95% HPD |
| Baseline rates | λ0 | 7.50E-01 | [0.0609, 2.0271] | 4.34E-01 | [0.0264, 1.3038] |
| | μ0 | 4.45E-01 | [3.2729E-3, 1.457] | 3.91E-01 | [8.7064E-3, 1.2124] |
| Correlation parameters to origination | Gλ0_0 | **-37.9458** | **[-48.5942, -28.5582]** | -0.0606 | [-13.7949, 12.0589] |
| | Gλ0_1 | 0.00E+00 | [-0.0597, 0.0536] | 3.49E-04 | [-0.0306, 0.0629] |
| | Gλ0_2 | 0.8311 | [-12.4818, 10.5103] | -17.1473 | [-52.3015, 5.6113] |
| | Gλ0_3 | **57.6394** | **[18.4766, 94.5182]** | 16.8529 | [-8.8735, 54.6688] |
| | Gλ0_4 | 0.0368 | [-0.3139, 0.8081] | -0.1268 | [-0.3938, 0.0843] |
| | Gλ0_5 | **-7.66E-01** | **[-1.5594, -0.1332]** | 1.19E-02 | [-0.4718, 0.4953] |
| | Gλ0_6 | **-22.3631** | **[-30.5017, -12.6251]** | **-27.8941** | **[-56.1601, -2.6505]** |
| | Gλ0_7 | **-1.1992** | **[-2.5923, -0.4367]** | 0.1539 | [-0.0062, 0.2996] |
| | Gλ0_8 | 2.39E-01 | [-7.1358, 4.6997] | 2.90E-01 | [-2.3466, 3.7192] |
| | Gλ0_9 | **0.1362** | **[0.0234, 0.2517]** | 0.3865 | [-0.1259, 1.0502] |
| Correlation parameters to extinction | Gμ0_0 | -0.9665 | [-16.833, 6.4564] | -8.2343 | [-36.4178, 4.7263] |
| | Gμ0_1 | 2.12E-05 | [-0.0481, 0.0546] | -2.70E-03 | [-0.2124, 0.0395] |
| | Gμ0_2 | **-12.047** | **[-18.801, -5.5176]** | 3.4498 | [-18.0498, 54.1039] |
| | Gμ0_3 | 0.5424 | [-25.1948, 30.3803] | 25.382 | [-12.007, 82.6306] |
| | Gμ0_4 | 0.1596 | [-0.079, 0.4026] | -0.0989 | [-0.6279, 0.5302] |
| | Gμ0_5 | 2.95E-01 | [-0.0551, 0.6477] | -1.02E+00 | [-1.8845, 0.0127] |
| | Gμ0_6 | 6.0109 | [-1.1225, 14.0159] | 0.3444 | [-37.4211, 29.2052] |
| | Gμ0_7 | 7.89E-02 | [-0.1827, 0.7246] | 1.04E-02 | [-0.1371, 0.2319] |
| | Gμ0_8 | 2.7134 | [-1.275, 7.5859] | -1.0993 | [-4.4665, 0.9145] |
| | Gμ0_9 | **-1.63E-01** | **[-0.2615, -0.0641]** | -8.24E-02 | [-0.7192, 0.1655] |
| Shrinkage weights (origination) | ωλ0_0 | **0.9991** | **[0.9956, 1]** | 0.9192 | [0.0594, 1] |
| | ωλ0_1 | 0.9691 | [0.1428, 1] | 0.9498 | [0.0757, 1] |
| | ωλ0_2 | 0.9189 | [0.0645, 1] | 0.9858 | [0.1666, 1] |
| | ωλ0_3 | **0.9938** | **[0.9526, 1]** | 0.9377 | [0.0868, 1] |
| | ωλ0_4 | 0.959 | [0.1175, 1] | 0.9468 | [0.1177, 1] |
| | ωλ0_5 | **0.9783** | **[0.8013, 1]** | 0.7893 | [0.0226, 1] |
| | ωλ0_6 | **0.9911** | **[0.9514, 1]** | **0.9916** | **[0.856, 1]** |
| | ωλ0_7 | **0.9994** | **[0.9948, 1]** | 0.97 | [0.4465, 1] |
| | ωλ0_8 | 0.8811 | [0.0499, 1] | 0.7708 | [0.0209, 1] |
| | ωλ0_9 | **0.9618** | **[0.6634, 1]** | 0.988 | [0.1253, 1] |
| Shrinkage weights (extinction) | ωμ0_0 | 0.9449 | [0.0922, 1] | 0.99 | [0.1526, 1] |
| | ωμ0_1 | 0.9686 | [0.1441, 1] | 0.9905 | [0.1568, 1] |
| | ωμ0_2 | **0.983** | **[0.893, 1]** | 0.9662 | [0.1122, 1] |
| | ωμ0_3 | 0.9112 | [0.0687, 1] | 0.9638 | [0.1458, 1] |
| | ωμ0_4 | 0.9624 | [0.2377, 1] | 0.9795 | [0.5569, 1] |
| | ωμ0_5 | 0.927 | [0.2319, 1] | 0.9818 | [0.7108, 1] |
| | ωμ0_6 | 0.9462 | [0.2929, 1] | 0.9684 | [0.1791, 1] |
| | ωμ0_7 | 0.9764 | [0.2004, 1] | 0.8882 | [0.0516, 1] |
| | ωμ0_8 | 0.9391 | [0.1636, 1] | 0.8309 | [0.0424, 1] |
| | ωμ0_9 | **0.9705** | **[0.7933, 1]** | 0.9498 | [0.0798, 1] |

**table S2.** Continued.

| Parameters | | Upper Cretaceous | | Cenozoic | |
|---|---|---|---|---|---|
| | | Median | 95% HPD | Median | 95% HPD |
| Baseline rates | λ0 | 3.83E-01 | [5.4694E-3, 1.2327] | 1.83E-04 | [5.6061E-7, 3.0865E-3] |
| | μ0 | 3.87E-01 | [0.0126, 1.234] | 1.84E-01 | [5.2219E-4, 0.7853] |
| Correlation parameters to origination | Gλ0_0 | -0.3578 | [-9.0066, 4.2206] | -1.08 | [-3.7303, 0.8366] |
| | Gλ0_1 | 6.00E-03 | [-0.0325, 0.1143] | **8.64E-03** | **[3.9814E-3, 0.0144]** |
| | Gλ0_2 | **106.2083** | **[51.2536, 200.7612]** | 29.9594 | [-1.7412, 65.2241] |
| | Gλ0_3 | **196.0201** | **[74.0702, 333.3077]** | **-37.6355** | **[-49.0691, -19.1376]** |
| | Gλ0_4 | **-0.8088** | **[-1.4578, -0.2483]** | 0.0144 | [-0.0307, 0.0639] |
| | Gλ0_5 | -2.50E-01 | [-0.7159, 0.1493] | 2.38E-01 | [-0.2904, 0.9738] |
| | Gλ0_6 | -246.3837 | [-627.3973, 15.7266] | **21.9103** | **[8.8458, 33.4077]** |
| | Gλ0_7 | -0.0446 | [-0.4809, 0.3273] | **0.2684** | **[0.183, 0.345]** |
| | Gλ0_8 | **-3.70E+00** | **[-7.3915, -0.3768]** | -1.96E+00 | [-2.9407, 5.3669E-3] |
| | Gλ0_9 | **-0.5349** | **[-0.8318, -0.2263]** | 0.1256 | [-0.0095, 0.2668] |
| Correlation parameters to extinction | Gμ0_0 | **13.2285** | **[4.9518, 22.14]** | **-5.0096** | **[-11.4696, -1.1818]** |
| | Gμ0_1 | 2.84E-02 | [-0.032, 0.1808] | 7.77E-04 | [-0.0038, 0.0149] |
| | Gμ0_2 | 0.3278 | [-63.6939, 141.9848] | -13.7988 | [-60.704, 7.2509] |
| | Gμ0_3 | -256.5281 | [-517.6593, 26.2255] | 1.7674 | [-8.5568, 18.1802] |
| | Gμ0_4 | **2.1444** | **[0.9276, 3.7635]** | **0.1092** | **[0.0199, 0.1806]** |
| | Gμ0_5 | -9.14E-01 | [-2.4458, 0.522] | -3.29E-01 | [-1.3783, 0.1476] |
| | Gμ0_6 | -258.4748 | [-1060.7066, 68.0065] | 3.626 | [-7.92, 32.1627] |
| | Gμ0_7 | **1.19E+00** | **[0.4533, 2.2871]** | 1.06E-01 | [-0.0171, 0.2936] |
| | Gμ0_8 | 0.5703 | [-1.778, 3.8533] | 0.9102 | [-1.6087, 4.3845] |
| | Gμ0_9 | 8.02E-02 | [-0.6198, 1.3789] | **-3.05E-01** | **[-0.4766, -0.1181]** |
| Shrinkage weights (origination) | ωλ0_0 | 0.9619 | [0.139, 1] | 0.7694 | [0.0303, 1] |
| | ωλ0_1 | 0.9956 | [0.5133, 1] | **0.9654** | **[0.7996, 1]** |
| | ωλ0_2 | **0.999** | **[0.9929, 1]** | 0.9556 | [0.4707, 1] |
| | ωλ0_3 | **0.9989** | **[0.9913, 1]** | **0.9668** | **[0.8248, 1]** |
| | ωλ0_4 | **0.9978** | **[0.9785, 1]** | 0.5436 | [1.0169E-8, 0.9769] |
| | ωλ0_5 | 0.9478 | [0.1345, 1] | 0.7941 | [0.0453, 1] |
| | ωλ0_6 | 0.9992 | [0.9824, 1] | **0.9286** | **[0.6341, 1]** |
| | ωλ0_7 | 0.9698 | [0.1969, 1] | **0.938** | **[0.7384, 1]** |
| | ωλ0_8 | **0.9795** | **[0.7204, 1]** | 0.8078 | [0.2044, 1] |
| | ωλ0_9 | **0.9964** | **[0.9728, 1]** | 0.874 | [0.1633, 1] |
| Shrinkage weights (extinction) | ωμ0_0 | **0.9963** | **[0.9707, 1]** | **0.9652** | **[0.7586, 1]** |
| | ωμ0_1 | 0.9986 | [0.6694, 1] | 0.7836 | [0.0198, 1] |
| | ωμ0_2 | 0.9903 | [0.3983, 1] | 0.8908 | [0.0417, 1] |
| | ωμ0_3 | 0.9992 | [0.9269, 1] | 0.6073 | [3.7264E-8, 0.9821] |
| | ωμ0_4 | **0.9996** | **[0.9972, 1]** | **0.8986** | **[0.4676, 1]** |
| | ωμ0_5 | 0.9877 | [0.7709, 1] | 0.8351 | [0.0375, 1] |
| | ωμ0_6 | 0.9994 | [0.9127, 1] | 0.7117 | [0.0124, 1] |
| | ωμ0_7 | **0.9981** | **[0.9846, 1]** | 0.8132 | [0.0921, 1] |
| | ωμ0_8 | 0.9201 | [0.0574, 1] | 0.7154 | [0.0202, 0.9999] |
| | ωμ0_9 | 0.9927 | [0.4014, 1] | **0.9628** | **[0.7975, 1]** |

**table S3.**

Posterior parameter estimates for the MBD model applied to Polyphaga genera, considering singletons and amber occurrences, across multiple temporal windows. The MBD model estimates the baseline origination and extinction rates ($\lambda 0$ and $\mu 0$), the correlation parameters ($G\lambda$ and $G\mu$) for each variable, and the shrinkage weights ($\omega$) of the correlation parameters. A variable was considered to have a significant effect (positive or negative depending on the sign of $G\lambda$ or $G\mu$) when its shrinkage weight exceeded 0.5 and when the 95% HPD interval of the corresponding correlation parameter did not overlap with zero (values highlighted in bold). The drivers are numbered as follows: (0) diversity of Polyphaga genera through time, (1) angiosperms diversity through time, (2) global variation of atmospheric $CO_2$ through time, (3) continental fragmentation through time, (4) gymnosperms diversity through time, (5) global variation in $\delta^{34}S$ through time (used here as an inverted proxy for global magmatic activity), (6) global variation of atmospheric $O_2$ through time, (7) Pteridophytes diversity through time, (8) Sea level fluctuations through time, and (9) variation of the global mean temperature through time. "All" corresponds to the time window encompassing the entire evolutionary history of Polyphaga genera, around 257 Ma to the present. "Before Upper Cretaceous" spans 257–100.5 Ma. The other time intervals are defined as follows: Triassic (251.902–201.4 Ma), Jurassic (201.4–143.1 Ma), Lower Cretaceous (143.1–100.5 Ma), Upper Cretaceous (100.5–66 Ma), and Cenozoic (66 Ma to the present).

| Parameters | | All | | Before Upper Cretaceous | |
|---|---|---|---|---|---|
| | | Median | 95% HPD | Median | 95% HPD |
| Baseline rates | λ0 | 1.55E-06 | [2.4487E-8, 6.0894E-6] | 5.04E-01 | [0.0357, 1.3682] |
| | μ0 | 4.21E-07 | [5.0582E-9, 4.4573E-6] | 2.10E-01 | [7.5291E-4, 0.8867] |
| Correlation parameters to origination | Gλ0_0 | -0.0536 | [-0.6095, 0.4624] | -4.5273 | [-15.2151, 1.3031] |
| | Gλ0_1 | **6.16E-03** | **[4.738E-3, 7.566E-3]** | -2.40E-03 | [-0.0328, 7.8985E-3] |
| | Gλ0_2 | **6.2193** | **[3.1413, 10.022]** | 2.5267 | [-0.4554, 6.5399] |
| | Gλ0_3 | **-14.0047** | **[-17.8435, -10.7373]** | -4.5668 | [-18.435, 3.2678] |
| | Gλ0_4 | **0.1041** | **[0.0913, 0.1159]** | -0.0158 | [-0.0843, 0.0263] |
| | Gλ0_5 | 0.018 | [-0.0243, 0.0771] | 0.0361 | [-0.0439, 0.1555] |
| | Gλ0_6 | **2.8687** | **[1.1172, 4.9964]** | 0.9389 | [-0.9244, 3.9587] |
| | Gλ0_7 | **0.0849** | **[0.0736, 0.096]** | 0.0384 | [-0.0023, 0.0722] |
| | Gλ0_8 | -0.6295 | [-1.6715, 0.0434] | 0.2769 | [-0.8931, 2.0259] |
| | Gλ0_9 | **0.1306** | **[0.1015, 0.1687]** | **-0.1387** | **[-0.2161, -0.0687]** |
| Correlation parameters to extinction | Gμ0_0 | **5.7127** | **[4.6166, 7.0982]** | 4.8067 | [-1.671, 15.53] |
| | Gμ0_1 | 3.83E-04 | [-0.0018, 3.2281E-3] | 6.18E-04 | [-0.0146, 0.0812] |
| | Gμ0_2 | **5.3322** | **[2.2382, 8.4226]** | -4.7204 | [-12.1466, 0.5981] |
| | Gμ0_3 | 0.9523 | [-3.6978, 6.9314] | 13.8962 | [-1.6158, 24.9016] |
| | Gμ0_4 | **0.1388** | **[0.1062, 0.1688]** | 0.1256 | [-0.0015, 0.3163] |
| | Gμ0_5 | **-0.3033** | **[-0.4068, -0.2113]** | **-0.4617** | **[-0.6907, -0.272]** |
| | Gμ0_6 | 0.1523 | [-1.8571, 3.4557] | 0.0834 | [-3.1084, 4.8101] |
| | Gμ0_7 | **0.0372** | **[0.0194, 0.0593]** | **-0.083** | **[-0.1513, -0.0232]** |
| | Gμ0_8 | 0.3387 | [-0.5472, 1.3843] | 0.1899 | [-2.1076, 2.2448] |
| | Gμ0_9 | **0.1893** | **[0.1137, 0.2748]** | -0.0781 | [-0.2539, 0.0229] |
| Shrinkage weights (origination) | ωλ0_0 | 0.33 | [3.3585E-8, 0.9684] | 0.9573 | [0.1135, 1] |
| | ωλ0_1 | **0.9621** | **[0.836, 1]** | 0.9391 | [0.0545, 1] |
| | ωλ0_2 | **0.9444** | **[0.7184, 1]** | 0.8252 | [0.0696, 1] |
| | ωλ0_3 | **0.9357** | **[0.7201, 1]** | 0.7957 | [0.0221, 0.9999] |
| | ωλ0_4 | **0.9686** | **[0.866, 1]** | 0.7834 | [0.0252, 1] |
| | ωλ0_5 | 0.5722 | [7.446E-8, 0.9807] | 0.7317 | [0.0205, 0.9999] |
| | ωλ0_6 | **0.8917** | **[0.4993, 1]** | 0.689 | [0.0163, 1] |
| | ωλ0_7 | **0.9519** | **[0.7983, 1]** | 0.8479 | [0.1375, 1] |
| | ωλ0_8 | 0.6344 | [0.0388, 1] | 0.54 | [4.0271E-7, 0.98] |
| | ωλ0_9 | **0.9598** | **[0.8239, 1]** | **0.9618** | **[0.8131, 1]** |
| Shrinkage weights (extinction) | ωμ0_0 | **0.9717** | **[0.8752, 1]** | 0.9635 | [0.1448, 1] |
| | ωμ0_1 | 0.676 | [0.0167, 1] | 0.9126 | [0.0392, 1] |
| | ωμ0_2 | **0.9275** | **[0.6401, 1]** | 0.9092 | [0.1146, 1] |
| | ωμ0_3 | 0.5807 | [2.2395E-9, 0.9798] | 0.9223 | [0.4272, 1] |
| | ωμ0_4 | **0.9803** | **[0.9108, 1]** | 0.976 | [0.743, 1] |
| | ωμ0_5 | **0.9751** | **[0.8863, 1]** | **0.9878** | **[0.9365, 1]** |
| | ωμ0_6 | 0.669 | [2.0722E-11, 0.9852] | 0.6791 | [0.014, 1] |
| | ωμ0_7 | **0.8679** | **[0.4567, 1]** | **0.9444** | **[0.6259, 1]** |
| | ωμ0_8 | 0.5291 | [2.1012E-9, 0.9781] | 0.6467 | [0.0171, 1] |
| | ωμ0_9 | **0.9779** | **[0.8913, 1]** | 0.9181 | [0.0834, 1] |

**table S3.** Continued.

| Parameters | | Triassic | | Jurassic | |
|---|---|---|---|---|---|
| | | Median | 95% HPD | Median | 95% HPD |
| Baseline rates | λ0 | 3.44E-01 | [0.0191, 1.1531] | 4.07E-01 | [3.1923E-3, 1.2277] |
| | μ0 | 3.37E-01 | [1.0306E-3, 1.17] | 5.18E-01 | [0.0226, 1.4237] |
| Correlation parameters to origination | Gλ0_0 | 2.39E-03 | [-8.4835, 9.6352] | **-3.25E+01** | **[-49.3023, -18.0023]** |
| | Gλ0_1 | 5.79E-06 | [-0.0119, 0.0108] | -1.00E-04 | [-0.8467, 0.0634] |
| | Gλ0_2 | -0.1479 | [-7.2326, 4.3434] | 5.8793 | [-1.5876, 12.1215] |
| | Gλ0_3 | 0.1073 | [-23.1222, 30.6236] | 0.0157 | [-42.0501, 55.04] |
| | Gλ0_4 | 1.86E-04 | [-0.0429, 0.0572] | 3.18E-02 | [-0.1227, 0.4096] |
| | Gλ0_5 | -0.0015 | [-0.1845, 0.1348] | -0.4342 | [-1.0829, 0.031] |
| | Gλ0_6 | -1.0603 | [-7.7184, 1.6973] | -7.1882 | [-16.7946, 1.4726] |
| | Gλ0_7 | -0.001 | [-0.1349, 0.088] | -0.1079 | [-1.0073, 0.1155] |
| | Gλ0_8 | 4.75E-03 | [-2.7428, 2.9807] | -3.80E-03 | [-2.2443, 2.2657] |
| | Gλ0_9 | -0.0154 | [-0.1254, 0.0306] | -0.0753 | [-0.1988, 0.0279] |
| Correlation parameters to extinction | Gμ0_0 | -0.0044 | [-8.8659, 9.0174] | 0.063 | [-6.5017, 8.8144] |
| | Gμ0_1 | 4.54E-08 | [-0.0137, 0.0119] | 0.00E+00 | [-0.0349, 0.0264] |
| | Gμ0_2 | -0.4756 | [-28.7706, 8.1695] | -2.7478 | [-7.4865, 1.1703] |
| | Gμ0_3 | -0.007 | [-44.4633, 43.4824] | 2.042 | [-11.1858, 25.0709] |
| | Gμ0_4 | -0.0011 | [-0.1499, 0.113] | 3.63E-03 | [-0.2057, 0.2191] |
| | Gμ0_5 | -0.0034 | [-0.8298, 0.3651] | -0.0193 | [-0.3193, 0.1847] |
| | Gμ0_6 | -0.7971 | [-31.9, 9.0041] | 1.7798 | [-2.6764, 9.0869] |
| | Gμ0_7 | -0.0006 | [-0.266, 0.169] | 0.3575 | [-0.0694, 0.9898] |
| | Gμ0_8 | -0.0717 | [-12.9965, 8.8045] | 0.1496 | [-2.4073, 3.5753] |
| | Gμ0_9 | -0.0289 | [-0.5139, 0.1459] | **-0.1384** | **[-0.2417, -0.042]** |
| Shrinkage weights (origination) | ωλ0_0 | 0.4156 | [2.3734E-9, 0.9936] | **0.9987** | **[0.9926, 1]** |
| | ωλ0_1 | 0.4161 | [1.3952E-8, 0.9925] | 0.8968 | [0.0329, 1] |
| | ωλ0_2 | 0.3684 | [5.5861E-10, 0.9721] | 0.9275 | [0.1925, 1] |
| | ωλ0_3 | 0.398 | [1.4203E-9, 0.9865] | 0.9411 | [0.0855, 1] |
| | ωλ0_4 | 0.2697 | [3.5147E-9, 0.9562] | 0.8616 | [0.0287, 1] |
| | ωλ0_5 | 0.3093 | [3.2336E-10, 0.9565] | 0.9312 | [0.3311, 1] |
| | ωλ0_6 | 0.515 | [2.1259E-8, 0.9777] | 0.9302 | [0.0953, 1] |
| | ωλ0_7 | 0.3797 | [3.8877E-10, 0.9826] | 0.971 | [0.0733, 1] |
| | ωλ0_8 | 0.3018 | [1.083E-8, 0.9566] | 0.578 | [3.9666E-7, 0.9803] |
| | ωλ0_9 | 0.4983 | [8.7394E-8, 0.9767] | 0.8739 | [0.0659, 1] |
| Shrinkage weights (extinction) | ωμ0_0 | 0.416 | [1.3797E-11, 0.9926] | 0.802 | [0.0272, 1] |
| | ωμ0_1 | 0.4321 | [1.2471E-8, 0.9944] | 0.8802 | [0.0356, 1] |
| | ωμ0_2 | 0.5982 | [3.8909E-3, 1] | 0.8311 | [0.067, 0.9999] |
| | ωμ0_3 | 0.4224 | [9.0695E-9, 0.9943] | 0.7853 | [0.0232, 0.9999] |
| | ωμ0_4 | 0.3958 | [1.0152E-10, 0.9874] | 0.8211 | [0.0293, 1] |
| | ωμ0_5 | 0.4323 | [2.9146E-9, 0.9949] | 0.5803 | [3.2195E-7, 0.9788] |
| | ωμ0_6 | 0.663 | [4.1781E-3, 1] | 0.7595 | [0.0242, 1] |
| | ωμ0_7 | 0.4162 | [4.6185E-7, 0.9947] | 0.9927 | [0.2139, 1] |
| | ωμ0_8 | 0.4358 | [1.1214E-9, 0.9956] | 0.6572 | [1.1786E-8, 0.9851] |
| | ωμ0_9 | 0.7624 | [5.9348E-3, 1] | **0.9429** | **[0.6625, 1]** |

**table S3.** Continued.

| Parameters | | **Lower Cretaceous** | | **Upper Cretaceous** | |
|---|---|---|---|---|---|
| | | Median | 95% HPD | Median | 95% HPD |
| Baseline rates | λ0 | 4.94E-01 | [0.0185, 1.5098] | 3.48E-01 | [6.2541E-3, 1.1191] |
| | μ0 | 5.24E-01 | [0.0339, 1.5329] | 3.43E-01 | [5.9635E-3, 1.1553] |
| Correlation parameters to origination | Gλ0_0 | -3.62E+00 | [-22.1658, 4.8063] | **6.14E+00** | **[3.0957, 9.3621]** |
| | Gλ0_1 | -1.14E-01 | [-0.2483, 9.9348E-3] | -8.30E-03 | [-0.0684, 0.0211] |
| | Gλ0_2 | -43.0353 | [-70.0592, 0.5047] | **68.8523** | **[24.5806, 98.3292]** |
| | Gλ0_3 | 3.7247 | [-11.3589, 29.2811] | **147.3873** | **[83.7458, 221.2349]** |
| | Gλ0_4 | 8.71E-02 | [-0.0914, 0.338] | **-7.86E-01** | **[-1.1929, -0.4199]** |
| | Gλ0_5 | 0.7228 | [-0.0339, 1.3312] | -0.2234 | [-0.5703, 0.1372] |
| | Gλ0_6 | -5.1987 | [-29.7604, 5.6431] | -145.5074 | [-306.9865, 41.8687] |
| | Gλ0_7 | 0.0631 | [-0.045, 0.277] | 0 | [-0.3899, 0.3297] |
| | Gλ0_8 | 2.25E-01 | [-1.9204, 2.5583] | **-2.83E+00** | **[-4.2807, -1.3632]** |
| | Gλ0_9 | 0.6645 | [-0.1499, 1.2681] | **-0.3759** | **[-0.6089, -0.1364]** |
| Correlation parameters to extinction | Gμ0_0 | 0.3042 | [-9.3493, 12.7255] | **11.5938** | **[7.9772, 15.3463]** |
| | Gμ0_1 | 0.00E+00 | [-0.0569, 0.1875] | 1.06E-01 | [-0.0018, 0.1995] |
| | Gμ0_2 | -1.461 | [-14.4794, 7.1453] | 25.5088 | [-27.5073, 103.8881] |
| | Gμ0_3 | **74.5851** | **[46.1414, 122.8364]** | **-227.1958** | **[-342.6143, -110.321]** |
| | Gμ0_4 | 5.65E-03 | [-0.1651, 0.2168] | **1.47E+00** | **[0.785, 2.0639]** |
| | Gμ0_5 | **-1.7032** | **[-2.9325, -0.9761]** | **-1.1986** | **[-2.1565, -0.2031]** |
| | Gμ0_6 | **-28.753** | **[-62.864, -10.1489]** | -185.8702 | [-468.458, 8.8167] |
| | Gμ0_7 | 0.049 | [-0.0483, 0.2877] | 0.1616 | [-0.2953, 0.7477] |
| | Gμ0_8 | 1.3685 | [-1.5719, 6.6283] | 1.2039 | [-2.1279, 4.9] |
| | Gμ0_9 | -0.0121 | [-0.2606, 0.215] | **0.5693** | **[0.082, 1.1266]** |
| Shrinkage weights (origination) | ωλ0_0 | 0.9687 | [0.113, 1] | **0.9897** | **[0.9081, 1]** |
| | ωλ0_1 | 0.9998 | [0.3888, 1] | 0.9946 | [0.566, 1] |
| | ωλ0_2 | 0.9967 | [0.8772, 1] | **0.9975** | **[0.9813, 1]** |
| | ωλ0_3 | 0.8263 | [0.0329, 1] | **0.9982** | **[0.99, 1]** |
| | ωλ0_4 | 0.9245 | [0.0905, 1] | **0.9976** | **[0.9855, 1]** |
| | ωλ0_5 | 0.968 | [0.3228, 1] | 0.9353 | [0.134, 1] |
| | ωλ0_6 | 0.9444 | [0.0737, 1] | 0.998 | [0.8985, 1] |
| | ωλ0_7 | 0.9307 | [0.0983, 1] | 0.9652 | [0.1823, 1] |
| | ωλ0_8 | 0.7406 | [0.0192, 1] | **0.9731** | **[0.699, 1]** |
| | ωλ0_9 | 0.9953 | [0.6339, 1] | **0.9941** | **[0.9483, 1]** |
| Shrinkage weights (extinction) | ωμ0_0 | 0.938 | [0.0845, 1] | **0.9953** | **[0.9711, 1]** |
| | ωμ0_1 | 0.9684 | [0.1093, 1] | 0.9998 | [0.9962, 1] |
| | ωμ0_2 | 0.8744 | [0.0404, 1] | 0.9933 | [0.4788, 1] |
| | ωμ0_3 | **0.9923** | **[0.9573, 1]** | **0.9991** | **[0.9944, 1]** |
| | ωμ0_4 | 0.8366 | [0.0322, 1] | **0.9991** | **[0.9949, 1]** |
| | ωμ0_5 | **0.9927** | **[0.9587, 1]** | **0.9904** | **[0.8972, 1]** |
| | ωμ0_6 | **0.9932** | **[0.9451, 1]** | 0.9987 | [0.9462, 1] |
| | ωμ0_7 | 0.9069 | [0.0637, 1] | 0.9809 | [0.3414, 1] |
| | ωμ0_8 | 0.8766 | [0.0626, 1] | 0.947 | [0.1197, 1] |
| | ωμ0_9 | 0.882 | [0.0505, 1] | **0.9967** | **[0.97, 1]** |

**table S3.** Continued.

| Parameters | | Cenozoic | |
|---|---|---|---|
| | | Median | 95% HPD |
| Baseline rates | λ0 | 1.24E-03 | [8.72E-6, 0.0133] |
| | μ0 | 1.91E-04 | [1.3525E-8, 0.0845] |
| Correlation parameters to origination | Gλ0_0 | 1.60E-05 | [-1.2549, 1.3619] |
| | Gλ0_1 | **5.64E-03** | **[8.2844E-4, 0.0109]** |
| | Gλ0_2 | 3.2776 | [-13.7253, 27.2287] |
| | Gλ0_3 | **-54.2668** | **[-66.7087, -34.8029]** |
| | Gλ0_4 | **7.84E-02** | **[0.0332, 0.1184]** |
| | Gλ0_5 | -0.0745 | [-0.5274, 0.3425] |
| | Gλ0_6 | **20.5679** | **[7.1407, 33.5522]** |
| | Gλ0_7 | **0.2903** | **[0.2034, 0.365]** |
| | Gλ0_8 | **-1.40E+00** | **[-2.5769, -0.3131]** |
| | Gλ0_9 | **0.3385** | **[0.2046, 0.441]** |
| Correlation parameters to extinction | Gμ0_0 | **5.2594** | **[2.6563, 7.4295]** |
| | Gμ0_1 | 1.83E-03 | [-0.003, 9.3369E-3] |
| | Gμ0_2 | -15.9399 | [-97.155, 23.0536] |
| | Gμ0_3 | **22.8815** | **[6.6477, 47.3726]** |
| | Gμ0_4 | **2.16E-01** | **[0.1115, 0.3605]** |
| | Gμ0_5 | **-0.982** | **[-2.3366, -0.2483]** |
| | Gμ0_6 | -26.9064 | [-46.4, 1.053] |
| | Gμ0_7 | **0.1901** | **[0.0754, 0.3297]** |
| | Gμ0_8 | 0.9982 | [-0.4637, 2.7242] |
| | Gμ0_9 | -0.0097 | [-0.1963, 0.28] |
| Shrinkage weights (origination) | ωλ0_0 | 0.4915 | [3.181E-8, 0.9792] |
| | ωλ0_1 | **0.9374** | **[0.5366, 1]** |
| | ωλ0_2 | 0.7533 | [0.0268, 1] |
| | ωλ0_3 | **0.9825** | **[0.9169, 1]** |
| | ωλ0_4 | **0.8721** | **[0.4491, 1]** |
| | ωλ0_5 | 0.6657 | [0.0157, 0.9999] |
| | ωλ0_6 | **0.9308** | **[0.6097, 1]** |
| | ωλ0_7 | **0.9489** | **[0.7679, 1]** |
| | ωλ0_8 | **0.7855** | **[0.2093, 1]** |
| | ωλ0_9 | **0.9706** | **[0.8607, 1]** |
| Shrinkage weights (extinction) | ωμ0_0 | **0.9657** | **[0.8254, 1]** |
| | ωμ0_1 | 0.8429 | [0.0398, 1] |
| | ωμ0_2 | 0.9402 | [0.0681, 1] |
| | ωμ0_3 | **0.9423** | **[0.6058, 1]** |
| | ωμ0_4 | **0.9675** | **[0.8134, 1]** |
| | ωμ0_5 | **0.9638** | **[0.6999, 1]** |
| | ωμ0_6 | 0.9464 | [0.3768, 1] |
| | ωμ0_7 | **0.9183** | **[0.568, 1]** |
| | ωμ0_8 | 0.7127 | [0.0272, 0.9996] |
| | ωμ0_9 | 0.8401 | [0.0738, 1] |

**table S4.**

Posterior parameter estimates for the MBD model applied to Polyphaga genera, excluding singletons, but considering amber occurrences, across multiple temporal windows. The MBD model estimates the baseline origination and extinction rates (λ0 and μ0), the correlation parameters (Gλ and Gμ) for each variable, and the shrinkage weights (ω) of the correlation parameters. A variable was considered to have a significant effect (positive or negative depending on the sign of Gλ or Gμ) when its shrinkage weight exceeded 0.5 and when the 95% HPD interval of the corresponding correlation parameter did not overlap with zero (values highlighted in bold). The drivers are numbered as follows: (0) diversity of Polyphaga genera through time, (1) angiosperms diversity through time, (2) global variation of atmospheric $CO_2$ through time, (3) continental fragmentation through time, (4) gymnosperms diversity through time, (5) global variation in $\delta^{34}S$ through time (used here as an inverted proxy for global magmatic activity), (6) global variation of atmospheric $O_2$ through time, (7) Pteridophytes diversity through time, (8) Sea level fluctuations through time, and (9) variation of the global mean temperature through time. "All" corresponds to the time window encompassing the entire evolutionary history of Polyphaga genera, around 257 Ma to the present. "Before Upper Cretaceous" spans 257–100.5 Ma. The other time intervals are defined as follows: Triassic (251.902–201.4 Ma), Jurassic (201.4–143.1 Ma), Lower Cretaceous (143.1–100.5 Ma), Upper Cretaceous (100.5–66 Ma), and Cenozoic (66 Ma to the present).

| Parameters | | All | | Before Upper Cretaceous | |
|---|---|---|---|---|---|
| | | Median | 95% HPD | Median | 95% HPD |
| Baseline rates | λ0 | 2.76E-05 | [1.8313E-6, 1.1236E-4] | 7.15E-01 | [0.1031, 1.8303] |
| | μ0 | 7.69E-05 | [4.4151E-7, 7.4382E-4] | 1.34E-01 | [1.0342E-5, 0.7128] |
| Correlation parameters to origination | Gλ0_0 | **-3.3788** | **[-4.2541, -2.4828]** | -26.0746 | [-40.287, 0.4252] |
| | Gλ0_1 | **6.69E-03** | **[5.4588E-3, 7.9418E-3]** | -3.30E-02 | [-0.0931, 0.0375] |
| | Gλ0_2 | **4.9514** | **[2.6043, 7.4319]** | **8.1674** | **[2.5387, 14.5149]** |
| | Gλ0_3 | **-15.3008** | **[-18.4094, -12.1574]** | 0.9026 | [-7.0427, 12.0988] |
| | Gλ0_4 | **0.0808** | **[0.0647, 0.0945]** | **-0.1186** | **[-0.1932, -0.0428]** |
| | Gλ0_5 | 0.0742 | [-0.0007, 0.1376] | -0.0051 | [-0.1221, 0.1077] |
| | Gλ0_6 | **4.0095** | **[2.0775, 6.0275]** | -0.2899 | [-5.8347, 3.9328] |
| | Gλ0_7 | **0.0733** | **[0.0608, 0.0845]** | **0.0687** | **[9.1964E-3, 0.1229]** |
| | Gλ0_8 | **-0.9806** | **[-1.6774, -0.1363]** | 1.5815 | [-0.671, 5.355] |
| | Gλ0_9 | 0.0425 | [-0.0039, 0.084] | -0.1249 | [-0.2315, 0.0173] |
| Correlation parameters to extinction | Gμ0_0 | **2.6582** | **[1.3569, 4.0338]** | -4.086 | [-34.6338, 6.0271] |
| | Gμ0_1 | **-0.0048** | **[-0.008, -0.0015]** | 0.0263 | [-0.012, 0.1707] |
| | Gμ0_2 | 3.8964 | [-0.0505, 7.3563] | **-18.8791** | **[-28.9493, -8.6574]** |
| | Gμ0_3 | -0.5067 | [-5.0867, 3.3747] | 15.4391 | [-34.4039, 38.1366] |
| | Gμ0_4 | **0.0842** | **[0.0522, 0.1223]** | **0.4391** | **[0.1793, 0.8738]** |
| | Gμ0_5 | **-0.1743** | **[-0.3076, -0.0565]** | **-0.4636** | **[-0.77, -0.1702]** |
| | Gμ0_6 | **6.0421** | **[3.3802, 8.7101]** | **12.6827** | **[3.349, 23.1936]** |
| | Gμ0_7 | 4.17E-03 | [-0.0129, 0.0279] | **-2.62E-01** | **[-0.3905, -0.135]** |
| | Gμ0_8 | -0.0837 | [-1.5215, 1.0126] | 1.6126 | [-1.0325, 4.8203] |
| | Gμ0_9 | 0.0239 | [-0.0163, 0.0882] | **-0.474** | **[-1.0629, -0.0971]** |
| Shrinkage weights (origination) | ωλ0_0 | **0.9349** | **[0.7262, 1]** | 0.9979 | [0.9355, 1] |
| | ωλ0_1 | **0.9665** | **[0.851, 1]** | 0.9985 | [0.778, 1] |
| | ωλ0_2 | **0.9194** | **[0.6326, 1]** | **0.9774** | **[0.8053, 1]** |
| | ωλ0_3 | **0.9398** | **[0.7433, 0.9999]** | 0.8368 | [0.0343, 1] |
| | ωλ0_4 | **0.9526** | **[0.8026, 1]** | **0.9834** | **[0.8733, 1]** |
| | ωλ0_5 | 0.8287 | [0.2037, 1] | 0.8374 | [0.0255, 1] |
| | ωλ0_6 | **0.922** | **[0.6479, 1]** | 0.8999 | [0.0591, 1] |
| | ωλ0_7 | **0.9401** | **[0.7407, 1]** | **0.9633** | **[0.631, 1]** |
| | ωλ0_8 | **0.7185** | **[0.1415, 0.9999]** | 0.9166 | [0.136, 1] |
| | ωλ0_9 | 0.8283 | [0.162, 0.9999] | 0.9735 | [0.4365, 1] |
| Shrinkage weights (extinction) | ωμ0_0 | **0.9129** | **[0.5991, 1]** | 0.9847 | [0.2676, 1] |
| | ωμ0_1 | **0.9449** | **[0.6875, 1]** | 0.9977 | [0.5123, 1] |
| | ωμ0_2 | 0.8924 | [0.3324, 1] | **0.9925** | **[0.9506, 1]** |
| | ωμ0_3 | 0.4525 | [1.9236E-7, 0.975] | 0.973 | [0.4843, 1] |
| | ωμ0_4 | **0.9576** | **[0.7952, 1]** | **0.998** | **[0.9853, 1]** |
| | ωμ0_5 | **0.9446** | **[0.6752, 1]** | **0.9913** | **[0.9382, 1]** |
| | ωμ0_6 | **0.9581** | **[0.7928, 1]** | **0.9914** | **[0.9165, 1]** |
| | ωμ0_7 | 0.4859 | [1.1306E-8, 0.9773] | **0.9941** | **[0.9633, 1]** |
| | ωμ0_8 | 0.5004 | [2.5659E-7, 0.9771] | 0.9195 | [0.1017, 1] |
| | ωμ0_9 | 0.7403 | [0.0301, 1] | **0.9966** | **[0.9625, 1]** |

**table S4.** Continued.

| Parameters | | Triassic Median | Triassic 95% HPD | Jurassic Median | Jurassic 95% HPD |
|---|---|---|---|---|---|
| Baseline rates | λ0 | 2.21E-01 | [0.0108, 0.9181] | 4.78E-01 | [0.0217, 1.526] |
| | μ0 | 2.99E-01 | [1.8704E-3, 1.0886] | 5.39E-01 | [0.0228, 1.6788] |
| Correlation parameters to origination | Gλ0_0 | -0.0038 | [-10.6793, 8.6734] | **-67.138** | **[-88.3445, -49.036]** |
| | Gλ0_1 | 8.64E-07 | [-0.0099, 0.0122] | 2.18E-04 | [-0.1246, 0.4631] |
| | Gλ0_2 | -0.3971 | [-9.1481, 3.226] | 13.0076 | [-0.6283, 21.0183] |
| | Gλ0_3 | -0.0148 | [-25.6159, 23.5578] | **65.7389** | **[18.3455, 111.3595]** |
| | Gλ0_4 | -0.0006 | [-0.0567, 0.0401] | -0.3491 | [-1.0481, 0.1879] |
| | Gλ0_5 | 2.59E-04 | [-0.1579, 0.176] | -3.38E-01 | [-1.5043, 0.3368] |
| | Gλ0_6 | -0.3011 | [-5.4275, 2.4101] | **-18.578** | **[-33.8032, -2.775]** |
| | Gλ0_7 | -0.0005 | [-0.1104, 0.0896] | -0.2984 | [-1.6624, 0.2768] |
| | Gλ0_8 | -0.025 | [-2.7377, 2.4139] | 2.4373 | [-1.8966, 14.1531] |
| | Gλ0_9 | -0.0064 | [-0.1062, 0.0356] | -0.0217 | [-0.2096, 0.1313] |
| Correlation parameters to extinction | Gμ0_0 | 1.73E-03 | [-8.464, 8.7555] | 1.42E+01 | [-6.158, 57.1351] |
| | Gμ0_1 | 0 | [-0.0109, 0.0112] | 3.60E-06 | [-0.0788, 0.0974] |
| | Gμ0_2 | -0.4588 | [-24.1485, 7.114] | **-25.7884** | **[-46.179, -7.7802]** |
| | Gμ0_3 | -0.0286 | [-47.7688, 37.4676] | 9.6244 | [-51.4902, 134.1075] |
| | Gμ0_4 | -0.0008 | [-0.1499, 0.085] | 0.1435 | [-0.2338, 0.7755] |
| | Gμ0_5 | -0.003 | [-0.8086, 0.3296] | **1.0598** | **[0.0307, 1.9835]** |
| | Gμ0_6 | -0.5989 | [-21.4366, 7.1889] | **25.2368** | **[2.6897, 52.5827]** |
| | Gμ0_7 | -4.00E-04 | [-0.3146, 0.1667] | 4.18E-01 | [-0.298, 2.4374] |
| | Gμ0_8 | -0.0384 | [-8.9944, 4.4306] | 3.256 | [-2.5543, 10.5433] |
| | Gμ0_9 | -0.0313 | [-0.5236, 0.1024] | -0.2883 | [-0.6304, 0.0169] |
| Shrinkage weights (origination) | ωλ0_0 | 0.3731 | [1.0193E-10, 0.9934] | **0.9997** | **[0.9986, 1]** |
| | ωλ0_1 | 0.3721 | [1.7159E-10, 0.9923] | 0.9902 | [0.3261, 1] |
| | ωλ0_2 | 0.4062 | [1.1987E-10, 0.9801] | 0.9876 | [0.6991, 1] |
| | ωλ0_3 | 0.3354 | [2.5058E-9, 0.9841] | **0.996** | **[0.9666, 1]** |
| | ωλ0_4 | 0.2731 | [5.6601E-11, 0.951] | 0.9899 | [0.4088, 1] |
| | ωλ0_5 | 0.2551 | [6.876E-9, 0.9598] | 0.9678 | [0.1649, 1] |
| | ωλ0_6 | 0.3205 | [2.1294E-9, 0.9564] | **0.9911** | **[0.908, 1]** |
| | ωλ0_7 | 0.3515 | [4.2583E-8, 0.9785] | 0.9967 | [0.482, 1] |
| | ωλ0_8 | 0.2788 | [2.6005E-9, 0.9493] | 0.9627 | [0.1482, 1] |
| | ωλ0_9 | 0.3639 | [2.6901E-11, 0.9637] | 0.9325 | [0.0821, 1] |
| Shrinkage weights (extinction) | ωμ0_0 | 0.3781 | [6.4403E-10, 0.9914] | 0.996 | [0.5447, 1] |
| | ωμ0_1 | 0.3768 | [6.4041E-11, 0.9922] | 0.9881 | [0.281, 1] |
| | ωμ0_2 | 0.5512 | [9.88E-11, 0.9962] | **0.996** | **[0.9682, 1]** |
| | ωμ0_3 | 0.3708 | [7.1122E-10, 0.9936] | 0.9848 | [0.3041, 1] |
| | ωμ0_4 | 0.3582 | [2.2795E-10, 0.9853] | 0.9817 | [0.2657, 1] |
| | ωμ0_5 | 0.3753 | [9.5234E-9, 0.9947] | **0.9894** | **[0.8718, 1]** |
| | ωμ0_6 | 0.5929 | [1.8508E-9, 0.9957] | **0.9945** | **[0.9397, 1]** |
| | ωμ0_7 | 0.3844 | [3.882E-9, 0.9945] | 0.9973 | [0.5359, 1] |
| | ωμ0_8 | 0.3739 | [6.1143E-9, 0.9872] | 0.9695 | [0.2704, 1] |
| | ωμ0_9 | 0.7481 | [4.7967E-3, 1] | 0.9901 | [0.8512, 1] |

**table S4.** Continued.

| Parameters | | Lower Cretaceous | | Upper Cretaceous | |
|---|---|---|---|---|---|
| | | Median | 95% HPD | Median | 95% HPD |
| Baseline rates | λ0 | 5.23E-01 | [0.0246, 1.5159] | 3.80E-01 | [4.0132E-3, 1.2795] |
| | μ0 | 4.03E-01 | [8.8295E-3, 1.3048] | 3.83E-01 | [7.0983E-3, 1.2843] |
| Correlation parameters to origination | Gλ0_0 | -0.12 | [-25.5395, 14.1019] | -0.3924 | [-7.6251, 6.7996] |
| | Gλ0_1 | 6.74E-04 | [-0.0302, 0.0909] | 2.79E-03 | [-0.0427, 0.08] |
| | Gλ0_2 | -7.3316 | [-57.4706, 10.9711] | **112.4052** | **[34.577, 188.0156]** |
| | Gλ0_3 | 1.7331 | [-19.477, 36.7482] | **229.0668** | **[71.8229, 373.9341]** |
| | Gλ0_4 | -0.2595 | [-0.4817, 7.8176E-3] | **-0.8174** | **[-1.3932, -0.1964]** |
| | Gλ0_5 | -4.85E-02 | [-0.6654, 0.3575] | -3.20E-01 | [-0.9492, 0.2397] |
| | Gλ0_6 | **-28.953** | **[-61.4481, -5.1014]** | -309.2292 | [-649.5878, 14.7876] |
| | Gλ0_7 | **0.2218** | **[0.0572, 0.4158]** | 8.59E-03 | [-0.4358, 0.4186] |
| | Gλ0_8 | 0.1 | [-2.0017, 2.6309] | -3.4842 | [-6.3677, 0.0205] |
| | Gλ0_9 | 0.31 | [-0.0515, 1.1872] | **-0.5389** | **[-0.9504, -0.1757]** |
| Correlation parameters to extinction | Gμ0_0 | -3.03E-01 | [-35.4991, 11.9685] | 1.09E+01 | [-0.2777, 21.0758] |
| | Gμ0_1 | -1.60E-03 | [-0.149, 0.1373] | 1.04E-01 | [-0.0322, 0.4189] |
| | Gμ0_2 | 1.2485 | [-12.3145, 50.4372] | 53.5208 | [-43.0351, 229.8193] |
| | Gμ0_3 | 19.6975 | [-28.385, 73.5007] | -314.5638 | [-658.1034, 73.7577] |
| | Gμ0_4 | -0.0437 | [-0.6239, 0.3902] | **2.8163** | **[0.7125, 4.6536]** |
| | Gμ0_5 | -0.8274 | [-2.0124, 0.0691] | -0.989 | [-3.7018, 0.3866] |
| | Gμ0_6 | 0.2109 | [-25.1526, 25.5951] | -695.9619 | [-1292.0602, 13.8204] |
| | Gμ0_7 | 1.65E-03 | [-0.1167, 0.2058] | **1.71E+00** | **[0.4104, 3.1337]** |
| | Gμ0_8 | -0.4005 | [-4.5548, 2.3911] | 0.4113 | [-2.1636, 4.0952] |
| | Gμ0_9 | -0.1007 | [-0.7585, 0.1321] | 0.3427 | [-0.7072, 2.1951] |
| Shrinkage weights (origination) | ωλ0_0 | 0.907 | [0.039, 1] | 0.9731 | [0.1804, 1] |
| | ωλ0_1 | 0.9397 | [0.0515, 1] | 0.9949 | [0.5565, 1] |
| | ωλ0_2 | 0.9727 | [0.0598, 1] | **0.999** | **[0.9919, 1]** |
| | ωλ0_3 | 0.8044 | [0.0222, 1] | **0.9992** | **[0.9923, 1]** |
| | ωλ0_4 | 0.974 | [0.6793, 1] | **0.9979** | **[0.9785, 1]** |
| | ωλ0_5 | 0.7708 | [0.0221, 1] | 0.9676 | [0.2354, 1] |
| | ωλ0_6 | **0.9919** | **[0.8773, 1]** | 0.9995 | [0.9863, 1] |
| | ωλ0_7 | **0.9809** | **[0.8116, 1]** | 0.9724 | [0.1886, 1] |
| | ωλ0_8 | 0.6665 | [0.0129, 1] | 0.9813 | [0.6567, 1] |
| | ωλ0_9 | 0.9874 | [0.1598, 1] | **0.9969** | **[0.9711, 1]** |
| Shrinkage weights (extinction) | ωμ0_0 | 0.9199 | [0.0425, 1] | 0.9954 | [0.9253, 1] |
| | ωμ0_1 | 0.9825 | [0.0748, 1] | 0.9998 | [0.8455, 1] |
| | ωμ0_2 | 0.8916 | [0.0341, 1] | 0.9977 | [0.7136, 1] |
| | ωμ0_3 | 0.9426 | [0.1573, 1] | 0.9995 | [0.916, 1] |
| | ωμ0_4 | 0.9082 | [0.0376, 1] | **0.9997** | **[0.9971, 1]** |
| | ωμ0_5 | 0.9741 | [0.6832, 1] | 0.992 | [0.654, 1] |
| | ωμ0_6 | 0.8849 | [0.0401, 1] | 0.9999 | [0.992, 1] |
| | ωμ0_7 | 0.7638 | [0.019, 1] | **0.9989** | **[0.9906, 1]** |
| | ωμ0_8 | 0.7664 | [0.0225, 1] | 0.9354 | [0.0636, 1] |
| | ωμ0_9 | 0.9403 | [0.0776, 1] | 0.9973 | [0.6689, 1] |

**table S4.** Continued.

| Parameters | | Cenozoic Median | Cenozoic 95% HPD |
|---|---|---|---|
| Baseline rates | λ0 | 1.96E-03 | [6.7429E-6, 0.0453] |
| | μ0 | 2.25E-01 | [9.5831E-4, 0.8809] |
| Correlation parameters to origination | Gλ0_0 | -3.0476 | [-5.7806, 1.0368] |
| | Gλ0_1 | 8.92E-03 | [-0.0001, 0.016] |
| | Gλ0_2 | 0.7396 | [-41.37, 68.4875] |
| | Gλ0_3 | **-45.3853** | **[-59.6256, -32.9653]** |
| | Gλ0_4 | 0.0329 | [-0.0107, 0.0913] |
| | Gλ0_5 | -1.32E-01 | [-1.0611, 1.1073] |
| | Gλ0_6 | 23.742 | [9.2484, 42.2323] |
| | Gλ0_7 | **3.17E-01** | **[0.1412, 0.4337]** |
| | Gλ0_8 | **-1.6509** | **[-3.0629, -0.1885]** |
| | Gλ0_9 | **0.2639** | **[-0.0255, 0.452]** |
| Correlation parameters to extinction | Gμ0_0 | **-5.66E+00** | **[-11.5234, -2.249]** |
| | Gμ0_1 | 1.80E-03 | [-0.0035, 0.0161] |
| | Gμ0_2 | -15.8796 | [-63.4943, 7.8747] |
| | Gμ0_3 | 1.6911 | [-12.21, 21.0937] |
| | Gμ0_4 | **0.1388** | **[0.0326, 0.2213]** |
| | Gμ0_5 | -0.3838 | [-1.5547, 0.1297] |
| | Gμ0_6 | 3.0371 | [-10.4895, 35.2223] |
| | Gμ0_7 | 9.94E-02 | [-0.0267, 0.2963] |
| | Gμ0_8 | 0.1202 | [-1.9736, 2.7395] |
| | Gμ0_9 | **-0.3216** | **[-0.4962, -0.0722]** |
| Shrinkage weights (origination) | ωλ0_0 | 0.9202 | [0.2226, 1] |
| | ωλ0_1 | 0.9662 | [0.5461, 1] |
| | ωλ0_2 | 0.9036 | [0.0464, 1] |
| | ωλ0_3 | **0.9771** | **[0.8954, 1]** |
| | ωλ0_4 | 0.6967 | [0.0264, 0.9999] |
| | ωλ0_5 | 0.8536 | [0.0796, 1] |
| | ωλ0_6 | 0.9414 | [0.6501, 1] |
| | ωλ0_7 | **0.9531** | **[0.7366, 1]** |
| | ωλ0_8 | **0.7994** | **[0.2015, 1]** |
| | ωλ0_9 | **0.9497** | **[0.3233, 1]** |
| Shrinkage weights (extinction) | ωμ0_0 | **0.9716** | **[0.8168, 1]** |
| | ωμ0_1 | 0.8623 | [0.0304, 1] |
| | ωμ0_2 | 0.9111 | [0.0569, 1] |
| | ωμ0_3 | 0.6864 | [0.015, 0.9998] |
| | ωμ0_4 | **0.9312** | **[0.6159, 1]** |
| | ωμ0_5 | 0.8713 | [0.0504, 1] |
| | ωμ0_6 | 0.7533 | [0.0174, 1] |
| | ωμ0_7 | 0.8213 | [0.074, 1] |
| | ωμ0_8 | 0.5919 | [2.2057E-9, 0.9827] |
| | ωμ0_9 | **0.9645** | **[0.7661, 1]** |

**table S5.**

Posterior parameter estimates for the MBD model applied to Adephaga genera, considering singletons and amber occurrences, across multiple temporal windows. The MBD model estimates the baseline origination and extinction rates ($\lambda 0$ and $\mu 0$), the correlation parameters ($G\lambda$ and $G\mu$) for each variable, and the shrinkage weights ($\omega$) of the correlation parameters. A variable was considered to have a significant effect (positive or negative depending on the sign of $G\lambda$ or $G\mu$) when its shrinkage weight exceeded 0.5 and when the 95% HPD interval of the corresponding correlation parameter did not overlap with zero (values highlighted in bold). The drivers are numbered as follows: (0) diversity of Adephaga genera through time, (1) angiosperms diversity through time, (2) global variation of atmospheric $CO_2$ through time, (3) continental fragmentation through time, (4) gymnosperms diversity through time, (5) global variation in $\delta^{34}S$ through time (used here as an inverted proxy for global magmatic activity), (6) global variation of atmospheric $O_2$ through time, (7) Pteridophytes diversity through time, (8) Sea level fluctuations through time, and (9) variation of the global mean temperature through time. "All" corresponds to the time window encompassing the entire evolutionary history of Adephaga genera, around 254 Ma to the present. "Before Upper Cretaceous" spans 254–100.5 Ma. The other time intervals are defined as follows: Triassic (251.902–201.4 Ma), Jurassic (201.4–143.1 Ma), Lower Cretaceous (143.1–100.5 Ma), Upper Cretaceous (100.5–66 Ma), and Cenozoic (66 Ma to the present).

| Parameters | | All | | Before Upper Cretaceous | |
|---|---|---|---|---|---|
| | | Median | 95% HPD | Median | 95% HPD |
| Baseline rates | λ0 | 0.0213 | [6.9007E-4, 0.0847] | 0.4656 | [0.0501, 1.1573] |
| | μ0 | 0.0899 | [2.1906E-3, 0.3676] | 0.1528 | [7.9642E-4, 0.5941] |
| Correlation parameters to origination | Gλ0_0 | **-3.2927** | **[-4.4344, -2.1699]** | -0.0899 | [-2.9665, 1.1858] |
| | Gλ0_1 | **2.61E-03** | **[6.2135E-4, 4.5184E-3]** | 3.48E-05 | [-0.0046, 8.5424E-3] |
| | Gλ0_2 | 0.1163 | [-1.4112, 2.2633] | 0.1663 | [-1.2617, 2.8596] |
| | Gλ0_3 | **-10.5229** | **[-15.6379, -5.1237]** | 0.2094 | [-3.837, 5.5336] |
| | Gλ0_4 | **0.0336** | **[2.9641E-3, 0.0608]** | **-0.0616** | **[-0.1066, -0.0189]** |
| | Gλ0_5 | 0.0226 | [-0.0372, 0.1398] | -0.0006 | [-0.0676, 0.055] |
| | Gλ0_6 | 0.8592 | [-0.4526, 2.535] | -0.1402 | [-1.9755, 0.8362] |
| | Gλ0_7 | **0.0482** | **[0.026, 0.0707]** | 0.0109 | [-0.0065, 0.0416] |
| | Gλ0_8 | -1.1736 | [-2.8733, 0.2551] | 0.0214 | [-1.2661, 1.6319] |
| | Gλ0_9 | 2.63E-04 | [-0.0403, 0.043] | 3.12E-03 | [-0.019, 0.0484] |
| Correlation parameters to extinction | Gμ0_0 | 0.7958 | [-0.474, 2.68] | 1.50E-04 | [-1.5301, 1.7236] |
| | Gμ0_1 | **-0.0053** | **[-0.0085, -0.0026]** | 2.50E-05 | [-0.0062, 6.6492E-3] |
| | Gμ0_2 | 0.1856 | [-1.7169, 2.9368] | -0.0819 | [-2.6441, 1.7689] |
| | Gμ0_3 | 0.8166 | [-4.46, 8.8675] | 5.8389 | [-2.0818, 15.2856] |
| | Gμ0_4 | -0.0313 | [-0.064, 2.2687E-3] | -0.0397 | [-0.084, 4.2344E-3] |
| | Gμ0_5 | 1.19E-03 | [-0.0628, 0.0724] | 5.25E-04 | [-0.0564, 0.0709] |
| | Gμ0_6 | 1.7013 | [-0.2381, 3.7416] | 0.4795 | [-0.774, 2.8952] |
| | Gμ0_7 | 0.031 | [-0.0007, 0.0581] | 0.0112 | [-0.0111, 0.0531] |
| | Gμ0_8 | -1.7501 | [-3.3881, 0.047] | -0.7886 | [-4.0345, 0.5352] |
| | Gμ0_9 | 3.88E-03 | [-0.0312, 0.0517] | 8.04E-03 | [-0.0178, 0.076] |
| Shrinkage weights (origination) | ωλ0_0 | **0.9077** | **[0.6554, 1]** | 0.2172 | [8.0567E-11, 0.9357] |
| | ωλ0_1 | **0.8218** | **[0.3433, 1]** | 0.2829 | [2.0844E-8, 0.9794] |
| | ωλ0_2 | 0.2863 | [4.3516E-12, 0.9369] | 0.1704 | [8.2025E-10, 0.8956] |
| | ωλ0_3 | **0.8578** | **[0.4968, 1]** | 0.1496 | [6.8872E-10, 0.8684] |
| | ωλ0_4 | **0.781** | **[0.2202, 1]** | **0.8865** | **[0.4858, 1]** |
| | ωλ0_5 | 0.5219 | [4.5074E-8, 0.9694] | 0.1521 | [9.2564E-8, 0.8823] |
| | ωλ0_6 | 0.4665 | [1.0304E-7, 0.958] | 0.1316 | [1.7187E-11, 0.8506] |
| | ωλ0_7 | **0.8471** | **[0.474, 1]** | 0.3159 | [2.5771E-10, 0.9271] |
| | ωλ0_8 | 0.6551 | [0.0363, 1] | 0.1547 | [1.3085E-9, 0.8752] |
| | ωλ0_9 | 0.3441 | [6.3749E-9, 0.9408] | 0.1682 | [6.3139E-9, 0.8832] |
| Shrinkage weights (extinction) | ωμ0_0 | 0.5872 | [3.4458E-10, 0.9728] | 0.1763 | [3.7523E-8, 0.8985] |
| | ωμ0_1 | **0.9383** | **[0.7147, 1]** | 0.2686 | [5.1872E-8, 0.9783] |
| | ωμ0_2 | 0.3527 | [1.0634E-8, 0.9484] | 0.1619 | [2.0223E-9, 0.8763] |
| | ωμ0_3 | 0.4164 | [5.9333E-8, 0.9555] | 0.619 | [1.5121E-8, 0.9741] |
| | ωμ0_4 | 0.7622 | [0.0855, 0.9999] | 0.7693 | [0.0249, 1] |
| | ωμ0_5 | 0.3264 | [2.8006E-8, 0.9412] | 0.1545 | [1.3838E-8, 0.8759] |
| | ωμ0_6 | 0.6588 | [0.0557, 0.9999] | 0.2458 | [3.3676E-10, 0.9149] |
| | ωμ0_7 | 0.7325 | [0.1302, 1] | 0.3877 | [1.9238E-8, 0.944] |
| | ωμ0_8 | 0.7619 | [0.14, 1] | 0.4828 | [7.6705E-8, 0.9689] |
| | ωμ0_9 | 0.3721 | [8.9381E-10, 0.947] | 0.2812 | [1.3207E-7, 0.9363] |

**table S5.** Continued.

| Parameters | | Triassic | | Jurassic | |
|---|---|---|---|---|---|
| | | Median | 95% HPD | Median | 95% HPD |
| Baseline rates | λ0 | 0.3742 | [0.0134, 1.0511] | 0.1932 | [0.0219, 0.7318] |
| | μ0 | 0.2775 | [4.2091E-3, 0.9472] | 0.1248 | [6.7698E-3, 0.5522] |
| Correlation parameters to origination | Gλ0_0 | -2.9935 | [-65.4126, 3.9133] | -0.5121 | [-19.63, 2.6042] |
| | Gλ0_1 | 3.68E-06 | [-0.0137, 0.0114] | 0.00E+00 | [-0.0092, 8.6502E-3] |
| | Gλ0_2 | 0.1425 | [-2.9149, 6.6609] | 6.41E-03 | [-2.3983, 2.8287] |
| | Gλ0_3 | -0.106 | [-35.8585, 31.7768] | -0.8236 | [-23.7524, 8.0688] |
| | Gλ0_4 | -0.0026 | [-0.0986, 0.0458] | -0.0013 | [-0.1296, 0.1591] |
| | Gλ0_5 | 0.0101 | [-0.0895, 0.2222] | -0.1646 | [-0.7119, 0.0661] |
| | Gλ0_6 | -0.5676 | [-8.5294, 2.3277] | -1.8464 | [-15.5571, 1.8214] |
| | Gλ0_7 | -0.0004 | [-0.1223, 0.0873] | -0.0001 | [-0.1221, 0.1005] |
| | Gλ0_8 | -0.1717 | [-5.2848, 2.444] | 0.0213 | [-2.2126, 4.271] |
| | Gλ0_9 | 2.31E-03 | [-0.0437, 0.0811] | -2.10E-03 | [-0.0642, 0.0334] |
| Correlation parameters to extinction | Gμ0_0 | -9.00E-03 | [-9.0464, 7.8318] | 3.82E-04 | [-3.4279, 3.9216] |
| | Gμ0_1 | 0.00E+00 | [-0.017, 0.0162] | 0.00E+00 | [-0.0099, 0.0107] |
| | Gμ0_2 | -0.0831 | [-7.0542, 4.2113] | -1.4216 | [-6.1231, 0.8339] |
| | Gμ0_3 | 0.1374 | [-25.4265, 34.9965] | 0.2025 | [-8.6761, 17.8718] |
| | Gμ0_4 | -0.0051 | [-0.119, 0.0438] | -0.0012 | [-0.0954, 0.0767] |
| | Gμ0_5 | 1.61E-01 | [-0.0205, 0.3772] | -1.74E-02 | [-0.3297, 0.1133] |
| | Gμ0_6 | -0.4901 | [-8.4747, 2.4365] | -0.0308 | [-4.6645, 3.2892] |
| | Gμ0_7 | -0.0009 | [-0.1679, 0.1075] | 3.66E-04 | [-0.101, 0.1295] |
| | Gμ0_8 | -0.8166 | [-9.1315, 1.9063] | -0.079 | [-3.6297, 1.814] |
| | Gμ0_9 | -1.00E-04 | [-0.0739, 0.0727] | -3.50E-03 | [-0.0735, 0.0312] |
| Shrinkage weights (origination) | ωλ0_0 | 0.9466 | [0.0102, 1] | 0.5275 | [7.3567E-9, 0.9977] |
| | ωλ0_1 | 0.4583 | [1.0414E-8, 0.9938] | 0.2545 | [1.3484E-8, 0.9884] |
| | ωλ0_2 | 0.3292 | [1.0063E-9, 0.9647] | 0.1516 | [2.1239E-9, 0.9128] |
| | ωλ0_3 | 0.4578 | [5.1196E-8, 0.99] | 0.3233 | [1.7603E-9, 0.9789] |
| | ωλ0_4 | 0.4048 | [2.1674E-9, 0.9735] | 0.247 | [1.7882E-9, 0.9691] |
| | ωλ0_5 | 0.3694 | [1.4607E-7, 0.9613] | 0.6106 | [5.1938E-11, 0.9865] |
| | ωλ0_6 | 0.4761 | [1.6918E-8, 0.9776] | 0.542 | [5.6276E-9, 0.9906] |
| | ωλ0_7 | 0.3917 | [6.2098E-9, 0.9808] | 0.2314 | [9.9818E-9, 0.975] |
| | ωλ0_8 | 0.4593 | [8.2786E-8, 0.9755] | 0.2083 | [1.265E-8, 0.96] |
| | ωλ0_9 | 0.3036 | [8.2505E-11, 0.952] | 0.1581 | [1.8224E-8, 0.8988] |
| Shrinkage weights (extinction) | ωμ0_0 | 0.4485 | [6.4318E-9, 0.9919] | 0.2213 | [3.0575E-11, 0.9687] |
| | ωμ0_1 | 0.4607 | [1.0996E-8, 0.9962] | 0.2514 | [7.113E-9, 0.9918] |
| | ωμ0_2 | 0.3602 | [4.5921E-10, 0.9693] | 0.4722 | [5.8996E-8, 0.9668] |
| | ωμ0_3 | 0.432 | [8.5677E-9, 0.9898] | 0.2055 | [1.509E-8, 0.9647] |
| | ωμ0_4 | 0.5035 | [2.7863E-8, 0.9826] | 0.184 | [8.325E-11, 0.9411] |
| | ωμ0_5 | 0.8467 | [0.0279, 0.9999] | 0.2004 | [3.1156E-8, 0.9366] |
| | ωμ0_6 | 0.4523 | [2.9115E-10, 0.9761] | 0.1788 | [4.2846E-8, 0.9257] |
| | ωμ0_7 | 0.423 | [2.3026E-13, 0.9859] | 0.2267 | [2.5328E-9, 0.9748] |
| | ωμ0_8 | 0.669 | [3.6893E-8, 0.9915] | 0.2265 | [1.4356E-8, 0.9534] |
| | ωμ0_9 | 0.3029 | [3.3699E-9, 0.9567] | 0.1754 | [1.4457E-8, 0.9188] |

**table S5.** Continued.

| Parameters | | Lower Cretaceous | | Upper Cretaceous | |
|---|---|---|---|---|---|
| | | Median | 95% HPD | Median | 95% HPD |
| Baseline rates | λ0 | 0.3101 | [0.0125, 1.1255] | 0.3946 | [0.0133, 1.3522] |
| | μ0 | 0.3005 | [0.0116, 1.0297] | 0.3676 | [3.813E-3, 1.2985] |
| Correlation parameters to origination | Gλ0_0 | -0.0515 | [-3.4509, 2.0152] | -0.1048 | [-23.736, 5.9804] |
| | Gλ0_1 | 1.90E-05 | [-0.0058, 8.8047E-3] | 4.11E-05 | [-0.007, 9.41E-3] |
| | Gλ0_2 | -1.59E+00 | [-9.9213, 1.9131] | -9.17E-01 | [-18.9877, 8.7977] |
| | Gλ0_3 | -0.3003 | [-16.0877, 8.9011] | -0.0002 | [-24.4137, 22.6159] |
| | Gλ0_4 | -0.0384 | [-0.1726, 0.0353] | -0.0016 | [-0.2839, 0.1689] |
| | Gλ0_5 | 8.32E-03 | [-0.1485, 0.2823] | -1.55E-01 | [-1.3187, 0.2732] |
| | Gλ0_6 | 0.0333 | [-3.0146, 3.7908] | -0.0698 | [-36.9837, 32.0064] |
| | Gλ0_7 | 1.39E-03 | [-0.0296, 0.0479] | 7.01E-03 | [-0.1659, 0.4061] |
| | Gλ0_8 | 0.0501 | [-1.1984, 2.2667] | -0.6235 | [-8.8921, 2.4902] |
| | Gλ0_9 | 6.37E-05 | [-0.0776, 0.0926] | -7.30E-03 | [-0.2029, 0.1003] |
| Correlation parameters to extinction | Gμ0_0 | -6.05E-01 | [-5.8897, 0.8601] | -1.00E-02 | [-7.1771, 6.1786] |
| | Gμ0_1 | 0.00E+00 | [-0.008, 7.1787E-3] | -9.00E-04 | [-0.0457, 7.0773E-3] |
| | Gμ0_2 | -0.0394 | [-4.6904, 3.5711] | 0.3899 | [-10.8893, 20.5129] |
| | Gμ0_3 | 4.6845 | [-5.9518, 33.6673] | -19.747 | [-204.0447, 11.5154] |
| | Gμ0_4 | -0.0057 | [-0.1225, 0.0504] | 0.0157 | [-0.1995, 1.9538] |
| | Gμ0_5 | **-7.30E-01** | **[-1.2929, -0.2248]** | -2.59E-02 | [-0.7076, 0.2649] |
| | Gμ0_6 | 0.0366 | [-7.3579, 6.2796] | -0.5588 | [-89.0522, 45.0422] |
| | Gμ0_7 | -1.10E-03 | [-0.0538, 0.0341] | -3.70E-03 | [-0.3377, 0.2234] |
| | Gμ0_8 | -0.0204 | [-2.2292, 1.6578] | -0.0422 | [-3.9382, 3.1015] |
| | Gμ0_9 | -3.10E-03 | [-0.1243, 0.0491] | 7.08E-03 | [-0.1184, 0.3164] |
| Shrinkage weights (origination) | ωλ0_0 | 0.2451 | [1.2688E-9, 0.9535] | 0.5465 | [3.1685E-3, 1] |
| | ωλ0_1 | 0.258 | [2.4393E-9, 0.9842] | 0.4578 | [7.5173E-8, 0.9868] |
| | ωλ0_2 | 0.5363 | [5.0708E-8, 0.9804] | 0.5074 | [7.5768E-9, 0.9837] |
| | ωλ0_3 | 0.1734 | [1.8962E-9, 0.941] | 0.3757 | [9.0403E-9, 0.977] |
| | ωλ0_4 | 0.547 | [3.6301E-9, 0.9757] | 0.4228 | [1.1321E-8, 0.9874] |
| | ωλ0_5 | 0.17 | [2.7219E-10, 0.9143] | 0.6493 | [1.8067E-8, 0.99] |
| | ωλ0_6 | 0.1428 | [1.5148E-8, 0.9001] | 0.4226 | [1.2444E-8, 0.9862] |
| | ωλ0_7 | 0.1403 | [1.3871E-8, 0.8921] | 0.4374 | [6.5528E-9, 0.9874] |
| | ωλ0_8 | 0.1743 | [7.3134E-10, 0.9083] | 0.6283 | [1.2289E-9, 0.9913] |
| | ωλ0_9 | 0.1886 | [1.9512E-8, 0.9344] | 0.5076 | [8.4074E-11, 0.9821] |
| Shrinkage weights (extinction) | ωμ0_0 | 0.5037 | [1.4711E-9, 0.9839] | 0.458 | [1.8722E-9, 0.9891] |
| | ωμ0_1 | 0.2685 | [5.1914E-9, 0.9815] | 0.7986 | [6.3519E-3, 1] |
| | ωμ0_2 | 0.1938 | [2.3976E-8, 0.9327] | 0.46 | [1.9267E-9, 0.9836] |
| | ωμ0_3 | 0.4744 | [2.4445E-8, 0.981] | 0.9088 | [0.0109, 1] |
| | ωμ0_4 | 0.2626 | [7.893E-10, 0.94] | 0.6729 | [4.0738E-3, 1] |
| | ωμ0_5 | **0.9517** | **[0.7062, 1]** | 0.3535 | [1.6085E-7, 0.9738] |
| | ωμ0_6 | 0.2206 | [7.796E-8, 0.9504] | 0.4991 | [5.766E-10, 0.9957] |
| | ωμ0_7 | 0.1661 | [6.2608E-9, 0.9036] | 0.4165 | [7.2468E-8, 0.9848] |
| | ωμ0_8 | 0.1906 | [3.0076E-10, 0.9174] | 0.3782 | [4.1728E-11, 0.9712] |
| | ωμ0_9 | 0.2212 | [1.5744E-8, 0.9488] | 0.47 | [1.6261E-9, 0.9912] |

**table S5.** Continued.

| Parameters | | Cenozoic Median | Cenozoic 95% HPD |
|---|---|---|---|
| Baseline rates | λ0 | 0.115 | [1.0171E-3, 0.5048] |
| | μ0 | 0.2154 | [2.1218E-3, 0.8581] |
| Correlation parameters to origination | Gλ0_0 | -1.1996 | [-3.9879, 0.4347] |
| | Gλ0_1 | 1.21E-04 | [-0.0028, 4.6538E-3] |
| | Gλ0_2 | -1.07E+00 | [-24.6767, 10.5583] |
| | Gλ0_3 | **-30.6562** | **[-51.1014, -10.485]** |
| | Gλ0_4 | **0.1069** | **[0.0234, 0.188]** |
| | Gλ0_5 | 4.99E-02 | [-0.2011, 0.4789] |
| | Gλ0_6 | -1.442 | [-17.4392, 9.4547] |
| | Gλ0_7 | 8.42E-02 | [-0.019, 0.2492] |
| | Gλ0_8 | -0.4827 | [-4.6378, 1.1759] |
| | Gλ0_9 | 1.28E-01 | [-0.0258, 0.3675] |
| Correlation parameters to extinction | Gμ0_0 | -1.08E-02 | [-3.2215, 2.8017] |
| | Gμ0_1 | -5.20E-03 | [-0.0111, 6.7499E-4] |
| | Gμ0_2 | -2.3851 | [-40.7566, 11.9307] |
| | Gμ0_3 | 0.2069 | [-15.5914, 19.3637] |
| | Gμ0_4 | 2.51E-03 | [-0.0496, 0.0753] |
| | Gμ0_5 | -2.34E-02 | [-0.7868, 0.4061] |
| | Gμ0_6 | 1.57 | [-6.4299, 16.9297] |
| | Gμ0_7 | 1.84E-02 | [-0.0769, 0.2421] |
| | Gμ0_8 | -0.1466 | [-3.6682, 2.3559] |
| | Gμ0_9 | -1.76E-01 | [-0.385, 0.0243] |
| Shrinkage weights (origination) | ωλ0_0 | 0.6708 | [3.7656E-8, 0.9794] |
| | ωλ0_1 | 0.378 | [2.0014E-8, 0.9586] |
| | ωλ0_2 | 0.4078 | [4.376E-8, 0.9681] |
| | ωλ0_3 | **0.9405** | **[0.6799, 1]** |
| | ωλ0_4 | **0.8578** | **[0.4069, 1]** |
| | ωλ0_5 | 0.355 | [1.7554E-9, 0.9499] |
| | ωλ0_6 | 0.4359 | [1.172E-7, 0.9589] |
| | ωλ0_7 | 0.645 | [8.8693E-10, 0.9788] |
| | ωλ0_8 | 0.4888 | [1.4316E-7, 0.9744] |
| | ωλ0_9 | 0.8143 | [0.0336, 1] |
| Shrinkage weights (extinction) | ωμ0_0 | 0.4442 | [1.566E-8, 0.9676] |
| | ωμ0_1 | 0.8901 | [0.1104, 1] |
| | ωμ0_2 | 0.5366 | [1.8063E-9, 0.9856] |
| | ωμ0_3 | 0.417 | [1.4425E-9, 0.9618] |
| | ωμ0_4 | 0.2875 | [2.4048E-9, 0.9386] |
| | ωμ0_5 | 0.3977 | [2.7422E-8, 0.9686] |
| | ωμ0_6 | 0.3891 | [1.24E-7, 0.9571] |
| | ωμ0_7 | 0.4092 | [2.9314E-9, 0.9651] |
| | ωμ0_8 | 0.4092 | [4.552E-8, 0.9586] |
| | ωμ0_9 | 0.8808 | [0.0981, 1] |

**table S6.**

Posterior parameter estimates for the MBD model applied to Adephaga genera, excluding singletons, but considering amber occurrences, across multiple temporal windows. The MBD model estimates the baseline origination and extinction rates ($\lambda 0$ and $\mu 0$), the correlation parameters ($G\lambda$ and $G\mu$) for each variable, and the shrinkage weights ($\omega$) of the correlation parameters. A variable was considered to have a significant effect (positive or negative depending on the sign of $G\lambda$ or $G\mu$) when its shrinkage weight exceeded 0.5 and when the 95% HPD interval of the corresponding correlation parameter did not overlap with zero (values highlighted in bold). The drivers are numbered as follows: (0) diversity of Adephaga genera through time, (1) angiosperms diversity through time, (2) global variation of atmospheric $CO_2$ through time, (3) continental fragmentation through time, (4) gymnosperms diversity through time, (5) global variation in $\delta^{34}S$ through time (used here as an inverted proxy for global magmatic activity), (6) global variation of atmospheric $O_2$ through time, (7) Pteridophytes diversity through time, (8) Sea level fluctuations through time, and (9) variation of the global mean temperature through time. "All" corresponds to the time window encompassing the entire evolutionary history of Adephaga genera, around 254 Ma to the present. "Before Upper Cretaceous" spans 254–100.5 Ma. The other time intervals are defined as follows: Triassic (251.902–201.4 Ma), Jurassic (201.4–143.1 Ma), Lower Cretaceous (143.1–100.5 Ma), Upper Cretaceous (100.5–66 Ma), and Cenozoic (66 Ma to the present).

| Parameters | | All | | Before Upper Cretaceous | |
|---|---|---|---|---|---|
| | | Median | 95% HPD | Median | 95% HPD |
| Baseline rates | λ0 | 0.061 | [2.1201E-3, 0.2774] | 0.536 | [0.0358, 1.4466] |
| | μ0 | 0.2325 | [4.896E-3, 0.7254] | 0.3045 | [7.2413E-3, 0.9403] |
| Correlation parameters to origination | Gλ0_0 | **-5.6964** | **[-7.3133, -4.2017]** | -0.0341 | [-3.9707, 2.4925] |
| | Gλ0_1 | **5.15E-03** | **[2.5385E-3, 7.4309E-3]** | -3.00E-04 | [-0.2378, 0.0152] |
| | Gλ0_2 | 0.1544 | [-1.5737, 2.4908] | 1.6327 | [-0.6697, 5.081] |
| | Gλ0_3 | **-7.4852** | **[-13.9896, -0.8352]** | -0.3868 | [-9.7336, 4.8141] |
| | Gλ0_4 | 0.0225 | [-0.0062, 0.0556] | **-0.0749** | **[-0.1268, -0.0147]** |
| | Gλ0_5 | 0.0573 | [-0.0296, 0.1927] | -0.0003 | [-0.0808, 0.0716] |
| | Gλ0_6 | 0.1153 | [-1.3277, 2.2179] | -0.6357 | [-3.1023, 0.748] |
| | Gλ0_7 | 9.02E-03 | [-0.0103, 0.0405] | -5.00E-04 | [-0.0401, 0.0445] |
| | Gλ0_8 | -1.0631 | [-2.949, 0.3662] | -0.2486 | [-3.9112, 1.2818] |
| | Gλ0_9 | -0.0176 | [-0.0964, 0.0224] | 5.80E-04 | [-0.0346, 0.0443] |
| Correlation parameters to extinction | Gμ0_0 | 0.3938 | [-1.1705, 3.0818] | -0.0223 | [-2.8139, 2.2481] |
| | Gμ0_1 | **-0.0056** | **[-0.01, -0.0018]** | 2.35E-05 | [-0.0071, 0.011] |
| | Gμ0_2 | 0.9904 | [-1.4563, 5.3782] | -0.034 | [-3.3749, 2.771] |
| | Gμ0_3 | 7.1473 | [-1.1305, 16.7803] | **16.4099** | **[5.0536, 31.0661]** |
| | Gμ0_4 | **-0.0827** | **[-0.1254, -0.0373]** | **-0.0861** | **[-0.1351, -0.0198]** |
| | Gμ0_5 | 4.54E-03 | [-0.0654, 0.0975] | 1.26E-02 | [-0.0379, 0.1343] |
| | Gμ0_6 | 0.6144 | [-1.0208, 3.4211] | 0.0165 | [-1.8805, 2.3459] |
| | Gμ0_7 | 0.0185 | [-0.0092, 0.064] | -0.0025 | [-0.0588, 0.0324] |
| | Gμ0_8 | -0.9266 | [-3.9962, 0.8282] | -0.4429 | [-5.0306, 1.2998] |
| | Gμ0_9 | -0.0027 | [-0.063, 0.0362] | 4.17E-03 | [-0.0322, 0.0684] |
| Shrinkage weights (origination) | ωλ0_0 | **0.9637** | **[0.843, 1]** | 0.3403 | [2.1966E-8, 0.967] |
| | ωλ0_1 | **0.9345** | **[0.7104, 1]** | 0.6388 | [4.9036E-3, 1] |
| | ωλ0_2 | 0.3142 | [1.015E-7, 0.9415] | 0.5562 | [3.1865E-9, 0.969] |
| | ωλ0_3 | **0.7845** | **[0.2134, 1]** | 0.2834 | [3.8334E-9, 0.9329] |
| | ωλ0_4 | 0.683 | [0.0312, 0.9999] | **0.9224** | **[0.5259, 1]** |
| | ωλ0_5 | 0.7063 | [0.0193, 0.9999] | 0.2467 | [5.3842E-8, 0.9356] |
| | ωλ0_6 | 0.3071 | [4.4799E-9, 0.9398] | 0.3584 | [4.871E-8, 0.9384] |
| | ωλ0_7 | 0.4347 | [9.0365E-9, 0.9591] | 0.2851 | [4.55E-9, 0.9408] |
| | ωλ0_8 | 0.6346 | [6.9967E-10, 0.9733] | 0.3893 | [3.3815E-8, 0.964] |
| | ωλ0_9 | 0.5668 | [1.0846E-9, 0.9771] | 0.2301 | [9.2994E-9, 0.9138] |
| Shrinkage weights (extinction) | ωμ0_0 | 0.5184 | [1.2743E-9, 0.9722] | 0.3122 | [9.308E-9, 0.9532] |
| | ωμ0_1 | **0.9437** | **[0.6808, 1]** | 0.4348 | [4.3224E-9, 0.9897] |
| | ωμ0_2 | 0.5609 | [6.2482E-10, 0.9737] | 0.261 | [1.1935E-8, 0.9355] |
| | ωμ0_3 | 0.7741 | [0.0513, 1] | **0.9239** | **[0.5864, 0.9999]** |
| | ωμ0_4 | **0.9396** | **[0.7153, 1]** | **0.9369** | **[0.5974, 1]** |
| | ωμ0_5 | 0.3793 | [1.7848E-7, 0.954] | 0.3371 | [8.0869E-9, 0.9514] |
| | ωμ0_6 | 0.4485 | [1.7698E-8, 0.9617] | 0.2159 | [1.308E-8, 0.9106] |
| | ωμ0_7 | 0.6244 | [8.8552E-7, 0.9749] | 0.322 | [4.5982E-11, 0.9522] |
| | ωμ0_8 | 0.651 | [3.5412E-9, 0.9804] | 0.472 | [2.5401E-9, 0.9765] |
| | ωμ0_9 | 0.3945 | [1.5715E-8, 0.9559] | 0.2973 | [7.2188E-10, 0.9384] |

**table S6.** Continued.

| Parameters | | Triassic | | Jurassic | |
|---|---|---|---|---|---|
| | | Median | 95% HPD | Median | 95% HPD |
| Baseline rates | λ0 | 0.391 | [0.0197, 1.1221] | 0.1693 | [0.0157, 0.6857] |
| | μ0 | 0.3178 | [6.241E-3, 1.0175] | 0.0986 | [2.0958E-3, 0.5447] |
| Correlation parameters to origination | Gλ0_0 | -23.3632 | [-112.4116, 4.4802] | -0.192 | [-10.3794, 3.0666] |
| | Gλ0_1 | 1.28E-06 | [-0.0137, 0.0182] | 7.80E-07 | [-0.0114, 0.0106] |
| | Gλ0_2 | 0.1347 | [-4.2983, 6.3429] | 0.027 | [-2.3422, 3.5806] |
| | Gλ0_3 | -0.2899 | [-78.6475, 38.378] | -1.7241 | [-34.0023, 7.455] |
| | Gλ0_4 | -0.0022 | [-0.0988, 0.0565] | -0.004 | [-0.1315, 0.0871] |
| | Gλ0_5 | 7.81E-03 | [-0.1166, 0.2055] | -9.61E-02 | [-0.6196, 0.0873] |
| | Gλ0_6 | -0.048 | [-7.3503, 6.1584] | -0.4369 | [-10.1219, 2.8304] |
| | Gλ0_7 | -5.00E-04 | [-0.1388, 0.0989] | -1.40E-03 | [-0.245, 0.1268] |
| | Gλ0_8 | -0.2814 | [-8.5328, 3.5377] | 1.36E-03 | [-2.8164, 3.0853] |
| | Gλ0_9 | 1.35E-03 | [-0.0631, 0.0736] | -1.72E-02 | [-0.1078, 0.0237] |
| Correlation parameters to extinction | Gμ0_0 | -0.0097 | [-12.1762, 8.2847] | 0.0145 | [-4.5834, 6.6364] |
| | Gμ0_1 | 6.63E-06 | [-0.0144, 0.0153] | 2.78E-06 | [-0.0112, 0.0125] |
| | Gμ0_2 | -0.0967 | [-9.6229, 5.4266] | -1.9599 | [-10.8189, 1.0706] |
| | Gμ0_3 | 0.2438 | [-33.4851, 53.7364] | 0.8594 | [-8.6579, 38.5132] |
| | Gμ0_4 | -0.0236 | [-0.1895, 0.0388] | 1.78E-04 | [-0.1274, 0.1365] |
| | Gμ0_5 | 1.34E-01 | [-0.0377, 0.4592] | -2.86E-02 | [-0.6439, 0.1905] |
| | Gμ0_6 | -0.1451 | [-7.7905, 3.9236] | -0.4307 | [-12.5602, 3.6166] |
| | Gμ0_7 | -0.0071 | [-0.5223, 0.095] | 1.07E-03 | [-0.134, 0.2668] |
| | Gμ0_8 | -0.4266 | [-9.6117, 2.659] | -0.0586 | [-4.7874, 2.7094] |
| | Gμ0_9 | -1.70E-03 | [-0.114, 0.0723] | -7.80E-03 | [-0.1191, 0.0342] |
| Shrinkage weights (origination) | ωλ0_0 | 0.9973 | [0.0227, 1] | 0.4419 | [7.0286E-8, 0.9922] |
| | ωλ0_1 | 0.5487 | [4.3358E-3, 1] | 0.3367 | [9.13E-10, 0.9915] |
| | ωλ0_2 | 0.397 | [8.2084E-9, 0.9717] | 0.1963 | [9.5787E-9, 0.9346] |
| | ωλ0_3 | 0.5616 | [4.4876E-3, 1] | 0.4798 | [2.5164E-9, 0.9886] |
| | ωλ0_4 | 0.4495 | [8.3877E-9, 0.9788] | 0.314 | [6.5884E-9, 0.9568] |
| | ωλ0_5 | 0.3987 | [2.1843E-7, 0.9685] | 0.4778 | [5.4179E-10, 0.9795] |
| | ωλ0_6 | 0.4805 | [2.6884E-7, 0.9758] | 0.3186 | [2.757E-10, 0.9747] |
| | ωλ0_7 | 0.4749 | [1.0655E-7, 0.9853] | 0.3766 | [8.4334E-9, 0.9887] |
| | ωλ0_8 | 0.6339 | [4.4634E-8, 0.9895] | 0.2717 | [4.5541E-9, 0.9574] |
| | ωλ0_9 | 0.3426 | [4.2429E-8, 0.9598] | 0.394 | [5.811E-9, 0.9599] |
| Shrinkage weights (extinction) | ωμ0_0 | 0.5284 | [4.1823E-10, 0.9943] | 0.3194 | [1.6096E-10, 0.9849] |
| | ωμ0_1 | 0.5293 | [4.8382E-3, 1] | 0.3359 | [7.1571E-9, 0.9928] |
| | ωμ0_2 | 0.4525 | [3.0893E-9, 0.9821] | 0.6275 | [1.7906E-9, 0.988] |
| | ωμ0_3 | 0.5233 | [4.0241E-9, 0.9941] | 0.399 | [7.017E-11, 0.9895] |
| | ωμ0_4 | 0.7867 | [0.0104, 1] | 0.2704 | [7.0997E-9, 0.964] |
| | ωμ0_5 | 0.8302 | [0.0131, 0.9999] | 0.346 | [5.9418E-10, 0.9749] |
| | ωμ0_6 | 0.4163 | [1.1579E-8, 0.974] | 0.3526 | [4.5975E-10, 0.9814] |
| | ωμ0_7 | 0.6658 | [6.2653E-3, 1] | 0.3396 | [9.7005E-9, 0.9918] |
| | ωμ0_8 | 0.6005 | [5.0526E-8, 0.9913] | 0.2987 | [2.9446E-9, 0.9725] |
| | ωμ0_9 | 0.4141 | [9.0709E-10, 0.9727] | 0.3071 | [3.2668E-10, 0.9559] |

**table S6.** Continued.

| Parameters | | Lower Cretaceous | | Upper Cretaceous | |
|---|---|---|---|---|---|
| | | Median | 95% HPD | Median | 95% HPD |
| Baseline rates | λ0 | 0.3718 | [6.3846E-3, 1.277] | 0.3303 | [9.5199E-3, 1.0637] |
| | μ0 | 0.3289 | [1.1519E-3, 1.1478] | 0.2721 | [3.5036E-3, 0.9778] |
| Correlation parameters to origination | Gλ0_0 | -0.1194 | [-9.9261, 5.0718] | -0.03 | [-13.7543, 6.834] |
| | Gλ0_1 | 0.00E+00 | [-0.017, 0.0131] | 1.70E-05 | [-0.006, 7.1304E-3] |
| | Gλ0_2 | -0.1213 | [-9.0719, 7.5397] | -2.1748 | [-24.8523, 4.4104] |
| | Gλ0_3 | -1.8097 | [-44.3363, 17.4551] | 0.033 | [-16.4686, 16.9667] |
| | Gλ0_4 | -0.0541 | [-0.3276, 0.0732] | -0.0002 | [-0.1679, 0.1385] |
| | Gλ0_5 | 4.77E-02 | [-0.2256, 0.792] | -8.56E-02 | [-1.2587, 0.2676] |
| | Gλ0_6 | -4.7888 | [-30.1776, 4.391] | -0.2255 | [-30.9541, 21.6019] |
| | Gλ0_7 | 1.53E-03 | [-0.0831, 0.2041] | 1.24E-05 | [-0.1672, 0.1823] |
| | Gλ0_8 | 5.34E-03 | [-3.9481, 4.2225] | -1.22E-01 | [-5.3423, 2.4145] |
| | Gλ0_9 | -1.57E-02 | [-0.2606, 0.0945] | -5.30E-03 | [-0.1857, 0.0853] |
| Correlation parameters to extinction | Gμ0_0 | -0.2625 | [-7.8027, 3.1359] | -0.0192 | [-9.5473, 6.8976] |
| | Gμ0_1 | 0.00E+00 | [-0.0135, 0.014] | 0.00E+00 | [-0.0078, 6.0851E-3] |
| | Gμ0_2 | -0.2224 | [-10.4016, 7.0268] | -0.2544 | [-16.7421, 9.7391] |
| | Gμ0_3 | 6.7675 | [-9.0194, 41.0382] | -0.6887 | [-28.1627, 14.3774] |
| | Gμ0_4 | -6.27E-02 | [-0.2612, 0.051] | -2.00E-04 | [-0.1972, 0.198] |
| | Gμ0_5 | -1.71E-01 | [-0.9614, 0.1261] | -9.00E-03 | [-0.6682, 0.369] |
| | Gμ0_6 | 1.8547 | [-5.08, 13.7331] | -1.0528 | [-49.3906, 17.8544] |
| | Gμ0_7 | -1.20E-03 | [-0.0938, 0.074] | -1.02E-02 | [-0.3021, 0.1119] |
| | Gμ0_8 | -0.3183 | [-6.0482, 2.6034] | -0.062 | [-4.8316, 2.7378] |
| | Gμ0_9 | -8.10E-03 | [-0.2355, 0.0917] | -2.70E-03 | [-0.1521, 0.0912] |
| Shrinkage weights (origination) | ωλ0_0 | 0.5626 | [4.771E-10, 0.991] | 0.3035 | [2.4526E-8, 0.9942] |
| | ωλ0_1 | 0.5713 | [5.5886E-3, 1] | 0.2871 | [1.9527E-9, 0.9798] |
| | ωλ0_2 | 0.5164 | [4.3993E-8, 0.9817] | 0.5444 | [1.6938E-8, 0.9879] |
| | ωλ0_3 | 0.5691 | [9.4578E-8, 0.9876] | 0.2347 | [9.1964E-9, 0.9508] |
| | ωλ0_4 | 0.7716 | [0.0138, 1] | 0.2494 | [1.0402E-8, 0.9685] |
| | ωλ0_5 | 0.5449 | [7.3598E-9, 0.9817] | 0.4709 | [9.3233E-9, 0.9868] |
| | ωλ0_6 | 0.8236 | [0.0192, 1] | 0.2739 | [1.0939E-8, 0.9743] |
| | ωλ0_7 | 0.4636 | [2.5738E-11, 0.9871] | 0.2485 | [6.6853E-10, 0.9655] |
| | ωλ0_8 | 0.4587 | [7.4225E-10, 0.9749] | 0.3092 | [9.5993E-10, 0.9748] |
| | ωλ0_9 | 0.6168 | [7.3405E-9, 0.9866] | 0.3511 | [3.9168E-10, 0.975] |
| Shrinkage weights (extinction) | ωμ0_0 | 0.6185 | [5.1936E-8, 0.9885] | 0.3004 | [2.5545E-8, 0.9911] |
| | ωμ0_1 | 0.552 | [2.3399E-9, 0.9946] | 0.2863 | [5.6306E-9, 0.9827] |
| | ωμ0_2 | 0.5129 | [2.955E-10, 0.9837] | 0.2951 | [3.2512E-8, 0.9717] |
| | ωμ0_3 | 0.7098 | [1.2717E-8, 0.9895] | 0.3129 | [6.3701E-9, 0.9686] |
| | ωμ0_4 | 0.7788 | [0.0145, 0.9999] | 0.2842 | [3.8529E-8, 0.9783] |
| | ωμ0_5 | 0.7319 | [3.7253E-7, 0.9898] | 0.2287 | [2.6347E-8, 0.9581] |
| | ωμ0_6 | 0.6702 | [1.5332E-7, 0.9866] | 0.3701 | [4.2754E-8, 0.986] |
| | ωμ0_7 | 0.4342 | [5.2823E-11, 0.9698] | 0.3572 | [1.8742E-8, 0.9751] |
| | ωμ0_8 | 0.5708 | [2.1729E-8, 0.9839] | 0.2816 | [1.5775E-11, 0.9639] |
| | ωμ0_9 | 0.5248 | [2.1915E-9, 0.9862] | 0.3006 | [7.9777E-9, 0.9645] |

**table S6.** Continued.

| Parameters | | Cenozoic | |
|---|---|---|---|
| | | Median | 95% HPD |
| Baseline rates | λ0 | 0.1576 | [1.8934E-3, 0.6924] |
| | μ0 | 0.2335 | [1.8039E-3, 0.9848] |
| Correlation parameters to origination | Gλ0_0 | -0.9086 | [-5.2682, 0.873] |
| | Gλ0_1 | 6.07E-06 | [-0.0037, 4.1873E-3] |
| | Gλ0_2 | 1.1682 | [-9.3828, 19.8456] |
| | Gλ0_3 | **-31.8229** | **[-52.3329, -7.7694]** |
| | Gλ0_4 | **0.1105** | **[0.0316, 0.1885]** |
| | Gλ0_5 | 9.77E-03 | [-0.2673, 0.5034] |
| | Gλ0_6 | -2.2418 | [-20.0314, 6.9103] |
| | Gλ0_7 | 6.63E-02 | [-0.0195, 0.1802] |
| | Gλ0_8 | -3.81E-02 | [-1.7778, 1.6303] |
| | Gλ0_9 | 3.33E-02 | [-0.0548, 0.2489] |
| Correlation parameters to extinction | Gμ0_0 | -0.31 | [-5.4085, 1.811] |
| | Gμ0_1 | -3.50E-03 | [-0.0107, 1.255E-3] |
| | Gμ0_2 | -1.0032 | [-23.9829, 14.8065] |
| | Gμ0_3 | -2.0021 | [-30.6494, 11.5981] |
| | Gμ0_4 | -1.00E-04 | [-0.0715, 0.0729] |
| | Gμ0_5 | -4.77E-02 | [-0.8549, 0.3363] |
| | Gμ0_6 | 0.6071 | [-10.0847, 18.6828] |
| | Gμ0_7 | -2.30E-03 | [-0.1479, 0.1088] |
| | Gμ0_8 | -0.8516 | [-6.5721, 1.7001] |
| | Gμ0_9 | -4.13E-02 | [-0.3101, 0.0844] |
| Shrinkage weights (origination) | ωλ0_0 | 0.6201 | [1.3343E-7, 0.9803] |
| | ωλ0_1 | 0.3422 | [1.6283E-10, 0.9567] |
| | ωλ0_2 | 0.3821 | [3.4696E-7, 0.9584] |
| | ωλ0_3 | **0.9422** | **[0.6545, 1]** |
| | ωλ0_4 | **0.857** | **[0.4159, 1]** |
| | ωλ0_5 | 0.2733 | [7.4497E-11, 0.9365] |
| | ωλ0_6 | 0.4246 | [3.697E-9, 0.9572] |
| | ωλ0_7 | 0.5494 | [6.98E-12, 0.963] |
| | ωλ0_8 | 0.2422 | [4.4405E-9, 0.9149] |
| | ωλ0_9 | 0.5255 | [2.7415E-8, 0.9755] |
| Shrinkage weights (extinction) | ωμ0_0 | 0.5181 | [7.3809E-11, 0.9811] |
| | ωμ0_1 | 0.8156 | [0.0231, 1] |
| | ωμ0_2 | 0.4136 | [1.7756E-8, 0.9688] |
| | ωμ0_3 | 0.4624 | [2.7908E-8, 0.9757] |
| | ωμ0_4 | 0.2816 | [1.517E-8, 0.9302] |
| | ωμ0_5 | 0.4255 | [4.8031E-9, 0.965] |
| | ωμ0_6 | 0.321 | [9.3564E-10, 0.9547] |
| | ωμ0_7 | 0.2954 | [1.1899E-8, 0.9398] |
| | ωμ0_8 | 0.59 | [4.2878E-7, 0.9825] |
| | ωμ0_9 | 0.6156 | [2.6807E-8, 0.9824] |

**table S7.**

Posterior parameter estimates for the MBD model applied to Coleoptera genera without Polyphaga, considering singletons and amber occurrences, across multiple temporal windows. The MBD model estimates the baseline origination and extinction rates (λ0 and μ0), the correlation parameters (Gλ and Gμ) for each variable, and the shrinkage weights (ω) of the correlation parameters. A variable was considered to have a significant effect (positive or negative depending on the sign of Gλ or Gμ) when its shrinkage weight exceeded 0.5 and when the 95% HPD interval of the corresponding correlation parameter did not overlap with zero (values highlighted in bold). The drivers are numbered as follows: (0) diversity of Coleoptera genera without Polyphaga through time, (1) angiosperms diversity through time, (2) global variation of atmospheric $CO_2$ through time, (3) continental fragmentation through time, (4) gymnosperms diversity through time, (5) global variation in $\delta^{34}S$ through time (used here as an inverted proxy for global magmatic activity), (6) global variation of atmospheric $O_2$ through time, (7) Pteridophytes diversity through time, (8) Sea level fluctuations through time, and (9) variation of the global mean temperature through time. "All" corresponds to the time window encompassing the entire evolutionary history of Coleoptera genera without Polyphaga, around 300 Ma to the present. "Before Upper Cretaceous" spans 300–100.5 Ma. The other time intervals are defined as follows: Permian (298.9–251.902 Ma), Triassic (251.902–201.4 Ma), Jurassic (201.4–143.1 Ma), Lower Cretaceous (143.1–100.5 Ma), Upper Cretaceous (100.5–66 Ma), and Cenozoic (66 Ma to the present).

| Parameters | | All | | Before Upper Cretaceous | |
|---|---|---|---|---|---|
| | | Median | 95% HPD | Median | 95% HPD |
| Baseline rates | λ0 | 0.027 | [3.9048E-3, 0.0778] | 0.1826 | [0.0108, 0.5729] |
| | μ0 | 0.0283 | [2.2693E-3, 0.1308] | 0.1421 | [0.0101, 0.4015] |
| Correlation parameters to origination | Gλ0_0 | 0.7732 | [-0.0178, 1.4528] | **2.0118** | **[0.755, 3.2578]** |
| | Gλ0_1 | 2.84E-04 | [-0.0005, 1.794E-3] | 9.61E-04 | [-0.0052, 0.0309] |
| | Gλ0_2 | **-1.567** | **[-2.6453, -0.4698]** | **-1.6991** | **[-2.8893, -0.4065]** |
| | Gλ0_3 | -2.0339 | [-5.3174, 0.4164] | -1.3451 | [-7.2987, 2.4036] |
| | Gλ0_4 | 9.19E-04 | [-0.0097, 0.0154] | **-3.33E-02** | **[-0.058, -0.0074]** |
| | Gλ0_5 | **-0.0503** | **[-0.0968, -0.0043]** | -0.0247 | [-0.0739, 0.0113] |
| | Gλ0_6 | -0.8869 | [-1.7693, 0.0352] | -1.6842 | [-2.9559, 0.0133] |
| | Gλ0_7 | 3.33E-03 | [-0.0049, 0.0161] | 3.22E-04 | [-0.0144, 0.0167] |
| | Gλ0_8 | 6.56E-03 | [-0.7549, 0.688] | 8.17E-02 | [-0.7872, 1.3186] |
| | Gλ0_9 | **0.074** | **[0.0466, 0.1031]** | **0.0502** | **[9.5966E-3, 0.0854]** |
| Correlation parameters to extinction | Gμ0_0 | **2.5209** | **[1.7807, 3.3735]** | **3.9128** | **[2.6983, 5.0409]** |
| | Gμ0_1 | **-0.0059** | **[-0.008, -0.0039]** | **0.0598** | **[0.0376, 0.0815]** |
| | Gμ0_2 | **-2.692** | **[-4.0025, -1.335]** | **-3.0182** | **[-4.518, -1.5122]** |
| | Gμ0_3 | 0.3731 | [-2.1249, 4.1542] | -2.7745 | [-8.2504, 1.5278] |
| | Gμ0_4 | -0.0067 | [-0.0298, 7.0603E-3] | -0.0101 | [-0.0388, 7.9372E-3] |
| | Gμ0_5 | 0.0296 | [-0.0094, 0.0719] | 0.0571 | [-0.0001, 0.0961] |
| | Gμ0_6 | -0.1941 | [-1.2419, 0.4174] | 0.0548 | [-0.8627, 1.0988] |
| | Gμ0_7 | 0.0181 | [-0.0002, 0.0327] | -0.0044 | [-0.0296, 0.0141] |
| | Gμ0_8 | -1.0454 | [-2.0207, 6.2406E-3] | **-1.8858** | **[-3.1588, -0.4655]** |
| | Gμ0_9 | **0.0518** | **[0.0131, 0.0894]** | 4.86E-03 | [-0.0195, 0.0432] |
| Shrinkage weights (origination) | ωλ0_0 | 0.5029 | [0.0504, 0.9994] | **0.8292** | **[0.391, 0.9998]** |
| | ωλ0_1 | 0.2716 | [2.6721E-8, 0.9248] | 0.8121 | [0.0184, 1] |
| | ωλ0_2 | **0.5552** | **[0.1225, 0.9998]** | **0.6289** | **[0.1432, 0.9999]** |
| | ωλ0_3 | 0.3637 | [2.3487E-8, 0.9272] | 0.3914 | [3.2427E-8, 0.9495] |
| | ωλ0_4 | 0.2015 | [1.5446E-8, 0.9009] | **0.8413** | **[0.3576, 1]** |
| | ωλ0_5 | **0.6083** | **[0.1085, 0.9997]** | 0.4521 | [7.738E-8, 0.9559] |
| | ωλ0_6 | 0.541 | [0.0458, 0.9991] | 0.7748 | [0.1971, 0.9999] |
| | ωλ0_7 | 0.2199 | [4.3116E-9, 0.9048] | 0.2646 | [3.1818E-8, 0.9329] |
| | ωλ0_8 | 0.1709 | [3.7002E-9, 0.8933] | 0.3014 | [1.1569E-8, 0.9411] |
| | ωλ0_9 | **0.886** | **[0.6055, 1]** | **0.8183** | **[0.3245, 1]** |
| Shrinkage weights (extinction) | ωμ0_0 | **0.8594** | **[0.5397, 1]** | **0.9311** | **[0.7236, 1]** |
| | ωμ0_1 | **0.947** | **[0.7792, 1]** | **0.9994** | **[0.9968, 1]** |
| | ωμ0_2 | **0.7363** | **[0.3008, 1]** | **0.7893** | **[0.3593, 0.9999]** |
| | ωμ0_3 | 0.1943 | [9.1559E-9, 0.8896] | 0.5175 | [7.2052E-10, 0.9636] |
| | ωμ0_4 | 0.4018 | [5.6841E-11, 0.948] | 0.5578 | [4.7916E-10, 0.9703] |
| | ωμ0_5 | 0.4384 | [3.6347E-9, 0.9446] | 0.687 | [0.1312, 1] |
| | ωμ0_6 | 0.2449 | [1.3308E-10, 0.9053] | 0.2937 | [1.5105E-8, 0.9352] |
| | ωμ0_7 | 0.6107 | [0.0887, 0.9999] | 0.4301 | [1.2037E-8, 0.9537] |
| | ωμ0_8 | 0.602 | [0.0826, 0.9999] | **0.8047** | **[0.31, 1]** |
| | ωμ0_9 | **0.8075** | **[0.3174, 1]** | 0.3882 | [1.8102E-7, 0.954] |

**table S7.** Continued.

| Parameters | | Permian | | Triassic | |
|---|---|---|---|---|---|
| | | Median | 95% HPD | Median | 95% HPD |
| Baseline rates | λ0 | 0.6319 | [0.0748, 1.5512] | 0.5302 | [0.0142, 1.5697] |
| | μ0 | 0.6107 | [0.0371, 1.6571] | 0.4928 | [0.0164, 1.5595] |
| Correlation parameters to origination | Gλ0_0 | -10.7776 | [-18.8164, 0.3814] | -0.8886 | [-9.6018, 2.0696] |
| | Gλ0_1 | 1.66E-06 | [-0.0214, 0.0239] | 2.51E-05 | [-0.0355, 0.0343] |
| | Gλ0_2 | 0.3758 | [-2.6526, 5.5087] | 9.1752 | [-0.155, 18.6484] |
| | Gλ0_3 | -0.1575 | [-56.8224, 37.0506] | 38.7868 | [-8.1872, 98.037] |
| | Gλ0_4 | 1.63E-04 | [-0.1022, 0.1053] | 2.15E-02 | [-0.0517, 0.2623] |
| | Gλ0_5 | 0.0232 | [-0.0738, 0.2138] | 4.42E-03 | [-0.1736, 0.4148] |
| | Gλ0_6 | 0.1827 | [-1.5552, 5.8446] | **-12.2531** | **[-17.6024, -6.9918]** |
| | Gλ0_7 | -7.00E-04 | [-0.1465, 0.0867] | -6.24E-02 | [-0.3945, 0.0625] |
| | Gλ0_8 | 1.42E-01 | [-1.4966, 2.8231] | -2.66E-02 | [-3.3384, 3.3487] |
| | Gλ0_9 | -0.0097 | [-0.1325, 0.0326] | -0.015 | [-0.2274, 0.1216] |
| Correlation parameters to extinction | Gμ0_0 | -0.0956 | [-6.9727, 4.7567] | 3.067 | [-0.8716, 9.2918] |
| | Gμ0_1 | 9.16E-06 | [-0.0182, 0.0178] | 2.58E-05 | [-0.0534, 0.0651] |
| | Gμ0_2 | 0.4099 | [-3.1229, 7.8757] | -1.3914 | [-21.9136, 3.9162] |
| | Gμ0_3 | 0.7227 | [-24.7929, 50.3775] | 4.2482 | [-27.4006, 75.1171] |
| | Gμ0_4 | 8.18E-04 | [-0.1011, 0.1771] | 4.00E-03 | [-0.0755, 0.1799] |
| | Gμ0_5 | 0.0591 | [-0.0424, 0.2592] | 0.0581 | [-0.0908, 0.2497] |
| | Gμ0_6 | 4.0632 | [-0.5977, 9.8001] | **-8.6761** | **[-15.6233, -3.1229]** |
| | Gμ0_7 | -0.002 | [-0.247, 0.1141] | -0.101 | [-0.6843, 0.0565] |
| | Gμ0_8 | -0.0345 | [-2.8407, 1.9518] | -0.0722 | [-3.3435, 2.5887] |
| | Gμ0_9 | -1.51E-01 | [-0.2912, 9.8749E-3] | 4.66E-02 | [-0.0432, 0.3917] |
| Shrinkage weights (origination) | ωλ0_0 | 0.9879 | [0.7998, 1] | 0.8073 | [0.023, 1] |
| | ωλ0_1 | 0.6584 | [6.715E-3, 1] | 0.8698 | [0.0272, 1] |
| | ωλ0_2 | 0.4592 | [9.5384E-9, 0.9755] | 0.9652 | [0.6564, 1] |
| | ωλ0_3 | 0.6141 | [5.9375E-3, 1] | 0.9848 | [0.2099, 1] |
| | ωλ0_4 | 0.5725 | [6.7186E-8, 0.9908] | 0.8368 | [0.0234, 1] |
| | ωλ0_5 | 0.6211 | [3.3223E-8, 0.9843] | 0.6816 | [0.011, 0.9999] |
| | ωλ0_6 | 0.4863 | [2.809E-8, 0.9871] | **0.9785** | **[0.8853, 1]** |
| | ωλ0_7 | 0.5819 | [2.4935E-6, 0.9911] | 0.9436 | [0.054, 1] |
| | ωλ0_8 | 0.4425 | [3.8034E-7, 0.9702] | 0.6617 | [8.8563E-9, 0.9874] |
| | ωλ0_9 | 0.5292 | [9.1393E-8, 0.9851] | 0.7793 | [0.0223, 1] |
| Shrinkage weights (extinction) | ωμ0_0 | 0.6075 | [1.5784E-8, 0.9894] | 0.9251 | [0.0918, 1] |
| | ωμ0_1 | 0.6482 | [7.22E-3, 1] | 0.8809 | [0.0304, 1] |
| | ωμ0_2 | 0.5095 | [3.8945E-10, 0.9812] | 0.8023 | [0.0186, 1] |
| | ωμ0_3 | 0.686 | [6.2667E-3, 1] | 0.9072 | [0.032, 1] |
| | ωμ0_4 | 0.6381 | [8.3017E-3, 1] | 0.7246 | [0.0139, 1] |
| | ωμ0_5 | 0.7471 | [0.0109, 1] | 0.7738 | [0.0342, 1] |
| | ωμ0_6 | 0.9313 | [0.0418, 1] | **0.9613** | **[0.75, 1]** |
| | ωμ0_7 | 0.693 | [8.9681E-3, 1] | 0.965 | [0.0571, 1] |
| | ωμ0_8 | 0.4493 | [7.0966E-9, 0.9707] | 0.6554 | [3.9905E-10, 0.9876] |
| | ωμ0_9 | 0.9658 | [0.2809, 1] | 0.8663 | [0.0297, 1] |

**table S7.** Continued.

| Parameters | | Jurassic | | Lower Cretaceous | |
|---|---|---|---|---|---|
| | | Median | 95% HPD | Median | 95% HPD |
| Baseline rates | λ0 | 4.63E-04 | [2.5763E-9, 0.0399] | 9.43E-02 | [3.3407E-4, 0.5734] |
| | μ0 | 0.0362 | [4.935E-5, 0.3061] | 0.4699 | [0.0441, 1.291] |
| Correlation parameters to origination | Gλ0_0 | **4.8268** | **[3.0748, 6.5448]** | -0.128 | [-9.5252, 2.1236] |
| | Gλ0_1 | 0.00E+00 | [-0.0297, 0.0288] | 0.00E+00 | [-0.0142, 0.0116] |
| | Gλ0_2 | 4.5967 | [-3.1925, 13.5009] | -1.1032 | [-42.3351, 4.6016] |
| | Gλ0_3 | 6.7324 | [-28.9517, 66.8104] | -1.837 | [-23.2627, 10.4069] |
| | Gλ0_4 | -2.41E-01 | [-0.5325, 0.0167] | 9.91E-03 | [-0.1063, 1.032] |
| | Gλ0_5 | **-3.75E-01** | **[-0.708, -0.0566]** | -1.09E-01 | [-0.678, 0.1831] |
| | Gλ0_6 | -5.5924 | [-15.7992, 1.0186] | -0.0021 | [-6.5189, 7.8072] |
| | Gλ0_7 | -1.28E-02 | [-0.7936, 0.2796] | -1.62E-02 | [-0.2456, 0.0284] |
| | Gλ0_8 | 2.22E+00 | [-0.5894, 6.2776] | 5.56E-01 | [-1.1785, 4.0912] |
| | Gλ0_9 | **0.2125** | **[0.0422, 0.3859]** | 3.92E-03 | [-0.1055, 0.3593] |
| Correlation parameters to extinction | Gμ0_0 | **6.5686** | **[4.5083, 8.67]** | -0.3975 | [-3.3457, 1.0104] |
| | Gμ0_1 | 8.62E-06 | [-0.0368, 0.0359] | 0.00E+00 | [-0.0181, 0.0139] |
| | Gμ0_2 | -3.3111 | [-10.6383, 1.4406] | 0.3707 | [-4.008, 18.3997] |
| | Gμ0_3 | 10.8661 | [-6.3274, 44.9643] | 11.9429 | [-4.3299, 52.1344] |
| | Gμ0_4 | -1.04E-01 | [-0.371, 0.1304] | -1.23E-02 | [-0.284, 0.1131] |
| | Gμ0_5 | 0.1093 | [-0.0651, 0.3841] | **-0.8413** | **[-1.8293, -0.3286]** |
| | Gμ0_6 | 0.6189 | [-2.3955, 5.5057] | -0.6324 | [-18.52, 4.4955] |
| | Gμ0_7 | -0.0244 | [-0.7276, 0.205] | -0.0159 | [-0.1088, 0.0446] |
| | Gμ0_8 | -0.4823 | [-4.696, 1.8903] | -0.0428 | [-2.4615, 2.411] |
| | Gμ0_9 | 1.16E-02 | [-0.0646, 0.1251] | -2.40E-03 | [-0.2125, 0.1057] |
| Shrinkage weights (origination) | ωλ0_0 | **0.9621** | **[0.8194, 1]** | 0.4857 | [2.3876E-7, 0.9938] |
| | ωλ0_1 | 0.888 | [0.0385, 1] | 0.5157 | [3.9727E-9, 0.9952] |
| | ωλ0_2 | 0.9181 | [0.1296, 1] | 0.5874 | [3.46E-3, 1] |
| | ωλ0_3 | 0.9486 | [0.0871, 1] | 0.4453 | [3.2093E-9, 0.9754] |
| | ωλ0_4 | 0.9722 | [0.5529, 1] | 0.4827 | [2.7705E-3, 1] |
| | ωλ0_5 | **0.916** | **[0.497, 1]** | 0.6235 | [7.0543E-8, 0.9826] |
| | ωλ0_6 | 0.9136 | [0.1752, 1] | 0.3574 | [8.5134E-8, 0.9743] |
| | ωλ0_7 | 0.9073 | [0.0462, 1] | 0.462 | [1.3014E-9, 0.9912] |
| | ωλ0_8 | 0.8843 | [0.0917, 1] | 0.5067 | [6.019E-9, 0.978] |
| | ωλ0_9 | **0.971** | **[0.7655, 1]** | 0.4528 | [1.8812E-8, 0.9922] |
| Shrinkage weights (extinction) | ωμ0_0 | **0.9767** | **[0.8897, 1]** | 0.478 | [9.641E-9, 0.9766] |
| | ωμ0_1 | 0.8911 | [0.042, 1] | 0.5087 | [3.3634E-9, 0.9966] |
| | ωμ0_2 | 0.8774 | [0.106, 1] | 0.4072 | [2.4356E-8, 0.9923] |
| | ωμ0_3 | 0.919 | [0.0704, 1] | 0.7983 | [7.5875E-3, 1] |
| | ωμ0_4 | 0.9212 | [0.1263, 1] | 0.4817 | [2.0826E-8, 0.9878] |
| | ωμ0_5 | 0.7131 | [0.0233, 1] | **0.9678** | **[0.7965, 1]** |
| | ωμ0_6 | 0.6319 | [5.0136E-10, 0.9852] | 0.5172 | [8.4935E-8, 0.9899] |
| | ωμ0_7 | 0.9176 | [0.046, 1] | 0.5503 | [2.9571E-8, 0.9772] |
| | ωμ0_8 | 0.744 | [0.0192, 1] | 0.3722 | [1.2812E-7, 0.966] |
| | ωμ0_9 | 0.7037 | [0.0177, 1] | 0.3577 | [8.3686E-8, 0.9839] |

**table S7.** Continued.

| Parameters | | Upper Cretaceous | | Cenozoic | |
|---|---|---|---|---|---|
| | | Median | 95% HPD | Median | 95% HPD |
| Baseline rates | λ0 | 3.73E-01 | [9.5499E-3, 1.1382] | 7.82E-02 | [1.9445E-4, 0.3927] |
| | μ0 | 0.3677 | [4.5969E-3, 1.1639] | 0.1138 | [1.9144E-3, 0.5277] |
| Correlation parameters to origination | Gλ0_0 | -1.9661 | [-10.1387, 1.4739] | -0.5528 | [-3.9807, 0.8767] |
| | Gλ0_1 | -1.80E-03 | [-0.0428, 0.0123] | 2.16E-04 | [-0.003, 7.7146E-3] |
| | Gλ0_2 | 16.723 | [-6.5959, 49.5654] | -3.8058 | [-34.9798, 7.2289] |
| | Gλ0_3 | -0.8833 | [-47.4349, 44.1231] | **-32.4125** | **[-50.1102, -15.0317]** |
| | Gλ0_4 | -1.00E-02 | [-0.434, 0.2594] | **1.12E-01** | **[0.0381, 0.1793]** |
| | Gλ0_5 | -9.74E-01 | [-1.6394, 0.0169] | 1.55E-03 | [-0.566, 0.3946] |
| | Gλ0_6 | -1.8202 | [-94.458, 46.1331] | -0.3329 | [-12.2088, 10.9711] |
| | Gλ0_7 | 1.12E-02 | [-0.3202, 0.5611] | 8.71E-02 | [-0.0191, 0.2502] |
| | Gλ0_8 | -9.79E-02 | [-6.8559, 5.0125] | -1.43E-01 | [-4.9228, 1.7274] |
| | Gλ0_9 | -7.86E-02 | [-0.5684, 0.1121] | 1.47E-01 | [-0.0279, 0.4253] |
| Correlation parameters to extinction | Gμ0_0 | 1.5401 | [-1.6128, 7.9998] | -0.5629 | [-4.1984, 0.9586] |
| | Gμ0_1 | -1.00E-04 | [-0.0315, 0.0254] | -2.00E-03 | [-0.0073, 1.3083E-3] |
| | Gμ0_2 | -1.0369 | [-40.7272, 20.2493] | -0.5487 | [-17.7817, 9.5009] |
| | Gμ0_3 | -120.8352 | [-247.0163, 19.5044] | 0.8227 | [-9.9375, 20.0337] |
| | Gμ0_4 | 1.06E+00 | [-0.0491, 2.1664] | 9.61E-03 | [-0.03, 0.0797] |
| | Gμ0_5 | -0.0755 | [-1.7046, 0.514] | -0.0086 | [-0.4317, 0.3123] |
| | Gμ0_6 | -9.1048 | [-328.9778, 58.5026] | 1.2207 | [-6.1629, 13.4716] |
| | Gμ0_7 | 6.66E-03 | [-0.3596, 0.5051] | 1.10E-03 | [-0.0873, 0.0888] |
| | Gμ0_8 | -0.0315 | [-3.2099, 2.8833] | 0.0199 | [-2.0472, 2.6143] |
| | Gμ0_9 | 5.43E-02 | [-0.15, 0.9108] | -1.70E-01 | [-0.3311, 6.3881E-3] |
| Shrinkage weights (origination) | ωλ0_0 | 0.8999 | [0.0495, 1] | 0.4745 | [9.0755E-9, 0.974] |
| | ωλ0_1 | 0.9386 | [0.0577, 1] | 0.3797 | [2.4303E-8, 0.9775] |
| | ωλ0_2 | 0.9613 | [0.1812, 1] | 0.5103 | [1.2172E-7, 0.9831] |
| | ωλ0_3 | 0.8269 | [0.0245, 1] | **0.9457** | **[0.7473, 1]** |
| | ωλ0_4 | 0.8432 | [0.0257, 1] | **0.8641** | **[0.4701, 1]** |
| | ωλ0_5 | 0.966 | [0.6865, 1] | 0.2748 | [7.5713E-8, 0.9533] |
| | ωλ0_6 | 0.857 | [0.0314, 1] | 0.3038 | [1.826E-7, 0.942] |
| | ωλ0_7 | 0.8091 | [0.0244, 1] | 0.635 | [0.0251, 0.9999] |
| | ωλ0_8 | 0.8349 | [0.028, 1] | 0.3964 | [3.3996E-9, 0.9687] |
| | ωλ0_9 | 0.9219 | [0.0402, 1] | 0.8546 | [0.0256, 1] |
| Shrinkage weights (extinction) | ωμ0_0 | 0.8714 | [0.0403, 1] | 0.5274 | [4.9332E-8, 0.9762] |
| | ωμ0_1 | 0.8838 | [0.0356, 1] | 0.6944 | [6.912E-8, 0.982] |
| | ωμ0_2 | 0.8166 | [0.0249, 1] | 0.3339 | [7.0104E-7, 0.9529] |
| | ωμ0_3 | 0.9956 | [0.4271, 1] | 0.3605 | [9.365E-8, 0.9575] |
| | ωμ0_4 | 0.9977 | [0.7497, 1] | 0.3095 | [1.1478E-9, 0.9364] |
| | ωμ0_5 | 0.8283 | [0.026, 1] | 0.2807 | [4.0503E-8, 0.9411] |
| | ωμ0_6 | 0.9412 | [0.0459, 1] | 0.3239 | [1.3461E-8, 0.9448] |
| | ωμ0_7 | 0.8077 | [0.0201, 1] | 0.223 | [7.2326E-8, 0.9186] |
| | ωμ0_8 | 0.6667 | [1.113E-8, 0.9871] | 0.306 | [7.5518E-8, 0.9413] |
| | ωμ0_9 | 0.9205 | [0.0387, 1] | 0.8773 | [0.1861, 0.9999] |

**table S8.**

Posterior parameter estimates for the MBD model applied to Coleoptera genera without Polyphaga, excluding singletons, but considering amber occurrences, across multiple temporal windows. The MBD model estimates the baseline origination and extinction rates ($\lambda 0$ and $\mu 0$), the correlation parameters ($G\lambda$ and $G\mu$) for each variable, and the shrinkage weights ($\omega$) of the correlation parameters. A variable was considered to have a significant effect (positive or negative depending on the sign of $G\lambda$ or $G\mu$) when its shrinkage weight exceeded 0.5 and when the 95% HPD interval of the corresponding correlation parameter did not overlap with zero (values highlighted in bold). The drivers are numbered as follows: (0) diversity of Coleoptera genera without Polyphaga through time, (1) angiosperms diversity through time, (2) global variation of atmospheric $CO_2$ through time, (3) continental fragmentation through time, (4) gymnosperms diversity through time, (5) global variation in $\delta^{34}S$ through time (used here as an inverted proxy for global magmatic activity), (6) global variation of atmospheric $O_2$ through time, (7) Pteridophytes diversity through time, (8) Sea level fluctuations through time, and (9) variation of the global mean temperature through time. "All" corresponds to the time window encompassing the entire evolutionary history of Coleoptera genera without Polyphaga, around 300 Ma to the present. "Before Upper Cretaceous" spans 300–100.5 Ma. The other time intervals are defined as follows: Permian (298.9–251.902 Ma), Triassic (251.902–201.4 Ma), Jurassic (201.4–143.1 Ma), Lower Cretaceous (143.1–100.5 Ma), Upper Cretaceous (100.5–66 Ma), and Cenozoic (66 Ma to the present).

| Parameters | | All | | Before Upper Cretaceous | |
|---|---|---|---|---|---|
| | | Median | 95% HPD | Median | 95% HPD |
| Baseline rates | λ0 | 0.1552 | [0.0179, 0.411] | 0.3891 | [0.0566, 1.1233] |
| | μ0 | 0.0425 | [9.2736E-4, 0.1462] | 0.0666 | [8.2221E-4, 0.2632] |
| Correlation parameters to origination | Gλ0_0 | **-4.0426** | **[-5.018, -3.0538]** | **-6.1203** | **[-8.2237, -4.1876]** |
| | Gλ0_1 | **4.05E-03** | **[2.4344E-3, 5.7226E-3]** | 0.00E+00 | [-0.0082, 6.5273E-3] |
| | Gλ0_2 | 1.4035 | [-0.2194, 3.1273] | 2.0974 | [-0.0609, 4.0068] |
| | Gλ0_3 | **-4.4431** | **[-7.7777, -0.3934]** | 2.8193 | [-1.891, 11.8456] |
| | Gλ0_4 | 1.91E-04 | [-0.0115, 0.0146] | -2.40E-03 | [-0.0307, 0.0164] |
| | Gλ0_5 | -0.0166 | [-0.0674, 0.0141] | -0.041 | [-0.1, 6.6774E-3] |
| | Gλ0_6 | 0.2065 | [-0.4775, 1.4337] | -0.7468 | [-2.3212, 0.2761] |
| | Gλ0_7 | -0.0026 | [-0.0243, 9.5E-3] | -0.019 | [-0.059, 4.8619E-3] |
| | Gλ0_8 | -0.3961 | [-1.466, 0.3094] | 0.0192 | [-0.7925, 1.1315] |
| | Gλ0_9 | -0.0019 | [-0.0333, 0.0206] | 4.63E-04 | [-0.0263, 0.034] |
| Correlation parameters to extinction | Gμ0_0 | -0.9541 | [-2.2831, 0.2249] | **-3.326** | **[-5.2895, -1.014]** |
| | Gμ0_1 | -0.0017 | [-0.0045, 3.9558E-4] | 6.40E-05 | [-0.0084, 0.0115] |
| | Gμ0_2 | 2.433 | [-0.0876, 4.425] | 0.5818 | [-0.7497, 2.9414] |
| | Gμ0_3 | 0.566 | [-2.98, 6.7155] | 8.2475 | [-0.0205, 14.4809] |
| | Gμ0_4 | **-0.0392** | **[-0.0659, -0.0072]** | -0.0038 | [-0.0346, 0.0131] |
| | Gμ0_5 | 0.0202 | [-0.0136, 0.0766] | 0.037 | [-0.0108, 0.0926] |
| | Gμ0_6 | 0.0575 | [-1.0203, 1.5151] | -0.1507 | [-1.9596, 0.898] |
| | Gμ0_7 | **0.0326** | **[5.2494E-3, 0.0543]** | 8.87E-04 | [-0.019, 0.0282] |
| | Gμ0_8 | -0.5514 | [-1.9976, 0.3837] | -0.9521 | [-2.7641, 0.3511] |
| | Gμ0_9 | 4.89E-03 | [-0.0206, 0.0522] | 6.37E-03 | [-0.0218, 0.0622] |
| Shrinkage weights (origination) | ωλ0_0 | **0.9306** | **[0.7321, 1]** | **0.9662** | **[0.8508, 1]** |
| | ωλ0_1 | **0.8975** | **[0.6228, 1]** | 0.4586 | [3.2361E-10, 0.9858] |
| | ωλ0_2 | 0.5064 | [1.9325E-7, 0.9484] | 0.6487 | [0.0892, 0.9996] |
| | ωλ0_3 | **0.5867** | **[0.1156, 0.9995]** | 0.4983 | [8.7993E-8, 0.963] |
| | ωλ0_4 | 0.1855 | [3.2372E-8, 0.8916] | 0.3352 | [5.4565E-8, 0.9352] |
| | ωλ0_5 | 0.3018 | [2.1231E-10, 0.9245] | 0.5365 | [3.7436E-7, 0.9616] |
| | ωλ0_6 | 0.2533 | [1.5643E-9, 0.9159] | 0.4843 | [3.4567E-7, 0.9617] |
| | ωλ0_7 | 0.2467 | [9.6589E-13, 0.9133] | 0.6371 | [3.2712E-8, 0.9768] |
| | ωλ0_8 | 0.3235 | [2.2969E-8, 0.9242] | 0.2087 | [3.6388E-11, 0.9033] |
| | ωλ0_9 | 0.2309 | [4.8882E-8, 0.9025] | 0.2444 | [3.8982E-10, 0.9145] |
| Shrinkage weights (extinction) | ωμ0_0 | 0.5662 | [1.569E-6, 0.9623] | **0.902** | **[0.5327, 1]** |
| | ωμ0_1 | 0.6787 | [0.0256, 1] | 0.4934 | [4.1649E-7, 0.9906] |
| | ωμ0_2 | 0.6823 | [0.086, 0.9999] | 0.3127 | [9.875E-9, 0.9267] |
| | ωμ0_3 | 0.2594 | [1.2298E-8, 0.918] | 0.7828 | [0.2245, 1] |
| | ωμ0_4 | **0.8545** | **[0.3736, 1]** | 0.3537 | [5.398E-9, 0.9507] |
| | ωμ0_5 | 0.3552 | [4.2873E-9, 0.93] | 0.5023 | [3.2532E-8, 0.9572] |
| | ωμ0_6 | 0.2509 | [7.2832E-8, 0.9158] | 0.2979 | [1.9134E-8, 0.9395] |
| | ωμ0_7 | **0.7779** | **[0.2535, 0.9999]** | 0.274 | [8.8334E-10, 0.9261] |
| | ωμ0_8 | 0.4205 | [8.7185E-8, 0.9465] | 0.568 | [2.1678E-9, 0.9667] |
| | ωμ0_9 | 0.3186 | [2.0168E-7, 0.938] | 0.3583 | [4.8726E-8, 0.9492] |

**table S8.** Continued.

| Parameters | | Permian | | Triassic | |
|---|---|---|---|---|---|
| | | Median | 95% HPD | Median | 95% HPD |
| Baseline rates | λ0 | 0.292 | [0.013, 0.8962] | 0.2848 | [6.7129E-4, 1.0173] |
| | μ0 | 0.2352 | [0.0133, 0.9936] | 0.296 | [2.2478E-3, 1.0197] |
| Correlation parameters to origination | Gλ0_0 | -8.2257 | [-16.1083, 0.761] | -1.9124 | [-10.6082, 1.5145] |
| | Gλ0_1 | 4.38E-06 | [-0.0094, 8.5552E-3] | 1.88E-06 | [-0.017, 0.0169] |
| | Gλ0_2 | -0.0485 | [-3.9885, 3.0813] | 5.6852 | [-1.4716, 23.0846] |
| | Gλ0_3 | 0.0224 | [-20.0598, 17.2355] | -7.1114 | [-81.6785, 12.4182] |
| | Gλ0_4 | 5.12E-05 | [-0.0568, 0.0773] | -6.50E-03 | [-0.1059, 0.0475] |
| | Gλ0_5 | 4.38E-03 | [-0.0658, 0.1535] | 5.95E-04 | [-0.1437, 0.1482] |
| | Gλ0_6 | -0.0391 | [-1.9357, 1.1728] | -3.1128 | [-8.0754, 0.9556] |
| | Gλ0_7 | 3.49E-05 | [-0.0658, 0.0812] | 9.07E-04 | [-0.1172, 0.1999] |
| | Gλ0_8 | -0.0099 | [-1.9083, 1.632] | -0.1582 | [-4.4723, 2.5143] |
| | Gλ0_9 | 3.92E-05 | [-0.0417, 0.042] | -2.14E-02 | [-0.2567, 0.05] |
| Correlation parameters to extinction | Gμ0_0 | -0.0744 | [-6.5983, 2.4213] | 0.374 | [-2.4109, 7.5744] |
| | Gμ0_1 | 0.00E+00 | [-0.0087, 8.898E-3] | 1.81E-07 | [-0.0185, 0.0184] |
| | Gμ0_2 | -0.015 | [-3.3752, 2.932] | -0.3874 | [-10.6629, 3.9666] |
| | Gμ0_3 | 0.033 | [-15.7849, 17.9864] | -0.4427 | [-41.2575, 17.5528] |
| | Gμ0_4 | 0 | [-0.0586, 0.0554] | -0.0205 | [-0.1302, 0.0358] |
| | Gμ0_5 | 9.81E-03 | [-0.0559, 0.2099] | 8.11E-02 | [-0.0571, 0.2824] |
| | Gμ0_6 | -0.1236 | [-3.094, 1.2916] | -4.2329 | [-10.0007, 0.9651] |
| | Gμ0_7 | -2.00E-04 | [-0.0691, 0.0611] | -1.70E-03 | [-0.1652, 0.0957] |
| | Gμ0_8 | -0.0165 | [-2.7923, 2.0411] | -0.4181 | [-5.5309, 2.26] |
| | Gμ0_9 | -2.41E-02 | [-0.1051, 0.016] | 9.44E-03 | [-0.0456, 0.1354] |
| Shrinkage weights (origination) | ωλ0_0 | 0.9768 | [0.0192, 1] | 0.8494 | [0.0163, 1] |
| | ωλ0_1 | 0.2667 | [1.3124E-8, 0.987] | 0.6506 | [7.552E-3, 1] |
| | ωλ0_2 | 0.1941 | [7.7495E-9, 0.9288] | 0.9179 | [0.0375, 1] |
| | ωλ0_3 | 0.2333 | [5.6081E-9, 0.9705] | 0.8723 | [0.0178, 1] |
| | ωλ0_4 | 0.2437 | [2.1096E-8, 0.97] | 0.556 | [6.7253E-8, 0.9849] |
| | ωλ0_5 | 0.2261 | [8.4894E-10, 0.953] | 0.4388 | [2.5544E-7, 0.9697] |
| | ωλ0_6 | 0.1576 | [2.7303E-8, 0.9112] | 0.7979 | [0.04, 1] |
| | ωλ0_7 | 0.2461 | [3.5322E-11, 0.9711] | 0.5816 | [3.0684E-8, 0.9916] |
| | ωλ0_8 | 0.1692 | [4.1184E-9, 0.9071] | 0.5465 | [7.1687E-8, 0.9762] |
| | ωλ0_9 | 0.1481 | [5.5021E-9, 0.9124] | 0.693 | [7.9918E-3, 1] |
| Shrinkage weights (extinction) | ωμ0_0 | 0.2715 | [1.1628E-9, 0.9786] | 0.632 | [2.4179E-8, 0.9903] |
| | ωμ0_1 | 0.2558 | [9.7301E-10, 0.9876] | 0.6528 | [8.616E-3, 1] |
| | ωμ0_2 | 0.167 | [4.7062E-9, 0.9144] | 0.5352 | [3.9832E-8, 0.9859] |
| | ωμ0_3 | 0.2346 | [4.8949E-9, 0.9688] | 0.6113 | [7.9136E-9, 0.9913] |
| | ωμ0_4 | 0.2217 | [2.1232E-9, 0.9675] | 0.7353 | [0.0126, 1] |
| | ωμ0_5 | 0.3112 | [4.9638E-9, 0.967] | 0.7278 | [0.0158, 0.9999] |
| | ωμ0_6 | 0.2487 | [2.6655E-8, 0.9468] | 0.8542 | [0.0614, 1] |
| | ωμ0_7 | 0.2297 | [6.9094E-8, 0.9669] | 0.5658 | [3.1471E-8, 0.9887] |
| | ωμ0_8 | 0.1975 | [6.2977E-10, 0.9425] | 0.6246 | [4.3642E-8, 0.9838] |
| | ωμ0_9 | 0.5458 | [8.7253E-11, 0.9765] | 0.5147 | [2.9504E-11, 0.9817] |

**table S8.** Continued.

| Parameters | | Jurassic | | Lower Cretaceous | |
|---|---|---|---|---|---|
| | | Median | 95% HPD | Median | 95% HPD |
| Baseline rates | λ0 | 0.3539 | [0.014, 1.2305] | 0.4912 | [0.0239, 1.3886] |
| | μ0 | 0.0645 | [1.9837E-3, 0.5904] | 0.4347 | [0.0246, 1.2527] |
| Correlation parameters to origination | Gλ0_0 | -5.5984 | [-12.4392, 0.5599] | -0.3245 | [-11.838, 3.9456] |
| | Gλ0_1 | 0.00E+00 | [-0.0218, 0.018] | 1.00E-04 | [-0.0265, 0.0912] |
| | Gλ0_2 | 2.4521 | [-0.8696, 7.8903] | 0.0483 | [-34.4637, 15.7922] |
| | Gλ0_3 | 0.0457 | [-20.2185, 29.7649] | -1.4989 | [-38.5453, 15.5811] |
| | Gλ0_4 | 1.61E-03 | [-0.0935, 0.1471] | -2.26E-01 | [-0.4936, 0.0638] |
| | Gλ0_5 | -4.14E-01 | [-0.9667, 0.016] | 2.09E-01 | [-0.1577, 0.99] |
| | Gλ0_6 | -2.7629 | [-12.0327, 2.0297] | -1.0068 | [-28.5507, 5.9804] |
| | Gλ0_7 | -4.00E-04 | [-0.3329, 0.201] | 2.25E-04 | [-0.0746, 0.106] |
| | Gλ0_8 | 0.0545 | [-1.7033, 3.1791] | 0.2694 | [-1.638, 4.6602] |
| | Gλ0_9 | -5.60E-03 | [-0.1137, 0.0804] | 9.02E-03 | [-0.1522, 0.6874] |
| Correlation parameters to extinction | Gμ0_0 | -0.343 | [-7.8503, 1.9717] | -1.9316 | [-8.7556, 2.0423] |
| | Gμ0_1 | 5.88E-06 | [-0.0158, 0.0118] | 0.00E+00 | [-0.0215, 0.0141] |
| | Gμ0_2 | -1.2352 | [-6.7519, 1.0926] | 0.5583 | [-5.3367, 20.0869] |
| | Gμ0_3 | 0.6772 | [-9.3166, 20.9169] | 0.8412 | [-12.4863, 21.1039] |
| | Gμ0_4 | 3.14E-03 | [-0.0922, 0.1574] | -5.35E-02 | [-0.3306, 0.0826] |
| | Gμ0_5 | -6.40E-03 | [-0.2339, 0.1507] | -2.52E-01 | [-0.9223, 0.1267] |
| | Gμ0_6 | -0.38 | [-7.4685, 2.8268] | 0.8921 | [-3.6769, 12.8298] |
| | Gμ0_7 | 3.02E-03 | [-0.1413, 0.3751] | -5.40E-03 | [-0.1124, 0.0461] |
| | Gμ0_8 | 0.0687 | [-2.3103, 3.8872] | -0.3729 | [-4.2069, 1.3646] |
| | Gμ0_9 | -1.80E-02 | [-0.1262, 0.0322] | -9.00E-04 | [-0.2552, 0.1462] |
| Shrinkage weights (origination) | ωλ0_0 | 0.9573 | [0.0256, 1] | 0.6905 | [6.1488E-3, 1] |
| | ωλ0_1 | 0.5093 | [2.8411E-3, 1] | 0.6131 | [5.8418E-3, 1] |
| | ωλ0_2 | 0.7096 | [3.7406E-7, 0.986] | 0.548 | [5.386E-3, 1] |
| | ωλ0_3 | 0.4871 | [2.3886E-8, 0.9869] | 0.4904 | [3.2371E-8, 0.9857] |
| | ωλ0_4 | 0.3709 | [6.6965E-9, 0.9673] | 0.9545 | [0.1146, 1] |
| | ωλ0_5 | 0.8889 | [0.1075, 1] | 0.7568 | [0.0114, 1] |
| | ωλ0_6 | 0.7066 | [6.5677E-9, 0.9887] | 0.5419 | [3.9553E-7, 0.994] |
| | ωλ0_7 | 0.5046 | [1.0323E-8, 0.9937] | 0.3789 | [1.1137E-8, 0.9739] |
| | ωλ0_8 | 0.3316 | [1.6851E-9, 0.9595] | 0.4875 | [1.0661E-7, 0.9756] |
| | ωλ0_9 | 0.4766 | [1.4521E-8, 0.9708] | 0.5333 | [4.5743E-3, 1] |
| Shrinkage weights (extinction) | ωμ0_0 | 0.5421 | [6.892E-10, 0.9896] | 0.8401 | [0.017, 1] |
| | ωμ0_1 | 0.5061 | [4.5538E-10, 0.9952] | 0.5839 | [5.8101E-3, 1] |
| | ωμ0_2 | 0.5421 | [1.1646E-8, 0.9774] | 0.5406 | [1.9817E-7, 0.993] |
| | ωμ0_3 | 0.4215 | [4.5398E-10, 0.9789] | 0.4258 | [9.9216E-10, 0.9731] |
| | ωμ0_4 | 0.3513 | [1.8992E-8, 0.97] | 0.7758 | [0.0133, 0.9999] |
| | ωμ0_5 | 0.2598 | [3.9673E-10, 0.9431] | 0.7919 | [0.0194, 1] |
| | ωμ0_6 | 0.3898 | [2.5776E-9, 0.9696] | 0.5193 | [2.3054E-7, 0.9837] |
| | ωμ0_7 | 0.5187 | [2.5351E-7, 0.9952] | 0.4355 | [1.3795E-7, 0.9751] |
| | ωμ0_8 | 0.3655 | [4.9606E-10, 0.9702] | 0.5128 | [4.9731E-9, 0.977] |
| | ωμ0_9 | 0.4875 | [1.0744E-7, 0.9715] | 0.4353 | [1.5006E-8, 0.9876] |

**table S8.** Continued.

| Parameters | | Upper Cretaceous | | Cenozoic | |
|---|---|---|---|---|---|
| | | Median | 95% HPD | Median | 95% HPD |
| Baseline rates | λ0 | 0.3921 | [5.2596E-3, 1.2989] | 0.0864 | [1.2307E-3, 0.3548] |
| | μ0 | 0.365 | [5.2333E-3, 1.2097] | 0.1343 | [1.4738E-3, 0.6031] |
| Correlation parameters to origination | Gλ0_0 | -2.2849 | [-42.9143, 4.1799] | -0.3094 | [-3.5712, 0.9408] |
| | Gλ0_1 | 7.28E-05 | [-0.0111, 0.013] | 5.82E-05 | [-0.0028, 4.3149E-3] |
| | Gλ0_2 | -3.9148 | [-33.1911, 9.0135] | -0.0112 | [-11.1485, 12.5144] |
| | Gλ0_3 | 0.4883 | [-25.2586, 33.7439] | **-32.3419** | **[-54.7003, -8.438]** |
| | Gλ0_4 | 5.99E-03 | [-0.1712, 0.3638] | **1.33E-01** | **[0.0514, 0.2055]** |
| | Gλ0_5 | -1.33E-01 | [-1.0853, 0.3168] | 4.01E-03 | [-0.2863, 0.3048] |
| | Gλ0_6 | -0.481 | [-64.415, 50.476] | -2.4592 | [-21.5585, 6.1081] |
| | Gλ0_7 | 1.75E-02 | [-0.1462, 0.4533] | 4.70E-02 | [-0.0304, 0.1698] |
| | Gλ0_8 | -0.4244 | [-8.3888, 2.9158] | -0.0081 | [-1.9504, 1.7502] |
| | Gλ0_9 | -2.40E-03 | [-0.2113, 0.1714] | 4.53E-02 | [-0.0542, 0.2499] |
| Correlation parameters to extinction | Gμ0_0 | -0.0664 | [-10.5049, 8.1778] | -0.9903 | [-5.6315, 0.9774] |
| | Gμ0_1 | -2.00E-04 | [-0.032, 0.018] | -1.60E-03 | [-0.0081, 1.5797E-3] |
| | Gμ0_2 | -0.4753 | [-30.4705, 18.0104] | -0.6647 | [-19.8589, 9.0159] |
| | Gμ0_3 | -86.9427 | [-252.2411, 13.3971] | -1.9797 | [-27.5178, 7.8182] |
| | Gμ0_4 | 8.82E-01 | [-0.0943, 2.5657] | 9.92E-04 | [-0.0512, 0.0652] |
| | Gμ0_5 | -1.37E-01 | [-0.9646, 0.2418] | -1.35E-02 | [-0.4581, 0.3804] |
| | Gμ0_6 | -3.0299 | [-160.8936, 45.9853] | 0.2807 | [-8.6148, 13.0246] |
| | Gμ0_7 | 1.08E-04 | [-0.3331, 0.3284] | -2.70E-03 | [-0.1119, 0.0865] |
| | Gμ0_8 | 0.011 | [-3.7062, 4.5102] | -0.3252 | [-4.7834, 1.6281] |
| | Gμ0_9 | 1.88E-04 | [-0.2839, 0.2512] | -2.25E-02 | [-0.257, 0.0628] |
| Shrinkage weights (origination) | ωλ0_0 | 0.9253 | [0.0166, 1] | 0.3607 | [1.4871E-7, 0.96] |
| | ωλ0_1 | 0.6736 | [7.9203E-3, 1] | 0.2583 | [1.7292E-8, 0.9476] |
| | ωλ0_2 | 0.8038 | [0.0188, 1] | 0.2512 | [3.7965E-9, 0.9204] |
| | ωλ0_3 | 0.5741 | [2.1335E-7, 0.987] | **0.9432** | **[0.6378, 1]** |
| | ωλ0_4 | 0.6397 | [1.8545E-7, 0.9925] | **0.8884** | **[0.5602, 1]** |
| | ωλ0_5 | 0.7062 | [9.5963E-3, 0.9997] | 0.2055 | [1.0748E-8, 0.902] |
| | ωλ0_6 | 0.6616 | [8.6791E-3, 1] | 0.3975 | [2.9183E-9, 0.9576] |
| | ωλ0_7 | 0.6489 | [8.7683E-3, 1] | 0.4333 | [2.7882E-8, 0.9556] |
| | ωλ0_8 | 0.7182 | [4.6229E-9, 0.9915] | 0.2162 | [2.525E-8, 0.9082] |
| | ωλ0_9 | 0.6418 | [1.8429E-8, 0.9894] | 0.5658 | [1.6374E-10, 0.9772] |
| Shrinkage weights (extinction) | ωμ0_0 | 0.6891 | [9.9116E-3, 1] | 0.6471 | [2.6806E-8, 0.9854] |
| | ωμ0_1 | 0.7933 | [0.0107, 1] | 0.6285 | [8.7187E-8, 0.982] |
| | ωμ0_2 | 0.6867 | [8.291E-3, 1] | 0.3069 | [8.0198E-9, 0.9451] |
| | ωμ0_3 | 0.9914 | [0.0903, 1] | 0.3957 | [1.3795E-8, 0.9646] |
| | ωμ0_4 | 0.9964 | [0.0412, 1] | 0.2057 | [8.7058E-8, 0.903] |
| | ωμ0_5 | 0.679 | [7.2358E-11, 0.9878] | 0.266 | [2.7175E-9, 0.9302] |
| | ωμ0_6 | 0.7809 | [9.2917E-3, 1] | 0.2296 | [2.9882E-8, 0.9227] |
| | ωμ0_7 | 0.6175 | [2.1302E-8, 0.9904] | 0.2163 | [6.1629E-10, 0.9064] |
| | ωμ0_8 | 0.5693 | [1.8857E-9, 0.9847] | 0.379 | [6.5685E-7, 0.9594] |
| | ωμ0_9 | 0.6675 | [8.1468E-3, 1] | 0.4358 | [8.4812E-11, 0.9686] |

**table S9.**

Posterior parameter estimates for the MBD model applied to Coleoptera genera, considering singletons and excluding amber occurrences, across multiple temporal windows. The MBD model estimates the baseline origination and extinction rates (λ0 and μ0), the correlation parameters (Gλ and Gμ) for each variable, and the shrinkage weights (ω) of the correlation parameters. A variable was considered to have a significant effect (positive or negative depending on the sign of Gλ or Gμ) when its shrinkage weight exceeded 0.5 and when the 95% HPD interval of the corresponding correlation parameter did not overlap with zero (values highlighted in bold). The drivers are numbered as follows: (0) diversity of Coleoptera genera through time, (1) angiosperms diversity through time, (2) global variation of atmospheric $CO_2$ through time, (3) continental fragmentation through time, (4) gymnosperms diversity through time, (5) global variation in $\delta^{34}S$ through time (used here as an inverted proxy for global magmatic activity), (6) global variation of atmospheric $O_2$ through time, (7) Pteridophytes diversity through time, (8) Sea level fluctuations through time, and (9) variation of the global mean temperature through time. "All" corresponds to the time window encompassing the entire evolutionary history of Coleoptera genera, around 300 Ma to the present. "Before Upper Cretaceous" spans 300–100.5 Ma. The other time intervals are defined as follows: Permian (298.9–251.902 Ma), Triassic (251.902–201.4 Ma), Jurassic (201.4–143.1 Ma), Lower Cretaceous (143.1–100.5 Ma), Upper Cretaceous (100.5–66 Ma), and Cenozoic (66 Ma to the present).

| Parameters | | All | | Before Upper Cretaceous | |
|---|---|---|---|---|---|
| | | Median | 95% HPD | Median | 95% HPD |
| Baseline rates | λ0 | 4.40E-03 | [5.9763E-4, 0.0157] | 5.46E-01 | [0.0851, 1.3269] |
| | μ0 | 0.0103 | [1.4971E-3, 0.0282] | 0.0123 | [3.5686E-4, 0.0658] |
| Correlation parameters to origination | Gλ0_0 | **-2.0439** | **[-2.9154, -1.2134]** | **4.7324** | **[0.8475, 8.5792]** |
| | Gλ0_1 | **4.77E-03** | **[3.6859E-3, 5.9678E-3]** | -1.89E-02 | [-0.0393, 1.5834E-3] |
| | Gλ0_2 | 1.0049 | [-0.0339, 1.9333] | -0.0796 | [-1.1744, 0.7706] |
| | Gλ0_3 | **-11.1884** | **[-13.2884, -9.2458]** | -0.4602 | [-5.7664, 3.4157] |
| | Gλ0_4 | **0.0392** | **[0.0271, 0.0533]** | **-0.0501** | **[-0.081, -0.0259]** |
| | Gλ0_5 | **-0.1357** | **[-0.1978, -0.0784]** | -0.0537 | [-0.1058, 1.3727E-3] |
| | Gλ0_6 | **1.0363** | **[0.2163, 1.971]** | **-1.7333** | **[-3.1527, -0.5191]** |
| | Gλ0_7 | **0.0353** | **[0.0278, 0.0428]** | 0.0195 | [-0.0001, 0.0362] |
| | Gλ0_8 | -0.1673 | [-0.7151, 0.193] | 0.2762 | [-0.6092, 1.8098] |
| | Gλ0_9 | **0.0477** | **[9.9913E-3, 0.0803]** | 0.0211 | [-0.0044, 0.0526] |
| Correlation parameters to extinction | Gμ0_0 | **1.5294** | **[0.4852, 2.5811]** | **8.749** | **[4.1674, 12.0787]** |
| | Gμ0_1 | **-0.003** | **[-0.0045, -0.0014]** | 0.0213 | [-0.0008, 0.0403] |
| | Gμ0_2 | -0.2104 | [-1.2826, 0.5355] | **-2.0271** | **[-3.3735, -0.6525]** |
| | Gμ0_3 | -2.1908 | [-4.7265, 0.1903] | 1.4772 | [-1.5681, 5.7478] |
| | Gμ0_4 | 0.0148 | [-0.0001, 0.03] | 0.0105 | [-0.0102, 0.0436] |
| | Gμ0_5 | **-0.0739** | **[-0.1174, -0.0258]** | 8.01E-04 | [-0.033, 0.0417] |
| | Gμ0_6 | 0.0136 | [-0.6738, 0.7335] | -0.1223 | [-1.3995, 0.7358] |
| | Gμ0_7 | **0.019** | **[8.8564E-3, 0.0295]** | -0.0039 | [-0.0239, 0.01] |
| | Gμ0_8 | -0.15 | [-0.8368, 0.3418] | -1.0495 | [-2.3566, 0.0809] |
| | Gμ0_9 | **0.058** | **[0.0308, 0.0841]** | **0.0743** | **[0.0287, 0.1225]** |
| Shrinkage weights (origination) | ωλ0_0 | **0.8423** | **[0.4698, 1]** | **0.95** | **[0.5957, 1]** |
| | ωλ0_1 | **0.931** | **[0.7283, 1]** | 0.9934 | [0.2406, 1] |
| | ωλ0_2 | 0.4766 | [1.176E-6, 0.9547] | 0.1966 | [9.3897E-9, 0.9245] |
| | ωλ0_3 | **0.8781** | **[0.5923, 0.9999]** | 0.3471 | [2.3738E-8, 0.9525] |
| | ωλ0_4 | **0.8803** | **[0.5787, 1]** | **0.9177** | **[0.6394, 1]** |
| | ωλ0_5 | **0.8934** | **[0.6016, 1]** | 0.6859 | [0.0908, 0.9999] |
| | ωλ0_6 | **0.6802** | **[0.15, 1]** | **0.8001** | **[0.2969, 1]** |
| | ωλ0_7 | **0.8409** | **[0.5025, 1]** | 0.6978 | [0.1262, 1] |
| | ωλ0_8 | 0.2326 | [8.61E-8, 0.9328] | 0.3895 | [2.9119E-8, 0.9608] |
| | ωλ0_9 | **0.8124** | **[0.2978, 1]** | 0.5917 | [4.4158E-8, 0.9734] |
| Shrinkage weights (extinction) | ωμ0_0 | **0.7715** | **[0.2918, 0.9999]** | **0.9828** | **[0.8961, 1]** |
| | ωμ0_1 | **0.8636** | **[0.5154, 0.9999]** | 0.9947 | [0.3901, 1] |
| | ωμ0_2 | 0.2343 | [1.2813E-8, 0.9239] | **0.6946** | **[0.1908, 1]** |
| | ωμ0_3 | 0.4493 | [3.8054E-7, 0.9557] | 0.377 | [7.6255E-10, 0.9551] |
| | ωμ0_4 | 0.6609 | [0.0909, 1] | 0.5981 | [9.3452E-9, 0.9774] |
| | ωμ0_5 | **0.776** | **[0.3129, 1]** | 0.2519 | [5.4999E-8, 0.9404] |
| | ωμ0_6 | 0.2422 | [9.6703E-9, 0.9324] | 0.3384 | [1.8729E-8, 0.9486] |
| | ωμ0_7 | **0.6964** | **[0.221, 0.9998]** | 0.3649 | [3.5375E-8, 0.9567] |
| | ωμ0_8 | 0.2651 | [8.0097E-11, 0.9327] | 0.6646 | [0.0535, 1] |
| | ωμ0_9 | **0.8606** | **[0.4968, 1]** | **0.9011** | **[0.5239, 1]** |

**table S9.** Continued.

| Parameters | | Permian | | Triassic | |
|---|---|---|---|---|---|
| | | Median | 95% HPD | Median | 95% HPD |
| Baseline rates | λ0 | 6.03E-01 | [0.0469, 1.5344] | 5.61E-01 | [0.0154, 1.5996] |
| | μ0 | 0.5569 | [0.0345, 1.5204] | 0.458 | [0.0125, 1.3453] |
| Correlation parameters to origination | Gλ0_0 | -67.9291 | [-113.3647, 3.1578] | -0.3082 | [-21.3892, 8.7525] |
| | Gλ0_1 | 0.00E+00 | [-0.0197, 0.0234] | -1.00E-04 | [-0.4844, 0.1215] |
| | Gλ0_2 | 0.658 | [-2.3253, 6.1588] | 7.0421 | [-0.4073, 14.4159] |
| | Gλ0_3 | 0.3559 | [-27.8836, 63.0201] | 36.264 | [-8.0737, 92.1906] |
| | Gλ0_4 | -0.0002 | [-0.1153, 0.0982] | 0.0203 | [-0.0431, 0.1622] |
| | Gλ0_5 | 0.0248 | [-0.0592, 0.1799] | -0.0012 | [-0.2014, 0.2071] |
| | Gλ0_6 | 0.1881 | [-1.6285, 6.5213] | **-11.6018** | **[-16.0001, -7.157]** |
| | Gλ0_7 | 0 | [-0.1147, 0.1048] | -0.1168 | [-0.4025, 0.0365] |
| | Gλ0_8 | 0.0909 | [-2.3577, 3.4349] | -0.0241 | [-3.0848, 3.2153] |
| | Gλ0_9 | -0.0113 | [-0.2006, 0.0293] | -0.0005 | [-0.1068, 0.106] |
| Correlation parameters to extinction | Gμ0_0 | -0.0271 | [-21.2445, 17.3715] | 2.1887 | [-4.7786, 48.1117] |
| | Gμ0_1 | 1.17E-06 | [-0.0246, 0.0172] | 2.71E-05 | [-0.0416, 0.0457] |
| | Gμ0_2 | 0.2863 | [-3.5057, 5.8661] | -1.3591 | [-19.424, 3.9132] |
| | Gμ0_3 | 3.0297 | [-17.9714, 76.9405] | 2.553 | [-28.6382, 67.7093] |
| | Gμ0_4 | 1.49E-03 | [-0.1046, 0.1647] | 9.18E-04 | [-0.0822, 0.1117] |
| | Gμ0_5 | 4.71E-02 | [-0.0528, 0.2301] | 7.32E-02 | [-0.0783, 0.2611] |
| | Gμ0_6 | 3.3938 | [-1.0001, 10.7315] | **-8.2089** | **[-14.753, -3.0042]** |
| | Gμ0_7 | -0.0009 | [-0.2253, 0.1551] | -0.0389 | [-0.4564, 0.0637] |
| | Gμ0_8 | 0.023 | [-2.7681, 3.1212] | -0.0238 | [-3.1002, 2.7973] |
| | Gμ0_9 | -0.1465 | [-0.3126, 3.0275E-3] | 0.0501 | [-0.0366, 0.3302] |
| Shrinkage weights (origination) | ωλ0_0 | 0.9996 | [0.3721, 1] | 0.8517 | [0.0244, 1] |
| | ωλ0_1 | 0.6841 | [8.8878E-3, 1] | 0.8859 | [0.0256, 1] |
| | ωλ0_2 | 0.5396 | [8.2197E-9, 0.9794] | 0.9466 | [0.4382, 1] |
| | ωλ0_3 | 0.6909 | [9.4896E-3, 1] | 0.9826 | [0.1772, 1] |
| | ωλ0_4 | 0.6169 | [4.439E-9, 0.9912] | 0.8254 | [0.0245, 1] |
| | ωλ0_5 | 0.6108 | [1.6361E-8, 0.9833] | 0.647 | [4.8588E-8, 0.9866] |
| | ωλ0_6 | 0.5162 | [4.2675E-9, 0.987] | **0.9762** | **[0.8829, 1]** |
| | ωλ0_7 | 0.6157 | [8.4724E-10, 0.9905] | 0.9685 | [0.0879, 1] |
| | ωλ0_8 | 0.5175 | [8.5023E-10, 0.9776] | 0.6502 | [6.6519E-9, 0.9849] |
| | ωλ0_9 | 0.5998 | [2.7764E-9, 0.9902] | 0.6859 | [0.0127, 0.9997] |
| Shrinkage weights (extinction) | ωμ0_0 | 0.7125 | [0.0102, 1] | 0.9472 | [0.0368, 1] |
| | ωμ0_1 | 0.6747 | [7.2872E-3, 1] | 0.8596 | [0.0264, 1] |
| | ωμ0_2 | 0.5172 | [1.5716E-7, 0.9772] | 0.8239 | [0.0157, 1] |
| | ωμ0_3 | 0.8156 | [0.013, 1] | 0.8825 | [0.0317, 1] |
| | ωμ0_4 | 0.6661 | [7.831E-3, 1] | 0.7273 | [0.0143, 1] |
| | ωμ0_5 | 0.7118 | [0.0139, 0.9998] | 0.7934 | [0.0396, 0.9999] |
| | ωμ0_6 | 0.9175 | [0.03, 1] | **0.9579** | **[0.7301, 1]** |
| | ωμ0_7 | 0.7075 | [8.7424E-3, 1] | 0.914 | [0.0359, 1] |
| | ωμ0_8 | 0.5205 | [2.5618E-7, 0.9777] | 0.6283 | [5.8454E-10, 0.9857] |
| | ωμ0_9 | 0.9667 | [0.4738, 1] | 0.8716 | [0.028, 1] |

**table S9.** Continued.

| Parameters | | Jurassic | | Lower Cretaceous | |
|---|---|---|---|---|---|
| | | Median | 95% HPD | Median | 95% HPD |
| Baseline rates | λ0 | 2.12E-03 | [2.3044E-8, 0.1236] | 2.37E-01 | [3.2764E-4, 1.0346] |
| | μ0 | 0.0157 | [8.4841E-7, 0.1994] | 0.677 | [0.0755, 1.8487] |
| Correlation parameters to origination | Gλ0_0 | -0.0983 | [-8.8795, 7.8858] | 7.2248 | [-1.5093, 21.1432] |
| | Gλ0_1 | 1.53E-05 | [-0.0548, 0.0486] | -1.00E-01 | [-0.2494, 7.3809E-3] |
| | Gλ0_2 | 0.0841 | [-8.417, 14.4123] | -20.5756 | [-55.3232, 1.7352] |
| | Gλ0_3 | 38.1171 | [-1.4835, 68.311] | -3.0707 | [-24.4926, 24.4108] |
| | Gλ0_4 | -0.0037 | [-0.3808, 0.2385] | 0.0407 | [-0.2033, 0.3875] |
| | Gλ0_5 | -0.3233 | [-0.7151, 0.0258] | 0.3199 | [-0.2499, 1.4616] |
| | Gλ0_6 | **-12.519** | **[-17.8037, -8.259]** | 2.0048 | [-15.273, 40.7814] |
| | Gλ0_7 | **-0.8956** | **[-1.4929, -0.1144]** | -0.051 | [-0.2776, 0.0876] |
| | Gλ0_8 | 1.3731 | [-0.855, 5.1284] | 0.9422 | [-0.9715, 3.8695] |
| | Gλ0_9 | **0.2896** | **[0.17, 0.4477]** | 0.2342 | [-0.1829, 0.9205] |
| Correlation parameters to extinction | Gμ0_0 | **11.4598** | **[3.9558, 19.6938]** | 0.2977 | [-3.6063, 5.861] |
| | Gμ0_1 | 0.00E+00 | [-0.0327, 0.0372] | -1.50E-03 | [-0.0796, 0.0135] |
| | Gμ0_2 | -4.7972 | [-11.613, 0.8198] | 1.2677 | [-6.1805, 21.7073] |
| | Gμ0_3 | 8.8733 | [-7.0684, 39.2067] | **49.4971** | **[27.7763, 70.8638]** |
| | Gμ0_4 | 1.87E-03 | [-0.128, 0.2422] | 2.62E-03 | [-0.1489, 0.1643] |
| | Gμ0_5 | 2.91E-02 | [-0.1253, 0.3471] | **-1.46E+00** | **[-1.9518, -0.9577]** |
| | Gμ0_6 | 0.3962 | [-3.1442, 6.5637] | -12.7393 | [-24.9504, 2.2811] |
| | Gμ0_7 | -0.1234 | [-0.8236, 0.0876] | -0.0073 | [-0.096, 0.0478] |
| | Gμ0_8 | 0.0408 | [-2.1053, 3.2765] | 0.4119 | [-1.075, 2.3621] |
| | Gμ0_9 | 0.1038 | [-0.0045, 0.2614] | -0.095 | [-0.4626, 0.0561] |
| Shrinkage weights (origination) | ωλ0_0 | 0.8851 | [0.0449, 1] | 0.9813 | [0.2373, 1] |
| | ωλ0_1 | 0.9243 | [0.056, 1] | 0.9997 | [0.1717, 1] |
| | ωλ0_2 | 0.8577 | [0.0407, 1] | 0.9896 | [0.4919, 1] |
| | ωλ0_3 | 0.9856 | [0.648, 1] | 0.8046 | [0.0336, 1] |
| | ωλ0_4 | 0.8945 | [0.0596, 1] | 0.895 | [0.0434, 1] |
| | ωλ0_5 | 0.9075 | [0.234, 1] | 0.9229 | [0.0388, 1] |
| | ωλ0_6 | **0.9737** | **[0.8712, 1]** | 0.9285 | [0.0384, 1] |
| | ωλ0_7 | **0.9987** | **[0.9784, 1]** | 0.9129 | [0.0471, 1] |
| | ωλ0_8 | 0.8362 | [0.0563, 1] | 0.7804 | [0.0289, 1] |
| | ωλ0_9 | **0.9835** | **[0.9174, 1]** | 0.9805 | [0.0805, 1] |
| Shrinkage weights (extinction) | ωμ0_0 | **0.9913** | **[0.9415, 1]** | 0.8109 | [0.0244, 1] |
| | ωμ0_1 | 0.9168 | [0.0538, 1] | 0.9548 | [0.0569, 1] |
| | ωμ0_2 | 0.9238 | [0.2481, 1] | 0.8562 | [0.0274, 1] |
| | ωμ0_3 | 0.9072 | [0.0596, 1] | **0.9827** | **[0.9098, 1]** |
| | ωμ0_4 | 0.7712 | [0.0223, 0.9999] | 0.7387 | [0.0168, 1] |
| | ωμ0_5 | 0.6246 | [2.9276E-8, 0.9883] | **0.9895** | **[0.9478, 1]** |
| | ωμ0_6 | 0.6674 | [0.015, 0.9998] | 0.9666 | [0.3225, 1] |
| | ωμ0_7 | 0.9713 | [0.1004, 1] | 0.7141 | [0.0156, 1] |
| | ωμ0_8 | 0.6754 | [0.0127, 0.9999] | 0.6501 | [0.0127, 1] |
| | ωμ0_9 | 0.9304 | [0.3612, 1] | 0.9287 | [0.0745, 1] |

**table S9.** Continued.

| Parameters | | Upper Cretaceous | | Cenozoic | |
|---|---|---|---|---|---|
| | | Median | 95% HPD | Median | 95% HPD |
| Baseline rates | λ0 | 3.75E-01 | [0.0163, 1.1805] | 3.58E-05 | [2.7483E-8, 3.6033E-4] |
| | μ0 | 0.3188 | [6.0073E-3, 1.0633] | 2.14E-03 | [7.9235E-7, 0.0274] |
| Correlation parameters to origination | Gλ0_0 | -0.0093 | [-6.3305, 4.5706] | **6.5265** | **[4.2018, 8.6043]** |
| | Gλ0_1 | 1.75E-05 | [-0.004, 5.6668E-3] | 4.20E-03 | [-0.0008, 0.0119] |
| | Gλ0_2 | -0.0293 | [-6.142, 5.5885] | **39.0952** | **[22.1275, 66.1798]** |
| | Gλ0_3 | -1.0862 | [-18.7955, 9.149] | **-31.1468** | **[-45.0306, -18.825]** |
| | Gλ0_4 | -0.0143 | [-0.2099, 0.0596] | 0.0835 | [-0.0009, 0.1273] |
| | Gλ0_5 | -0.3063 | [-0.5733, 5.4053E-3] | 0.0156 | [-0.2703, 0.4714] |
| | Gλ0_6 | -0.0594 | [-19.5208, 20.8678] | 7.068 | [-4.4968, 31.9162] |
| | Gλ0_7 | 3.38E-03 | [-0.0884, 0.1611] | **1.90E-01** | **[0.1254, 0.2645]** |
| | Gλ0_8 | -0.1122 | [-2.7824, 1.5406] | **-1.8439** | **[-2.8522, -0.7932]** |
| | Gλ0_9 | -0.0112 | [-0.1499, 0.0426] | **0.163** | **[0.0749, 0.2587]** |
| Correlation parameters to extinction | Gμ0_0 | -0.1041 | [-22.0394, 6.1209] | 4.2282 | [-0.356, 12.2287] |
| | Gμ0_1 | -2.00E-04 | [-0.0112, 4.494E-3] | -3.00E-03 | [-0.015, 2.3409E-3] |
| | Gμ0_2 | 0.2363 | [-5.9304, 10.7382] | 0.1216 | [-21.694, 19.545] |
| | Gμ0_3 | -0.5125 | [-40.3621, 17.8889] | 12.8939 | [-1.3377, 31.9302] |
| | Gμ0_4 | 1.84E-01 | [-0.05, 0.6334] | 8.11E-02 | [-0.001, 0.1739] |
| | Gμ0_5 | -1.00E-04 | [-0.2129, 0.2427] | -1.65E-01 | [-0.5622, 0.1207] |
| | Gμ0_6 | -1.5943 | [-76.457, 18.6755] | -15.291 | [-43.6591, 2.2978] |
| | Gμ0_7 | -0.2544 | [-0.5299, 0.0192] | 0.0673 | [-0.0442, 0.1742] |
| | Gμ0_8 | 3.75E-03 | [-2.0883, 2.001] | 1.29E+00 | [-0.4428, 3.9245] |
| | Gμ0_9 | 0.0102 | [-0.0439, 0.1454] | -0.0381 | [-0.2017, 0.0523] |
| Shrinkage weights (origination) | ωλ0_0 | 0.389 | [1.1954E-7, 0.985] | **0.9736** | **[0.8812, 1]** |
| | ωλ0_1 | 0.3454 | [2.2234E-7, 0.9719] | 0.8998 | [0.1299, 1] |
| | ωλ0_2 | 0.2386 | [1.3765E-9, 0.9309] | **0.9711** | **[0.8509, 1]** |
| | ωλ0_3 | 0.3132 | [3.6883E-9, 0.9537] | **0.9553** | **[0.7922, 1]** |
| | ωλ0_4 | 0.4518 | [2.7178E-9, 0.9783] | 0.8502 | [0.2732, 1] |
| | ωλ0_5 | 0.7362 | [0.1104, 0.9999] | 0.5291 | [3.4894E-9, 0.9773] |
| | ωλ0_6 | 0.3229 | [2.3919E-11, 0.9673] | 0.7613 | [0.0273, 0.9999] |
| | ωλ0_7 | 0.2851 | [4.7154E-8, 0.9527] | **0.8987** | **[0.6016, 1]** |
| | ωλ0_8 | 0.3174 | [5.0618E-8, 0.9457] | **0.8015** | **[0.3091, 0.9999]** |
| | ωλ0_9 | 0.4253 | [7.5129E-8, 0.967] | **0.9042** | **[0.5856, 1]** |
| Shrinkage weights (extinction) | ωμ0_0 | 0.5005 | [3.1111E-3, 1] | 0.9518 | [0.237, 1] |
| | ωμ0_1 | 0.4486 | [1.8646E-7, 0.9883] | 0.8792 | [0.0311, 1] |
| | ωμ0_2 | 0.3159 | [1.0658E-7, 0.9514] | 0.6501 | [2.5407E-10, 0.9853] |
| | ωμ0_3 | 0.4092 | [2.3442E-9, 0.9807] | 0.8539 | [0.0968, 1] |
| | ωμ0_4 | 0.9332 | [0.0144, 1] | 0.86 | [0.2214, 1] |
| | ωμ0_5 | 0.2055 | [1.4E-7, 0.918] | 0.6638 | [0.0183, 1] |
| | ωμ0_6 | 0.5 | [6.6827E-8, 0.9939] | 0.8905 | [0.0915, 1] |
| | ωμ0_7 | 0.935 | [0.1124, 1] | 0.6983 | [0.0354, 1] |
| | ωμ0_8 | 0.269 | [2.1554E-8, 0.9354] | 0.7281 | [0.0295, 1] |
| | ωμ0_9 | 0.3942 | [7.898E-9, 0.9665] | 0.6658 | [2.1056E-7, 0.9854] |

**table S10.**

Posterior parameter estimates for the MBD model applied to Coleoptera genera, excluding singletons and amber occurrences, across multiple temporal windows. The MBD model estimates the baseline origination and extinction rates ($\lambda 0$ and $\mu 0$), the correlation parameters ($G\lambda$ and $G\mu$) for each variable, and the shrinkage weights ($\omega$) of the correlation parameters. A variable was considered to have a significant effect (positive or negative depending on the sign of $G\lambda$ or $G\mu$) when its shrinkage weight exceeded 0.5 and when the 95% HPD interval of the corresponding correlation parameter did not overlap with zero (values highlighted in bold). The drivers are numbered as follows: (0) diversity of Coleoptera genera through time, (1) angiosperms diversity through time, (2) global variation of atmospheric $CO_2$ through time, (3) continental fragmentation through time, (4) gymnosperms diversity through time, (5) global variation in $\delta^{34}S$ through time (used here as an inverted proxy for global magmatic activity), (6) global variation of atmospheric $O_2$ through time, (7) Pteridophytes diversity through time, (8) Sea level fluctuations through time, and (9) variation of the global mean temperature through time. "All" corresponds to the time window encompassing the entire evolutionary history of Coleoptera genera, around 300 Ma to the present. "Before Upper Cretaceous" spans 300–100.5 Ma. The other time intervals are defined as follows: Permian (298.9–251.902 Ma), Triassic (251.902–201.4 Ma), Jurassic (201.4–143.1 Ma), Lower Cretaceous (143.1–100.5 Ma), Upper Cretaceous (100.5–66 Ma), and Cenozoic (66 Ma to the present).

| Parameters | | All | | Before Upper Cretaceous | |
|---|---|---|---|---|---|
| | | Median | 95% HPD | Median | 95% HPD |
| Baseline rates | λ0 | 1.97E-03 | [1.3171E-4, 6.4831E-3] | 5.66E-01 | [0.0588, 1.4404] |
| | μ0 | 1.68E-03 | [2.4662E-5, 0.0129] | 2.52E-04 | [4.1283E-7, 5.9749E-3] |
| Correlation parameters to origination | Gλ0_0 | **-5.249** | **[-6.0878, -4.3902]** | **-11.4895** | **[-15.9557, -6.0664]** |
| | Gλ0_1 | **7.49E-03** | **[6.3949E-3, 8.5427E-3]** | **-1.13E-01** | **[-0.1583, -0.0767]** |
| | Gλ0_2 | **4.3463** | **[2.8648, 5.8367]** | **4.7028** | **[2.8289, 6.7431]** |
| | Gλ0_3 | **-11.7564** | **[-13.9424, -9.6792]** | **10.6** | **[4.5311, 16.504]** |
| | Gλ0_4 | **0.0281** | **[0.0137, 0.0432]** | **-0.0622** | **[-0.0886, -0.0298]** |
| | Gλ0_5 | -0.0317 | [-0.0899, 0.0109] | -0.0361 | [-0.0875, 8.5382E-3] |
| | Gλ0_6 | **3.6819** | **[2.6949, 4.7296]** | 0.1464 | [-1.1313, 2.2115] |
| | Gλ0_7 | **0.0344** | **[0.0235, 0.0456]** | **0.0336** | **[6.2089E-3, 0.058]** |
| | Gλ0_8 | -0.5919 | [-1.2611, 0.0304] | -0.0239 | [-1.8516, 1.3119] |
| | Gλ0_9 | -0.012 | [-0.046, 0.015] | **-0.0534** | **[-0.0979, -0.0103]** |
| Correlation parameters to extinction | Gμ0_0 | -0.098 | [-1.7106, 1.1923] | -0.6037 | [-4.5776, 1.5811] |
| | Gμ0_1 | -0.0023 | [-0.0046, 8.7162E-5] | 1.63E-04 | [-0.0102, 0.016] |
| | Gμ0_2 | **3.1914** | **[0.5652, 5.6255]** | **0.8439** | **[-0.6825, 3.0899]** |
| | Gμ0_3 | 0.6806 | [-3.086, 5.7136] | 11.2604 | [5.0064, 17.7353] |
| | Gμ0_4 | 8.73E-04 | [-0.0257, 0.031] | 2.02E-02 | [-0.0222, 0.1033] |
| | Gμ0_5 | 0.012 | [-0.0256, 0.0616] | 0.054 | [-0.0031, 0.108] |
| | Gμ0_6 | **2.2252** | **[0.2973, 3.8793]** | 0.6838 | [-1.1622, 4.1019] |
| | Gμ0_7 | 0.0209 | [-0.0008, 0.0407] | -0.0044 | [-0.0288, 0.0129] |
| | Gμ0_8 | -0.5055 | [-2.4416, 0.6414] | -0.3728 | [-3.9634, 1.2965] |
| | Gμ0_9 | 0.027 | [-0.0082, 0.0684] | **0.1034** | **[0.0336, 0.1717]** |
| Shrinkage weights (origination) | ωλ0_0 | **0.9603** | **[0.8379, 1]** | **0.9899** | **[0.9484, 1]** |
| | ωλ0_1 | **0.9679** | **[0.8671, 1]** | **0.9998** | **[0.9991, 1]** |
| | ωλ0_2 | **0.8769** | **[0.566, 0.9999]** | **0.9012** | **[0.5983, 1]** |
| | ωλ0_3 | **0.8928** | **[0.6202, 1]** | **0.8859** | **[0.5287, 1]** |
| | ωλ0_4 | **0.8286** | **[0.4118, 1]** | **0.9436** | **[0.7417, 1]** |
| | ωλ0_5 | 0.5628 | [1.4708E-7, 0.9703] | 0.6341 | [0.0328, 0.9999] |
| | ωλ0_6 | **0.9313** | **[0.7295, 1]** | 0.5101 | [3.7176E-8, 0.9731] |
| | ωλ0_7 | **0.8435** | **[0.4815, 1]** | **0.853** | **[0.3652, 1]** |
| | ωλ0_8 | 0.509 | [3.358E-7, 0.9615] | 0.4808 | [8.2967E-9, 0.9725] |
| | ωλ0_9 | 0.5144 | [2.6883E-9, 0.9654] | **0.8677** | **[0.3905, 1]** |
| Shrinkage weights (extinction) | ωμ0_0 | 0.4358 | [1.2833E-7, 0.9619] | 0.7137 | [0.0183, 1] |
| | ωμ0_1 | 0.8295 | [0.2294, 1] | 0.8092 | [0.0227, 1] |
| | ωμ0_2 | **0.8136** | **[0.2815, 1]** | **0.5421** | **[1.1203E-7, 0.974]** |
| | ωμ0_3 | 0.3718 | [1.5306E-7, 0.9536] | 0.8964 | [0.5459, 1] |
| | ωμ0_4 | 0.5432 | [6.5545E-8, 0.9704] | 0.8434 | [0.0268, 1] |
| | ωμ0_5 | 0.3924 | [8.9245E-9, 0.9567] | 0.7321 | [0.0943, 1] |
| | ωμ0_6 | **0.8526** | **[0.3078, 1]** | 0.7487 | [0.0163, 1] |
| | ωμ0_7 | 0.7214 | [0.1059, 1] | 0.4854 | [3.5888E-9, 0.9734] |
| | ωμ0_8 | 0.5602 | [7.4435E-8, 0.9733] | 0.686 | [0.0123, 1] |
| | ωμ0_9 | 0.6908 | [0.0379, 0.9999] | **0.9465** | **[0.7016, 1]** |

**table S10.** Continued.

| Parameters | | Permian | | Triassic | |
|---|---|---|---|---|---|
| | | Median | 95% HPD | Median | 95% HPD |
| Baseline rates | λ0 | 2.42E-01 | [0.0107, 0.7721] | 2.31E-01 | [6.3996E-4, 0.7639] |
| | μ0 | 1.98E-01 | [0.0112, 0.8403] | 2.33E-01 | [9.0473E-3, 0.7696] |
| Correlation parameters to origination | Gλ0_0 | -10.8813 | [-105.6254, 4.289] | -0.2738 | [-23.7892, 4.3604] |
| | Gλ0_1 | 0.00E+00 | [-0.0089, 9.3665E-3] | 1.94E-06 | [-0.0146, 0.0148] |
| | Gλ0_2 | -0.1239 | [-5.1153, 2.6337] | 1.5856 | [-2.0463, 12.8196] |
| | Gλ0_3 | 0.1697 | [-15.442, 36.0395] | -13.5158 | [-93.206, 8.2491] |
| | Gλ0_4 | -0.0003 | [-0.0952, 0.0609] | -0.0082 | [-0.0755, 0.0313] |
| | Gλ0_5 | 3.70E-03 | [-0.0664, 0.1401] | 4.04E-03 | [-0.0923, 0.1382] |
| | Gλ0_6 | -0.0108 | [-1.9245, 1.9113] | -2.3391 | [-7.0196, 0.9335] |
| | Gλ0_7 | -0.0003 | [-0.1183, 0.07] | -0.0004 | [-0.0884, 0.0855] |
| | Gλ0_8 | -0.004 | [-3.6559, 2.4043] | -0.3643 | [-3.8907, 1.5187] |
| | Gλ0_9 | -0.0002 | [-0.0543, 0.0456] | -0.0037 | [-0.123, 0.0476] |
| Correlation parameters to extinction | Gμ0_0 | -0.0107 | [-10.0311, 6.0752] | 0.0222 | [-7.4572, 10.2156] |
| | Gμ0_1 | 1.06E-06 | [-0.0112, 9.974E-3] | 1.81E-06 | [-0.0137, 0.0166] |
| | Gμ0_2 | -0.0565 | [-4.06, 2.9353] | -0.1572 | [-6.4195, 3.799] |
| | Gμ0_3 | 0.0864 | [-15.4744, 25.2516] | -0.2443 | [-29.7501, 18.3189] |
| | Gμ0_4 | -2.00E-04 | [-0.0761, 0.0578] | -5.50E-03 | [-0.1299, 0.0596] |
| | Gμ0_5 | 3.40E-03 | [-0.0869, 0.1947] | 9.68E-02 | [-0.063, 0.3432] |
| | Gμ0_6 | -0.0744 | [-3.0267, 1.6981] | -4.2721 | [-10.5168, 0.7556] |
| | Gμ0_7 | -0.0004 | [-0.0887, 0.0621] | 1.09E-05 | [-0.1044, 0.109] |
| | Gμ0_8 | -0.0012 | [-2.8492, 2.4918] | -0.4379 | [-5.4976, 2.0175] |
| | Gμ0_9 | -0.0225 | [-0.1079, 0.0157] | 5.75E-03 | [-0.0395, 0.1028] |
| Shrinkage weights (origination) | ωλ0_0 | 0.9908 | [2.7725E-3, 1] | 0.6306 | [6.897E-3, 1] |
| | ωλ0_1 | 0.2636 | [2.7424E-10, 0.9881] | 0.5282 | [2.2533E-10, 0.9953] |
| | ωλ0_2 | 0.2372 | [1.9365E-9, 0.9493] | 0.6556 | [4.8354E-11, 0.9913] |
| | ωλ0_3 | 0.2757 | [1.3449E-8, 0.9849] | 0.9244 | [0.017, 1] |
| | ωλ0_4 | 0.2656 | [2.9275E-11, 0.9793] | 0.4685 | [2.3569E-8, 0.9724] |
| | ωλ0_5 | 0.2091 | [5.1415E-10, 0.9489] | 0.3444 | [3.5794E-11, 0.9531] |
| | ωλ0_6 | 0.1579 | [2.4654E-11, 0.9226] | 0.7081 | [5.7088E-8, 0.9837] |
| | ωλ0_7 | 0.2634 | [4.6239E-9, 0.9815] | 0.4324 | [3.4519E-9, 0.9775] |
| | ωλ0_8 | 0.2059 | [3.2873E-9, 0.9572] | 0.4862 | [2.0685E-8, 0.969] |
| | ωλ0_9 | 0.173 | [2.5227E-8, 0.9348] | 0.3776 | [6.4362E-9, 0.9742] |
| Shrinkage weights (extinction) | ωμ0_0 | 0.2623 | [3.1237E-8, 0.9913] | 0.5108 | [9.6766E-9, 0.9929] |
| | ωμ0_1 | 0.2712 | [1.0569E-13, 0.9929] | 0.5388 | [4.6719E-3, 1] |
| | ωμ0_2 | 0.1759 | [1.4715E-9, 0.9285] | 0.3723 | [1.252E-8, 0.9702] |
| | ωμ0_3 | 0.237 | [1.1621E-10, 0.9775] | 0.4755 | [7.9146E-8, 0.9848] |
| | ωμ0_4 | 0.2461 | [1.7462E-9, 0.975] | 0.5882 | [5.8834E-8, 0.9843] |
| | ωμ0_5 | 0.2495 | [1.0618E-9, 0.9604] | 0.7498 | [0.0177, 1] |
| | ωμ0_6 | 0.2352 | [2.5656E-11, 0.9425] | 0.8391 | [0.0288, 0.9999] |
| | ωμ0_7 | 0.2378 | [5.4586E-10, 0.9757] | 0.4588 | [2.2825E-8, 0.9828] |
| | ωμ0_8 | 0.2041 | [6.0589E-9, 0.9524] | 0.5787 | [7.857E-10, 0.9798] |
| | ωμ0_9 | 0.5318 | [5.5614E-9, 0.9771] | 0.3941 | [9.9243E-8, 0.9684] |

**table S10.** Continued.

| Parameters | | Jurassic | | Lower Cretaceous | |
|---|---|---|---|---|---|
| | | Median | 95% HPD | Median | 95% HPD |
| Baseline rates | λ0 | 6.22E-01 | [0.0594, 1.6965] | 4.81E-01 | [0.0188, 1.456] |
| | μ0 | 4.36E-01 | [7.9667E-3, 1.293] | 4.81E-01 | [0.0208, 1.4557] |
| Correlation parameters to origination | Gλ0_0 | **-25.9743** | **[-33.137, -18.6355]** | 1.3872 | [-5.9345, 25.625] |
| | Gλ0_1 | 5.67E-06 | [-0.0645, 0.0724] | -8.00E-04 | [-0.1727, 0.0244] |
| | Gλ0_2 | -0.1124 | [-8.9523, 8.1858] | -0.2188 | [-15.4681, 12.7811] |
| | Gλ0_3 | 61.5065 | [-1.0815, 111.5307] | 5.1663 | [-17.3159, 52.2067] |
| | Gλ0_4 | 0.0795 | [-0.3642, 0.8208] | -0.1707 | [-0.5621, 0.1412] |
| | Gλ0_5 | -6.67E-01 | [-2.2202, 0.0195] | -9.50E-02 | [-0.6832, 0.4638] |
| | Gλ0_6 | **-23.7224** | **[-33.9603, -13.0921]** | -17.3298 | [-36.6452, 3.1153] |
| | Gλ0_7 | **-1.4728** | **[-2.6148, -0.3939]** | 0.1256 | [-0.0137, 0.2555] |
| | Gλ0_8 | -0.0567 | [-4.7757, 5.0478] | 0.1919 | [-2.8899, 3.5443] |
| | Gλ0_9 | 0.1791 | [-0.0017, 0.2949] | -0.0056 | [-0.3527, 0.221] |
| Correlation parameters to extinction | Gμ0_0 | -0.8809 | [-10.0508, 3.878] | -1.1445 | [-23.6645, 37.4191] |
| | Gμ0_1 | 6.16E-05 | [-0.0627, 0.0686] | -1.20E-03 | [-0.2194, 0.0199] |
| | Gμ0_2 | **-11.6971** | **[-18.6654, -5.4787]** | 3.5972 | [-10.1669, 31.284] |
| | Gμ0_3 | -0.0184 | [-26.4647, 26.0737] | 19.827 | [-18.1581, 98.014] |
| | Gμ0_4 | 1.62E-01 | [-0.1118, 0.4492] | -1.01E-01 | [-0.7373, 0.3454] |
| | Gμ0_5 | 2.55E-01 | [-0.0339, 0.6059] | -7.91E-01 | [-2.2356, 0.0844] |
| | Gμ0_6 | 5.0017 | [-1.8111, 13.73] | -0.2363 | [-36.398, 42.819] |
| | Gμ0_7 | 1.23E-01 | [-0.2144, 0.9509] | 1.39E-03 | [-0.2562, 0.1654] |
| | Gμ0_8 | 2.5954 | [-0.9318, 6.8945] | -0.2369 | [-4.7245, 1.8134] |
| | Gμ0_9 | **-1.68E-01** | **[-0.2769, -0.0576]** | -1.38E-02 | [-0.5905, 0.307] |
| Shrinkage weights (origination) | ωλ0_0 | **0.9982** | **[0.9909, 1]** | 0.9325 | [0.0529, 1] |
| | ωλ0_1 | 0.9709 | [0.1439, 1] | 0.9467 | [0.0618, 1] |
| | ωλ0_2 | 0.9047 | [0.0773, 1] | 0.8487 | [0.0314, 1] |
| | ωλ0_3 | 0.9942 | [0.8981, 1] | 0.8606 | [0.0425, 1] |
| | ωλ0_4 | 0.973 | [0.1752, 1] | 0.9527 | [0.1514, 1] |
| | ωλ0_5 | 0.9774 | [0.6958, 1] | 0.8256 | [0.0395, 1] |
| | ωλ0_6 | **0.992** | **[0.9545, 1]** | 0.9789 | [0.3623, 1] |
| | ωλ0_7 | **0.9995** | **[0.994, 1]** | 0.9507 | [0.2609, 1] |
| | ωλ0_8 | 0.8973 | [0.0698, 1] | 0.7176 | [0.0167, 0.9998] |
| | ωλ0_9 | 0.9701 | [0.5771, 1] | 0.8569 | [0.033, 1] |
| Shrinkage weights (extinction) | ωμ0_0 | 0.9252 | [0.0839, 1] | 0.9924 | [0.3324, 1] |
| | ωμ0_1 | 0.9703 | [0.1479, 1] | 0.9545 | [0.0553, 1] |
| | ωμ0_2 | **0.9831** | **[0.8917, 1]** | 0.9271 | [0.0525, 1] |
| | ωμ0_3 | 0.897 | [0.0622, 1] | 0.9478 | [0.0873, 1] |
| | ωμ0_4 | 0.963 | [0.2332, 1] | 0.9713 | [0.1797, 1] |
| | ωμ0_5 | 0.9177 | [0.1934, 1] | 0.9728 | [0.3006, 1] |
| | ωμ0_6 | 0.9359 | [0.1745, 1] | 0.9461 | [0.0539, 1] |
| | ωμ0_7 | 0.9849 | [0.2048, 1] | 0.8385 | [0.0238, 1] |
| | ωμ0_8 | 0.9361 | [0.1743, 1] | 0.7294 | [0.0147, 1] |
| | ωμ0_9 | **0.9698** | **[0.7839, 1]** | 0.9002 | [0.0447, 1] |

**table S10.** Continued.

| Parameters | | Upper Cretaceous | | Cenozoic | |
|---|---|---|---|---|---|
| | | Median | 95% HPD | Median | 95% HPD |
| Baseline rates | λ0 | 4.91E-01 | [0.0186, 1.4808] | 1.00E-04 | [2.1232E-7, 5.063E-4] |
| | μ0 | 3.02E-01 | [2.902E-3, 1.0949] | 1.20E-01 | [4.0913E-4, 0.6164] |
| Correlation parameters to origination | Gλ0_0 | 0.0191 | [-11.0552, 12.8408] | **7.3942** | **[5.3741, 9.3833]** |
| | Gλ0_1 | 3.46E-04 | [-0.0051, 0.0186] | 8.62E-04 | [-0.0021, 5.7716E-3] |
| | Gλ0_2 | -0.143 | [-13.9726, 12.5311] | **43.178** | **[25.1698, 68.6457]** |
| | Gλ0_3 | 0.0821 | [-21.4739, 23.3493] | **-24.0793** | **[-35.6746, -14.8032]** |
| | Gλ0_4 | -0.006 | [-0.2863, 0.0962] | **0.0626** | **[0.0173, 0.1007]** |
| | Gλ0_5 | 2.62E-03 | [-0.4518, 0.3916] | 8.34E-03 | [-0.317, 0.5195] |
| | Gλ0_6 | 0.6434 | [-23.374, 43.7784] | 1.5292 | [-5.9445, 14.2009] |
| | Gλ0_7 | 4.73E-03 | [-0.1266, 0.2419] | **1.83E-01** | **[0.1156, 0.2598]** |
| | Gλ0_8 | -0.2419 | [-5.7923, 1.9447] | **-1.3012** | **[-2.5703, -0.1384]** |
| | Gλ0_9 | -0.1344 | [-0.3266, 0.0397] | 0.0569 | [-0.0156, 0.1481] |
| Correlation parameters to extinction | Gμ0_0 | -0.0303 | [-22.9725, 14.3093] | -5.7341 | [-9.5833, 0.2214] |
| | Gμ0_1 | -1.00E-04 | [-0.0148, 6.5955E-3] | -2.00E-04 | [-0.0067, 3.9862E-3] |
| | Gμ0_2 | 0.0118 | [-14.3678, 11.7637] | -13.001 | [-33.6179, 3.5959] |
| | Gμ0_3 | -1.0012 | [-60.8501, 27.7469] | 1.1946 | [-9.3298, 19.1957] |
| | Gμ0_4 | 1.45E-01 | [-0.1065, 0.9685] | 1.48E-02 | [-0.0278, 0.0793] |
| | Gμ0_5 | -1.69E-02 | [-0.5021, 0.2841] | 9.27E-02 | [-0.2115, 0.5669] |
| | Gμ0_6 | -4.5868 | [-135.3297, 22.1109] | -1.1906 | [-14.8884, 7.2335] |
| | Gμ0_7 | -1.11E-01 | [-0.6409, 0.0715] | 8.73E-03 | [-0.0609, 0.1129] |
| | Gμ0_8 | 8.73E-03 | [-4.0784, 3.4635] | -1.12E-02 | [-2.1138, 2.0737] |
| | Gμ0_9 | -4.00E-04 | [-0.1238, 0.1319] | -2.89E-02 | [-0.1749, 0.0543] |
| Shrinkage weights (origination) | ωλ0_0 | 0.5099 | [8.7749E-10, 0.9957] | **0.977** | **[0.8988, 1]** |
| | ωλ0_1 | 0.5737 | [2.1962E-8, 0.9953] | 0.5612 | [2.2215E-7, 0.9752] |
| | ωλ0_2 | 0.4659 | [5.9648E-9, 0.9771] | **0.9716** | **[0.864, 1]** |
| | ωλ0_3 | 0.3538 | [9.972E-13, 0.9734] | **0.9208** | **[0.674, 1]** |
| | ωλ0_4 | 0.4351 | [1.6626E-7, 0.9851] | **0.7527** | **[0.2701, 1]** |
| | ωλ0_5 | 0.3389 | [1.3833E-11, 0.9589] | 0.412 | [3.3644E-8, 0.9567] |
| | ωλ0_6 | 0.4451 | [6.9824E-8, 0.986] | 0.4089 | [1.1739E-8, 0.9605] |
| | ωλ0_7 | 0.3694 | [6.7119E-8, 0.9748] | **0.871** | **[0.5434, 1]** |
| | ωλ0_8 | 0.4724 | [1.3023E-9, 0.981] | **0.6365** | **[0.1045, 1]** |
| | ωλ0_9 | 0.9053 | [0.0483, 1] | 0.632 | [0.0271, 1] |
| Shrinkage weights (extinction) | ωμ0_0 | 0.5241 | [3.2144E-3, 1] | 0.9611 | [0.3903, 1] |
| | ωμ0_1 | 0.4777 | [8.3808E-10, 0.9933] | 0.5166 | [1.233E-7, 0.9766] |
| | ωμ0_2 | 0.3871 | [1.937E-8, 0.9745] | 0.8111 | [0.0358, 1] |
| | ωμ0_3 | 0.4825 | [4.7411E-8, 0.9924] | 0.4545 | [4.9956E-7, 0.969] |
| | ωμ0_4 | 0.9246 | [7.64E-3, 1] | 0.429 | [3.0814E-8, 0.9584] |
| | ωμ0_5 | 0.3235 | [6.0341E-10, 0.9619] | 0.4922 | [3.1196E-7, 0.9668] |
| | ωμ0_6 | 0.7288 | [4.6545E-3, 1] | 0.3948 | [7.851E-8, 0.9612] |
| | ωμ0_7 | 0.8247 | [9.9277E-3, 1] | 0.3457 | [2.7816E-8, 0.9456] |
| | ωμ0_8 | 0.3831 | [2.7676E-8, 0.9739] | 0.3709 | [1.5352E-8, 0.9534] |
| | ωμ0_9 | 0.3516 | [9.1575E-8, 0.9724] | 0.5471 | [2.9108E-8, 0.9724] |

**table S11.**

Posterior parameter estimates for the MBD model applied to Polyphaga genera, considering singletons and excluding amber occurrences, across multiple temporal windows. The MBD model estimates the baseline origination and extinction rates ($\lambda 0$ and $\mu 0$), the correlation parameters ($G\lambda$ and $G\mu$) for each variable, and the shrinkage weights ($\omega$) of the correlation parameters. A variable was considered to have a significant effect (positive or negative depending on the sign of $G\lambda$ or $G\mu$) when its shrinkage weight exceeded 0.5 and when the 95% HPD interval of the corresponding correlation parameter did not overlap with zero (values highlighted in bold). The drivers are numbered as follows: (0) diversity of Polyphaga genera through time, (1) angiosperms diversity through time, (2) global variation of atmospheric $CO_2$ through time, (3) continental fragmentation through time, (4) gymnosperms diversity through time, (5) global variation in $\delta^{34}S$ through time (used here as an inverted proxy for global magmatic activity), (6) global variation of atmospheric $O_2$ through time, (7) Pteridophytes diversity through time, (8) Sea level fluctuations through time, and (9) variation of the global mean temperature through time. "All" corresponds to the time window encompassing the entire evolutionary history of Polyphaga genera, around 257 Ma to the present. "Before Upper Cretaceous" spans 257–100.5 Ma. The other time intervals are defined as follows: Triassic (251.902–201.4 Ma), Jurassic (201.4–143.1 Ma), Lower Cretaceous (143.1–100.5 Ma), Upper Cretaceous (100.5–66 Ma), and Cenozoic (66 Ma to the present).

| Parameters | | All | | Before Upper Cretaceous | |
|---|---|---|---|---|---|
| | | Median | 95% HPD | Median | 95% HPD |
| Baseline rates | λ0 | 4.87E-04 | [1.7783E-5, 1.4691E-3] | 4.61E-01 | [0.0313, 1.2795] |
| | μ0 | 4.91E-03 | [2.9766E-4, 0.0157] | 1.53E-01 | [8.172E-4, 0.6978] |
| Correlation parameters to origination | Gλ0_0 | **-2.7719** | **[-3.6635, -1.8941]** | -2.138 | [-6.5291, 1.0397] |
| | Gλ0_1 | **6.02E-03** | **[4.8049E-3, 7.5071E-3]** | **-6.53E-02** | **[-0.1002, -0.0321]** |
| | Gλ0_2 | **3.2849** | **[1.2722, 5.2481]** | **5.1266** | **[1.5594, 8.255]** |
| | Gλ0_3 | **-13.3294** | **[-16.5384, -10.332]** | -3.1147 | [-16.4848, 4.5295] |
| | Gλ0_4 | **7.73E-02** | **[0.0626, 0.0917]** | -4.94E-02 | [-0.0999, 6.8019E-3] |
| | Gλ0_5 | -0.0259 | [-0.0939, 0.0241] | -0.0233 | [-0.144, 0.0507] |
| | Gλ0_6 | -0.032 | [-1.6371, 1.2699] | 1.54 | [-0.7548, 4.8018] |
| | Gλ0_7 | **0.0662** | **[0.0547, 0.0775]** | **0.0578** | **[0.0203, 0.0959]** |
| | Gλ0_8 | -0.1064 | [-1.0029, 0.4893] | 0.0799 | [-1.4322, 2.1912] |
| | Gλ0_9 | 9.41E-03 | [-0.0214, 0.0586] | **-1.20E-01** | **[-0.2059, -0.0451]** |
| Correlation parameters to extinction | Gμ0_0 | 1.2105 | [-0.0888, 2.4692] | 3.7472 | [-1.0541, 10.2274] |
| | Gμ0_1 | -3.00E-04 | [-0.0022, 6.9411E-4] | 1.61E-03 | [-0.0091, 0.0467] |
| | Gμ0_2 | 0.7854 | [-0.7002, 2.9775] | -2.3795 | [-7.3844, 1.0605] |
| | Gμ0_3 | -0.1663 | [-3.8914, 2.9115] | 10.9867 | [-0.9308, 21.4195] |
| | Gμ0_4 | **0.0719** | **[0.051, 0.0933]** | **0.1031** | **[0.0246, 0.199]** |
| | Gμ0_5 | **-0.2739** | **[-0.3774, -0.1605]** | **-0.4811** | **[-0.6955, -0.2958]** |
| | Gμ0_6 | -0.3072 | [-2.186, 0.8391] | 6.68E-03 | [-2.6769, 3.4442] |
| | Gμ0_7 | 9.33E-03 | [-0.0028, 0.0248] | -5.72E-02 | [-0.1106, -0.0067] |
| | Gμ0_8 | 0.6493 | [-0.3003, 1.7482] | -0.0469 | [-1.6995, 1.4054] |
| | Gμ0_9 | 6.97E-03 | [-0.0208, 0.0477] | -4.23E-02 | [-0.1768, 0.0349] |
| Shrinkage weights (origination) | ωλ0_0 | **0.887** | **[0.6097, 1]** | 0.8893 | [0.0724, 1] |
| | ωλ0_1 | **0.9506** | **[0.8072, 1]** | **0.9995** | **[0.9967, 1]** |
| | ωλ0_2 | **0.8135** | **[0.376, 1]** | **0.9188** | **[0.5637, 1]** |
| | ωλ0_3 | **0.9056** | **[0.6636, 1]** | 0.747 | [0.0223, 1] |
| | ωλ0_4 | **0.9397** | **[0.7693, 1]** | 0.9061 | [0.2094, 1] |
| | ωλ0_5 | 0.494 | [6.5572E-9, 0.9613] | 0.6854 | [0.018, 1] |
| | ωλ0_6 | 0.3216 | [2.8498E-9, 0.9457] | 0.8006 | [0.0467, 0.9999] |
| | ωλ0_7 | **0.9105** | **[0.6804, 1]** | **0.9156** | **[0.5704, 1]** |
| | ωλ0_8 | 0.2679 | [4.8549E-8, 0.9369] | 0.5965 | [2.0382E-11, 0.9825] |
| | ωλ0_9 | 0.4642 | [5.1088E-7, 0.9635] | **0.956** | **[0.737, 1]** |
| Shrinkage weights (extinction) | ωμ0_0 | 0.6944 | [0.0919, 1] | 0.9434 | [0.1302, 1] |
| | ωμ0_1 | 0.4043 | [7.345E-10, 0.956] | 0.9345 | [0.0565, 1] |
| | ωμ0_2 | 0.4441 | [3.9436E-8, 0.9575] | 0.8121 | [0.0391, 1] |
| | ωμ0_3 | 0.2549 | [3.6575E-8, 0.9297] | 0.8999 | [0.2459, 1] |
| | ωμ0_4 | **0.9318** | **[0.7304, 1]** | **0.9691** | **[0.7735, 1]** |
| | ωμ0_5 | **0.9644** | **[0.8355, 1]** | **0.9888** | **[0.9417, 1]** |
| | ωμ0_6 | 0.4008 | [1.5085E-7, 0.9567] | 0.653 | [2.814E-8, 0.9851] |
| | ωμ0_7 | 0.4253 | [4.1155E-7, 0.9514] | 0.9155 | [0.4513, 1] |
| | ωμ0_8 | 0.521 | [7.4865E-10, 0.9667] | 0.5336 | [7.2593E-8, 0.9792] |
| | ωμ0_9 | 0.4218 | [8.5267E-8, 0.9554] | 0.8539 | [0.0483, 1] |

**table S11.** Continued.

| Parameters | | Triassic | | Jurassic | |
|---|---|---|---|---|---|
| | | Median | 95% HPD | Median | 95% HPD |
| Baseline rates | λ0 | 0.3038 | [0.0127, 1.0079] | 4.40E-01 | [0.0133, 1.3999] |
| | μ0 | 0.2832 | [2.3253E-3, 1.0045] | 4.73E-01 | [0.0293, 1.4435] |
| Correlation parameters to origination | Gλ0_0 | 0.0101 | [-8.9894, 8.623] | **-22.8466** | **[-31.8773, -13.8267]** |
| | Gλ0_1 | 0 | [-0.0143, 0.0144] | 0.00E+00 | [-0.0282, 0.0286] |
| | Gλ0_2 | -0.089 | [-6.5735, 4.6982] | **8.0903** | **[2.3285, 12.2394]** |
| | Gλ0_3 | 0.1095 | [-23.6948, 32.1599] | 1.3907 | [-23.0733, 37.648] |
| | Gλ0_4 | 4.25E-04 | [-0.046, 0.0643] | 9.91E-04 | [-0.1504, 0.1704] |
| | Gλ0_5 | -0.002 | [-0.1829, 0.1353] | **-0.6947** | **[-1.1949, -0.2386]** |
| | Gλ0_6 | -1.2145 | [-7.7754, 1.5926] | -8.3558 | [-15.7415, 0.3382] |
| | Gλ0_7 | -0.0013 | [-0.159, 0.0799] | -0.0217 | [-0.6412, 0.1654] |
| | Gλ0_8 | -0.0145 | [-3.231, 2.8188] | 6.94E-03 | [-2.2918, 2.1195] |
| | Gλ0_9 | -0.0137 | [-0.1236, 0.0334] | -8.43E-02 | [-0.1763, 4.7991E-3] |
| Correlation parameters to extinction | Gμ0_0 | 0 | [-8.5227, 10.3117] | 0.1344 | [-4.2856, 6.4519] |
| | Gμ0_1 | 6.83E-06 | [-0.0156, 0.0168] | 0.00E+00 | [-0.0369, 0.0318] |
| | Gμ0_2 | -0.479 | [-24.5129, 8.196] | -3.2839 | [-7.4963, 0.8501] |
| | Gμ0_3 | -0.0766 | [-54.5921, 50.8483] | 6.0105 | [-8.5328, 34.3214] |
| | Gμ0_4 | -0.001 | [-0.1337, 0.101] | 0.0231 | [-0.1024, 0.2328] |
| | Gμ0_5 | -0.0043 | [-1.0509, 0.3733] | -0.0051 | [-0.2888, 0.2414] |
| | Gμ0_6 | -0.6816 | [-23.3197, 6.4202] | 7.42E-01 | [-3.8087, 8.0038] |
| | Gμ0_7 | -0.001 | [-0.3171, 0.1746] | 1.29E-01 | [-0.0979, 0.6951] |
| | Gμ0_8 | -0.0261 | [-8.4692, 5.9767] | -0.0843 | [-3.0995, 2.2253] |
| | Gμ0_9 | -0.0288 | [-0.3817, 0.0822] | **-1.11E-01** | **[-0.2082, -0.0085]** |
| Shrinkage weights (origination) | ωλ0_0 | 0.411 | [3.998E-10, 0.9929] | **0.9974** | **[0.9866, 1]** |
| | ωλ0_1 | 0.4222 | [1.9648E-8, 0.9953] | 0.8465 | [0.0285, 1] |
| | ωλ0_2 | 0.3515 | [7.7131E-9, 0.9697] | **0.954** | **[0.7325, 1]** |
| | ωλ0_3 | 0.3949 | [6.5747E-10, 0.9882] | 0.8465 | [0.0327, 1] |
| | ωλ0_4 | 0.2797 | [9.976E-9, 0.9567] | 0.6615 | [2.9868E-8, 0.9863] |
| | ωλ0_5 | 0.3058 | [3.8563E-8, 0.9611] | **0.9612** | **[0.7654, 1]** |
| | ωλ0_6 | 0.5456 | [6.3331E-11, 0.9774] | 0.9407 | [0.4386, 1] |
| | ωλ0_7 | 0.3736 | [6.1824E-9, 0.9854] | 0.8897 | [0.0311, 1] |
| | ωλ0_8 | 0.2992 | [1.5501E-8, 0.965] | 0.5681 | [1.4997E-7, 0.9794] |
| | ωλ0_9 | 0.4721 | [2.4577E-8, 0.975] | 0.8817 | [0.2056, 1] |
| Shrinkage weights (extinction) | ωμ0_0 | 0.4229 | [6.3478E-8, 0.9935] | 0.7537 | [0.0196, 0.9999] |
| | ωμ0_1 | 0.4164 | [3.4701E-9, 0.9963] | 0.8579 | [0.0329, 1] |
| | ωμ0_2 | 0.5796 | [3.5138E-3, 1] | 0.8459 | [0.0811, 0.9999] |
| | ωμ0_3 | 0.4279 | [2.4955E-9, 0.996] | 0.852 | [0.0383, 1] |
| | ωμ0_4 | 0.399 | [1.3381E-9, 0.9849] | 0.7939 | [0.0257, 1] |
| | ωμ0_5 | 0.4308 | [1.9559E-8, 0.9969] | 0.5528 | [1.1142E-7, 0.9774] |
| | ωμ0_6 | 0.6057 | [4.0679E-3, 1] | 0.6847 | [0.0152, 1] |
| | ωμ0_7 | 0.4441 | [1.4404E-8, 0.9955] | 0.9724 | [0.0719, 1] |
| | ωμ0_8 | 0.3907 | [2.3022E-9, 0.9886] | 0.6031 | [2.9974E-8, 0.9813] |
| | ωμ0_9 | 0.7512 | [5.3892E-3, 1] | **0.9198** | **[0.4808, 1]** |

**table S11.** Continued.

| Parameters | | Lower Cretaceous | | Upper Cretaceous | |
|---|---|---|---|---|---|
| | | Median | 95% HPD | Median | 95% HPD |
| Baseline rates | λ0 | 4.33E-01 | [0.0157, 1.3664] | 4.54E-01 | [0.0159, 1.3056] |
| | μ0 | 4.73E-01 | [0.0165, 1.4209] | 4.10E-01 | [0.0105, 1.3388] |
| Correlation parameters to origination | Gλ0_0 | -0.5986 | [-9.0962, 4.8076] | -0.0198 | [-8.4186, 5.5482] |
| | Gλ0_1 | **-1.77E-01** | **[-0.3521, -0.0422]** | 3.94E-05 | [-0.0053, 6.7782E-3] |
| | Gλ0_2 | -13.8973 | [-67.7052, 4.0561] | 0.1456 | [-7.9678, 16.6453] |
| | Gλ0_3 | 3.6166 | [-21.4534, 33.4128] | -0.6651 | [-21.0872, 12.9124] |
| | Gλ0_4 | -2.04E-02 | [-0.3074, 0.1884] | -2.39E-02 | [-0.2993, 0.063] |
| | Gλ0_5 | 0.3922 | [-0.1093, 1.3122] | -0.1162 | [-0.4205, 0.0992] |
| | Gλ0_6 | 0.9756 | [-16.9911, 25.1313] | 0.1741 | [-22.5212, 36.7228] |
| | Gλ0_7 | 0.0659 | [-0.0539, 0.2178] | 0.0165 | [-0.0793, 0.2257] |
| | Gλ0_8 | 3.07E-01 | [-1.3499, 2.7484] | -1.90E-01 | [-3.5811, 1.6156] |
| | Gλ0_9 | 1.33E-01 | [-0.1745, 1.2534] | -4.70E-02 | [-0.2525, 0.051] |
| Correlation parameters to extinction | Gμ0_0 | 0.9039 | [-5.3403, 11.664] | -0.0625 | [-15.5448, 7.3913] |
| | Gμ0_1 | -7.00E-04 | [-0.0621, 0.0185] | 0.00E+00 | [-0.0092, 7.7672E-3] |
| | Gμ0_2 | 1.959 | [-8.9478, 35.7916] | -0.0092 | [-10.2727, 10.3112] |
| | Gμ0_3 | **49.9833** | **[15.4082, 91.3823]** | 2.6848 | [-14.2196, 53.5366] |
| | Gμ0_4 | 0.0107 | [-0.1594, 0.2312] | 0.0963 | [-0.0578, 0.5727] |
| | Gμ0_5 | **-1.4244** | **[-2.1446, -0.664]** | -0.0709 | [-0.6509, 0.1473] |
| | Gμ0_6 | -8.18E+00 | [-37.0389, 6.0428] | -1.91E-01 | [-58.029, 38.269] |
| | Gμ0_7 | -8.20E-03 | [-0.1172, 0.1062] | **-4.41E-01** | **[-0.8207, -0.0505]** |
| | Gμ0_8 | 0.4379 | [-1.7062, 3.4278] | 0.1303 | [-3.424, 6.2872] |
| | Gμ0_9 | -1.42E-01 | [-0.8185, 0.0788] | 1.62E-02 | [-0.0608, 0.2437] |
| Shrinkage weights (origination) | ωλ0_0 | 0.878 | [0.0402, 1] | 0.4861 | [6.035E-11, 0.9888] |
| | ωλ0_1 | **0.9999** | **[0.9989, 1]** | 0.4365 | [1.1433E-9, 0.9813] |
| | ωλ0_2 | 0.9847 | [0.1901, 1] | 0.4102 | [1.0426E-8, 0.9747] |
| | ωλ0_3 | 0.8378 | [0.0406, 1] | 0.3731 | [3.0869E-9, 0.9674] |
| | ωλ0_4 | 0.876 | [0.0479, 1] | 0.5889 | [1.5338E-8, 0.9889] |
| | ωλ0_5 | 0.9308 | [0.0844, 1] | 0.4919 | [2.6789E-8, 0.9622] |
| | ωλ0_6 | 0.8846 | [0.0386, 1] | 0.4299 | [7.6904E-11, 0.9807] |
| | ωλ0_7 | 0.9095 | [0.0914, 1] | 0.4471 | [1.6447E-9, 0.9727] |
| | ωλ0_8 | 0.6661 | [0.0127, 1] | 0.4051 | [1.7056E-8, 0.9653] |
| | ωλ0_9 | 0.9717 | [0.0777, 1] | 0.7171 | [1.5763E-8, 0.9882] |
| Shrinkage weights (extinction) | ωμ0_0 | 0.9108 | [0.059, 1] | 0.5544 | [4.7543E-3, 1] |
| | ωμ0_1 | 0.9282 | [0.0512, 1] | 0.461 | [1.1468E-8, 0.9877] |
| | ωμ0_2 | 0.905 | [0.0356, 1] | 0.359 | [4.1318E-9, 0.966] |
| | ωμ0_3 | **0.9828** | **[0.87, 1]** | 0.5886 | [7.2104E-9, 0.99] |
| | ωμ0_4 | 0.8096 | [0.029, 0.9999] | 0.8759 | [0.0125, 1] |
| | ωμ0_5 | **0.9889** | **[0.931, 1]** | 0.437 | [1.6787E-9, 0.9708] |
| | ωμ0_6 | 0.9535 | [0.1063, 1] | 0.495 | [7.6144E-11, 0.9905] |
| | ωμ0_7 | 0.7919 | [0.0284, 1] | **0.9762** | **[0.7391, 1]** |
| | ωμ0_8 | 0.7163 | [0.0181, 1] | 0.5162 | [1.638E-8, 0.9841] |
| | ωμ0_9 | 0.9585 | [0.0924, 1] | 0.5158 | [1.3701E-8, 0.9856] |

**table S11.** Continued.

| Parameters | | Cenozoic Median | Cenozoic 95% HPD |
|---|---|---|---|
| Baseline rates | λ0 | 9.89E-06 | [5.2604E-8, 5.2529E-4] |
| | μ0 | 5.54E-03 | [7.7131E-6, 0.1053] |
| Correlation parameters to origination | Gλ0_0 | **3.6611** | **[1.4104, 6.1192]** |
| | Gλ0_1 | **9.89E-03** | **[5.8754E-6, 0.0163]** |
| | Gλ0_2 | **38.6265** | **[13.2547, 61.0914]** |
| | Gλ0_3 | **-37.8095** | **[-50.1931, -26.6074]** |
| | Gλ0_4 | 3.38E-02 | [-0.0111, 0.0867] |
| | Gλ0_5 | -0.0042 | [-0.4627, 0.431] |
| | Gλ0_6 | **22.8815** | **[5.4722, 37.1435]** |
| | Gλ0_7 | **0.2751** | **[0.1939, 0.3486]** |
| | Gλ0_8 | **-1.97E+00** | **[-3.5033, -0.5202]** |
| | Gλ0_9 | **2.29E-01** | **[0.1271, 0.3367]** |
| Correlation parameters to extinction | Gμ0_0 | 2.1673 | [-0.618, 6.926] |
| | Gμ0_1 | -1.80E-03 | [-0.0107, 2.4806E-3] |
| | Gμ0_2 | -2.0827 | [-40.0086, 15.3646] |
| | Gμ0_3 | 12.8322 | [-1.8843, 31.3017] |
| | Gμ0_4 | **0.0894** | **[0.0197, 0.1643]** |
| | Gμ0_5 | -0.208 | [-0.8735, 0.1406] |
| | Gμ0_6 | -1.44E+01 | [-35.8109, 1.3272] |
| | Gμ0_7 | 6.41E-02 | [-0.0288, 0.1672] |
| | Gμ0_8 | 1.1583 | [-0.5646, 3.7203] |
| | Gμ0_9 | -4.48E-02 | [-0.1908, 0.0441] |
| Shrinkage weights (origination) | ωλ0_0 | **0.9389** | **[0.6806, 1]** |
| | ωλ0_1 | **0.969** | **[0.6513, 1]** |
| | ωλ0_2 | **0.9677** | **[0.8114, 1]** |
| | ωλ0_3 | **0.9671** | **[0.8555, 1]** |
| | ωλ0_4 | 0.6855 | [0.0322, 1] |
| | ωλ0_5 | 0.5723 | [9.4232E-8, 0.9796] |
| | ωλ0_6 | **0.9302** | **[0.5735, 1]** |
| | ωλ0_7 | **0.9391** | **[0.7354, 1]** |
| | ωλ0_8 | **0.8226** | **[0.3117, 0.9999]** |
| | ωλ0_9 | **0.9432** | **[0.7354, 1]** |
| Shrinkage weights (extinction) | ωμ0_0 | 0.884 | [0.0742, 1] |
| | ωμ0_1 | 0.8152 | [0.0284, 1] |
| | ωμ0_2 | 0.7348 | [0.019, 1] |
| | ωμ0_3 | 0.8551 | [0.0953, 1] |
| | ωμ0_4 | **0.8731** | **[0.3841, 1]** |
| | ωμ0_5 | 0.737 | [0.0306, 0.9999] |
| | ωμ0_6 | 0.8795 | [0.1369, 1] |
| | ωμ0_7 | 0.6955 | [0.0329, 0.9999] |
| | ωμ0_8 | 0.711 | [0.027, 1] |
| | ωμ0_9 | 0.7073 | [0.0203, 1] |

**table S12.**

Posterior parameter estimates for the MBD model applied to Polyphaga genera, excluding singletons and amber occurrences, across multiple temporal windows. The MBD model estimates the baseline origination and extinction rates ($\lambda 0$ and $\mu 0$), the correlation parameters ($G\lambda$ and $G\mu$) for each variable, and the shrinkage weights ($\omega$) of the correlation parameters. A variable was considered to have a significant effect (positive or negative depending on the sign of $G\lambda$ or $G\mu$) when its shrinkage weight exceeded 0.5 and when the 95% HPD interval of the corresponding correlation parameter did not overlap with zero (values highlighted in bold). The drivers are numbered as follows: (0) diversity of Polyphaga genera through time, (1) angiosperms diversity through time, (2) global variation of atmospheric $CO_2$ through time, (3) continental fragmentation through time, (4) gymnosperms diversity through time, (5) global variation in $\delta^{34}S$ through time (used here as an inverted proxy for global magmatic activity), (6) global variation of atmospheric $O_2$ through time, (7) Pteridophytes diversity through time, (8) Sea level fluctuations through time, and (9) variation of the global mean temperature through time. “All” corresponds to the time window encompassing the entire evolutionary history of Polyphaga genera, around 257 Ma to the present. “Before Upper Cretaceous” spans 257–100.5 Ma. The other time intervals are defined as follows: Triassic (251.902–201.4 Ma), Jurassic (201.4–143.1 Ma), Lower Cretaceous (143.1–100.5 Ma), Upper Cretaceous (100.5–66 Ma), and Cenozoic (66 Ma to the present).

| Parameters | | All | | Before Upper Cretaceous | |
|---|---|---|---|---|---|
| | | Median | 95% HPD | Median | 95% HPD |
| Baseline rates | λ0 | 8.50E-04 | [3.9735E-5, 3.2075E-3] | 5.32E-01 | [0.0432, 1.4446] |
| | μ0 | 6.45E-03 | [2.6791E-5, 0.0426] | 1.18E-01 | [2.0324E-5, 0.6202] |
| Correlation parameters to origination | Gλ0_0 | **-4.4537** | **[-5.5965, -3.2062]** | **-23.0296** | **[-31.3122, -14.0414]** |
| | Gλ0_1 | **7.70E-03** | **[6.3723E-3, 9.101E-3]** | **-1.49E-01** | **[-0.2068, -0.0885]** |
| | Gλ0_2 | 1.9737 | [-0.1675, 4.2143] | **12.7702** | **[6.8191, 18.0083]** |
| | Gλ0_3 | **-13.9865** | **[-17.5364, -10.2115]** | 3.6303 | [-5.1827, 14.8448] |
| | Gλ0_4 | **0.075** | **[0.0575, 0.0937]** | **-0.1145** | **[-0.1848, -0.0425]** |
| | Gλ0_5 | -0.0429 | [-0.1408, 0.0216] | -0.0763 | [-0.2493, 0.0811] |
| | Gλ0_6 | -0.2155 | [-2.1468, 1.1528] | 3.1044 | [-1.1348, 8.0023] |
| | Gλ0_7 | **0.0571** | **[0.0426, 0.0718]** | **0.1036** | **[0.0348, 0.1599]** |
| | Gλ0_8 | 1.42E-03 | [-1.1293, 0.8562] | 5.76E-01 | [-1.4038, 3.5222] |
| | Gλ0_9 | -0.0001 | [-0.0453, 0.0548] | **-0.1771** | **[-0.2888, -0.0767]** |
| Correlation parameters to extinction | Gμ0_0 | 0.592 | [-0.7803, 2.9724] | -0.9155 | [-25.6916, 9.4221] |
| | Gμ0_1 | -0.0032 | [-0.0064, 2.6368E-4] | 0.0375 | [-0.009, 0.1577] |
| | Gμ0_2 | 0.1189 | [-2.024, 2.8874] | **-16.696** | **[-27.8722, -6.1596]** |
| | Gμ0_3 | 0.3562 | [-3.9849, 5.8757] | 14.2178 | [-29.0497, 38.9388] |
| | Gμ0_4 | 0.0414 | [-0.0035, 0.0967] | **0.3137** | **[0.0738, 0.8681]** |
| | Gμ0_5 | -0.1808 | [-0.36, 4.5076E-3] | -0.3536 | [-0.6638, 2.9263E-3] |
| | Gμ0_6 | 0.9052 | [-0.5703, 3.5925] | 8.2124 | [-0.7846, 25.0763] |
| | Gμ0_7 | -0.0007 | [-0.0292, 0.0232] | **-0.2277** | **[-0.4183, -0.0789]** |
| | Gμ0_8 | 0.9212 | [-1.2405, 3.3855] | 1.8688 | [-2.2274, 4.9987] |
| | Gμ0_9 | -0.0128 | [-0.0821, 0.0244] | -0.3262 | [-1.1926, 0.0179] |
| Shrinkage weights (origination) | ωλ0_0 | **0.9465** | **[0.7826, 1]** | **0.9978** | **[0.9883, 1]** |
| | ωλ0_1 | **0.9694** | **[0.8692, 1]** | **0.9999** | **[0.9994, 1]** |
| | ωλ0_2 | 0.6934 | [0.0663, 0.9999] | **0.9875** | **[0.9138, 1]** |
| | ωλ0_3 | **0.9132** | **[0.6771, 1]** | 0.8918 | [0.0551, 1] |
| | ωλ0_4 | **0.9397** | **[0.7684, 1]** | **0.9848** | **[0.8802, 1]** |
| | ωλ0_5 | 0.6482 | [2.2149E-7, 0.9771] | 0.9372 | [0.1386, 1] |
| | ωλ0_6 | 0.426 | [6.5149E-10, 0.962] | 0.957 | [0.2425, 1] |
| | ωλ0_7 | **0.8895** | **[0.6164, 1]** | **0.98** | **[0.8324, 1]** |
| | ωλ0_8 | 0.3195 | [4.887E-9, 0.9482] | 0.8789 | [0.0401, 1] |
| | ωλ0_9 | 0.4538 | [1.5536E-7, 0.9643] | **0.9865** | **[0.9031, 1]** |
| Shrinkage weights (extinction) | ωμ0_0 | 0.5884 | [1.0606E-7, 0.975] | 0.9749 | [0.1955, 1] |
| | ωμ0_1 | 0.8732 | [0.18, 0.9999] | 0.9987 | [0.6746, 1] |
| | ωμ0_2 | 0.3858 | [2.0722E-8, 0.9595] | 0.9913 | [0.9289, 1] |
| | ωμ0_3 | 0.3491 | [5.0864E-8, 0.9504] | 0.9716 | [0.4475, 1] |
| | ωμ0_4 | 0.8647 | [0.1534, 1] | 0.9972 | [0.9652, 1] |
| | ωμ0_5 | 0.9311 | [0.2973, 1] | 0.9881 | [0.8341, 1] |
| | ωμ0_6 | 0.631 | [4.0412E-7, 0.9781] | 0.9891 | [0.6387, 1] |
| | ωμ0_7 | 0.3676 | [9.2581E-10, 0.9517] | 0.9934 | [0.9467, 1] |
| | ωμ0_8 | 0.6778 | [0.0214, 0.9992] | 0.9423 | [0.254, 1] |
| | ωμ0_9 | 0.5533 | [1.626E-7, 0.9756] | 0.9953 | [0.8149, 1] |

**table S12.** Continued.

| Parameters | | Triassic Median | Triassic 95% HPD | Jurassic Median | Jurassic 95% HPD |
|---|---|---|---|---|---|
| Baseline rates | λ0 | 2.02E-01 | [9.0466E-3, 0.8465] | 5.39E-01 | [0.038, 1.5082] |
| | μ0 | 2.76E-01 | [3.1421E-4, 1.0624] | 5.62E-01 | [0.0244, 1.5583] |
| Correlation parameters to origination | Gλ0_0 | -0.0064 | [-9.7569, 7.9437] | **-45.1088** | **[-60.3006, -30.1675]** |
| | Gλ0_1 | 6.38E-06 | [-0.0124, 0.0118] | -2.00E-04 | [-0.0939, 0.082] |
| | Gλ0_2 | -0.3247 | [-9.2226, 3.0094] | 4.0794 | [-3.1312, 17.9503] |
| | Gλ0_3 | -0.2228 | [-43.3292, 24.8545] | **77.3598** | **[21.1944, 128.6725]** |
| | Gλ0_4 | -0.0011 | [-0.064, 0.04] | -0.0498 | [-0.8709, 0.5221] |
| | Gλ0_5 | 1.80E-03 | [-0.1427, 0.1655] | -1.55E-01 | [-2.1459, 0.4361] |
| | Gλ0_6 | -0.2676 | [-5.34, 2.6276] | **-19.699** | **[-41.2905, -5.6057]** |
| | Gλ0_7 | -0.001 | [-0.1416, 0.0805] | -1.2659 | [-2.4955, 0.2008] |
| | Gλ0_8 | -5.00E-04 | [-2.4354, 3.0388] | 3.14E+00 | [-1.5168, 8.4284] |
| | Gλ0_9 | -0.0082 | [-0.1114, 0.0404] | 0.0158 | [-0.1404, 0.2375] |
| Correlation parameters to extinction | Gμ0_0 | -0.0003 | [-10.353, 10.4927] | 7.5334 | [-5.0568, 34.3898] |
| | Gμ0_1 | 2.07E-06 | [-0.0166, 0.0148] | 9.49E-05 | [-0.1464, 0.1438] |
| | Gμ0_2 | -0.5919 | [-34.4128, 8.5414] | **-24.4548** | **[-44.1909, -5.1009]** |
| | Gμ0_3 | -0.0464 | [-46.4525, 39.7801] | 5.2179 | [-92.7588, 117.5806] |
| | Gμ0_4 | -0.0013 | [-0.1996, 0.1045] | 0.238 | [-0.2694, 0.9892] |
| | Gμ0_5 | -0.0026 | [-0.7017, 0.3627] | 1.0224 | [-0.0858, 1.8786] |
| | Gμ0_6 | -0.517 | [-34.2531, 12.2607] | 25.619 | [-1.2398, 50.5994] |
| | Gμ0_7 | -0.0008 | [-0.3135, 0.1577] | 0.5526 | [-0.2242, 2.0973] |
| | Gμ0_8 | -0.0422 | [-14.0765, 7.1721] | 2.5361 | [-1.8782, 8.9332] |
| | Gμ0_9 | -0.0269 | [-0.4547, 0.1141] | -0.3198 | [-0.6434, 8.5868E-3] |
| Shrinkage weights (origination) | ωλ0_0 | 0.4154 | [2.4325E-9, 0.993] | **0.9994** | **[0.997, 1]** |
| | ωλ0_1 | 0.3847 | [2.1465E-8, 0.9934] | 0.9874 | [0.273, 1] |
| | ωλ0_2 | 0.4065 | [6.1839E-9, 0.9779] | 0.9657 | [0.1691, 1] |
| | ωλ0_3 | 0.4044 | [1.0097E-7, 0.9919] | **0.9968** | **[0.9734, 1]** |
| | ωλ0_4 | 0.2944 | [5.852E-9, 0.9576] | 0.9873 | [0.4272, 1] |
| | ωλ0_5 | 0.2888 | [2.8482E-8, 0.9569] | 0.9441 | [0.0843, 1] |
| | ωλ0_6 | 0.3248 | [2.2374E-9, 0.9601] | **0.9924** | **[0.9359, 1]** |
| | ωλ0_7 | 0.3675 | [4.7706E-10, 0.9837] | 0.9993 | [0.8759, 1] |
| | ωλ0_8 | 0.2801 | [9.9871E-9, 0.9543] | 0.9637 | [0.2219, 1] |
| | ωλ0_9 | 0.4025 | [9.5719E-9, 0.9717] | 0.9426 | [0.1037, 1] |
| Shrinkage weights (extinction) | ωμ0_0 | 0.4037 | [3.6975E-8, 0.994] | 0.9909 | [0.4378, 1] |
| | ωμ0_1 | 0.4069 | [2.3121E-8, 0.9957] | 0.9888 | [0.2922, 1] |
| | ωμ0_2 | 0.619 | [2.9512E-3, 1] | **0.9954** | **[0.9561, 1]** |
| | ωμ0_3 | 0.4102 | [1.0769E-9, 0.9945] | 0.9884 | [0.3436, 1] |
| | ωμ0_4 | 0.4128 | [1.1211E-9, 0.9914] | 0.9875 | [0.456, 1] |
| | ωμ0_5 | 0.4013 | [1.1208E-9, 0.9936] | 0.9886 | [0.8284, 1] |
| | ωμ0_6 | 0.5751 | [3.3735E-3, 1] | 0.9943 | [0.9011, 1] |
| | ωμ0_7 | 0.4288 | [2.1686E-9, 0.9954] | 0.998 | [0.5476, 1] |
| | ωμ0_8 | 0.4021 | [9.2239E-9, 0.9947] | 0.9595 | [0.2106, 1] |
| | ωμ0_9 | 0.7513 | [4.613E-3, 1] | 0.9909 | [0.8752, 1] |

**table S12.** Continued.

| Parameters | | Lower Cretaceous | | Upper Cretaceous | |
|---|---|---|---|---|---|
| | | Median | 95% HPD | Median | 95% HPD |
| Baseline rates | λ0 | 5.31E-01 | [0.0362, 1.5682] | 3.42E-01 | [0.0112, 1.0414] |
| | μ0 | 4.51E-01 | [0.0145, 1.3839] | 2.97E-01 | [5.0767E-3, 1.0599] |
| Correlation parameters to origination | Gλ0_0 | -0.0427 | [-14.9615, 11.2713] | -0.017 | [-8.3432, 6.1287] |
| | Gλ0_1 | -1.50E-03 | [-0.2126, 0.0335] | 3.81E-05 | [-0.0048, 8.4941E-3] |
| | Gλ0_2 | -0.0514 | [-39.2469, 21.9484] | -6.4688 | [-21.7154, 2.7737] |
| | Gλ0_3 | 2.308 | [-22.4308, 41.8524] | 0.0805 | [-11.8951, 17.6143] |
| | Gλ0_4 | -0.2004 | [-0.5115, 0.0454] | -0.0021 | [-0.1617, 0.0886] |
| | Gλ0_5 | -2.87E-01 | [-1.1134, 0.2011] | -1.80E-02 | [-0.4143, 0.1951] |
| | Gλ0_6 | -24.0147 | [-62.7172, 6.4959] | -0.2016 | [-28.3218, 22.1605] |
| | Gλ0_7 | 0.2121 | [-0.0175, 0.4685] | 2.38E-03 | [-0.0963, 0.1973] |
| | Gλ0_8 | 1.29E-01 | [-2.7118, 4.6897] | -1.87E-02 | [-2.4853, 2.3789] |
| | Gλ0_9 | 0.0124 | [-0.2894, 0.7383] | -0.007 | [-0.1988, 0.0878] |
| Correlation parameters to extinction | Gμ0_0 | -0.0007 | [-22.2692, 22.7097] | -0.0118 | [-7.6617, 6.4505] |
| | Gμ0_1 | -3.00E-03 | [-0.3805, 0.0408] | 0.00E+00 | [-0.0063, 5.2374E-3] |
| | Gμ0_2 | 1.5943 | [-8.6023, 34.5295] | -0.1412 | [-11.0494, 7.694] |
| | Gμ0_3 | 2.9127 | [-23.7596, 47.9693] | -0.0293 | [-21.7389, 25.5313] |
| | Gμ0_4 | -0.1745 | [-0.7559, 0.2758] | 3.02E-03 | [-0.1701, 0.4748] |
| | Gμ0_5 | -0.3758 | [-1.4826, 0.3005] | -0.0046 | [-0.3628, 0.2767] |
| | Gμ0_6 | 4.5381 | [-10.1675, 49.1238] | -0.824 | [-54.7256, 23.038] |
| | Gμ0_7 | -0.0028 | [-0.2129, 0.1077] | -0.1463 | [-0.6413, 0.0661] |
| | Gμ0_8 | -0.3591 | [-5.146, 1.8351] | -0.0058 | [-3.5928, 3.285] |
| | Gμ0_9 | -0.0093 | [-0.7618, 0.3493] | 1.62E-04 | [-0.1136, 0.1022] |
| Shrinkage weights (origination) | ωλ0_0 | 0.8625 | [0.0315, 1] | 0.2868 | [6.1254E-9, 0.9888] |
| | ωλ0_1 | 0.9706 | [0.0701, 1] | 0.2805 | [2.0183E-8, 0.9808] |
| | ωλ0_2 | 0.8493 | [0.026, 1] | 0.7514 | [8.5626E-8, 0.9881] |
| | ωλ0_3 | 0.8103 | [0.0208, 0.9999] | 0.2017 | [1.2675E-8, 0.9405] |
| | ωλ0_4 | 0.962 | [0.2289, 1] | 0.2478 | [7.9121E-8, 0.9607] |
| | ωλ0_5 | 0.887 | [0.0301, 1] | 0.2211 | [1.3692E-9, 0.9276] |
| | ωλ0_6 | 0.9875 | [0.2193, 1] | 0.256 | [4.7146E-10, 0.9699] |
| | ωλ0_7 | 0.978 | [0.4706, 1] | 0.2168 | [3.1563E-9, 0.9536] |
| | ωλ0_8 | 0.701 | [0.0161, 0.9998] | 0.212 | [1.8252E-8, 0.9335] |
| | ωλ0_9 | 0.8435 | [0.0201, 1] | 0.4385 | [5.6891E-9, 0.9772] |
| Shrinkage weights (extinction) | ωμ0_0 | 0.9209 | [0.0308, 1] | 0.3005 | [3.1694E-9, 0.9901] |
| | ωμ0_1 | 0.9894 | [0.0535, 1] | 0.2618 | [1.4813E-8, 0.9754] |
| | ωμ0_2 | 0.8715 | [0.0307, 1] | 0.235 | [7.402E-10, 0.9507] |
| | ωμ0_3 | 0.8436 | [0.0283, 1] | 0.2742 | [6.8728E-9, 0.9689] |
| | ωμ0_4 | 0.9637 | [0.2272, 1] | 0.3454 | [1.3172E-8, 0.9917] |
| | ωμ0_5 | 0.9196 | [0.1247, 1] | 0.186 | [1.6425E-9, 0.9258] |
| | ωμ0_6 | 0.9318 | [0.0506, 1] | 0.374 | [1.7663E-7, 0.9885] |
| | ωμ0_7 | 0.7889 | [0.0198, 1] | 0.8394 | [7.0394E-3, 1] |
| | ωμ0_8 | 0.7203 | [0.0153, 1] | 0.2508 | [1.545E-9, 0.9565] |
| | ωμ0_9 | 0.8726 | [0.0245, 1] | 0.2242 | [6.1383E-10, 0.9531] |

**table S12.** Continued.

| Parameters | | Cenozoic Median | Cenozoic 95% HPD |
|---|---|---|---|
| Baseline rates | λ0 | 6.98E-05 | [8.857E-8, 9.0715E-4] |
| | μ0 | 1.88E-01 | [3.4185E-4, 0.854] |
| Correlation parameters to origination | Gλ0_0 | **6.1177** | **[3.8041, 8.4268]** |
| | Gλ0_1 | 3.71E-03 | [-0.0013, 0.0114] |
| | Gλ0_2 | **40.3079** | **[20.2036, 63.7696]** |
| | Gλ0_3 | **-27.7012** | **[-38.3796, -16.8971]** |
| | Gλ0_4 | 0.059 | [-0.0113, 0.109] |
| | Gλ0_5 | -1.30E-02 | [-0.4297, 0.3619] |
| | Gλ0_6 | 3.8282 | [-5.2514, 25.9518] |
| | Gλ0_7 | **2.09E-01** | **[0.1296, 0.2969]** |
| | Gλ0_8 | -1.52E+00 | [-3.5471, 0.0553] |
| | Gλ0_9 | **0.1345** | **[0.0118, 0.2472]** |
| Correlation parameters to extinction | Gμ0_0 | -6.935 | [-11.4812, 0.1566] |
| | Gμ0_1 | -2.00E-04 | [-0.0091, 4.9558E-3] |
| | Gμ0_2 | -22.2803 | [-48.9468, 2.3158] |
| | Gμ0_3 | 6.6509 | [-9.9561, 37.2664] |
| | Gμ0_4 | 4.88E-02 | [-0.0139, 0.1432] |
| | Gμ0_5 | 0.0793 | [-0.3453, 0.7015] |
| | Gμ0_6 | -3.3281 | [-27.6132, 8.4945] |
| | Gμ0_7 | -0.0028 | [-0.1233, 0.1017] |
| | Gμ0_8 | 0.0625 | [-2.3544, 2.9837] |
| | Gμ0_9 | -6.27E-02 | [-0.2577, 0.0456] |
| Shrinkage weights (origination) | ωλ0_0 | **0.9705** | **[0.8632, 1]** |
| | ωλ0_1 | 0.874 | [0.0438, 1] |
| | ωλ0_2 | **0.97** | **[0.8477, 1]** |
| | ωλ0_3 | **0.9437** | **[0.752, 1]** |
| | ωλ0_4 | 0.7615 | [0.1271, 1] |
| | ωλ0_5 | 0.4908 | [1.034E-7, 0.9698] |
| | ωλ0_6 | 0.6733 | [1.3381E-7, 0.9853] |
| | ωλ0_7 | **0.9052** | **[0.617, 1]** |
| | ωλ0_8 | 0.7477 | [0.0963, 1] |
| | ωλ0_9 | **0.8658** | **[0.325, 1]** |
| Shrinkage weights (extinction) | ωμ0_0 | 0.9733 | [0.6787, 1] |
| | ωμ0_1 | 0.6588 | [6.7008E-8, 0.9864] |
| | ωμ0_2 | 0.9207 | [0.1804, 1] |
| | ωμ0_3 | 0.763 | [0.016, 1] |
| | ωμ0_4 | 0.7374 | [0.04, 1] |
| | ωμ0_5 | 0.6246 | [1.8538E-8, 0.9805] |
| | ωμ0_6 | 0.6429 | [2.2871E-8, 0.9853] |
| | ωμ0_7 | 0.4444 | [1.2896E-9, 0.9689] |
| | ωμ0_8 | 0.5306 | [3.1702E-8, 0.9768] |
| | ωμ0_9 | 0.7437 | [0.0215, 1] |

**table S13.**

Posterior parameter estimates for the MBD model applied to Adephaga genera, considering singletons and excluding amber occurrences, across multiple temporal windows. The MBD model estimates the baseline origination and extinction rates ($\lambda 0$ and $\mu 0$), the correlation parameters ($G\lambda$ and $G\mu$) for each variable, and the shrinkage weights ($\omega$) of the correlation parameters. A variable was considered to have a significant effect (positive or negative depending on the sign of $G\lambda$ or $G\mu$) when its shrinkage weight exceeded 0.5 and when the 95% HPD interval of the corresponding correlation parameter did not overlap with zero (values highlighted in bold). The drivers are numbered as follows: (0) diversity of Adephaga genera through time, (1) angiosperms diversity through time, (2) global variation of atmospheric $CO_2$ through time, (3) continental fragmentation through time, (4) gymnosperms diversity through time, (5) global variation in $\delta^{34}S$ through time (used here as an inverted proxy for global magmatic activity), (6) global variation of atmospheric $O_2$ through time, (7) Pteridophytes diversity through time, (8) Sea level fluctuations through time, and (9) variation of the global mean temperature through time. "All" corresponds to the time window encompassing the entire evolutionary history of Adephaga genera, around 254 Ma to the present. "Before Upper Cretaceous" spans 254–100.5 Ma. The other time intervals are defined as follows: Triassic (251.902–201.4 Ma), Jurassic (201.4–143.1 Ma), Lower Cretaceous (143.1–100.5 Ma), Upper Cretaceous (100.5–66 Ma), and Cenozoic (66 Ma to the present).

| Parameters | | All | | Before Upper Cretaceous | |
|---|---|---|---|---|---|
| | | Median | 95% HPD | Median | 95% HPD |
| Baseline rates | λ0 | 0.1377 | [0.0173, 0.3934] | 0.2694 | [0.0328, 0.8125] |
| | μ0 | 0.1826 | [1.8625E-3, 0.6352] | 0.154 | [5.1131E-4, 0.5948] |
| Correlation parameters to origination | Gλ0_0 | **-1.6893** | **[-2.982, -0.3663]** | -0.006 | [-1.9271, 1.8763] |
| | Gλ0_1 | 9.66E-04 | [-0.0004, 3.009E-3] | -1.00E-04 | [-0.0283, 7.1089E-3] |
| | Gλ0_2 | 0.3089 | [-1.0165, 2.6082] | 0.0915 | [-1.6261, 2.9275] |
| | Gλ0_3 | **-5.926** | **[-10.8948, -0.6225]** | -0.4465 | [-9.3667, 3.5036] |
| | Gλ0_4 | -0.0035 | [-0.0271, 0.0116] | -0.0378 | [-0.0886, 4.5012E-3] |
| | Gλ0_5 | -0.0081 | [-0.1017, 0.0435] | 3.02E-04 | [-0.0669, 0.0785] |
| | Gλ0_6 | 0.0901 | [-0.9773, 1.4498] | -0.0745 | [-1.7044, 0.9336] |
| | Gλ0_7 | **0.0313** | **[5.92E-3, 0.0542]** | 9.00E-03 | [-0.0085, 0.0515] |
| | Gλ0_8 | -1.3225 | [-2.8155, 0.1237] | -0.3819 | [-2.7739, 0.7072] |
| | Gλ0_9 | -0.0006 | [-0.0344, 0.0281] | 4.50E-04 | [-0.0288, 0.0359] |
| Correlation parameters to extinction | Gμ0_0 | 0.7247 | [-0.5389, 2.8014] | 0.0332 | [-1.3256, 2.1463] |
| | Gμ0_1 | -0.0039 | [-0.0067, 2.105E-5] | 5.95E-05 | [-0.0062, 0.0174] |
| | Gμ0_2 | 1.296 | [-0.7253, 4.7342] | -0.0274 | [-2.3921, 1.9572] |
| | Gμ0_3 | 0.0809 | [-4.6382, 6.2784] | 7.1606 | [-1.0203, 16.9521] |
| | Gμ0_4 | **-0.0671** | **[-0.1052, -0.0294]** | -0.0383 | [-0.0883, 7.0552E-3] |
| | Gμ0_5 | -0.0115 | [-0.1149, 0.043] | -0.0003 | [-0.0723, 0.0661] |
| | Gμ0_6 | 0.2301 | [-0.956, 2.1398] | 0.5642 | [-0.7515, 2.9557] |
| | Gμ0_7 | **0.0456** | **[0.0173, 0.0783]** | 0.0112 | [-0.01, 0.0523] |
| | Gμ0_8 | -0.9335 | [-3.3485, 0.4787] | -1.4995 | [-5.1663, 0.5172] |
| | Gμ0_9 | 0.0124 | [-0.018, 0.073] | 0.0116 | [-0.0169, 0.0856] |
| Shrinkage weights (origination) | ωλ0_0 | **0.7556** | **[0.2474, 0.9999]** | 0.2043 | [1.6132E-8, 0.9222] |
| | ωλ0_1 | 0.5093 | [3.7958E-8, 0.9594] | 0.3386 | [1.5019E-9, 0.9977] |
| | ωλ0_2 | 0.2631 | [4.1618E-8, 0.9184] | 0.1708 | [2.5904E-8, 0.8927] |
| | ωλ0_3 | **0.684** | **[0.1458, 1]** | 0.1923 | [5.6791E-8, 0.9125] |
| | ωλ0_4 | 0.2677 | [4.9916E-9, 0.9225] | 0.7532 | [0.0202, 1] |
| | ωλ0_5 | 0.3261 | [1.1403E-9, 0.9407] | 0.1869 | [2.8834E-8, 0.9029] |
| | ωλ0_6 | 0.1904 | [1.5469E-10, 0.8944] | 0.1233 | [8.8811E-10, 0.8512] |
| | ωλ0_7 | **0.7114** | **[0.1957, 0.9999]** | 0.2917 | [3.2276E-8, 0.9407] |
| | ωλ0_8 | 0.6397 | [0.0487, 0.9999] | 0.3188 | [5.7391E-8, 0.9402] |
| | ωλ0_9 | 0.2373 | [4.8156E-9, 0.9108] | 0.1472 | [5.5224E-10, 0.8747] |
| Shrinkage weights (extinction) | ωμ0_0 | 0.5249 | [1.3625E-8, 0.9655] | 0.1928 | [3.2995E-8, 0.9233] |
| | ωμ0_1 | 0.8886 | [0.3467, 1] | 0.3159 | [1.4968E-7, 0.9938] |
| | ωμ0_2 | 0.5175 | [2.3219E-7, 0.9637] | 0.1533 | [7.195E-9, 0.8803] |
| | ωμ0_3 | 0.2499 | [9.1157E-9, 0.9208] | 0.7062 | [5.2193E-10, 0.9806] |
| | ωμ0_4 | **0.9115** | **[0.6062, 1]** | 0.7664 | [0.0152, 1] |
| | ωμ0_5 | 0.3475 | [3.1915E-8, 0.9476] | 0.1684 | [1.0921E-8, 0.8941] |
| | ωμ0_6 | 0.2531 | [2.1343E-8, 0.9117] | 0.283 | [5.3109E-9, 0.9168] |
| | ωμ0_7 | **0.8163** | **[0.361, 1]** | 0.3795 | [5.8323E-9, 0.9474] |
| | ωμ0_8 | 0.5638 | [1.5749E-7, 0.9688] | 0.6672 | [1.3262E-7, 0.9816] |
| | ωμ0_9 | 0.4199 | [7.9149E-8, 0.9532] | 0.3346 | [3.3744E-7, 0.9528] |

**table S13.** Continued.

| Parameters | | Triassic Median | Triassic 95% HPD | Jurassic Median | Jurassic 95% HPD |
|---|---|---|---|---|---|
| Baseline rates | λ0 | 0.3842 | [0.0298, 1.1076] | 0.2061 | [0.025, 0.8182] |
| | μ0 | 0.2839 | [8.9213E-3, 0.9435] | 0.1127 | [9.3086E-3, 0.5866] |
| Correlation parameters to origination | Gλ0_0 | -7.1996 | [-56.7444, 3.605] | -0.9541 | [-20.6988, 1.9107] |
| | Gλ0_1 | 0.00E+00 | [-0.0123, 0.0118] | 8.87E-07 | [-0.0085, 9.4748E-3] |
| | Gλ0_2 | 0.1073 | [-3.4677, 5.9473] | 0.0336 | [-2.1316, 3.1613] |
| | Gλ0_3 | -0.1004 | [-35.6173, 28.0415] | -0.2041 | [-19.6959, 12.7794] |
| | Gλ0_4 | -0.0017 | [-0.0907, 0.0483] | -0.0009 | [-0.1253, 0.225] |
| | Gλ0_5 | 1.66E-02 | [-0.0864, 0.2513] | -1.22E-01 | [-0.6944, 0.0665] |
| | Gλ0_6 | -0.6039 | [-8.0013, 2.4933] | -1.3069 | [-15.3301, 2.233] |
| | Gλ0_7 | -4.00E-04 | [-0.1088, 0.0961] | -1.00E-04 | [-0.1081, 0.1216] |
| | Gλ0_8 | -0.1624 | [-6.1407, 2.6647] | -0.0315 | [-2.7289, 2.0144] |
| | Gλ0_9 | 2.13E-03 | [-0.0483, 0.0779] | -2.10E-03 | [-0.0702, 0.0383] |
| Correlation parameters to extinction | Gμ0_0 | -0.0069 | [-7.152, 8.2253] | -0.0002 | [-3.6761, 3.3171] |
| | Gμ0_1 | 3.50E-06 | [-0.0109, 0.0121] | 4.25E-06 | [-0.0083, 0.0106] |
| | Gμ0_2 | -0.1024 | [-7.295, 4.5543] | -1.1337 | [-6.2543, 0.9131] |
| | Gμ0_3 | 0.1103 | [-29.373, 40.3132] | 0.2158 | [-8.0029, 17.8439] |
| | Gμ0_4 | -0.0053 | [-0.1169, 0.0411] | -0.0005 | [-0.0944, 0.0807] |
| | Gμ0_5 | 0.1813 | [-0.0276, 0.4106] | -0.0196 | [-0.303, 0.1107] |
| | Gμ0_6 | -0.7475 | [-8.2958, 2.5326] | -0.026 | [-4.5559, 3.0712] |
| | Gμ0_7 | -0.0012 | [-0.1835, 0.0934] | 8.17E-04 | [-0.1051, 0.1707] |
| | Gμ0_8 | -0.8472 | [-8.8254, 1.6167] | -0.0888 | [-4.6424, 2.1994] |
| | Gμ0_9 | -0.0001 | [-0.0834, 0.0669] | -0.0042 | [-0.0804, 0.0303] |
| Shrinkage weights (origination) | ωλ0_0 | 0.9806 | [0.0158, 1] | 0.6349 | [1.0308E-8, 0.9981] |
| | ωλ0_1 | 0.4651 | [2.4784E-11, 0.9939] | 0.2377 | [1.6894E-9, 0.9894] |
| | ωλ0_2 | 0.3418 | [3.1583E-9, 0.9669] | 0.1506 | [6.3114E-8, 0.9125] |
| | ωλ0_3 | 0.4535 | [1.4904E-7, 0.9901] | 0.2574 | [1.1424E-8, 0.9695] |
| | ωλ0_4 | 0.4029 | [5.5127E-10, 0.9728] | 0.2606 | [3.4855E-9, 0.9763] |
| | ωλ0_5 | 0.4179 | [4.5018E-9, 0.9727] | 0.506 | [9.0797E-10, 0.9819] |
| | ωλ0_6 | 0.4786 | [2.0302E-7, 0.9777] | 0.4532 | [2.3061E-10, 0.9902] |
| | ωλ0_7 | 0.4247 | [4.7896E-8, 0.9786] | 0.2228 | [1.1782E-7, 0.9753] |
| | ωλ0_8 | 0.4975 | [1.818E-7, 0.98] | 0.2031 | [1.4314E-8, 0.9334] |
| | ωλ0_9 | 0.313 | [9.2956E-10, 0.9568] | 0.1662 | [2.4358E-10, 0.9037] |
| Shrinkage weights (extinction) | ωμ0_0 | 0.465 | [1.3965E-9, 0.9898] | 0.2028 | [1.1605E-11, 0.9663] |
| | ωμ0_1 | 0.4653 | [2.2158E-8, 0.9934] | 0.2443 | [1.9537E-8, 0.9888] |
| | ωμ0_2 | 0.3762 | [1.274E-9, 0.9732] | 0.4215 | [1.4106E-8, 0.9688] |
| | ωμ0_3 | 0.4604 | [2.466E-8, 0.9917] | 0.2045 | [1.4267E-8, 0.9604] |
| | ωμ0_4 | 0.4987 | [1.1258E-7, 0.9799] | 0.1579 | [2.3412E-9, 0.9309] |
| | ωμ0_5 | 0.8752 | [0.0384, 1] | 0.1894 | [1.5675E-9, 0.9302] |
| | ωμ0_6 | 0.5111 | [7.0901E-8, 0.9795] | 0.1589 | [1.0138E-10, 0.917] |
| | ωμ0_7 | 0.456 | [2.2942E-9, 0.9882] | 0.2335 | [3.55E-8, 0.9823] |
| | ωμ0_8 | 0.6827 | [1.7706E-7, 0.9913] | 0.2415 | [3.3381E-8, 0.9621] |
| | ωμ0_9 | 0.3322 | [1.7489E-8, 0.9596] | 0.1861 | [4.8352E-9, 0.9185] |

**table S13.** Continued.

| Parameters | | Lower Cretaceous | | Upper Cretaceous | |
|---|---|---|---|---|---|
| | | Median | 95% HPD | Median | 95% HPD |
| Baseline rates | λ0 | 0.2471 | [5.5906E-3, 1.0443] | 0.3316 | [0.0111, 1.1144] |
| | μ0 | 0.3577 | [0.0202, 1.1978] | 0.2949 | [6.4132E-3, 1.0773] |
| Correlation parameters to origination | Gλ0_0 | 3.85E-03 | [-2.1707, 2.2058] | -1.80E-02 | [-7.7571, 4.4194] |
| | Gλ0_1 | 0.00E+00 | [-0.021, 9.2081E-3] | 8.11E-06 | [-0.0044, 5.5626E-3] |
| | Gλ0_2 | -0.6986 | [-8.7498, 2.5462] | -0.4146 | [-12.9314, 4.8185] |
| | Gλ0_3 | -1.4355 | [-23.0444, 6.4095] | -0.153 | [-15.8841, 11.932] |
| | Gλ0_4 | 8.07E-05 | [-0.093, 0.1194] | -9.00E-04 | [-0.1529, 0.1108] |
| | Gλ0_5 | -6.00E-04 | [-0.1981, 0.2056] | -2.29E-01 | [-1.1342, 0.1385] |
| | Gλ0_6 | -0.1156 | [-5.798, 3.7511] | -0.2666 | [-27.1215, 16.0093] |
| | Gλ0_7 | 7.29E-03 | [-0.0207, 0.0796] | 1.16E-04 | [-0.1471, 0.1706] |
| | Gλ0_8 | -0.0223 | [-2.3559, 1.7812] | -0.1758 | [-5.2085, 1.7882] |
| | Gλ0_9 | -3.29E-02 | [-0.2446, 0.0397] | -4.40E-03 | [-0.1502, 0.0565] |
| Correlation parameters to extinction | Gμ0_0 | -0.2589 | [-4.4497, 1.1992] | -0.0041 | [-6.5625, 5.7909] |
| | Gμ0_1 | 9.82E-06 | [-0.0069, 8.9269E-3] | -1.00E-04 | [-0.0125, 5.0592E-3] |
| | Gμ0_2 | -0.0884 | [-5.3248, 3.7274] | 9.33E-03 | [-9.181, 8.4312] |
| | Gμ0_3 | 2.4973 | [-6.8466, 22.8372] | -1.7341 | [-35.3211, 9.2941] |
| | Gμ0_4 | -0.0167 | [-0.1461, 0.0404] | -0.0015 | [-0.2194, 0.1311] |
| | Gμ0_5 | **-0.6749** | **[-1.2062, -0.1197]** | 0.0113 | [-0.2748, 0.4884] |
| | Gμ0_6 | 0.28 | [-3.7809, 6.9869] | -0.6161 | [-40.5462, 18.6326] |
| | Gμ0_7 | -2.00E-04 | [-0.0495, 0.0387] | -1.31E-02 | [-0.3313, 0.0874] |
| | Gμ0_8 | -0.1551 | [-3.1423, 1.1218] | -0.0336 | [-4.2299, 2.6792] |
| | Gμ0_9 | -0.0008 | [-0.0883, 0.0728] | 2.67E-04 | [-0.0962, 0.1038] |
| Shrinkage weights (origination) | ωλ0_0 | 0.2065 | [1.927E-9, 0.9359] | 0.2581 | [5.6145E-8, 0.986] |
| | ωλ0_1 | 0.3053 | [1.7697E-8, 0.9965] | 0.2444 | [1.2181E-9, 0.9653] |
| | ωλ0_2 | 0.383 | [3.1904E-9, 0.9705] | 0.2564 | [6.0187E-12, 0.9561] |
| | ωλ0_3 | 0.2851 | [3.2789E-9, 0.9568] | 0.1935 | [2.9824E-10, 0.936] |
| | ωλ0_4 | 0.2191 | [1.7306E-10, 0.9433] | 0.2285 | [7.2208E-8, 0.9578] |
| | ωλ0_5 | 0.1644 | [3.8304E-9, 0.8961] | 0.6721 | [7.976E-8, 0.9879] |
| | ωλ0_6 | 0.193 | [1.5276E-9, 0.9356] | 0.2377 | [7.8001E-8, 0.9636] |
| | ωλ0_7 | 0.2441 | [1.7279E-9, 0.9422] | 0.2041 | [1.1248E-8, 0.9499] |
| | ωλ0_8 | 0.1989 | [6.0855E-11, 0.9287] | 0.3102 | [1.6212E-9, 0.9713] |
| | ωλ0_9 | 0.5862 | [1.1588E-8, 0.985] | 0.2846 | [8.6476E-8, 0.9624] |
| Shrinkage weights (extinction) | ωμ0_0 | 0.3532 | [7.7021E-11, 0.9669] | 0.2592 | [1.4751E-11, 0.9853] |
| | ωμ0_1 | 0.2918 | [1.4364E-9, 0.986] | 0.2745 | [1.2446E-9, 0.9883] |
| | ωμ0_2 | 0.2224 | [7.8111E-11, 0.9446] | 0.196 | [2.3253E-8, 0.9416] |
| | ωμ0_3 | 0.3597 | [7.629E-8, 0.9614] | 0.3709 | [8.9507E-11, 0.9776] |
| | ωμ0_4 | 0.3851 | [2.908E-9, 0.9635] | 0.2643 | [9.8436E-9, 0.9699] |
| | ωμ0_5 | **0.9448** | **[0.6107, 1]** | 0.2023 | [9.3496E-9, 0.9391] |
| | ωμ0_6 | 0.2574 | [1.0028E-8, 0.9485] | 0.2958 | [5.5375E-9, 0.9801] |
| | ωμ0_7 | 0.1661 | [3.9059E-9, 0.898] | 0.3678 | [5.2668E-8, 0.9781] |
| | ωμ0_8 | 0.2546 | [2.5886E-10, 0.9438] | 0.2438 | [1.2538E-8, 0.9633] |
| | ωμ0_9 | 0.2063 | [5.2721E-9, 0.926] | 0.2115 | [3.2357E-9, 0.9466] |

**table S13.** Continued.

| Parameters | | Cenozoic Median | Cenozoic 95% HPD |
|---|---|---|---|
| Baseline rates | λ0 | 0.1839 | [4.481E-3, 0.576] |
| | μ0 | 0.0525 | [6.6438E-4, 0.4373] |
| Correlation parameters to origination | Gλ0_0 | 7.28E-01 | [-1.0062, 6.6625] |
| | Gλ0_1 | -1.00E-04 | [-0.0058, 3.4219E-3] |
| | Gλ0_2 | 1.3985 | [-7.9306, 20.9489] |
| | Gλ0_3 | **-24.9926** | **[-43.328, -9.0891]** |
| | Gλ0_4 | **1.10E-01** | **[0.0241, 0.2423]** |
| | Gλ0_5 | -3.12E-01 | [-0.9476, 0.0784] |
| | Gλ0_6 | 0.2949 | [-7.2676, 11.1655] |
| | Gλ0_7 | 3.36E-03 | [-0.0533, 0.0855] |
| | Gλ0_8 | -0.5406 | [-3.3045, 0.655] |
| | Gλ0_9 | 3.42E-03 | [-0.0781, 0.1181] |
| Correlation parameters to extinction | Gμ0_0 | -0.026 | [-2.9459, 2.0374] |
| | Gμ0_1 | -1.10E-03 | [-0.0081, 1.4745E-3] |
| | Gμ0_2 | -2.16E+00 | [-28.3855, 7.3022] |
| | Gμ0_3 | -0.448 | [-16.4538, 9.7436] |
| | Gμ0_4 | -0.0053 | [-0.0947, 0.0361] |
| | Gμ0_5 | -0.0312 | [-0.6908, 0.2601] |
| | Gμ0_6 | -0.3023 | [-11.4702, 7.7246] |
| | Gμ0_7 | -1.00E-04 | [-0.1046, 0.1242] |
| | Gμ0_8 | -0.0411 | [-2.9545, 2.3161] |
| | Gμ0_9 | -8.00E-03 | [-0.1728, 0.08] |
| Shrinkage weights (origination) | ωλ0_0 | 0.5574 | [2.803E-9, 0.9881] |
| | ωλ0_1 | 0.2583 | [5.1789E-10, 0.9581] |
| | ωλ0_2 | 0.3299 | [8.2647E-9, 0.9558] |
| | ωλ0_3 | **0.913** | **[0.5916, 1]** |
| | ωλ0_4 | **0.8603** | **[0.3625, 1]** |
| | ωλ0_5 | 0.6676 | [8.7649E-9, 0.9801] |
| | ωλ0_6 | 0.1989 | [3.6767E-9, 0.9148] |
| | ωλ0_7 | 0.154 | [2.1486E-8, 0.8812] |
| | ωλ0_8 | 0.3475 | [2.64E-8, 0.9526] |
| | ωλ0_9 | 0.2383 | [7.0555E-9, 0.9248] |
| Shrinkage weights (extinction) | ωμ0_0 | 0.2619 | [1.4816E-8, 0.9476] |
| | ωμ0_1 | 0.5067 | [1.6873E-8, 0.98] |
| | ωμ0_2 | 0.3775 | [5.6178E-8, 0.9706] |
| | ωμ0_3 | 0.244 | [9.4786E-9, 0.9327] |
| | ωμ0_4 | 0.2405 | [1.4285E-11, 0.9281] |
| | ωμ0_5 | 0.2734 | [8.7901E-9, 0.9512] |
| | ωμ0_6 | 0.2073 | [3.2071E-9, 0.9124] |
| | ωμ0_7 | 0.2015 | [2.1057E-9, 0.9056] |
| | ωμ0_8 | 0.2414 | [1.803E-8, 0.9323] |
| | ωμ0_9 | 0.2927 | [4.4554E-9, 0.9463] |

**table S14.**

Posterior parameter estimates for the MBD model applied to Adephaga genera, excluding singletons and amber occurrences, across multiple temporal windows. The MBD model estimates the baseline origination and extinction rates ($\lambda 0$ and $\mu 0$), the correlation parameters ($G\lambda$ and $G\mu$) for each variable, and the shrinkage weights ($\omega$) of the correlation parameters. A variable was considered to have a significant effect (positive or negative depending on the sign of $G\lambda$ or $G\mu$) when its shrinkage weight exceeded 0.5 and when the 95% HPD interval of the corresponding correlation parameter did not overlap with zero (values highlighted in bold). The drivers are numbered as follows: (0) diversity of Adephaga genera through time, (1) angiosperms diversity through time, (2) global variation of atmospheric $CO_2$ through time, (3) continental fragmentation through time, (4) gymnosperms diversity through time, (5) global variation in $\delta^{34}S$ through time (used here as an inverted proxy for global magmatic activity), (6) global variation of atmospheric $O_2$ through time, (7) Pteridophytes diversity through time, (8) Sea level fluctuations through time, and (9) variation of the global mean temperature through time. "All" corresponds to the time window encompassing the entire evolutionary history of Adephaga genera, around 254 Ma to the present. "Before Upper Cretaceous" spans 254–100.5 Ma. The other time intervals are defined as follows: Triassic (251.902–201.4 Ma), Jurassic (201.4–143.1 Ma), Lower Cretaceous (143.1–100.5 Ma), Upper Cretaceous (100.5–66 Ma), and Cenozoic (66 Ma to the present).

| Parameters | | All | | Before Upper Cretaceous | |
|---|---|---|---|---|---|
| | | Median | 95% HPD | Median | 95% HPD |
| Baseline rates | λ0 | 0.1679 | [0.012, 0.567] | 0.3843 | [0.0244, 1.229] |
| | μ0 | 0.2395 | [7.1979E-3, 0.7899] | 0.3077 | [6.3625E-3, 0.9572] |
| Correlation parameters to origination | Gλ0_0 | **-4.9292** | **[-6.8319, -2.9424]** | -0.2822 | [-7.8897, 2.0648] |
| | Gλ0_1 | **5.40E-03** | **[2.6799E-3, 7.912E-3]** | -5.00E-04 | [-0.3022, 0.0123] |
| | Gλ0_2 | 1.948 | [-0.4851, 4.8224] | 1.5025 | [-1.0585, 5.936] |
| | Gλ0_3 | -4.3326 | [-9.7138, 0.9512] | -0.7078 | [-12.8085, 5.9501] |
| | Gλ0_4 | -0.0175 | [-0.0514, 6.8044E-3] | -0.0668 | [-0.1236, 3.2406E-3] |
| | Gλ0_5 | -0.0014 | [-0.0988, 0.071] | 1.51E-03 | [-0.09, 0.0996] |
| | Gλ0_6 | -0.1223 | [-1.945, 1.1665] | -0.6994 | [-3.3608, 0.8254] |
| | Gλ0_7 | 5.19E-03 | [-0.013, 0.0333] | 3.62E-05 | [-0.0415, 0.0599] |
| | Gλ0_8 | -0.5654 | [-2.4153, 0.5141] | -0.0356 | [-3.0786, 2.0849] |
| | Gλ0_9 | -0.0088 | [-0.0611, 0.0187] | -0.0016 | [-0.0605, 0.0389] |
| Correlation parameters to extinction | Gμ0_0 | 0.0556 | [-1.9857, 2.7215] | 1.66E-03 | [-2.5521, 2.624] |
| | Gμ0_1 | -0.0033 | [-0.0078, 4.9165E-4] | 3.37E-05 | [-0.0087, 0.0126] |
| | Gμ0_2 | 3.7989 | [-0.3009, 7.9812] | 6.31E-03 | [-3.1783, 3.301] |
| | Gμ0_3 | 4.5941 | [-2.4102, 15.5935] | **17.4843** | **[5.8896, 33.0879]** |
| | Gμ0_4 | **-0.1218** | **[-0.1692, -0.0778]** | **-0.0932** | **[-0.1438, -0.0332]** |
| | Gμ0_5 | -0.0029 | [-0.1036, 0.0673] | 6.67E-03 | [-0.059, 0.1233] |
| | Gμ0_6 | -0.0117 | [-2.0897, 1.8088] | -0.0032 | [-2.362, 2.1199] |
| | Gμ0_7 | 0.0431 | [-0.0058, 0.0847] | -0.0053 | [-0.0735, 0.0297] |
| | Gμ0_8 | -0.0564 | [-2.1039, 1.5596] | -0.3331 | [-3.6881, 1.1969] |
| | Gμ0_9 | 8.79E-04 | [-0.0401, 0.0468] | 1.02E-02 | [-0.0257, 0.0877] |
| Shrinkage weights (origination) | ωλ0_0 | **0.951** | **[0.7852, 1]** | 0.5015 | [6.1751E-10, 0.9874] |
| | ωλ0_1 | **0.9381** | **[0.7228, 1]** | 0.7253 | [7.6742E-3, 1] |
| | ωλ0_2 | 0.6397 | [3.1999E-8, 0.9757] | 0.5605 | [5.4958E-8, 0.9743] |
| | ωλ0_3 | 0.6038 | [7.8379E-8, 0.9669] | 0.3628 | [8.1329E-10, 0.9614] |
| | ωλ0_4 | 0.5902 | [5.2374E-8, 0.9685] | 0.9025 | [0.1977, 1] |
| | ωλ0_5 | 0.3542 | [4.6011E-8, 0.9463] | 0.3073 | [2.2925E-8, 0.9458] |
| | ωλ0_6 | 0.2683 | [1.7191E-8, 0.921] | 0.3772 | [2.7265E-8, 0.9519] |
| | ωλ0_7 | 0.3349 | [8.2765E-8, 0.9377] | 0.2972 | [1.3758E-10, 0.9537] |
| | ωλ0_8 | 0.4603 | [3.2462E-10, 0.9579] | 0.342 | [6.64E-9, 0.957] |
| | ωλ0_9 | 0.3949 | [2.0936E-7, 0.9528] | 0.2905 | [5.5222E-9, 0.9353] |
| Shrinkage weights (extinction) | ωμ0_0 | 0.433 | [1.2703E-7, 0.9598] | 0.3513 | [1.674E-7, 0.9596] |
| | ωμ0_1 | 0.8679 | [0.0994, 1] | 0.4673 | [3.4348E-9, 0.9922] |
| | ωμ0_2 | 0.8248 | [0.0734, 0.9999] | 0.2877 | [1.2046E-8, 0.9425] |
| | ωμ0_3 | 0.666 | [5.8961E-9, 0.9794] | **0.9345** | **[0.6573, 1]** |
| | ωμ0_4 | **0.9689** | **[0.8603, 1]** | **0.9457** | **[0.7211, 1]** |
| | ωμ0_5 | 0.3605 | [5.6835E-9, 0.9486] | 0.3266 | [2.277E-8, 0.9525] |
| | ωμ0_6 | 0.2838 | [1.6515E-9, 0.9316] | 0.2394 | [1.051E-9, 0.9258] |
| | ωμ0_7 | 0.7976 | [0.0854, 1] | 0.3939 | [4.3165E-9, 0.9644] |
| | ωμ0_8 | 0.3372 | [6.8924E-9, 0.9466] | 0.3984 | [2.4522E-9, 0.9663] |
| | ωμ0_9 | 0.3319 | [1.2263E-6, 0.949] | 0.4036 | [7.2145E-9, 0.9608] |

**table S14.** Continued.

| Parameters | | Triassic | | Jurassic | |
|---|---|---|---|---|---|
| | | Median | 95% HPD | Median | 95% HPD |
| Baseline rates | λ0 | 0.402 | [0.0125, 1.2343] | 0.2177 | [5.1542E-3, 0.9604] |
| | μ0 | 0.3435 | [5.0585E-3, 1.166] | 0.1255 | [8.8218E-4, 0.7394] |
| Correlation parameters to origination | Gλ0_0 | -1.4739 | [-72.7549, 4.8443] | -0.5932 | [-24.8193, 4.2757] |
| | Gλ0_1 | 0.00E+00 | [-0.0158, 0.0135] | 2.36E-06 | [-0.0126, 0.0146] |
| | Gλ0_2 | 0.1855 | [-4.0096, 7.1631] | 0.1743 | [-2.3896, 5.76] |
| | Gλ0_3 | -0.2725 | [-57.0086, 27.3253] | -2.1213 | [-63.2287, 24.3691] |
| | Gλ0_4 | -0.004 | [-0.1104, 0.0443] | -0.003 | [-0.19, 0.1723] |
| | Gλ0_5 | 1.46E-02 | [-0.1112, 0.249] | -1.04E-01 | [-0.7787, 0.1092] |
| | Gλ0_6 | -0.4877 | [-8.7759, 3.685] | -0.3681 | [-14.2081, 4.539] |
| | Gλ0_7 | -1.20E-03 | [-0.1413, 0.1031] | -2.30E-03 | [-0.3178, 0.1376] |
| | Gλ0_8 | -0.4745 | [-7.9838, 2.4969] | 0.1378 | [-2.845, 7.2093] |
| | Gλ0_9 | 9.64E-04 | [-0.0572, 0.0776] | -3.63E-02 | [-0.1574, 0.0283] |
| Correlation parameters to extinction | Gμ0_0 | -1.48E-02 | [-10.2383, 8.255] | 3.81E-02 | [-5.793, 8.4409] |
| | Gμ0_1 | 0.00E+00 | [-0.0178, 0.0138] | 4.05E-07 | [-0.013, 0.0145] |
| | Gμ0_2 | -1.20E-01 | [-10.1989, 5.1291] | -2.80E+00 | [-12.3242, 1.3868] |
| | Gμ0_3 | 0.0932 | [-33.2967, 39.3247] | 0.7124 | [-9.6775, 34.3539] |
| | Gμ0_4 | -0.0295 | [-0.1924, 0.0377] | 3.21E-04 | [-0.1282, 0.1803] |
| | Gμ0_5 | 1.37E-01 | [-0.0439, 0.4757] | -2.49E-02 | [-0.5649, 0.2485] |
| | Gμ0_6 | -0.1379 | [-7.2434, 3.8484] | -0.5077 | [-12.4909, 3.8448] |
| | Gμ0_7 | -0.0091 | [-0.5746, 0.0893] | 9.26E-04 | [-0.146, 0.2713] |
| | Gμ0_8 | -0.2048 | [-8.941, 2.8081] | -0.038 | [-4.7418, 3.5071] |
| | Gμ0_9 | -1.40E-03 | [-0.1228, 0.0713] | -1.76E-02 | [-0.1366, 0.0302] |
| Shrinkage weights (origination) | ωλ0_0 | 0.9004 | [9.9956E-3, 1] | 0.685 | [3.9405E-3, 1] |
| | ωλ0_1 | 0.524 | [3.0982E-9, 0.9954] | 0.4626 | [1.4931E-8, 0.9947] |
| | ωλ0_2 | 0.4025 | [5.5071E-8, 0.9742] | 0.3211 | [9.7385E-9, 0.9665] |
| | ωλ0_3 | 0.5147 | [3.3854E-10, 0.9945] | 0.6951 | [6.2637E-9, 0.9956] |
| | ωλ0_4 | 0.4863 | [1.0804E-8, 0.9789] | 0.4617 | [7.5349E-10, 0.9806] |
| | ωλ0_5 | 0.4554 | [9.4364E-8, 0.9724] | 0.5651 | [4.2533E-9, 0.9881] |
| | ωλ0_6 | 0.5344 | [9.5736E-9, 0.9832] | 0.4142 | [8.4826E-11, 0.9868] |
| | ωλ0_7 | 0.4618 | [1.1241E-8, 0.9853] | 0.5088 | [3.4093E-11, 0.9938] |
| | ωλ0_8 | 0.6217 | [1.6717E-8, 0.988] | 0.4408 | [7.0366E-9, 0.9854] |
| | ωλ0_9 | 0.3331 | [1.5604E-8, 0.9598] | 0.6176 | [5.9179E-9, 0.9803] |
| Shrinkage weights (extinction) | ωμ0_0 | 0.5157 | [2.781E-8, 0.9933] | 0.4266 | [1.0095E-9, 0.9915] |
| | ωμ0_1 | 0.5415 | [4.0826E-3, 1] | 0.4616 | [1.3504E-8, 0.9955] |
| | ωμ0_2 | 0.4624 | [1.0573E-8, 0.9809] | 0.7577 | [2.1081E-7, 0.9909] |
| | ωμ0_3 | 0.512 | [2.003E-10, 0.9919] | 0.4498 | [2.9217E-9, 0.9893] |
| | ωμ0_4 | 0.8157 | [0.0129, 1] | 0.3555 | [5.6262E-8, 0.9756] |
| | ωμ0_5 | 0.841 | [0.0106, 1] | 0.4087 | [5.4312E-8, 0.9746] |
| | ωμ0_6 | 0.4141 | [3.0592E-8, 0.9731] | 0.4405 | [7.2783E-10, 0.9853] |
| | ωμ0_7 | 0.7211 | [7.1461E-3, 1] | 0.4383 | [9.8212E-9, 0.9925] |
| | ωμ0_8 | 0.534 | [6.1641E-8, 0.9885] | 0.3815 | [2.2872E-9, 0.9768] |
| | ωμ0_9 | 0.4187 | [1.0562E-9, 0.9734] | 0.4715 | [5.9258E-8, 0.9755] |

**table S14.** Continued.

| Parameters | | Lower Cretaceous | | Upper Cretaceous | |
|---|---|---|---|---|---|
| | | Median | 95% HPD | Median | 95% HPD |
| Baseline rates | λ0 | 0.3529 | [9.0953E-3, 1.1011] | 0.3121 | [0.0132, 1.1064] |
| | μ0 | 0.3169 | [9.7074E-3, 1.0065] | 0.295 | [3.7053E-3, 1.1304] |
| Correlation parameters to origination | Gλ0_0 | -0.1794 | [-13.6519, 5.0363] | -0.0331 | [-9.9104, 6.2484] |
| | Gλ0_1 | -1.00E-04 | [-0.0405, 0.0217] | 1.34E-05 | [-0.0054, 6.8691E-3] |
| | Gλ0_2 | -0.1935 | [-10.5769, 8.6775] | -1.4327 | [-19.3458, 4.9286] |
| | Gλ0_3 | -2.6009 | [-48.7597, 19.1068] | 0.0251 | [-15.4937, 18.0556] |
| | Gλ0_4 | -0.0512 | [-0.3719, 0.092] | 1.77E-04 | [-0.1315, 0.159] |
| | Gλ0_5 | 1.11E-01 | [-0.2858, 1.2474] | -6.97E-02 | [-0.9452, 0.2119] |
| | Gλ0_6 | -5.9202 | [-36.3712, 3.9946] | -0.334 | [-31.006, 17.8057] |
| | Gλ0_7 | 6.39E-03 | [-0.0733, 0.2577] | -7.00E-04 | [-0.1799, 0.1289] |
| | Gλ0_8 | 0.0522 | [-3.5753, 5.3718] | -0.1151 | [-4.8146, 2.0787] |
| | Gλ0_9 | -1.73E-02 | [-0.2897, 0.1095] | -5.60E-03 | [-0.164, 0.0717] |
| Correlation parameters to extinction | Gμ0_0 | -5.51E-01 | [-9.2847, 2.3315] | -1.36E-02 | [-8.5888, 5.9233] |
| | Gμ0_1 | 1.24E-05 | [-0.0137, 0.0182] | 0.00E+00 | [-0.008, 6.1983E-3] |
| | Gμ0_2 | -7.99E-02 | [-7.9963, 6.2395] | -2.15E-01 | [-16.965, 9.2839] |
| | Gμ0_3 | 3.516 | [-10.7755, 38.0782] | -0.4586 | [-31.4129, 14.6915] |
| | Gμ0_4 | -6.88E-02 | [-0.2692, 0.0575] | -6.00E-04 | [-0.2084, 0.1972] |
| | Gμ0_5 | -1.58E-01 | [-0.8613, 0.1752] | -4.00E-03 | [-0.8774, 0.6001] |
| | Gμ0_6 | 2.2334 | [-3.4416, 15.1301] | -0.6878 | [-49.3355, 15.8431] |
| | Gμ0_7 | -7.70E-03 | [-0.1287, 0.0539] | -7.30E-03 | [-0.3613, 0.1304] |
| | Gμ0_8 | -0.4095 | [-4.3182, 1.4259] | -0.1615 | [-8.7107, 3.2797] |
| | Gμ0_9 | -6.00E-04 | [-0.1479, 0.1183] | -1.30E-03 | [-0.1617, 0.0951] |
| Shrinkage weights (origination) | ωλ0_0 | 0.6311 | [7.4208E-3, 1] | 0.3032 | [5.6829E-9, 0.9924] |
| | ωλ0_1 | 0.6159 | [6.4417E-3, 1] | 0.2562 | [4.0779E-10, 0.9758] |
| | ωλ0_2 | 0.5443 | [7.6746E-8, 0.9838] | 0.4366 | [5.7319E-10, 0.9794] |
| | ωλ0_3 | 0.6375 | [8.8832E-8, 0.99] | 0.2255 | [1.3466E-8, 0.9494] |
| | ωλ0_4 | 0.7815 | [0.0111, 1] | 0.2432 | [2.1938E-8, 0.9604] |
| | ωλ0_5 | 0.699 | [8.4679E-3, 0.9999] | 0.4085 | [3.3267E-10, 0.9791] |
| | ωλ0_6 | 0.872 | [0.0301, 1] | 0.2635 | [2.6176E-9, 0.9688] |
| | ωλ0_7 | 0.5659 | [4.222E-10, 0.9912] | 0.226 | [1.3122E-8, 0.9516] |
| | ωλ0_8 | 0.5027 | [7.8145E-8, 0.9802] | 0.2844 | [3.0396E-9, 0.9628] |
| | ωλ0_9 | 0.6622 | [0.0102, 1] | 0.3171 | [1.4805E-8, 0.9654] |
| Shrinkage weights (extinction) | ωμ0_0 | 0.7101 | [8.014E-3, 1] | 0.2884 | [3.7713E-9, 0.9903] |
| | ωμ0_1 | 0.6061 | [6.7784E-3, 1] | 0.273 | [4.5933E-11, 0.9814] |
| | ωμ0_2 | 0.4657 | [2.0548E-8, 0.978] | 0.2701 | [1.712E-8, 0.9681] |
| | ωμ0_3 | 0.604 | [3.089E-8, 0.9851] | 0.303 | [8.8525E-8, 0.9696] |
| | ωμ0_4 | 0.7945 | [0.0185, 0.9999] | 0.2695 | [4.5733E-9, 0.9789] |
| | ωμ0_5 | 0.7167 | [0.0135, 0.9998] | 0.2613 | [1.14E-8, 0.9699] |
| | ωμ0_6 | 0.6831 | [8.2999E-8, 0.9876] | 0.3502 | [5.5673E-8, 0.9853] |
| | ωμ0_7 | 0.5139 | [3.4255E-9, 0.9772] | 0.3461 | [1.9058E-9, 0.98] |
| | ωμ0_8 | 0.5218 | [1.5153E-8, 0.9758] | 0.3442 | [8.7663E-8, 0.9885] |
| | ωμ0_9 | 0.4756 | [2.2263E-7, 0.9762] | 0.2705 | [8.37E-10, 0.9639] |

**table S14.** Continued.

| Parameters | | Cenozoic | |
|---|---|---|---|
| | | Median | 95% HPD |
| Baseline rates | λ0 | 0.1792 | [9.5004E-5, 0.6029] |
| | μ0 | 0.1289 | [8.6225E-4, 0.6675] |
| Correlation parameters to origination | Gλ0_0 | 0.4886 | [-1.0905, 6.0202] |
| | Gλ0_1 | 0.00E+00 | [-0.0038, 3.6937E-3] |
| | Gλ0_2 | 1.2388 | [-9.5093, 21.0789] |
| | Gλ0_3 | **-24.6242** | **[-39.8623, -8.8976]** |
| | Gλ0_4 | 9.45E-02 | [-0.0021, 0.1867] |
| | Gλ0_5 | -3.39E-01 | [-0.9258, 0.0936] |
| | Gλ0_6 | -0.1662 | [-11.6579, 9.3176] |
| | Gλ0_7 | 3.43E-02 | [-0.0283, 0.1331] |
| | Gλ0_8 | -0.2809 | [-2.8806, 0.991] |
| | Gλ0_9 | 3.12E-03 | [-0.0828, 0.1351] |
| Correlation parameters to extinction | Gμ0_0 | -4.06E-01 | [-5.472, 1.7396] |
| | Gμ0_1 | -1.20E-03 | [-0.0092, 1.9783E-3] |
| | Gμ0_2 | -6.01E-01 | [-23.306, 13.3917] |
| | Gμ0_3 | -2.1462 | [-30.3749, 9.7426] |
| | Gμ0_4 | -4.70E-03 | [-0.1259, 0.0622] |
| | Gμ0_5 | -6.01E-02 | [-0.8513, 0.3345] |
| | Gμ0_6 | -3.6102 | [-26.1848, 7.145] |
| | Gμ0_7 | 3.13E-03 | [-0.1272, 0.1647] |
| | Gμ0_8 | -0.0855 | [-4.0586, 2.6162] |
| | Gμ0_9 | -1.91E-02 | [-0.3018, 0.0838] |
| Shrinkage weights (origination) | ωλ0_0 | 0.5094 | [2.0377E-7, 0.9832] |
| | ωλ0_1 | 0.3003 | [1.886E-8, 0.9469] |
| | ωλ0_2 | 0.3849 | [1.3047E-9, 0.959] |
| | ωλ0_3 | **0.9125** | **[0.6101, 1]** |
| | ωλ0_4 | 0.8247 | [0.2485, 1] |
| | ωλ0_5 | 0.6987 | [0.0246, 0.9999] |
| | ωλ0_6 | 0.2438 | [1.2275E-8, 0.9149] |
| | ωλ0_7 | 0.3608 | [1.2865E-9, 0.9347] |
| | ωλ0_8 | 0.3063 | [1.9419E-9, 0.9412] |
| | ωλ0_9 | 0.2831 | [6.7398E-10, 0.9348] |
| Shrinkage weights (extinction) | ωμ0_0 | 0.5397 | [5.4756E-13, 0.982] |
| | ωμ0_1 | 0.5981 | [2.6665E-7, 0.9833] |
| | ωμ0_2 | 0.3527 | [1.102E-9, 0.9655] |
| | ωμ0_3 | 0.4555 | [6.8674E-10, 0.9716] |
| | ωμ0_4 | 0.3512 | [7.1981E-8, 0.9512] |
| | ωμ0_5 | 0.431 | [3.9303E-9, 0.9647] |
| | ωμ0_6 | 0.5073 | [1.8938E-8, 0.9733] |
| | ωμ0_7 | 0.2818 | [3.6075E-8, 0.9454] |
| | ωμ0_8 | 0.3351 | [7.1842E-10, 0.9544] |
| | ωμ0_9 | 0.4706 | [7.9308E-8, 0.9754] |

**table S15.**

Posterior parameter estimates for the MBD model applied to Coleoptera genera without Polyphaga, considering singletons and excluding amber occurrences, across multiple temporal windows. The MBD model estimates the baseline origination and extinction rates ($\lambda 0$ and $\mu 0$), the correlation parameters ($G\lambda$ and $G\mu$) for each variable, and the shrinkage weights ($\omega$) of the correlation parameters. A variable was considered to have a significant effect (positive or negative depending on the sign of $G\lambda$ or $G\mu$) when its shrinkage weight exceeded 0.5 and when the 95% HPD interval of the corresponding correlation parameter did not overlap with zero (values highlighted in bold). The drivers are numbered as follows: (0) diversity of Coleoptera genera without Polyphaga through time, (1) angiosperms diversity through time, (2) global variation of atmospheric $CO_2$ through time, (3) continental fragmentation through time, (4) gymnosperms diversity through time, (5) global variation in $\delta^{34}S$ through time (used here as an inverted proxy for global magmatic activity), (6) global variation of atmospheric $O_2$ through time, (7) Pteridophytes diversity through time, (8) Sea level fluctuations through time, and (9) variation of the global mean temperature through time. “All” corresponds to the time window encompassing the entire evolutionary history of Coleoptera genera without Polyphaga, around 300 Ma to the present. “Before Upper Cretaceous” spans 300–100.5 Ma. The other time intervals are defined as follows: Permian (298.9–251.902 Ma), Triassic (251.902–201.4 Ma), Jurassic (201.4–143.1 Ma), Lower Cretaceous (143.1–100.5 Ma), Upper Cretaceous (100.5–66 Ma), and Cenozoic (66 Ma to the present).

| Parameters | | All | | Before Upper Cretaceous | |
|---|---|---|---|---|---|
| | | Median | 95% HPD | Median | 95% HPD |
| Baseline rates | λ0 | 0.1144 | [0.0141, 0.3451] | 0.2256 | [0.0113, 0.7068] |
| | μ0 | 0.0491 | [6.9687E-4, 0.2483] | 0.1783 | [7.4758E-3, 0.4925] |
| Correlation parameters to origination | Gλ0_0 | **0.8209** | **[0.1397, 1.4216]** | **1.4147** | **[0.4816, 2.4098]** |
| | Gλ0_1 | 5.55E-04 | [-0.0005, 2.1585E-3] | 2.78E-04 | [-0.0065, 0.0157] |
| | Gλ0_2 | **-1.6141** | **[-2.6979, -0.4629]** | **-1.5389** | **[-2.8174, -0.1947]** |
| | Gλ0_3 | -2.0168 | [-5.1497, 0.6198] | -0.7518 | [-6.0846, 2.6679] |
| | Gλ0_4 | -0.0116 | [-0.0287, 2.799E-3] | **-0.0358** | **[-0.0633, -0.01]** |
| | Gλ0_5 | **-0.0595** | **[-0.1084, -0.0135]** | -0.0379 | [-0.0857, 6.9314E-3] |
| | Gλ0_6 | **-1.3909** | **[-2.404, -0.3587]** | **-2.0851** | **[-3.3318, -0.7573]** |
| | Gλ0_7 | -0.0018 | [-0.0152, 6.9976E-3] | -0.0015 | [-0.0197, 0.012] |
| | Gλ0_8 | -0.0108 | [-0.67, 0.6387] | 0.2597 | [-0.5432, 1.4819] |
| | Gλ0_9 | **0.0547** | **[0.0296, 0.0835]** | **0.0547** | **[0.0227, 0.0899]** |
| Correlation parameters to extinction | Gμ0_0 | **2.0193** | **[1.4024, 2.7024]** | **3.0473** | **[2.1246, 4.0287]** |
| | Gμ0_1 | **-0.0046** | **[-0.0068, -0.0022]** | **0.0572** | **[0.034, 0.0783]** |
| | Gμ0_2 | **-2.4855** | **[-4.0049, -0.9071]** | **-2.9605** | **[-4.4498, -1.3308]** |
| | Gμ0_3 | 0.2417 | [-2.6537, 3.9627] | -1.7701 | [-7.9389, 2.3207] |
| | Gμ0_4 | -0.0148 | [-0.0411, 4.4258E-3] | -0.0089 | [-0.0364, 8.2584E-3] |
| | Gμ0_5 | 0.0228 | [-0.0112, 0.0672] | 0.0503 | [-0.0003, 0.0875] |
| | Gμ0_6 | -0.643 | [-1.8435, 0.2201] | -0.049 | [-1.1223, 0.7298] |
| | Gμ0_7 | 0.0124 | [-0.0024, 0.029] | -0.0118 | [-0.034, 5.5155E-3] |
| | Gμ0_8 | -0.711 | [-1.8313, 0.2164] | **-1.7863** | **[-3.0808, -0.4129]** |
| | Gμ0_9 | **0.0497** | **[8.3464E-3, 0.0922]** | 6.88E-03 | [-0.0193, 0.0471] |
| Shrinkage weights (origination) | ωλ0_0 | **0.5329** | **[0.0875, 0.9993]** | **0.7369** | **[0.2488, 1]** |
| | ωλ0_1 | 0.3935 | [2.3486E-8, 0.9491] | 0.6598 | [0.0108, 1] |
| | ωλ0_2 | **0.5667** | **[0.1236, 1]** | **0.5875** | **[0.1012, 0.9985]** |
| | ωλ0_3 | 0.3687 | [1.4307E-10, 0.929] | 0.3259 | [5.2942E-9, 0.9369] |
| | ωλ0_4 | 0.5118 | [1.4637E-7, 0.9564] | **0.8551** | **[0.3942, 1]** |
| | ωλ0_5 | **0.6721** | **[0.1878, 0.9999]** | 0.5673 | [0.0333, 0.9996] |
| | ωλ0_6 | **0.7052** | **[0.2032, 1]** | **0.8307** | **[0.3966, 1]** |
| | ωλ0_7 | 0.189 | [1.9094E-8, 0.8968] | 0.2732 | [2.8817E-9, 0.9284] |
| | ωλ0_8 | 0.1666 | [4.6349E-9, 0.8839] | 0.3509 | [9.1647E-9, 0.9375] |
| | ωλ0_9 | **0.8272** | **[0.4456, 1]** | **0.8376** | **[0.4237, 1]** |
| Shrinkage weights (extinction) | ωμ0_0 | **0.8073** | **[0.4443, 1]** | **0.9008** | **[0.6333, 0.9999]** |
| | ωμ0_1 | **0.9172** | **[0.6406, 1]** | **0.9993** | **[0.9965, 1]** |
| | ωμ0_2 | **0.7039** | **[0.2245, 1]** | **0.7773** | **[0.3381, 1]** |
| | ωμ0_3 | 0.2045 | [1.2888E-7, 0.8987] | 0.4365 | [1.9455E-7, 0.9559] |
| | ωμ0_4 | 0.6021 | [1.62E-7, 0.9711] | 0.519 | [1.8808E-8, 0.9664] |
| | ωμ0_5 | 0.3847 | [4.4948E-8, 0.9416] | 0.6438 | [0.1089, 1] |
| | ωμ0_6 | 0.4683 | [2.5249E-7, 0.9459] | 0.2671 | [5.7675E-8, 0.9297] |
| | ωμ0_7 | 0.4988 | [2.1051E-7, 0.9562] | 0.5328 | [9.1028E-9, 0.9623] |
| | ωμ0_8 | 0.4995 | [3.9071E-8, 0.9559] | **0.7914** | **[0.2955, 1]** |
| | ωμ0_9 | **0.8065** | **[0.3008, 0.9999]** | 0.4049 | [2.6465E-7, 0.9503] |

**table S15.** Continued.

| Parameters | | Permian | | Triassic | |
|---|---|---|---|---|---|
| | | Median | 95% HPD | Median | 95% HPD |
| Baseline rates | λ0 | 0.6332 | [0.0869, 1.5409] | 0.595 | [0.0378, 1.6629] |
| | μ0 | 0.6274 | [0.0588, 1.6537] | 0.586 | [0.0522, 1.6368] |
| Correlation parameters to origination | Gλ0_0 | **-10.8868** | **[-18.1094, -3.7856]** | -1.1072 | [-5.4605, 1.4895] |
| | Gλ0_1 | 0.00E+00 | [-0.0181, 0.0193] | 2.09E-05 | [-0.032, 0.03] |
| | Gλ0_2 | 0.313 | [-2.3903, 4.6149] | 8.5832 | [-0.0978, 17.0688] |
| | Gλ0_3 | 0.0366 | [-23.2995, 31.3627] | 41.0954 | [-6.7042, 94.8136] |
| | Gλ0_4 | 5.43E-04 | [-0.068, 0.1015] | 1.50E-02 | [-0.0578, 0.2187] |
| | Gλ0_5 | 0.0138 | [-0.0643, 0.1609] | 7.14E-03 | [-0.1526, 0.2745] |
| | Gλ0_6 | 0.0505 | [-1.4929, 2.2191] | **-11.7158** | **[-16.3693, -6.9738]** |
| | Gλ0_7 | 1.95E-04 | [-0.0814, 0.0914] | -4.22E-02 | [-0.3667, 0.076] |
| | Gλ0_8 | 0.0562 | [-1.5579, 2.6223] | -0.0004 | [-3.8288, 3.075] |
| | Gλ0_9 | -0.0023 | [-0.0683, 0.0351] | -0.0174 | [-0.164, 0.0788] |
| Correlation parameters to extinction | Gμ0_0 | -0.3148 | [-9.4099, 3.1662] | 2.4808 | [-0.8729, 8.2525] |
| | Gμ0_1 | 0 | [-0.0146, 0.015] | 3.93E-06 | [-0.0282, 0.0326] |
| | Gμ0_2 | 0.3118 | [-2.9299, 5.9137] | -2.1291 | [-20.3257, 4.0167] |
| | Gμ0_3 | 1.7911 | [-17.0287, 76.1994] | 9.386 | [-17.3299, 78.6351] |
| | Gμ0_4 | 2.95E-04 | [-0.1071, 0.1333] | 1.15E-02 | [-0.0561, 0.1596] |
| | Gμ0_5 | 0.0251 | [-0.0672, 0.2357] | 0.0418 | [-0.1358, 0.2575] |
| | Gμ0_6 | 2.2994 | [-1.0259, 7.5392] | **-9.4043** | **[-16.9967, -3.601]** |
| | Gμ0_7 | -0.0009 | [-0.1766, 0.1058] | -0.1287 | [-0.6585, 0.0519] |
| | Gμ0_8 | 0.0763 | [-2.3278, 3.1464] | -0.2689 | [-4.0885, 2.4819] |
| | Gμ0_9 | -1.20E-01 | [-0.2309, 4.9006E-3] | 7.14E-02 | [-0.0421, 0.3755] |
| Shrinkage weights (origination) | ωλ0_0 | **0.9885** | **[0.9077, 1]** | 0.8053 | [0.0283, 1] |
| | ωλ0_1 | 0.5677 | [6.1301E-3, 1] | 0.8732 | [0.0332, 1] |
| | ωλ0_2 | 0.4002 | [3.9505E-8, 0.9629] | 0.9608 | [0.6171, 1] |
| | ωλ0_3 | 0.5117 | [3.1801E-8, 0.9884] | 0.9858 | [0.258, 1] |
| | ωλ0_4 | 0.5028 | [8.7999E-10, 0.9863] | 0.8267 | [0.0215, 1] |
| | ωλ0_5 | 0.4825 | [8.3186E-8, 0.9702] | 0.6813 | [0.0136, 1] |
| | ωλ0_6 | 0.3213 | [1.9253E-9, 0.9517] | **0.9766** | **[0.8846, 1]** |
| | ωλ0_7 | 0.4984 | [6.1633E-9, 0.9844] | 0.9199 | [0.0483, 1] |
| | ωλ0_8 | 0.3585 | [3.4831E-8, 0.9605] | 0.6571 | [3.995E-9, 0.987] |
| | ωλ0_9 | 0.347 | [4.022E-10, 0.9595] | 0.791 | [0.0254, 1] |
| Shrinkage weights (extinction) | ωμ0_0 | 0.6394 | [1.6088E-7, 0.992] | 0.9064 | [0.0642, 1] |
| | ωμ0_1 | 0.5476 | [5.7921E-3, 1] | 0.8719 | [0.0267, 1] |
| | ωμ0_2 | 0.4342 | [3.2406E-9, 0.971] | 0.8678 | [0.0273, 1] |
| | ωμ0_3 | 0.7071 | [8.3872E-3, 1] | 0.9369 | [0.0559, 1] |
| | ωμ0_4 | 0.5522 | [4.5157E-10, 0.991] | 0.7959 | [0.0227, 1] |
| | ωμ0_5 | 0.585 | [7.1658E-7, 0.9825] | 0.7709 | [0.0323, 1] |
| | ωμ0_6 | 0.8451 | [0.0148, 1] | **0.966** | **[0.7888, 1]** |
| | ωμ0_7 | 0.5829 | [9.1511E-9, 0.992] | 0.9746 | [0.0844, 1] |
| | ωμ0_8 | 0.4284 | [2.3851E-8, 0.9673] | 0.6994 | [0.0138, 0.9999] |
| | ωμ0_9 | 0.947 | [0.2331, 1] | 0.9116 | [0.0414, 1] |

**table S15.** Continued.

| Parameters | | Jurassic | | Lower Cretaceous | |
|---|---|---|---|---|---|
| | | Median | 95% HPD | Median | 95% HPD |
| Baseline rates | λ0 | 1.25E-03 | [1.1526E-7, 0.0572] | 1.06E-01 | [1.1264E-4, 0.5632] |
| | μ0 | 0.0326 | [1.5935E-5, 0.2562] | 0.4973 | [0.0272, 1.348] |
| Correlation parameters to origination | Gλ0_0 | **4.2755** | **[2.8356, 5.8097]** | -3.1602 | [-10.8353, 1.0137] |
| | Gλ0_1 | 7.88E-06 | [-0.0459, 0.0487] | 0.00E+00 | [-0.0196, 0.0157] |
| | Gλ0_2 | 4.1552 | [-1.957, 12.5533] | -2.3098 | [-32.2717, 4.9463] |
| | Gλ0_3 | 13.1509 | [-31.6116, 69.7179] | -2.9061 | [-32.946, 12.7626] |
| | Gλ0_4 | -3.20E-01 | [-0.6397, 0.0155] | 1.54E-01 | [-0.0779, 0.8949] |
| | Gλ0_5 | -4.17E-01 | [-0.7017, 0.0107] | -1.13E-01 | [-0.6108, 0.2001] |
| | Gλ0_6 | -7.0496 | [-12.8362, 0.0254] | 0.1441 | [-6.9917, 11.8663] |
| | Gλ0_7 | -3.30E-03 | [-0.8613, 0.4862] | -4.85E-02 | [-0.2293, 0.0151] |
| | Gλ0_8 | 3.851 | [-0.0643, 7.3446] | 1.1707 | [-0.8381, 5.7496] |
| | Gλ0_9 | **0.1773** | **[0.0293, 0.3211]** | 1.36E-03 | [-0.1496, 0.2171] |
| Correlation parameters to extinction | Gμ0_0 | **5.7628** | **[3.8712, 7.4]** | -0.1625 | [-3.372, 2.6374] |
| | Gμ0_1 | 0.00E+00 | [-0.0336, 0.0318] | -2.00E-04 | [-0.1162, 0.0159] |
| | Gμ0_2 | -3.8783 | [-9.8316, 0.9335] | 3.8961 | [-3.1713, 36.4773] |
| | Gμ0_3 | 12.4499 | [-6.502, 51.0323] | 7.4315 | [-7.6617, 42.5314] |
| | Gμ0_4 | -9.42E-02 | [-0.3237, 0.0825] | -1.03E-01 | [-0.5963, 0.0598] |
| | Gμ0_5 | 0.1237 | [-0.0844, 0.4352] | **-0.7504** | **[-1.7113, -0.2265]** |
| | Gμ0_6 | 0.9953 | [-2.7289, 5.9472] | -0.014 | [-12.7373, 10.3887] |
| | Gμ0_7 | -0.0588 | [-0.5953, 0.1307] | -0.0027 | [-0.1047, 0.0991] |
| | Gμ0_8 | -0.3861 | [-3.9988, 1.9605] | -0.4725 | [-3.0798, 1.0711] |
| | Gμ0_9 | 1.39E-02 | [-0.054, 0.1161] | -3.70E-03 | [-0.5039, 0.1691] |
| Shrinkage weights (origination) | ωλ0_0 | **0.9559** | **[0.7912, 1]** | 0.926 | [0.0165, 1] |
| | ωλ0_1 | 0.9126 | [0.0464, 1] | 0.7106 | [9.1585E-3, 1] |
| | ωλ0_2 | 0.9081 | [0.1125, 1] | 0.8353 | [9.5278E-3, 1] |
| | ωλ0_3 | 0.9599 | [0.1603, 1] | 0.6342 | [2.2122E-7, 0.9869] |
| | ωλ0_4 | 0.9818 | [0.74, 1] | 0.9461 | [0.0125, 1] |
| | ωλ0_5 | 0.9256 | [0.4951, 1] | 0.6746 | [0.0147, 1] |
| | ωλ0_6 | 0.9352 | [0.482, 1] | 0.5778 | [1.5448E-7, 0.9877] |
| | ωλ0_7 | 0.9316 | [0.0568, 1] | 0.8272 | [0.0178, 1] |
| | ωλ0_8 | 0.9406 | [0.4288, 1] | 0.7279 | [7.7075E-9, 0.9906] |
| | ωλ0_9 | **0.9636** | **[0.6983, 1]** | 0.591 | [1.2608E-7, 0.9884] |
| Shrinkage weights (extinction) | ωμ0_0 | **0.971** | **[0.8632, 1]** | 0.5977 | [1.2897E-8, 0.9836] |
| | ωμ0_1 | 0.9083 | [0.0461, 1] | 0.772 | [8.7232E-3, 1] |
| | ωμ0_2 | 0.8984 | [0.1718, 1] | 0.8651 | [0.0129, 1] |
| | ωμ0_3 | 0.9345 | [0.0882, 1] | 0.7637 | [0.0147, 1] |
| | ωμ0_4 | 0.907 | [0.1014, 1] | 0.8873 | [0.0208, 1] |
| | ωμ0_5 | 0.7592 | [0.0241, 1] | **0.965** | **[0.7332, 1]** |
| | ωμ0_6 | 0.7001 | [0.0165, 0.9999] | 0.5707 | [2.6187E-8, 0.9878] |
| | ωμ0_7 | 0.9496 | [0.0647, 1] | 0.6262 | [9.1189E-8, 0.984] |
| | ωμ0_8 | 0.7284 | [0.0194, 1] | 0.5533 | [3.4919E-7, 0.9779] |
| | ωμ0_9 | 0.7037 | [0.0153, 0.9998] | 0.6315 | [6.2825E-3, 1] |

**table S15.** Continued.

| Parameters | | Upper Cretaceous | | Cenozoic | |
|---|---|---|---|---|---|
| | | Median | 95% HPD | Median | 95% HPD |
| Baseline rates | λ0 | 2.49E-01 | [8.0909E-3, 0.8507] | 1.35E-01 | [1.7998E-3, 0.4283] |
| | μ0 | 0.2719 | [3.0405E-3, 0.9146] | 0.0185 | [5.6006E-4, 0.142] |
| Correlation parameters to origination | Gλ0_0 | -0.044 | [-6.0008, 3.8039] | 0.1623 | [-0.8397, 4.8043] |
| | Gλ0_1 | 0.00E+00 | [-0.0066, 4.6453E-3] | 0.00E+00 | [-0.0027, 2.7757E-3] |
| | Gλ0_2 | -0.1929 | [-9.065, 5.3342] | 0.6238 | [-7.4744, 14.0159] |
| | Gλ0_3 | -0.4766 | [-19.2716, 10.6591] | **-21.89** | **[-37.3243, -7.9388]** |
| | Gλ0_4 | -8.00E-04 | [-0.1741, 0.1191] | 7.41E-02 | [-0.0026, 0.161] |
| | Gλ0_5 | -9.55E-01 | [-1.9681, 0.0227] | -1.65E-01 | [-0.6798, 0.0672] |
| | Gλ0_6 | -0.7661 | [-37.8886, 16.7157] | 0.1788 | [-5.613, 9.1885] |
| | Gλ0_7 | -6.00E-04 | [-0.1777, 0.158] | 2.53E-03 | [-0.0438, 0.079] |
| | Gλ0_8 | -0.0204 | [-2.9845, 2.5699] | -0.1578 | [-2.7013, 0.8294] |
| | Gλ0_9 | -4.00E-04 | [-0.0944, 0.0955] | 7.10E-03 | [-0.0526, 0.1382] |
| Correlation parameters to extinction | Gμ0_0 | 0.0245 | [-3.649, 5.8787] | -0.0401 | [-2.0536, 1.165] |
| | Gμ0_1 | -3.00E-04 | [-0.0253, 4.5031E-3] | -1.00E-04 | [-0.0044, 1.4742E-3] |
| | Gμ0_2 | 0.1937 | [-7.3547, 11.5612] | -0.2459 | [-13.1126, 5.697] |
| | Gμ0_3 | -10.4625 | [-57.7195, 10.6869] | -0.0999 | [-11.1931, 7.1489] |
| | Gμ0_4 | 5.58E-04 | [-0.2056, 0.2799] | -3.00E-04 | [-0.0435, 0.0369] |
| | Gμ0_5 | 0.0156 | [-0.196, 0.3715] | -0.005 | [-0.2981, 0.2125] |
| | Gμ0_6 | -0.6754 | [-56.8044, 21.496] | -0.4204 | [-10.0833, 4.3668] |
| | Gμ0_7 | -0.0042 | [-0.2392, 0.1264] | 3.94E-04 | [-0.0761, 0.0772] |
| | Gμ0_8 | -0.0505 | [-3.4018, 1.9175] | 7.37E-03 | [-1.7581, 1.782] |
| | Gμ0_9 | 6.53E-03 | [-0.0643, 0.2097] | -2.10E-03 | [-0.119, 0.0587] |
| Shrinkage weights (origination) | ωλ0_0 | 0.3284 | [5.2917E-8, 0.981] | 0.2108 | [5.9752E-9, 0.9664] |
| | ωλ0_1 | 0.2952 | [1.3347E-9, 0.9732] | 0.1162 | [4.6903E-9, 0.8797] |
| | ωλ0_2 | 0.2222 | [1.4383E-8, 0.944] | 0.1547 | [3.6003E-9, 0.9013] |
| | ωλ0_3 | 0.2622 | [2.0098E-10, 0.9544] | **0.8876** | **[0.5111, 1]** |
| | ωλ0_4 | 0.2877 | [2.0819E-9, 0.9643] | 0.7166 | [0.0779, 1] |
| | ωλ0_5 | 0.956 | [0.4864, 1] | 0.3906 | [4.8388E-11, 0.9549] |
| | ωλ0_6 | 0.3186 | [2.663E-8, 0.9793] | 0.0957 | [8.3843E-10, 0.8437] |
| | ωλ0_7 | 0.2669 | [1.1904E-7, 0.9647] | 0.0878 | [5.4477E-10, 0.7894] |
| | ωλ0_8 | 0.2514 | [8.4745E-10, 0.9491] | 0.1455 | [2.5766E-8, 0.8994] |
| | ωλ0_9 | 0.2442 | [5.0311E-8, 0.9513] | 0.1616 | [2.3889E-9, 0.9174] |
| Shrinkage weights (extinction) | ωμ0_0 | 0.3203 | [1.5667E-9, 0.979] | 0.1205 | [2.005E-9, 0.9011] |
| | ωμ0_1 | 0.5088 | [3.4395E-8, 0.9972] | 0.1397 | [3.0192E-9, 0.921] |
| | ωμ0_2 | 0.2751 | [5.051E-8, 0.9558] | 0.1072 | [5.3291E-15, 0.8826] |
| | ωμ0_3 | 0.7548 | [7.7861E-9, 0.9938] | 0.1019 | [9.1369E-10, 0.8546] |
| | ωμ0_4 | 0.3233 | [1.3805E-8, 0.9859] | 0.0858 | [1.3087E-9, 0.818] |
| | ωμ0_5 | 0.224 | [5.3043E-9, 0.9277] | 0.0945 | [7.4315E-9, 0.84] |
| | ωμ0_6 | 0.3723 | [8.2077E-11, 0.9866] | 0.0982 | [7.6992E-10, 0.8502] |
| | ωμ0_7 | 0.3013 | [3.8694E-9, 0.9648] | 0.0934 | [1.3286E-9, 0.8206] |
| | ωμ0_8 | 0.2745 | [7.6516E-10, 0.9486] | 0.1067 | [1.0603E-9, 0.8574] |
| | ωμ0_9 | 0.3359 | [3.8718E-8, 0.9753] | 0.1153 | [3.7839E-10, 0.8816] |

**table S16.**

Posterior parameter estimates for the MBD model applied to Coleoptera genera without Polyphaga, excluding singletons and amber occurrences, across multiple temporal windows. The MBD model estimates the baseline origination and extinction rates ($\lambda 0$ and $\mu 0$), the correlation parameters ($G\lambda$ and $G\mu$) for each variable, and the shrinkage weights ($\omega$) of the correlation parameters. A variable was considered to have a significant effect (positive or negative depending on the sign of $G\lambda$ or $G\mu$) when its shrinkage weight exceeded 0.5 and when the 95% HPD interval of the corresponding correlation parameter did not overlap with zero (values highlighted in bold). The drivers are numbered as follows: (0) diversity of Coleoptera genera without Polyphaga through time, (1) angiosperms diversity through time, (2) global variation of atmospheric $CO_2$ through time, (3) continental fragmentation through time, (4) gymnosperms diversity through time, (5) global variation in $\delta^{34}S$ through time (used here as an inverted proxy for global magmatic activity), (6) global variation of atmospheric $O_2$ through time, (7) Pteridophytes diversity through time, (8) Sea level fluctuations through time, and (9) variation of the global mean temperature through time. "All" corresponds to the time window encompassing the entire evolutionary history of Coleoptera genera without Polyphaga, around 300 Ma to the present. "Before Upper Cretaceous" spans 300–100.5 Ma. The other time intervals are defined as follows: Permian (298.9–251.902 Ma), Triassic (251.902–201.4 Ma), Jurassic (201.4–143.1 Ma), Lower Cretaceous (143.1–100.5 Ma), Upper Cretaceous (100.5–66 Ma), and Cenozoic (66 Ma to the present).

| Parameters | | All | | Before Upper Cretaceous | |
|---|---|---|---|---|---|
| | | Median | 95% HPD | Median | 95% HPD |
| Baseline rates | λ0 | 0.2857 | [0.0452, 0.7832] | 0.3527 | [0.0535, 1.0272] |
| | μ0 | 0.0613 | [4.6694E-4, 0.2764] | 0.085 | [7.8263E-4, 0.3845] |
| Correlation parameters to origination | Gλ0_0 | **-3.7517** | **[-4.8184, -2.6755]** | **-5.7505** | **[-7.991, -3.6154]** |
| | Gλ0_1 | **4.52E-03** | **[2.7848E-3, 6.2804E-3]** | **-1.07E-01** | **[-0.2166, -0.0251]** |
| | Gλ0_2 | **2.1573** | **[0.2173, 3.969]** | 1.737 | [-0.1109, 3.7548] |
| | Gλ0_3 | **-5.2402** | **[-9.306, -1.0398]** | 5.6843 | [-1.2333, 14.5587] |
| | Gλ0_4 | -0.0069 | [-0.0256, 6.0443E-3] | -0.0022 | [-0.0303, 0.0146] |
| | Gλ0_5 | -0.0472 | [-0.1007, 3.6143E-3] | -0.0366 | [-0.1001, 0.0102] |
| | Gλ0_6 | -0.0062 | [-1.0089, 0.9263] | -0.4266 | [-2.2221, 0.5973] |
| | Gλ0_7 | -0.0107 | [-0.0372, 7.245E-3] | -0.0016 | [-0.0322, 0.0194] |
| | Gλ0_8 | -0.079 | [-1.1262, 0.7466] | 0.0253 | [-1.584, 1.5807] |
| | Gλ0_9 | -0.0053 | [-0.0365, 0.0141] | -0.0017 | [-0.0359, 0.0244] |
| Correlation parameters to extinction | Gμ0_0 | -1.2083 | [-2.3996, 0.1247] | **-3.4656** | **[-5.7591, -0.6155]** |
| | Gμ0_1 | -0.0008 | [-0.0035, 7.5565E-4] | 4.32E-04 | [-0.0064, 0.0457] |
| | Gμ0_2 | **3.2037** | **[0.9185, 5.2962]** | 0.7431 | [-0.7608, 3.0226] |
| | Gμ0_3 | 0.1523 | [-4.0411, 5.2165] | 7.6919 | [-1.1308, 14.4984] |
| | Gμ0_4 | **-0.0578** | **[-0.0918, -0.0276]** | -0.0084 | [-0.0487, 0.0109] |
| | Gμ0_5 | 7.80E-03 | [-0.0244, 0.0552] | 2.73E-02 | [-0.0141, 0.0793] |
| | Gμ0_6 | -0.3554 | [-2.2506, 0.7065] | -0.5588 | [-2.722, 0.5673] |
| | Gμ0_7 | **0.035** | **[8.3993E-3, 0.0596]** | 9.41E-05 | [-0.0243, 0.0257] |
| | Gμ0_8 | 0.1448 | [-1.0839, 2.1692] | -0.224 | [-2.7706, 1.1981] |
| | Gμ0_9 | 9.47E-03 | [-0.0183, 0.0577] | 1.03E-02 | [-0.0186, 0.067] |
| Shrinkage weights (origination) | ωλ0_0 | **0.9234** | **[0.7083, 1]** | **0.9632** | **[0.8311, 1]** |
| | ωλ0_1 | **0.9157** | **[0.6664, 1]** | **0.9998** | **[0.998, 1]** |
| | ωλ0_2 | **0.6749** | **[0.1372, 0.9999]** | 0.6002 | [0.0447, 0.9998] |
| | ωλ0_3 | **0.6625** | **[0.1606, 1]** | 0.6924 | [0.0316, 1] |
| | ωλ0_4 | 0.4012 | [1.2118E-8, 0.943] | 0.3326 | [5.1822E-9, 0.943] |
| | ωλ0_5 | 0.5964 | [0.0466, 1] | 0.5145 | [6.3105E-9, 0.9589] |
| | ωλ0_6 | 0.2267 | [1.3626E-8, 0.9105] | 0.421 | [2.3078E-9, 0.9598] |
| | ωλ0_7 | 0.4897 | [6.6595E-7, 0.9614] | 0.3271 | [2.0763E-7, 0.9425] |
| | ωλ0_8 | 0.256 | [3.1926E-9, 0.9181] | 0.312 | [1.2944E-9, 0.94] |
| | ωλ0_9 | 0.302 | [2.7755E-9, 0.932] | 0.2843 | [1.8876E-8, 0.932] |
| Shrinkage weights (extinction) | ωμ0_0 | 0.654 | [0.0601, 0.9997] | **0.9068** | **[0.4762, 1]** |
| | ωμ0_1 | 0.5157 | [5.2507E-9, 0.968] | 0.7365 | [9.6589E-3, 1] |
| | ωμ0_2 | **0.7833** | **[0.3053, 1]** | 0.3829 | [5.2882E-9, 0.9478] |
| | ωμ0_3 | 0.2497 | [4.0632E-10, 0.9154] | 0.7623 | [0.0715, 1] |
| | ωμ0_4 | **0.9261** | **[0.6778, 1]** | 0.5145 | [1.0478E-7, 0.9708] |
| | ωμ0_5 | 0.2577 | [6.9467E-9, 0.9138] | 0.4447 | [3.3239E-8, 0.9535] |
| | ωμ0_6 | 0.4125 | [2.1346E-8, 0.9562] | 0.4839 | [1.4874E-7, 0.9643] |
| | ωμ0_7 | **0.8084** | **[0.3277, 1]** | 0.2999 | [1.6645E-11, 0.9327] |
| | ωμ0_8 | 0.3781 | [1.5812E-9, 0.9437] | 0.4247 | [1.9483E-8, 0.9593] |
| | ωμ0_9 | 0.439 | [7.3783E-8, 0.9589] | 0.4371 | [3.187E-9, 0.9628] |

**table S16.** Continued.

| Parameters | | Permian | | Triassic | |
|---|---|---|---|---|---|
| | | Median | 95% HPD | Median | 95% HPD |
| Baseline rates | λ0 | 0.2595 | [0.0137, 0.7349] | 0.2732 | [9.3176E-4, 1.0122] |
| | μ0 | 0.201 | [0.0131, 0.8207] | 0.2496 | [3.8735E-3, 0.9155] |
| Correlation parameters to origination | Gλ0_0 | -7.5168 | [-14.6342, 0.5776] | -2.1433 | [-8.7922, 1.1013] |
| | Gλ0_1 | 7.11E-07 | [-0.0088, 8.1645E-3] | 8.77E-06 | [-0.0156, 0.0186] |
| | Gλ0_2 | -0.0741 | [-5.0484, 2.7152] | 2.4688 | [-2.3351, 20.3104] |
| | Gλ0_3 | 0.0155 | [-15.9172, 21.2446] | -15.8271 | [-104.9632, 8.8675] |
| | Gλ0_4 | 2.03E-04 | [-0.0546, 0.087] | -5.20E-03 | [-0.0672, 0.0362] |
| | Gλ0_5 | 1.62E-03 | [-0.0786, 0.1152] | 1.16E-03 | [-0.1372, 0.1275] |
| | Gλ0_6 | -0.0115 | [-1.7446, 1.4196] | -1.7645 | [-7.1405, 1.5065] |
| | Gλ0_7 | 1.41E-04 | [-0.0736, 0.0864] | -4.00E-04 | [-0.0951, 0.097] |
| | Gλ0_8 | -0.0091 | [-2.3518, 1.8053] | -0.1364 | [-4.1691, 2.3313] |
| | Gλ0_9 | 6.25E-04 | [-0.0327, 0.0551] | -5.20E-03 | [-0.219, 0.0482] |
| Correlation parameters to extinction | Gμ0_0 | -0.0822 | [-6.0883, 2.6813] | 0.0329 | [-3.3815, 3.9451] |
| | Gμ0_1 | 9.62E-08 | [-0.0079, 8.0588E-3] | 0.00E+00 | [-0.0142, 0.0141] |
| | Gμ0_2 | -0.0048 | [-3.171, 3.2023] | -0.1385 | [-5.9999, 3.6742] |
| | Gμ0_3 | 0.0327 | [-15.5783, 19.0108] | -0.2455 | [-32.2288, 18.9404] |
| | Gμ0_4 | 0 | [-0.065, 0.0718] | -0.0068 | [-0.0838, 0.0314] |
| | Gμ0_5 | 6.22E-03 | [-0.0655, 0.2087] | 1.01E-01 | [-0.0256, 0.2454] |
| | Gμ0_6 | -0.1256 | [-3.122, 1.5901] | -4.367 | [-9.1886, 0.5173] |
| | Gμ0_7 | -1.00E-04 | [-0.0711, 0.0719] | -1.30E-03 | [-0.1117, 0.0818] |
| | Gμ0_8 | 0.0204 | [-1.9928, 3.5207] | -0.3908 | [-5.1151, 1.597] |
| | Gμ0_9 | -1.77E-02 | [-0.1008, 0.0174] | 4.46E-03 | [-0.0432, 0.094] |
| Shrinkage weights (origination) | ωλ0_0 | 0.9727 | [0.0317, 1] | 0.833 | [0.0188, 1] |
| | ωλ0_1 | 0.2627 | [4.5171E-10, 0.9869] | 0.5472 | [5.8432E-3, 1] |
| | ωλ0_2 | 0.2071 | [5.4295E-9, 0.9437] | 0.754 | [0.0113, 1] |
| | ωλ0_3 | 0.2376 | [3.4858E-8, 0.9734] | 0.9448 | [0.0198, 1] |
| | ωλ0_4 | 0.246 | [5.9059E-10, 0.9773] | 0.4479 | [9.5462E-10, 0.9681] |
| | ωλ0_5 | 0.1895 | [2.5397E-8, 0.9362] | 0.3664 | [2.697E-8, 0.9601] |
| | ωλ0_6 | 0.1486 | [4.775E-10, 0.9187] | 0.6532 | [2.0415E-8, 0.981] |
| | ωλ0_7 | 0.2352 | [2.1937E-9, 0.9746] | 0.4572 | [1.7042E-8, 0.9804] |
| | ωλ0_8 | 0.1782 | [3.047E-9, 0.9263] | 0.4445 | [3.0029E-8, 0.9695] |
| | ωλ0_9 | 0.1589 | [5.6918E-9, 0.9223] | 0.4649 | [1.7196E-8, 0.9875] |
| Shrinkage weights (extinction) | ωμ0_0 | 0.2723 | [4.1317E-10, 0.979] | 0.4467 | [5.2976E-8, 0.9751] |
| | ωμ0_1 | 0.257 | [6.2478E-9, 0.9856] | 0.5527 | [5.0841E-3, 1] |
| | ωμ0_2 | 0.1627 | [4.9575E-10, 0.91] | 0.3903 | [8.912E-11, 0.9704] |
| | ωμ0_3 | 0.2327 | [3.374E-9, 0.9699] | 0.5041 | [4.0076E-8, 0.9878] |
| | ωμ0_4 | 0.2308 | [2.517E-9, 0.9727] | 0.4931 | [2.693E-9, 0.9729] |
| | ωμ0_5 | 0.2718 | [3.2165E-8, 0.9658] | 0.7379 | [0.0274, 0.9998] |
| | ωμ0_6 | 0.2622 | [3.321E-9, 0.9514] | 0.8491 | [0.0751, 1] |
| | ωμ0_7 | 0.231 | [2.8031E-9, 0.9732] | 0.4731 | [5.8023E-9, 0.9814] |
| | ωμ0_8 | 0.2044 | [3.823E-9, 0.9502] | 0.5375 | [3.5347E-7, 0.9792] |
| | ωμ0_9 | 0.4738 | [7.6414E-9, 0.9742] | 0.3703 | [4.4639E-10, 0.9679] |

**table S16.** Continued.

| Parameters | | Jurassic | | Lower Cretaceous | |
|---|---|---|---|---|---|
| | | Median | 95% HPD | Median | 95% HPD |
| Baseline rates | λ0 | 0.4035 | [9.0622E-3, 1.3403] | 0.3852 | [7.8814E-3, 1.2443] |
| | μ0 | 0.0603 | [1.0619E-3, 0.4877] | 0.4769 | [0.0169, 1.4162] |
| Correlation parameters to origination | Gλ0_0 | -4.1371 | [-10.8327, 0.6824] | -0.3341 | [-8.146, 3.3127] |
| | Gλ0_1 | 0.00E+00 | [-0.0171, 0.014] | 0.00E+00 | [-0.0193, 0.0154] |
| | Gλ0_2 | 2.2662 | [-0.8986, 7.8083] | -0.1324 | [-9.86, 8.1986] |
| | Gλ0_3 | -0.7593 | [-38.855, 12.6175] | -1.8974 | [-46.5187, 19.3524] |
| | Gλ0_4 | 1.33E-03 | [-0.1037, 0.156] | -4.33E-02 | [-0.3158, 0.0926] |
| | Gλ0_5 | -4.74E-01 | [-1.0928, 0.0181] | 1.28E-01 | [-0.1988, 0.9899] |
| | Gλ0_6 | -3.4131 | [-13.9961, 2.7696] | -6.024 | [-20.7244, 3.5567] |
| | Gλ0_7 | 2.73E-04 | [-0.1498, 0.1757] | 1.08E-03 | [-0.0844, 0.098] |
| | Gλ0_8 | 0.3453 | [-2.6336, 5.6719] | 3.12E-03 | [-4.0911, 4.3746] |
| | Gλ0_9 | -1.96E-02 | [-0.177, 0.0594] | -1.84E-02 | [-0.2836, 0.0913] |
| Correlation parameters to extinction | Gμ0_0 | -0.5546 | [-7.6404, 1.6505] | -0.0743 | [-7.0396, 4.6789] |
| | Gμ0_1 | 1.21E-05 | [-0.0553, 0.0489] | 4.30E-05 | [-0.0189, 0.0209] |
| | Gμ0_2 | -0.7858 | [-5.9844, 1.2938] | 0.2986 | [-7.439, 17.2147] |
| | Gμ0_3 | 0.9507 | [-8.8126, 24.0449] | -0.649 | [-32.8312, 14.7471] |
| | Gμ0_4 | 2.05E-03 | [-0.0988, 0.1432] | -1.96E-01 | [-0.504, 0.0275] |
| | Gμ0_5 | -5.80E-03 | [-0.2607, 0.1893] | -2.62E-01 | [-0.8497, 0.1229] |
| | Gμ0_6 | -0.4791 | [-8.2208, 2.8206] | 0.6836 | [-4.7287, 11.939] |
| | Gμ0_7 | 3.14E-03 | [-0.127, 0.3151] | -2.40E-03 | [-0.0929, 0.0564] |
| | Gμ0_8 | 0.0608 | [-3.8867, 4.8311] | -0.0176 | [-3.5515, 3.8998] |
| | Gμ0_9 | -1.07E-02 | [-0.1138, 0.0436] | 1.84E-02 | [-0.0911, 0.2557] |
| Shrinkage weights (origination) | ωλ0_0 | 0.9252 | [0.0231, 1] | 0.6463 | [3.8998E-8, 0.9904] |
| | ωλ0_1 | 0.5289 | [1.1067E-7, 0.9958] | 0.6179 | [6.5225E-3, 1] |
| | ωλ0_2 | 0.6954 | [1.1642E-8, 0.9862] | 0.5352 | [4.7047E-8, 0.982] |
| | ωλ0_3 | 0.5365 | [1.0687E-9, 0.9908] | 0.6288 | [1.3953E-7, 0.9883] |
| | ωλ0_4 | 0.3962 | [1.9536E-8, 0.9705] | 0.7527 | [0.0116, 1] |
| | ωλ0_5 | 0.9108 | [0.1063, 1] | 0.702 | [0.0109, 1] |
| | ωλ0_6 | 0.7798 | [2.6238E-8, 0.9907] | 0.8673 | [0.0383, 1] |
| | ωλ0_7 | 0.4852 | [2.0171E-8, 0.9875] | 0.4508 | [7.873E-9, 0.9731] |
| | ωλ0_8 | 0.543 | [2.2026E-8, 0.9825] | 0.4887 | [2.3636E-8, 0.9768] |
| | ωλ0_9 | 0.6268 | [2.9175E-10, 0.9829] | 0.6653 | [8.3873E-8, 0.9898] |
| Shrinkage weights (extinction) | ωμ0_0 | 0.6089 | [1.266E-9, 0.9904] | 0.6436 | [1.2225E-10, 0.9885] |
| | ωμ0_1 | 0.5773 | [3.6682E-3, 1] | 0.6147 | [7.7432E-3, 1] |
| | ωμ0_2 | 0.4597 | [8.6317E-10, 0.9697] | 0.5726 | [1.2316E-7, 0.9902] |
| | ωμ0_3 | 0.4724 | [1.0277E-9, 0.9828] | 0.477 | [2.5261E-9, 0.9809] |
| | ωμ0_4 | 0.3778 | [7.1931E-9, 0.9706] | 0.945 | [0.1185, 1] |
| | ωμ0_5 | 0.2917 | [1.1012E-7, 0.9489] | 0.8056 | [0.0227, 1] |
| | ωμ0_6 | 0.4153 | [1.3416E-8, 0.9749] | 0.4979 | [1.6148E-8, 0.9814] |
| | ωμ0_7 | 0.5376 | [1.6796E-8, 0.994] | 0.3901 | [1.6773E-9, 0.9647] |
| | ωμ0_8 | 0.4509 | [1.8956E-8, 0.9803] | 0.4612 | [1.3903E-8, 0.9748] |
| | ωμ0_9 | 0.4415 | [1.601E-10, 0.9687] | 0.631 | [1.2929E-8, 0.9891] |

table S16. Continued.

| Parameters | | Upper Cretaceous | | Cenozoic | |
|---|---|---|---|---|---|
| | | Median | 95% HPD | Median | 95% HPD |
| Baseline rates | λ0 | 0.3582 | [6.9926E-3, 1.1714] | 0.1271 | [1.6426E-4, 0.461] |
| | μ0 | 0.3302 | [3.1494E-3, 1.152] | 0.0789 | [9.8232E-4, 0.532] |
| Correlation parameters to origination | Gλ0_0 | -0.1817 | [-20.5955, 3.9918] | 0.6007 | [-1.2479, 7.6488] |
| | Gλ0_1 | 4.36E-05 | [-0.0061, 8.1012E-3] | 0.00E+00 | [-0.0062, 4.1074E-3] |
| | Gλ0_2 | -2.7467 | [-22.6542, 4.3995] | 0.8899 | [-10.4815, 17.7944] |
| | Gλ0_3 | 0.1412 | [-14.1503, 21.5541] | **-25.2656** | **[-43.4148, -7.7383]** |
| | Gλ0_4 | 4.55E-04 | [-0.1484, 0.2101] | 8.98E-02 | [-0.004, 0.1966] |
| | Gλ0_5 | -2.99E-01 | [-2.8818, 0.2597] | -2.85E-01 | [-0.9981, 0.0881] |
| | Gλ0_6 | -0.3686 | [-34.8121, 22.4966] | -0.0291 | [-11.1883, 9.0887] |
| | Gλ0_7 | -4.00E-04 | [-0.195, 0.1823] | 3.83E-02 | [-0.0323, 0.1563] |
| | Gλ0_8 | -6.92E-02 | [-4.3525, 2.8404] | -1.44E-01 | [-2.427, 1.1656] |
| | Gλ0_9 | -6.70E-03 | [-0.1922, 0.0826] | 3.79E-03 | [-0.0929, 0.1343] |
| Correlation parameters to extinction | Gμ0_0 | -0.0756 | [-13.8077, 6.1278] | -0.5441 | [-5.6382, 1.2746] |
| | Gμ0_1 | -1.00E-04 | [-0.0128, 6.7749E-3] | -1.00E-03 | [-0.0085, 1.7588E-3] |
| | Gμ0_2 | 0.0823 | [-8.6964, 13.7341] | -0.3389 | [-17.2019, 10.729] |
| | Gμ0_3 | -0.7519 | [-30.6645, 15.6211] | -1.8179 | [-25.9472, 9.9758] |
| | Gμ0_4 | 7.89E-04 | [-0.1724, 0.3203] | -1.30E-03 | [-0.0784, 0.0609] |
| | Gμ0_5 | -1.80E-03 | [-0.6036, 0.4566] | -1.20E-02 | [-0.5811, 0.3232] |
| | Gμ0_6 | -0.9603 | [-56.1825, 22.9348] | -5.6631 | [-26.3322, 5.1184] |
| | Gμ0_7 | -6.32E-02 | [-0.522, 0.0891] | 3.36E-03 | [-0.084, 0.1463] |
| | Gμ0_8 | -0.0106 | [-3.748, 3.1793] | 8.54E-03 | [-2.3996, 2.7286] |
| | Gμ0_9 | 7.79E-04 | [-0.1109, 0.1455] | -4.30E-03 | [-0.167, 0.1025] |
| Shrinkage weights (origination) | ωλ0_0 | 0.4772 | [2.2921E-3, 1] | 0.5409 | [2.846E-9, 0.9896] |
| | ωλ0_1 | 0.3491 | [1.1723E-9, 0.981] | 0.2863 | [1.2637E-7, 0.9586] |
| | ωλ0_2 | 0.6002 | [2.9403E-8, 0.9873] | 0.3352 | [5.3804E-9, 0.9521] |
| | ωλ0_3 | 0.2711 | [9.8663E-9, 0.9609] | **0.9159** | **[0.6032, 1]** |
| | ωλ0_4 | 0.3121 | [2.2424E-9, 0.9767] | 0.8085 | [0.1564, 1] |
| | ωλ0_5 | 0.7783 | [6.1878E-3, 1] | 0.6507 | [7.7691E-8, 0.9797] |
| | ωλ0_6 | 0.3476 | [1.8517E-8, 0.9784] | 0.215 | [3.8707E-8, 0.9177] |
| | ωλ0_7 | 0.3013 | [4.8116E-10, 0.9693] | 0.3801 | [1.8624E-10, 0.9459] |
| | ωλ0_8 | 0.338 | [3.1024E-11, 0.9712] | 0.2306 | [2.0206E-8, 0.9249] |
| | ωλ0_9 | 0.4118 | [1.2342E-10, 0.9765] | 0.2672 | [4.7479E-8, 0.9355] |
| Shrinkage weights (extinction) | ωμ0_0 | 0.4254 | [1.0756E-8, 0.9943] | 0.5405 | [5.1984E-8, 0.9823] |
| | ωμ0_1 | 0.3832 | [3.3845E-9, 0.9905] | 0.5416 | [9.18E-10, 0.9792] |
| | ωμ0_2 | 0.3006 | [1.0285E-11, 0.9651] | 0.2673 | [2.2397E-8, 0.9391] |
| | ωμ0_3 | 0.3711 | [8.8379E-8, 0.9726] | 0.3987 | [1.6163E-7, 0.9682] |
| | ωμ0_4 | 0.3671 | [1.405E-8, 0.9866] | 0.2439 | [2.5803E-9, 0.9263] |
| | ωμ0_5 | 0.2727 | [5.19E-9, 0.9594] | 0.2721 | [2.8366E-10, 0.9366] |
| | ωμ0_6 | 0.4221 | [2.2143E-8, 0.9882] | 0.5594 | [3.0958E-7, 0.9753] |
| | ωμ0_7 | 0.6959 | [1.4446E-9, 0.9929] | 0.227 | [2.9803E-10, 0.9233] |
| | ωμ0_8 | 0.3079 | [8.6628E-10, 0.9642] | 0.2561 | [1.9847E-8, 0.9383] |
| | ωμ0_9 | 0.3081 | [2.8267E-10, 0.9698] | 0.2767 | [9.441E-8, 0.9472] |

**table S17.**

Estimates of extinction periods for Coleoptera and Polyphaga based on RJMCMC results. Using the results from RJMCMC analyses, we inferred major periods of extinction for both Coleoptera and Polyphaga. The frameworks considered are without singletons, and with or without amber occurrences. The table highlights the different most significant extinction intervals, along with their start and end. The number of extinct genera during the crisis was obtained from the *Te* of the corresponding RJMCMC analysis. The standing mean diversity at the start of the crisis was also obtained from the corresponding RJMCMC analysis. The start, end, and duration of the crises is in million years. The mean extinction rate at the top of the crisis, as well as the 95% HPD are in events/Million years/lineage. Abbreviation: HPD, Highest Posterior Density, HPD; Nb, Number.

| Clade | Amber | Geological time | Start | End | Duration of the crisis | Nb of extinct genera during the crisis | Standing diversity before the crisis |
|---|---|---|---|---|---|---|---|
| Coleoptera | no | Eocene–Oligocene | 32.670 | 31.570 | 1.101 | 5 | 470.6 |
| Coleoptera | yes | Eocene–Oligocene | 35.469 | 32.467 | 3.002 | 29 | 848.2 |
| Coleoptera | yes | lowermost-Albian | 113.610 | 110.808 | 2.801 | 48 | 144.9 |
| Coleoptera | no | lowermost-Albian | 113.721 | 110.920 | 2.802 | 27 | 137.1 |
| Coleoptera | no | Barremian–Aptian | 122.227 | 119.625 | 2.602 | 42 | 173.5 |
| Coleoptera | yes | Barremian–Aptian | 122.414 | 120.113 | 2.301 | 25 | 183.8 |
| Coleoptera | yes | Upper Jurassic | 154.431 | 151.630 | 2.801 | 62 | 141.2 |
| Coleoptera | no | Upper Jurassic | 155.247 | 151.645 | 3.602 | 65 | 146.3 |
| Coleoptera | yes | Ladinian–Carnian | 238.476 | 236.474 | 2.001 | 21 | 40.9 |
| Coleoptera | no | Ladinian–Carnian | 238.500 | 236.198 | 2.301 | 21 | 40.5 |
| Polyphaga | no | Eocene–Oligocene | 32.674 | 31.773 | 0.901 | 5 | 419 |
| Polyphaga | yes | Eocene–Oligocene | 35.277 | 32.575 | 2.702 | 24 | 766.8 |
| Polyphaga | no | lowermost-Albian | 114.835 | 111.032 | 3.803 | 11 | 83.1 |
| Polyphaga | yes | lowermost-Albian | 114.538 | 112.537 | 2.002 | 6 | 86.5 |
| Polyphaga | no | Barremian–Aptian | 122.841 | 120.639 | 2.202 | 7 | 97.8 |
| Polyphaga | yes | Barremian–Aptian | 122.144 | 119.942 | 2.202 | 23 | 106 |
| Polyphaga | yes | Upper Jurassic | 155.670 | 153.168 | 2.502 | 50 | 78.1 |
| Polyphaga | no | Upper Jurassic | 155.865 | 152.963 | 2.902 | 57 | 77.7 |

| Clade | Amber | Geological time | Percentage of extinction | Top of the crisis | Mean extinction rate at the top | Min 95% HPD | Max 95% HPD |
|---|---|---|---|---|---|---|---|
| Coleoptera | no | Eocene–Oligocene | 1.06% | 31.759 | 0.010 | 0.002 | 0.013 |
| Coleoptera | yes | Eocene–Oligocene | 3.42% | 33.256 | 0.039 | 0.012 | 0.058 |
| Coleoptera | yes | lowermost-Albian | 33.13% | 112.271 | 0.067 | 0.043 | 0.402 |
| Coleoptera | no | lowermost-Albian | 19.69% | 112.883 | 0.118 | 0.031 | 0.444 |
| Coleoptera | no | Barremian–Aptian | 24.21% | 121.385 | 0.070 | 0.034 | 0.171 |
| Coleoptera | yes | Barremian–Aptian | 13.60% | 121.673 | 0.068 | 0.046 | 0.179 |
| Coleoptera | yes | Upper Jurassic | 43.91% | 153.979 | 0.106 | 0.032 | 0.211 |
| Coleoptera | no | Upper Jurassic | 44.43% | 153.995 | 0.142 | 0.032 | 0.235 |

| | | | | | | | |
|---|---|---|---|---|---|---|---|
| Coleoptera | yes | Ladinian–Carnian | 51.34% | 236.895 | 0.280 | 0.057 | 0.426 |
| Coleoptera | no | Ladinian–Carnian | 51.85% | 237.119 | 0.302 | 0.143 | 0.513 |
| Polyphaga | no | Eocene–Oligocene | 1.19% | 32.261 | 0.010 | 0.002 | 0.013 |
| Polyphaga | yes | Eocene–Oligocene | 3.13% | 33.863 | 0.044 | 0.031 | 0.061 |
| Polyphaga | no | lowermost-Albian | 13.24% | 111.088 | 0.071 | 0.031 | 0.317 |
| Polyphaga | yes | lowermost-Albian | 6.94% | 113.093 | 0.065 | 0.037 | 0.434 |
| Polyphaga | no | Barremian–Aptian | 7.16% | 121.292 | 0.074 | 0.037 | 0.240 |
| Polyphaga | yes | Barremian–Aptian | 21.70% | 121.496 | 0.067 | 0.046 | 0.223 |
| Polyphaga | yes | Upper Jurassic | 64.02% | 153.708 | 0.227 | 0.065 | 0.475 |
| Polyphaga | no | Upper Jurassic | 73.36% | 154.103 | 0.258 | 0.114 | 0.450 |
